# Structural and evolutionary analyses support reclassification of glycopeptide antibiotics as xyclopeptides

**DOI:** 10.1101/2023.02.10.526856

**Authors:** Athina Gavriilidou, Noel Kubach, Martina Adamek, Jens-Peter Rodler, Susanna Kremer, Daniel H. Huson, Rosa Alduina, Gerard Wright, Mohammad Seyemsayadost, Wolfgang Wohlleben, Stefano Donadio, Margherita Sosio, Min Xu, Max J. Cryle, Evi Stegmann, Nadine Ziemert

## Abstract

Glycopeptide antibiotics (GPAs) are key agents against multidrug-resistant Gram-positive pathogens, yet both the term “glycopeptide” and the current GPA type I-V classification framework have become increasingly strained as structurally and mechanistically divergent members continue to be discovered. In particular, compounds historically grouped as “type V GPAs” differ from classical GPAs in features such as glycosylation, peptide length, and reported mode of action, raising the question of whether they belong to the same natural product class. Here, a curated dataset of GPA-associated biosynthetic gene clusters (BGCs) is analysed by combining fingerprint similarity of the products with phylogenetic analysis of the BGCs. Fingerprint-based structural similarity networks and BGC similarity comparisons reveal a pronounced separation between classical lipid II-binding GPAs (types I-IV) and type V GPAs. Multi-locus phylogenetic analyses of conserved biosynthetic components further support two deeply divergent evolutionary subclasses, consistent with subclass-specific biosynthetic signatures. Together, these results motivate a revised, unambiguous framework in which the broader class is termed xyclopeptides, comprising the subclasses dalabactins (legacy GPA types I–IV) and murobactins (legacy type V).

## Introduction

Glycopeptide antibiotics (GPA) have long been central to the treatment of infections caused by multidrug-resistant Gram-positive pathogens, exemplified by the discovery of vancomycin in 1953. Since then, dozens of natural products and numerous semi-synthetic derivatives have been described, several of which remain in clinical use. Accordingly, the biosynthesis and mechanisms of action of these compounds have been studied extensively over decades [1–9].

Historically, the term “glycopeptide antibiotic” has referred to a characteristic molecular architecture: a non-ribosomal peptide scaffold, typically built from a seven-amino acid backbone enriched in aromatic residues, which is conformationally constrained by oxidative crosslinks and frequently decorated by glycosylation and other tailoring reactions (e.g., halogenation, sulfation, methylation) (**Supplementary Table 1**) [1,10–13]. However, terminology around “glycopeptides” is inherently ambiguous. In broader chemical and biomedical usage, “glycopeptide” can denote any glycosylated peptide, including molecules that are biosynthetically and mechanistically unrelated to GPAs, and even structurally distinct natural products that simply carry sugar moieties. This overlap has long created confusion in the literature and across communities.

This confusion has been acknowledged before, especially for regulatory purposes in bringing new glycosylated peptides into clinical development. Accordingly, type I-IV GPAs were proposed to be named as “dalbaheptides” [14] back in 1989, a term which became the basis for naming dalbavancin a second-generation GPA [15]. However, the term dalbaheptides has not really caught on in the scientific literature. As genome mining and natural product discovery continue to expand the diversity of known glycosylated peptides, and some of them may in the future reach clinical development, the time is ripe for a revised nomenclature.

Indeed, recent discoveries have broadened the chemical and biological space of GPA-related crosslinked aromatic peptides and exposed limitations of extending the GPA label to increasingly divergent compounds. In particular, molecules previously grouped as “type V GPAs” [16,17] deviate from the classical GPA paradigm (**Figure 1**). They typically lack glycosylation, can feature variable backbone lengths (up to nine amino acids), and share distinctive structural motifs such as a tryptophane (Trp) residue crosslinked to a central 4-hydroxyphenylglycine (Hpg) [11,18]. Importantly, they also exhibit a different reported mode of action. Rather than sequestering lipid-II, they can act via inhibition of autolysins in peptidoglycan remodeling during cell division [19]. These differences raise a fundamental classification question: should these molecules be treated as an additional GPA type, or do they represent a distinct subclass within a broader class of related natural products?

**Figure 1:**
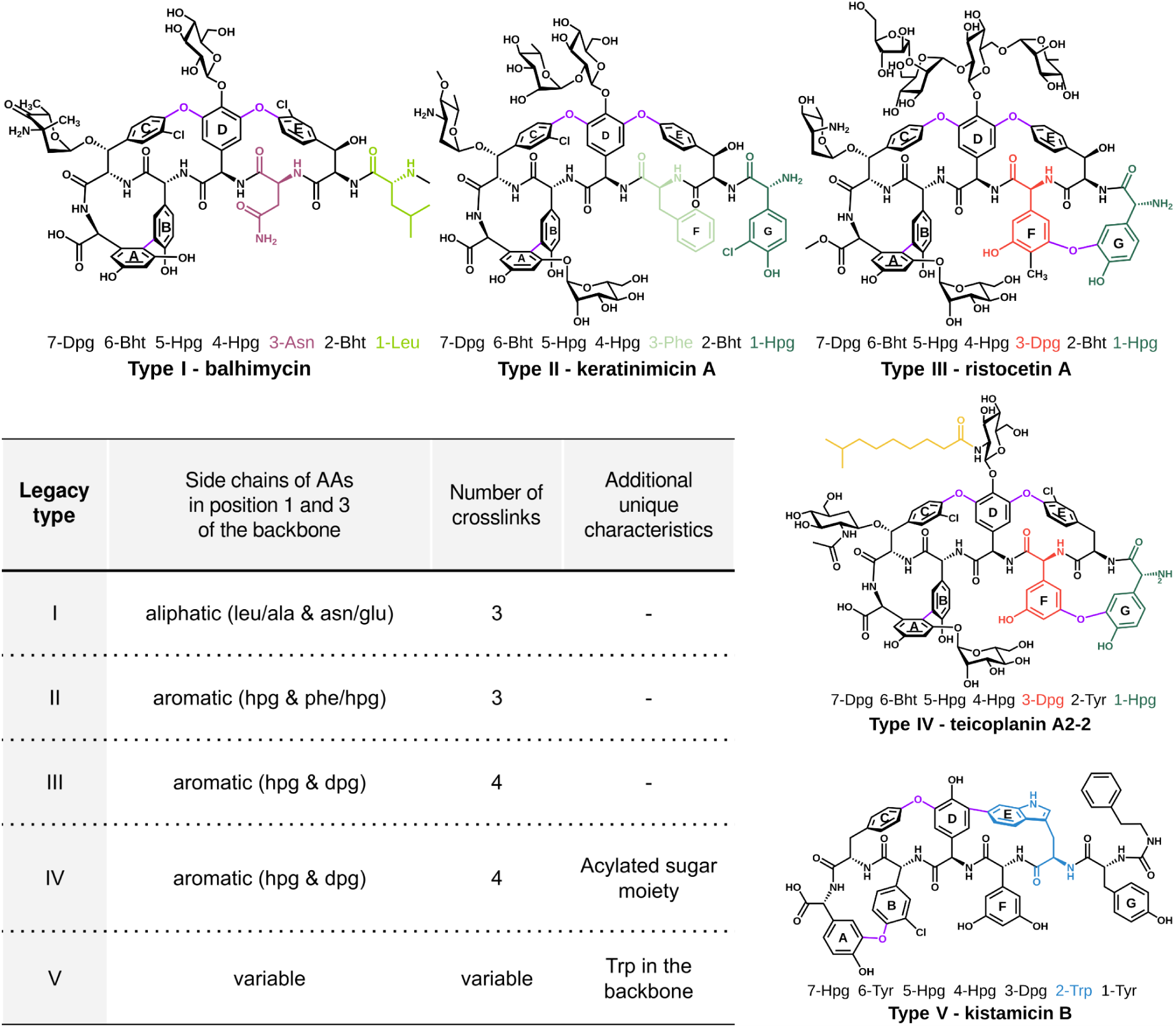
Structural characteristics of the legacy GPA types. The most important features for the classification of xyclopeptides are listed and accompanied by examples of structures. The most significant characteristics for each type are coloured. The aromatic rings of the AAs are labelled A-G based on prior publications [23]. Abbreviations: Tyr, tyrosine; Leu, leucine; Asn, asparagine; Bht, *β*-hydroxytyrosine; Hpg, 4-hydroxyphenylglycine; Dpg, 3,5-dihydroxyphenylglycine; Phe, phenylalanine; Trp, tryptophan. The SMILES of the structures can be found in **Supplementary Table 2**.

To address this and building on a community consensus emerging from recent comparative work, we propose a nomenclature and classification framework that separates family-level naming from historical clinical terminology. We use “xyclopeptides” as an umbrella term for the class of crosslinked, aromatic peptide natural products and subdivide it into two subclasses: (i) the classical lipid II–binding compounds previously classified as GPA types I–IV, which we refer to as “dalabactins”, and (ii) the subclass previously termed “type V GPAs” or glycopeptide-related peptides, which we refer to as “murobactins”. The class name is intended to reflect a defining combination of chemistry and biosynthesis: xyclopeptides share a characteristic conformationally constrained (“macrocyclic-like”) peptide core generated by oxidative crosslinking and a distinctive NRPS-associated X-domain [12,20–22], a hallmark of these pathways that mediates recruitment of cytochrome P450 enzymes responsible for installing the crosslinks. This terminology aims to be clear and unambiguous, reflect shared biosynthetic specificities (NRPS assembly and oxidative crosslinking), and avoid confusion with unrelated classes of glycosylated peptides. We use dalabactins for legacy glycopeptide types I–IV to reflect their canonical target interaction, binding the D-Ala–D-Ala terminus of lipid-linked peptidoglycan precursors. We use murobactins for legacy type V to reflect their direct interaction with murein (peptidoglycan), as exemplified by members that bind peptidoglycan and inhibit cell-wall remodelling by blocking autolysin access.

To evaluate and substantiate this proposed framework, we systematically integrate analyses across chemical, genomic, and evolutionary dimensions. Specifically, we combined structural analyses of known compounds with comparative analyses of GPA-associated biosynthetic gene clusters (BGCs) and multi-locus phylogenetic reconstructions of conserved biosynthetic components. By analysing concordance and discordance between chemical similarity, BGC architecture, and evolutionary relationships, we delineate the major types within the class, identify type-specific biosynthetic signatures, and provide a practical crosswalk between legacy GPA nomenclature and the proposed xyclopeptide terminology to support consistent usage and future discovery.

## Results

### Dataset curation of xyclopeptide-associated biosynthetic gene clusters

To compare chemical and evolutionary relationships across xyclopeptides we assembled a curated set of biosynthetic gene clusters (BGCs) associated with xyclopeptide biosynthesis. Candidate loci were first identified by automated BGC detection using antiSMASH [24] and grouped into gene cluster families to support consistent comparisons. We then manually refined BGC boundaries within each family by reviewing gene content and gene order and trimming unrelated flanking regions (see Methods: BGC dataset creation). This step was important because the antiSMASH boundary estimation can inflate apparent BGC-similarity [25] and complicate downstream analyses, especially phylogenetic reconstructions and gene-content comparisons.

After completeness checks and boundary refinement, the final dataset comprised 182 trimmed BGCs from at least nine bacterial genera (**Supplementary Table 3**). One outlier locus from *Streptomyces varsoviensis* could not be evaluated confidently from available assemblies; therefore, the strain was resequenced and the resulting assembly was added to the dataset.

Many BGCs in the dataset are not linked to a known compound. To place these BGCs in context, we predicted likely peptide backbones from NRPS adenylation (A) domain specificity codes. Where necessary, specificity assignments were refined by comparison to established signatures. These predicted backbones were used throughout the manuscript to describe BGC diversity and to support legacy type assignments (**Supplementary Table 4**).

We labelled BGCs according to the legacy GPA types I-V, based on similarity to reference BGCs and concordant features in gene content and predicted backbone composition. In this curated dataset, BGCs corresponding to legacy types I-IV (dalabactins) and type V (murobactins) already differ in multiple features, motivating the structural comparison, BGC similarity, and phylogenetic analyses presented below.

### Structure similarity analysis distinguishes dalabactins from murobactins

To assess how the known xyclopeptide structures relate to one another independently of biosynthetic gene content, we created a similarity network based on the pairwise structure similarities (**Figure 2**).

**Figure 2:**
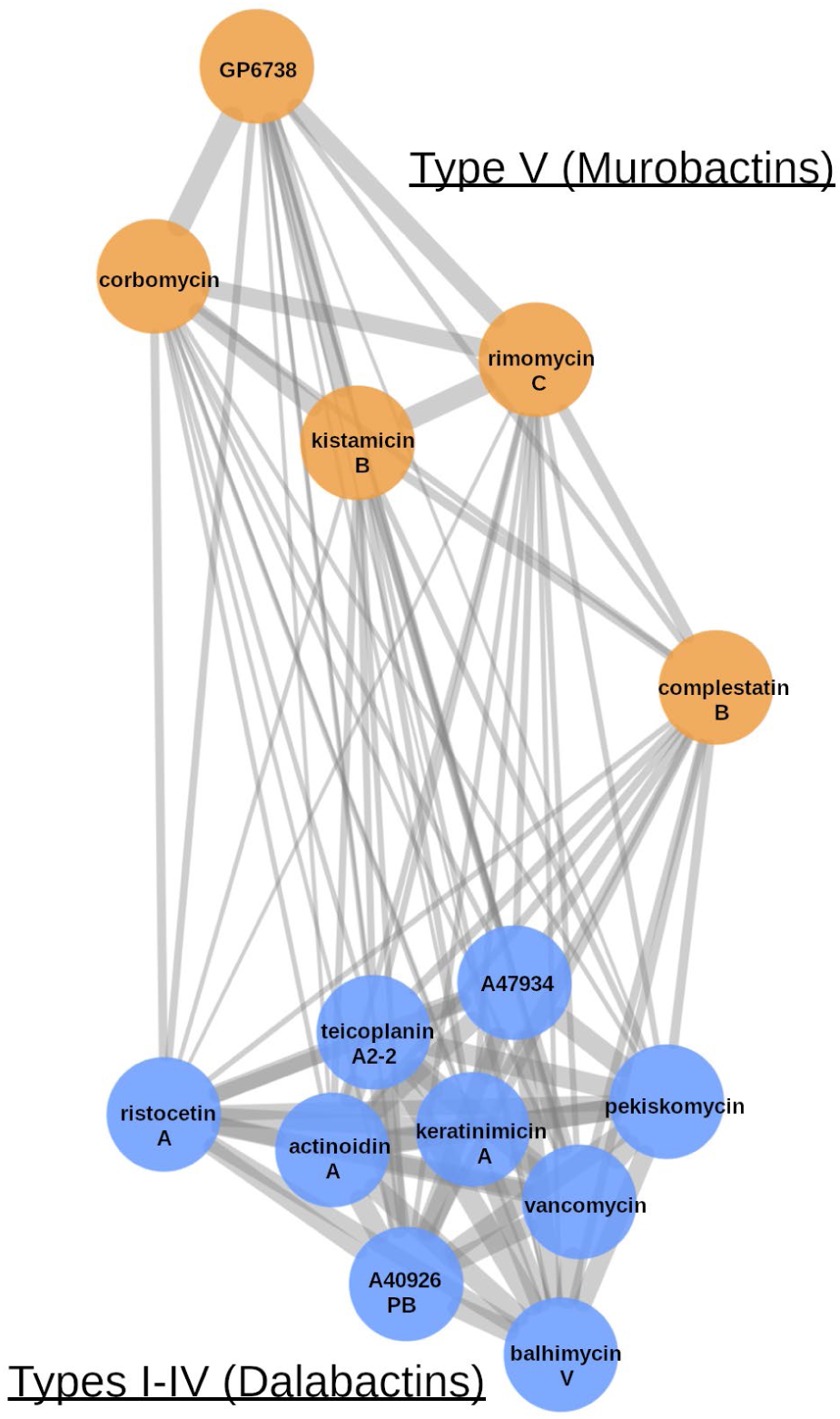
Structural and biosynthetic distance of murobactins. Tanimoto similarity network of the PubChem fingerprint of selected xyclopeptides structures visualized with the perfuse force-directed layout in Cytoscape [26] (see Methods, **Supplementary Table 2**). The murobactins are coloured orange, while the dalabactins are shown in blue. The width of the edges reflects the value of each similarity metric.

This analysis revealed a clear separation between compounds assigned to dalabactins and those assigned to murobactins. Dalabactin compounds formed a connected region in the network with similarity relationships that reflected the shared heptapeptide scaffolds and common glycosylation patterns. In contrast, murobactins clustered separately and showed limited similarity to the dalabactin cluster, consistent with the absence of glycosylation and variable peptide length. A wider spread of the murobactins across the similarity network suggests higher structural diversity.

Overall, the structure similarity network supports the view that murobactins are not simply a peripheral extension of the dalabactin structural continuum, but instead represent a distinct branch of glycopeptide-related structural diversity. This separation motivated subsequent analyses of BGC similarity and phylogeny to test whether genomic and evolutionary relationships recapitulate the same division.

### Phylogenetic analysis of full BGCs supports a reclassification

To test whether the structural separation between dalabactins and murobactins is mirrored by biosynthetic evolution, we carried out multi-locus phylogenetic analyses across homologous genes of our curated xyclopeptide BGCs. Because gene order is not consistently conserved across these BGCs, it cannot be used reliably for homology detection. Therefore, we first inferred fine-grained orthologous groups using zol [27] and refined ambiguous cases by phylogeny-guided curation (for example for closely related P450s; see methods section and **Supplementary Figure 1**). We then reconstructed maximum-likelihood phylogenies for individual genes based on their amino acid sequences.

Because NRPS genes are shaped by frequent domain and module rearrangements during evolution [28–33], we analysed NRPS genes at the domain level rather than as full-length genes. Phylogenies were built for each functional domain type and, where applicable, separately for each module position. This avoids comparing non-equivalent domains across modules and preserves the link between NRPS organisation and the predicted peptide backbone, including backbone length.

Gene and domain diversity was high across the dataset, and many loci were not shared broadly enough to be informative for tree building. We therefore focused on genes and domains present in at least half of the BGCs and summarised their phylogenetic signals in a super network [34] (**Figure 3**). Despite frequent incompatibilities among individual trees, a robust partition is evident: dalabactin BGCs occupy a compact region of the network, whereas murobactin BGCs span a broader phylogenetic space [35].

**Figure 3:**
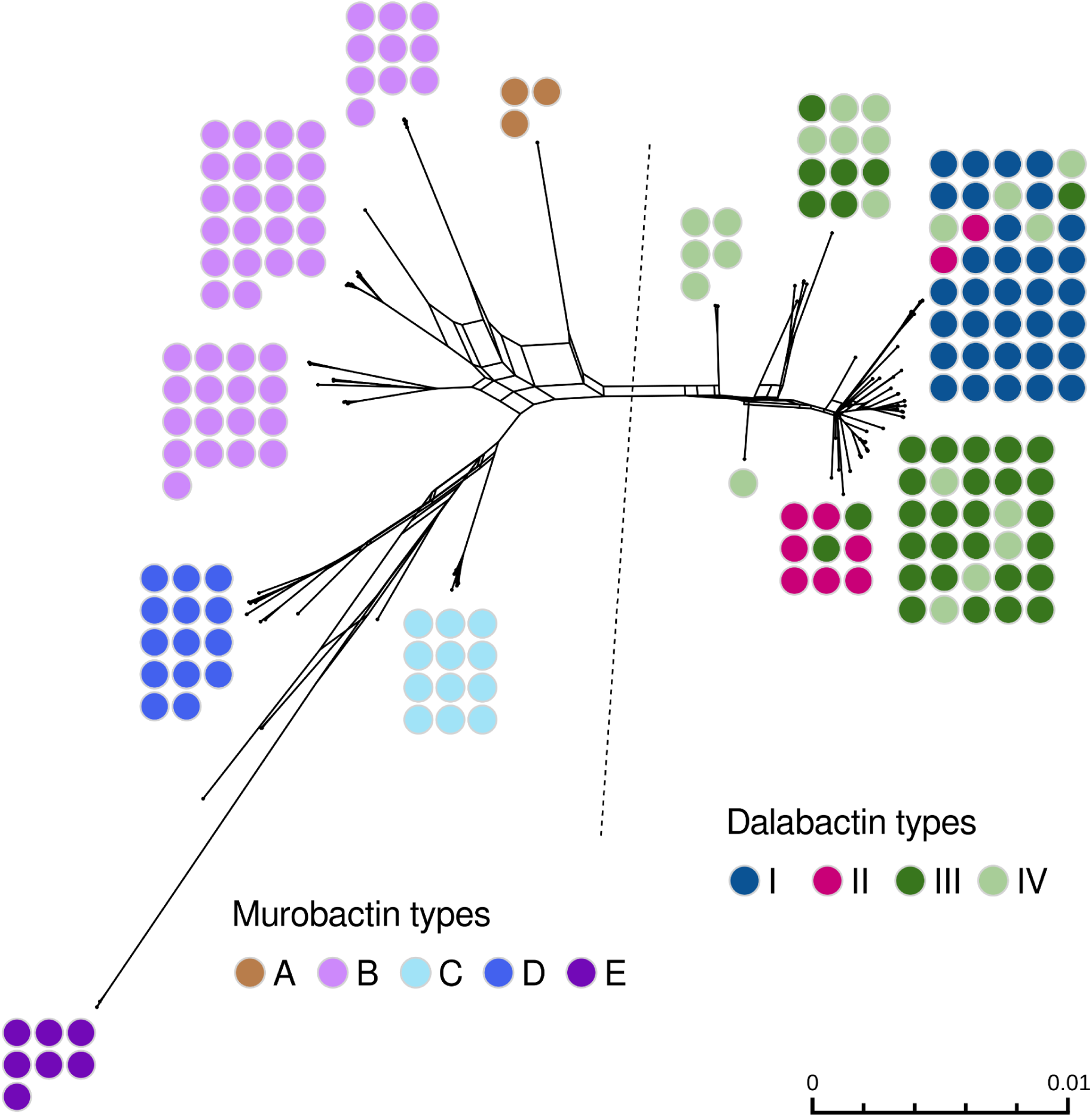
Phylogenetic network of xyclopeptide BGCs. Summarized super network constructed from maximum likelihood trees of all sufficiently present (>50%) genes and domains from xyclopeptide encoding BGCs (**Supplementary Table 3**) (seed=0), computed with the SplitsTree program [36], using “greedy weak compatibility” filtering to reduce visual complexity. Each node around the network corresponds to one BGC of a specific type, indicated by the colors. To simplify the visualization, close nodes are bundled in rectangular formations and a dotted line separates dalabactins and murobactins (detailed version in **Supplementary Data 1**). Most BGCs belong to one-type bundles. However, there are exceptions to this among the dalabactin BGCs, which can be explained in part by the multidimensional nature of the underlying data. Within dalabactins, some BGCs bundle closer to other types that are found in producers of the same species.

As an independent support, we also reconstructed a concatenated phylogeny using a subset of conserved genes and domains present in at least 90% of BGCs and passing a congruence filter (Methods). The concatenated tree is consistent with the super network, recovering the same deep split between dalabactins and murobactins and similar internal substructure (**Figure 4** and **Supplementary Figure 2**). We use this reference phylogeny to map gene presence and absence patterns and to highlight subclass- and type-associated modules described in the following sections.

**Figure 4:**
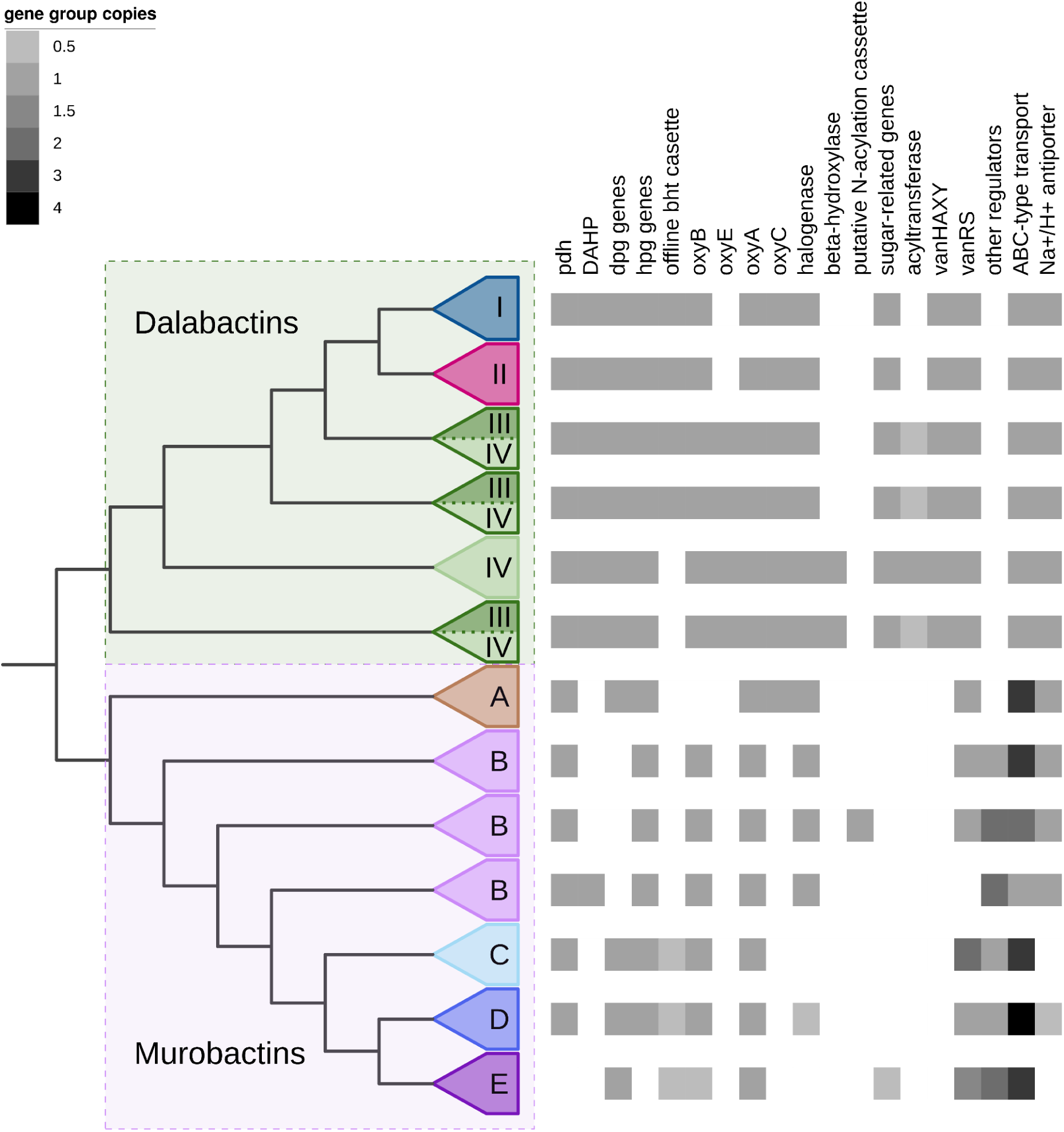
Phylogeny and gene patterns of xyclopeptide BGCs. Left: summarized concatenated phylogeny of xyclopeptide BGCs. Monophyletic clades were collapsed and branch lengths ignored. Colours correspond to dalabactin and murobactin types, which are labelled on the collapsed clades. Right: summarized gene presence/absense heatmap. Only genes or gene groups that form distinct patterns across xyclopeptide types are shown here. A detailed view of this figure can be seen in **Supplementary** Figure 2.

### Type-specific genomic signatures within dalabactins

To assess whether the established dalabactin subdivision into legacy types I–IV is supported by our expanded dataset, we compared multi-locus phylogeny, predicted backbone motifs, and gene-content patterns across curated xyclopeptide BGCs. Overall, dalabactin BGCs form a compact phylogenetic group, and legacy types are largely recovered as coherent subclades [37], with predicted backbone composition highly conserved within each type (**Figure 5**). In addition, dalabactin BGCs show the expected enrichment of glycosylation-related genes and canonical resistance genes whereas murobactin BGCs lack these genes, consistent with their established biosynthetic logic (**Supplementary Figure 2**, **Supplementary Table 1**, **Supplementary Table 5**). The main exception is the partial intermixing of legacy types III and IV, which is expected because their historical distinction is driven primarily by sugar acylation (typically a single acyltransferase) rather than broad pathway divergence. Given the overall concordance across phylogeny, backbone predictions, and key accessory modules, we retain the legacy dalabactin type I-IV classification as a practical and biologically meaningful framework.

**Figure 5:**
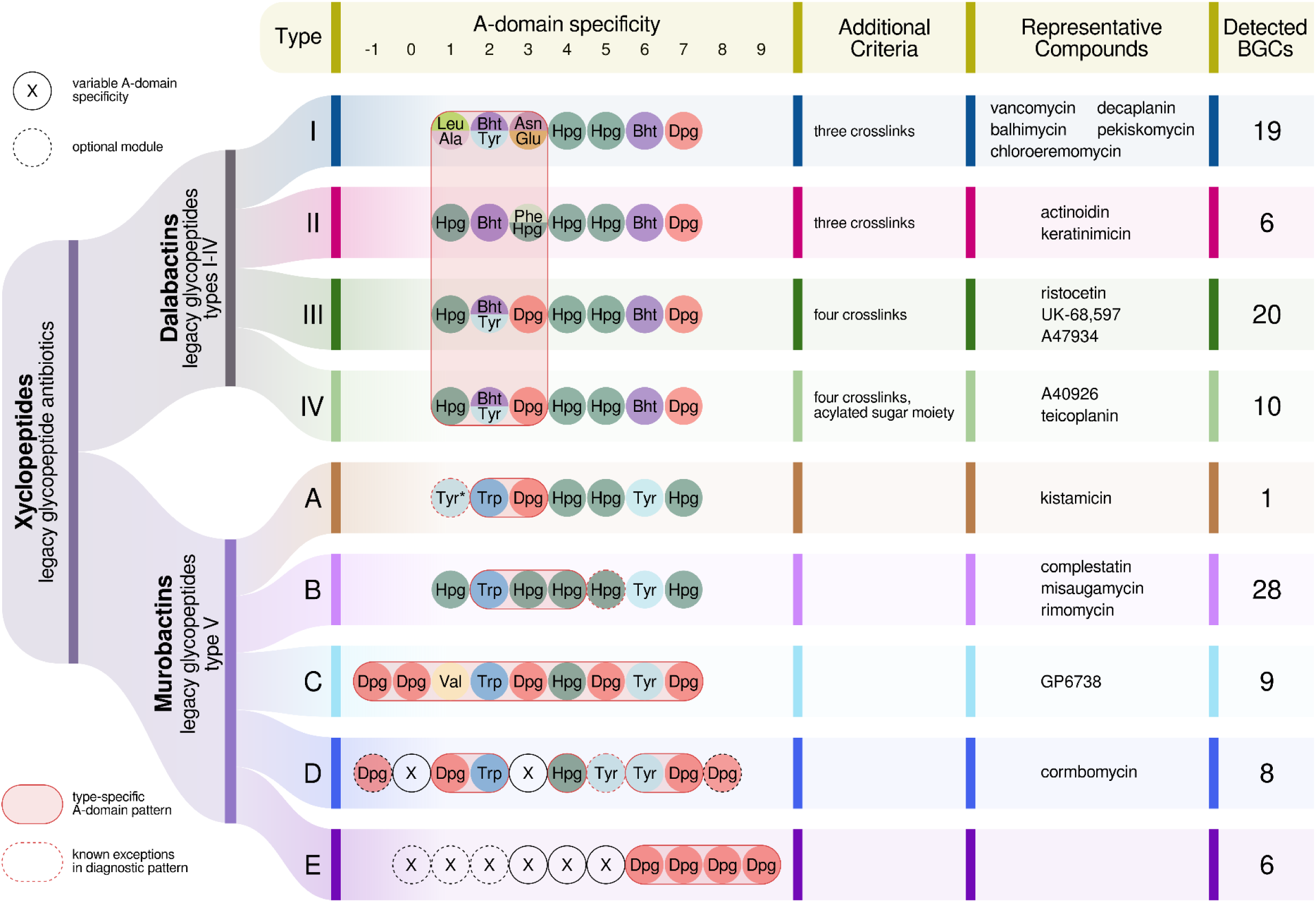
Overview showing the new classification of glycopeptide antibiotics (GPAs) - now xyclopeptides - into the D-Ala–D-Ala binding subclass dalabactins (legacy GPAs type I-IV) and the murein binding subclass murobactins (legacy GPAs type V). Types: new types within each subclass stay the same for types I-IV and further distinguish legacy type V. A-domain specificity: Main distinctive criterion for the type. Red outlines indicate type specific diagnostic patterns, red dashes mark known exceptions. Positions with an “X” mark variable A-domain specificities within one type and dashed black outlines represent optional modules. The incorporated amino acids are aligned in agreement with A-domain phylogeny and numbered according to the legacy seven amino acid GPA backbone. *The first tyrosine in the structure of kistamicin (type A) does not correspond to any NRPS module in the associated BGC. Additional criteria: criteria apart from the a-domain specificity that are important to tell apart the types. Detected BGCs: number of different (< 99% sequence identity) xyclopeptide BGCs found in public sequence data (total 107).

A small number of dalabactin BGCs deviates from the typical enrichment of sugar-associated modules, including type I BGCs most similar to pekiskomycin-like loci (StsY_A, StsAN), type II BGCs related to the recently discovered kineomicin [38] loci (AkAan_1, AkAan_2, UME), and type III BGCs related to the A47934 locus (StTo, StToN1) (**Supplementary Figure 2**). Notably, apart from these rare cases and the expected III/IV boundary blur driven by sugar acylation, the legacy dalabactin types show strong concordance between chemical classification (backbone motifs and crosslink patterns) and biosynthetic organisation, reinforcing their utility for comparative analyses. In contrast, murobactins (legacy type V) span a broader phylogenetic space and show greater variability in NRPS architecture and accessory gene content, motivating a separate fine-scale classification.

### Murobactins show distinct biosynthetic logic and form phylogeny-supported subclades

This divergence is also reflected in predicted backbone length and composition (**Figure 5**), supporting the separation of murobactins as a distinct subclass within xyclopeptides. In contrast to dalabactins, murobactin BGCs generally lack glycosylation-related genes and canonical resistance genes (**Supplementary Figure 2**), and instead show comparatively prominent regulatory and transport features. While tailoring capacity after backbone assembly appears more limited, several murobactin-specific genes remain unassigned. No murobactin containing Bht has been reported, although Bht-associated genes occur sporadically in some types (**Supplementary Table 1**). Murobactin NRPSs also display expanded and distinctive domain architectures, including additional E-domains and TIGR01720 domains and, in some cases, C-starter, N-methyltransferase, and terminal inactive A-domains. Finally, multi-locus phylogeny resolves reproducible subclades associated with characteristic backbone motifs (**Figure 5**), which we use as a practical fine-scale classification into murobactin types A-E.

### Murobactin type A

Type A comprises BGCs similar to kistamicin and is characterised by a Trp-Dpg motif at positions 2-3 (**Figure 5**). All identified type A BGCs encode seven NRPS modules; notably, the first module lacks an A-domain despite Tyr being present at position 1 in kistamicin. Type A most closely resembles dalabactins in overall gene content and forms an outgroup to the remaining murobactins. It uniquely includes all genes for Dpg biosynthesis and a dalabactin-like halogenase gene (**Supplementary Table 1**). Type A also encodes a distinctive set of P450 monooxygenases (Oxy) that lacks OxyB but includes a unique OxyC reported to catalyse two non-sequential crosslinks [5].

### Murobactin type B

Type B murobactins also encode heptapeptide backbones but differ from type A in the absence of Dpg-specific A-domains. Their characteristic motif is Trp–Hpg–Hpg–(Hpg) at positions 2-5 (**Figure 5**), with one small exception predicted to contain Dpg in place of Hpg at position 5. Surprisingly, genes related to the precursor supply of Dpg are absent even in the two BGCs that are predicted to include Dpg in their backbone. These two BGCs are unusually small in length and lack many other characteristic genes. One structure has been recently associated with these compounds, varsomycin, where a 3,5-dihydroxyphenylalanine is present at position 5 [39]. Type B includes complestatin-, rimomycin-, and misaugamycin-associated subclades that are distinguishable phylogenetically (**Figure 4**). Regulatory repertoires vary among these subclades, and some rimomycin- and complestatin-like BGCs encode N-methyltransferase domains in module 6. Misaugamycin-like BGCs can also include a four-gene cassette proposed to support N-acylation.

### Murobactin type C

Type C BGCs, represented by GP6738, are highly uniform, forming a clearly supported monophyletic group across phylogenetic representations (**Figures 3, 4**) and exhibiting consistent gene-content patterns (**Figure 4, Supplementary Figure 2**). Predicted backbones comprise nine residues and include a Val residue at position 3 followed by the conserved Trp motif typical of murobactins (**Figure 5**). Type C BGCs encode precursor supply genes for Dpg and Hpg and include an alpha/beta hydrolase associated with offline Bht hydroxylation, but lack the remaining genes required for Bht biosynthesis; accordingly, Bht-specific A-domain signatures are not detected and the known representative structure does not contain Bht. Type C BGCs encode only two *Oxy* genes, consistent with fewer crosslinks, and are unique among murobactins in encoding two copies of the VanR/VanS two-component system.

### Murobactin type D

Type D encompasses backbones ranging from eight to ten residues and includes corbomycin-like BGCs (**Figure 5**). Although more variable than types A-C, type D displays recurring motifs (**Supplementary Figure 3**), including a conserved Dpg-Trp motif at aligned positions 1-2 and a conserved Hpg at aligned position 4, which is shared across types A-D and is likely linked to the conserved Trp with a crosslink. A Tyr-Dpg motif at aligned positions 6-7 is also common. With respect to precursor supply, type D resembles type C, but it shows additional conserved regulatory and transport features, including a LysR-like regulator and two ABC transporter classes. A halogenase occurs in a subclade but is not universally conserved. Several recurrent genes remain unassigned and may encode type-specific functions.

### Murobactin type E

Type E is the most heterogeneous murobactin group and currently lacks associated structures. Predicted backbones range from seven to ten residues and are characterised by a Dpg-rich tetrapeptide tail (**Supplementary Figure 4**), while typically lacking Trp and Hpg. Type E BGCs consistently lack shikimate-pathway genes (**Supplementary Table 1**) and show substantial variation in NRPS domain organisation, mirroring backbone diversity. It is the only type to display two distinct oxy-gene patterns. A small subclade uniquely encodes a mannosyltransferase, and one “minimal” BGC resembles the compact architectures noted in type B, lacking several transport and regulatory genes common in other xyclopeptide clusters. Given the absence of experimentally characterised structures and limited functional data, type E should be considered provisional and is likely to be refined as additional murobactins are discovered and characterised.

## Discussion

The historical type I-V classification for glycopeptide antibiotics was built around chemical features and was invaluable for organising an emerging field. However, accumulating evidence shows that legacy type V BGCs and their corresponding compounds are consistently separated from legacy types I–IV in terms of structure, biosynthetic organization, and mechanism of action that targets peptidoglycan remodelling rather than binding lipid II. This has led to recurring confusion in the literature, including the fact that many legacy type V molecules are not glycosylated, and therefore do not fit the term “glycopeptide antibiotic” in a strict sense. Furthermore, the interpretation of glycopeptides as any glycosylated peptide does not reflect the 7-10 AA backbone with characteristic crosslinks and antibiotic activity. Against this backdrop, a nomenclature update is not cosmetic, it is necessary to align names with biology and to prevent continued overloading of “type V” as a catch-all bin for newly discovered outliers.

We therefore propose xyclopeptides as an umbrella framework that preserves relatedness while making the major split explicit. Within xyclopeptides, dalabactins correspond to legacy GPA types I-IV, and murobactins correspond to legacy type V. These names were chosen deliberately to be (i) distinctive and unambiguous, (ii) expandable as additional chemistry and BGC diversity is discovered, and (iii) compatible with legacy usage by retaining type labels where they remain informative. Importantly, the framework is intended to be practical for comparative genomics and discovery, it groups what is genuinely related while separating subclasses that differ in target biology and biosynthetic logic.

There have been other attempts to reclassify type I-IV GPAs in the past, such as the suggested “dalbaheptides”[14], which was the basis for naming the second-generation GPA dalbavancin. However, the term dalbaheptides has not really caught on in the scientific literature.

A central goal of this study was to test whether this nomenclature is supported beyond intuition from known structures and mechanisms. Across multiple, independent analyses, the dalabactin–murobactin split is consistently recovered from BGC evolution, and murobactins span a broader phylogenetic space than dalabactins, mirroring their broader variability in NRPS architecture and predicted backbone patterns. These evolutionary results align with chemical and functional distinctions reported for representative murobactins such as complestatin, corbomycin, and GP6738, reinforcing that the proposed names capture real biological structure rather than arbitrary clustering. At finer scale, the reproducible subclades within murobactins provide a pragmatic basis for the provisional A-E subdivision, which can be refined and expanded as additional structures and modes of action are characterised.

At the same time, we emphasise limitations. BGC boundary definition and orthology inference across diverse BGCs remain non-trivial, and reticulate evolution, including horizontal transfer [13,31,40], complicates any single “true” tree. Our approach mitigates this by combining locus-wise phylogenies, network representations that tolerate phylogenetic incongruence, and a conservative concatenation strategy. But ambiguous cases remain, particularly for murobactin type E, where no experimentally linked structures are currently available. The A-E scheme should therefore be viewed as a working classification that is designed to be updated, not a final taxonomy.

Finally, we want to stress that durable nomenclature changes require broad consensus. In developing this framework, we benefited from extensive discussions among researchers studying chemistry, biosynthesis, genomics, and mechanism of action. This collaborative approach matters because naming systems succeed when they encode shared agreement and remain straightforward to apply, including for non-specialists. A useful model comes from the RiPP field, where community-driven nomenclature and classification helped stabilise terminology across a rapidly expanding chemical space [41]. We hope xyclopeptides can play a similar role for dalabactins and murobactins by reducing ambiguity, strengthening links between chemical and biosynthetic classes, and providing a scalable structure for future discoveries. More broadly, we hope this process can serve as a template for future nomenclature efforts in other natural product families, and as an example of how collaborative, cross-disciplinary discussion can improve the long-term clarity and usability of scientific language.

## Methods

### BGC dataset creation

For the evolutionary analysis of xyclopeptide BGCs to be as complete as possible, we aimed to create a dataset of all sequenced clusters. The initial dataset contained all known clusters found in the literature [9,11,19,42–62] and was extended by searching in publicly available sequence databases (MIBiG [63], most of NCBI RefSeq and Genbank [64–66], JGI IMG DB [67], MGnify [68]) and some published projects [69–72]. Information on the origin of the BGCs, the date of the search and more metadata are available in **Supplementary Table 3**. **Supplementary Data 4** includes the accession numbers of all NCBI (RefSeq and Genbank) entries checked and whether they were a hit (True), not a hit (False) and the ones where the search failed due to lack of protein sequences associated with the accession number (Failed). The target of the search, accommodated via HMMER (v. 3.3.1, accessed from http://hmmer.org/, RRID:SCR_005305), was the X-domain in the last NRPS module (its HMM was extracted from antiSMASH v6 [73], RRID:SCR_022060), which has so far only been found in xyclopeptide encoding BGCs [72,73]. However, the X-domain has evolved from a condensation domain (C domain) and their sequences remain quite similar [6]. To avoid false positives, the known clusters were searched with the X-domain HMM and the lowest local score of the confirmed X-domains (300) was used as a minimal threshold. Lower local scores (up to 250) were manually checked for the MIBiG dataset and indeed no X-domains were found below the chosen threshold. The custom scripts used for this and all other parts of the analysis can be found in **Supplementary Data 5**.

The sequences that were a hit in the search were used as input for an antiSMASH v7 [24] (RRID:SCR_022060) analysis (default parameters + MIBiG cluster comparison) to detect the full BGC. At this stage, after manual inspection, some candidates were dropped due to low quality assemblies which led to obviously incomplete clusters, and due to a few false positives (where a C domain was mistakenly detected as an X-domain). The cluster regions that passed this inspection were assigned an ID reflecting their taxonomic placement (**Supplementary Table 3**) and their coding domains were re-annotated with bakta (v1.8.1) [74] to ensure homogeneity. Bakta was run with default parameters + skipping trna and tmrna detection (--skip-trna --skip-tmrna) and on metagenome mode when appropriate. The original contig headers were kept (--keep-contig-headers) but the assigned ID was used as a locus and locus tag prefix. The bakta-annotated regions were re-analyzed with antiSMASH v7 [24] with all features on (--fullhmmer –clusterhmmer –tigrfam –asf –cc-mibig –cb-general –cb-subclusters –cb-knownclusters –pfam2go –rre –smcog-trees –tfbs).

One exception to this process was done for a very unusual BGC from *Streptomyces varsoviensis*, which could not be conclusively labelled complete. Due to its interesting characteristics though, the genome of the strain was resequenced. For isolation of high molecular weight genomic DNA *S. varsoviensis* was cultured in R5 medium [75] for 2 days on a rotary shaker at 28 °C. DNA isolation was performed by using DNA isolation kit (NucleoBond® HMW DNA Kit, Machery-Nagel, Düren, Germany) following the manufacturer’s protocol. Genome sequencing was performed by the NGS competence center (NCC in Tübingen, Germany) on a PromethION (Nanopore) with 9.4.1 chemistry (details in **Supplementary Data 6**). Assembly was done with Unicycler [76] (for long reads) and corrected with medaka (accessed from https://github.com/nanoporetech/medaka) based on the sequencing parameters (**Supplementary Data 6**). The resulting genome sequence was searched for the X-domain as described above and BGCs were included in the analysis (**Supplementary Table 3**). The sequence was also submitted to NCBI and was assigned the accession number CP162614.

To overcome antiSMASH’s generous selection of BGC boundaries, a manual inspection was necessary, to ensure the quality of the evolutionary reconstruction, which would have been affected by the accidental inclusion of unrelated sequences. The process was sped up by annotating the BGCs in groups of GCFs as defined by a BiG-SCAPE analysis [77] (v1.0.1 2020-01-27, RRID:SCR_022561). The chosen threshold of 0.2 defined the largest possible groups that never have more than one type xyclopeptide (based on the MIBiG dataset). BiG-SCAPE was run on glocal mode, mixing all classes (as some BGCs are marked NRPS and others are marked other), but using the score weights of the NRPS class. The GCFs were then visualized with clinker [78] (v0.0.28). We meticulously manually curated each cluster by investigating every single gene for its function and possible role in the biosynthesis and the clinker visualization ensured uniform trimming of the most closely related BGCs (**Supplementary Figure 5**). Thus, the final dataset of 182 trimmed clusters from 9 different genera was created (**Supplementary Table 3**).

To put a number on how many BGCs per type we have in our dataset (**Figure 5**), we clustered the 182 trimmed BGCs on DNA level with MMseqs2 (version 18.8cc5c, RRID:SCR_022962) in the easy-cluster-mode with a minimal sequence identity and coverage of 99% (--min-seq-id 0.99 -c 0.99 --cov-mode 0), which resulted in 107 BGCs.

### Chemical structure similarity network

The structures of the glycopeptides (**Figure 1**, **Figure 2**) were collected from the original publications [11,17,19,42,43,50,51,53,79–88] (entries under database: MIBiG in **Supplementary Table 3**), formatted and converted to SMILES with ChemDraw (https://revvitysignals.com/products/research/chemdraw, RRID:SCR_016768) and RDKit (v. 2020.09.1, http://www.rdkit.org/, RRID:SCR_014274).

The examination of the structural variation within and among xyclopeptide types was done by use of the Tanimoto similarity metric [89]. For the PubChem fingerprint, the SMILES of known GPA compounds were inserted into the Cytoscape program (v. 3.9.1, RRID:SCR_003032) [26], which accommodates the calculation and visualisation of a structure similarity network (**Figure 2**) based on Tanimoto with *chemViz2* (fingerprint: Pubchem), a cheminformatics app for Cytoscape. The RDKit, Morgan [90] and MACCS 166 [91] fingerprints were calculated using RDKit with the default settings for the RDKit and MACCS fingerprint and a radius of two for the Morgan fingerprint. The Tanimoto similarity values were imported in Cytoscape and visualised in **Supplementary Figure 7**. The nodes of the resulting network were annotated by legacy GPA type (I-IV or V) and the edge width was adjusted to represent the degree of similarity among the connecting parts.

### A-domain specificity code analysis

A-domain specificity was predicted (**Supplementary Table 4**) using the 34 AA long specificity-conferring code determined by NRPSpredictor2, implemented in antiSMASH (v. 5.1.2, RRID:SCR_022060) [92–94]. Due to the fact that certain A-domain specificities could not be predicted *in silico* [42], all the known amino acid specificities (for glycopeptides whose structure is known) were used in a blastp (v2.14.0+) search [95] and best scoring alignments were used to determine the final annotations, *i.e.*, the ones used for the rest of the analysis, which sometimes differed from the antiSMASH assigned ones. A summary of A domain specificity was visualised in **Figure 5**, while the predicted backbone of all BGCs can be seen in **Supplementary Figure 2.**

### Detection of homologous genes

The identification of homologous groups of genes from the full dataset (**Supplementary Table 5**) was mostly carried out by the zol tool (v 1.3.9) [27], which infers phylogenetic orthology for comparative genomics of gene clusters. The platform (run with default parameters) generated 309 so-called orthogroups (OGs). There was one case of related genes with distinct functions being placed in the same Orthogroup, namely the oxyAs and oxyEs, which are known to be closely related [5,43]. Those proteins were used for a multiple sequence alignment (MSA) and their separation into groups was guided by their phylogenetic placement compared to proteins of known function. Cases of OGs with only a few clusters containing multiple copies of a gene were dealt with via a custom script, discarding some copies based on similarity criteria [96]. Finally, the NRPS domains were extracted from the genes and split into groups based on the position of their module compared to the rest.

The latter was achieved by building an MSA from the concatenated 34 AA A-domain specificity codes (as provided in the antiSMASH results [24,97] with the help of the NRPSpredictor2 tool [92]) of the underlying A-domains (**Supplementary Data 3**). The longest conserved region (positions 4-7 of balhimycin) was taken as a center and all internal gaps were rejected, since the positions directly correspond to the backbone of the compounds. The alignment was in agreement with the fact that positions 1-3 of the types I-IV glycopeptides are known to be the most variable among known compounds and their corresponding BGCs [37]. All module positions (00, 0, 1-9) were annotated based on this fixed MSA, with the balhimycin encoding BGC as a template for positions 1-7.

### Visualising complete evolutionary history

All multiple sequence alignments (MSAs) were performed with the mafft tool (v7.490, 2021/Oct/30, RRID:SCR_011811) with default parameters [98] (**Supplementary Data 7**). Phylogenetic trees for every occasion in this study (153 gene and domain trees, as well as concatenated phylogenies described below) were built with iqtree (multicore v2.2.0.3 COVID-edition for Linux 64-bit built Aug 2 2022) [99,100] (**Supplementary Data 7**). For the gene and domain trees, iqtree was first run only in model testing mode, checking for all bacteria-suitable evolutionary models (suitable models are listed in **Supplementary table 6**) and then in tree-building mode based on the best fitting model. The resulting trees can be seen in **Supplementary Data 1**.

A graphical summary of the adequately populated (n>50%) 47 separate (partial) gene and domain trees was calculated and visualised by the super network algorithm [101] (default options) implemented in the SplitsTree CE tool (version 6.0.10-beta) [36,102,103] (**Figure 3**) after greedily selecting a weakly compatible set of splits of maximum support (GreedyWeaklyCompatible splits filter). The super network (**Supplementary Data 1**) summarises the set of input trees, taking into account that many of the trees are incomplete. It is a splits network in which each band of parallel edges represents one of the splits or branches found in the set of input trees. Incompatibilities among the input trees give rise to parallelograms in the network. Edges in the network are scaled to represent the average relative length of the corresponding edges in the input trees. The set of splits computed by the super network method was greedily filtered by decreasing support (number of trees that contain a given split), so as to obtain a subset of “weakly compatible splits” that maintains major incompatibilities, while avoiding higher-dimensional edge configurations in the network, and thereby visual clutter.

### Concatenated phylogeny

A species phylogeny can be constructed from concatenated sequences of core genes, as long as their separate trees are congruent [104]. Following this concept, 22 genes or domains found in at least 90% of the BGCs in the dataset were identified. Duplications were considered acceptable up to a max of 5% (i.e. up to 5% of the clusters had more than one gene in the orthologous group) but otherwise, these groups were single-copy genes. These two thresholds were chosen based on a wide-scale phylogenetic study [104]. However, it was necessary to ensure the congruence of the participating genes before concatenating them. The underlying MSAs were first trimmed with trimAl (v1.4.rev15 build[2013-12-17], RRID:SCR_017334) [105] with default parameters and then the trees were recalculated (**Supplementary Data 7**). The latter were used in a congruence analysis as performed by Parks and colleagues [104]: The well supported splits were calculated by checking their existence in random subsampled concatenated phylogenies and then for each gene tree, their presence was used to calculate ‘normalised compatible split length’ ,a metric that reflects congruence of this tree to the rest in the group (**Supplementary Table 7**). This value was computed with a python script, implemented with Biopython [106] (RRID:SCR_007173), FastTree v2.1.11 [107] (RRID:SCR_015501) and a function of the (still in development) GeneTreeTk toolbox (accessed from https://github.com/dparks1134/GeneTreeTk). Eleven genes or domains passed a specific threshold (0.67) and were used for the concatenated phylogeny representing the evolutionary history of the xyclopeptide BGCs (**Supplementary Data 2**). The tree was then rooted with the MAD algorithm [108]. This tree, together with an absence/presence heatmap of the various genes and domains present in the clusters was visualised with iTOL [109,110] (RRID:SCR_018174) (**Supplementary Figure 2**). The tree in **Figure 4** was rooted between the dalabactins and murobactins. This alternative rooting is quite plausible due to the topology of the super network (**Figure 3**), supported also by the long branch leading to type A murobactins in the concatenated phylogeny (**Supplementary Figure 2**).

## Supplementary Material

<u>Supplementary Table 1:</u> Most common gene categories found in dalabactin-synthesis (legacy types I-IV) encoding BGCs. The genes and their known functions are summarised in this table, organised into categories and subcategories based on their role. An example gene or domain is provided in the corresponding columns, as well as references.

<u>Supplementary Table 2:</u> SMILES of selected known xyclopeptide structures. Also, pairwise tanimoto similarity values.

<u>Supplementary Table 3:</u> Dataset of xyclopeptide BGCs and related metadata.

<u>Supplementary Table 4:</u> Results of selectivity code analysis of xyclopeptide BGC A-domains.

<u>Supplementary Table 5:</u> Table of orthologous groups and gene presence/absence table from Supplementary Figure 2.

<u>Supplementary Table 6:</u> Table of ML models used for the phylogenetic trees of genes and domains.

<u>Supplementary Table 7:</u> Results of congruence analysis.

<u>Supplementary Data 1:</u> SplitsTree6 session file, containing all (154) calculated phylogenetic trees, as well as the super network of Figure 3.

<u>Supplementary Data 2:</u> Fasta file of the concatenated MSA and newick files (unrooted with bootstrap values and MAD rooted) containing the concatenated phylogeny of xyclopeptides.

<u>Supplementary Data 3:</u> A-domain selectivity code-based concatenated MSA.

<u>Supplementary Data 4:</u> Tables of accession numbers of NCBI RefSeq and Genbank entries included in the X-domain search and the result of the search.

<u>Supplementary Data 5:</u> Custom scripts used for the analysis.

<u>Supplementary Data 6:</u> Sequencing report of *S. varsoviensis*.

<u>Supplementary Data 7:</u> Fasta files of all gene and domain MSAs (and trimmed MSAs) and newick files of the phylogenetic trees they were used to build.

## Supplementary Figures

**Supplementary Figure 1:**
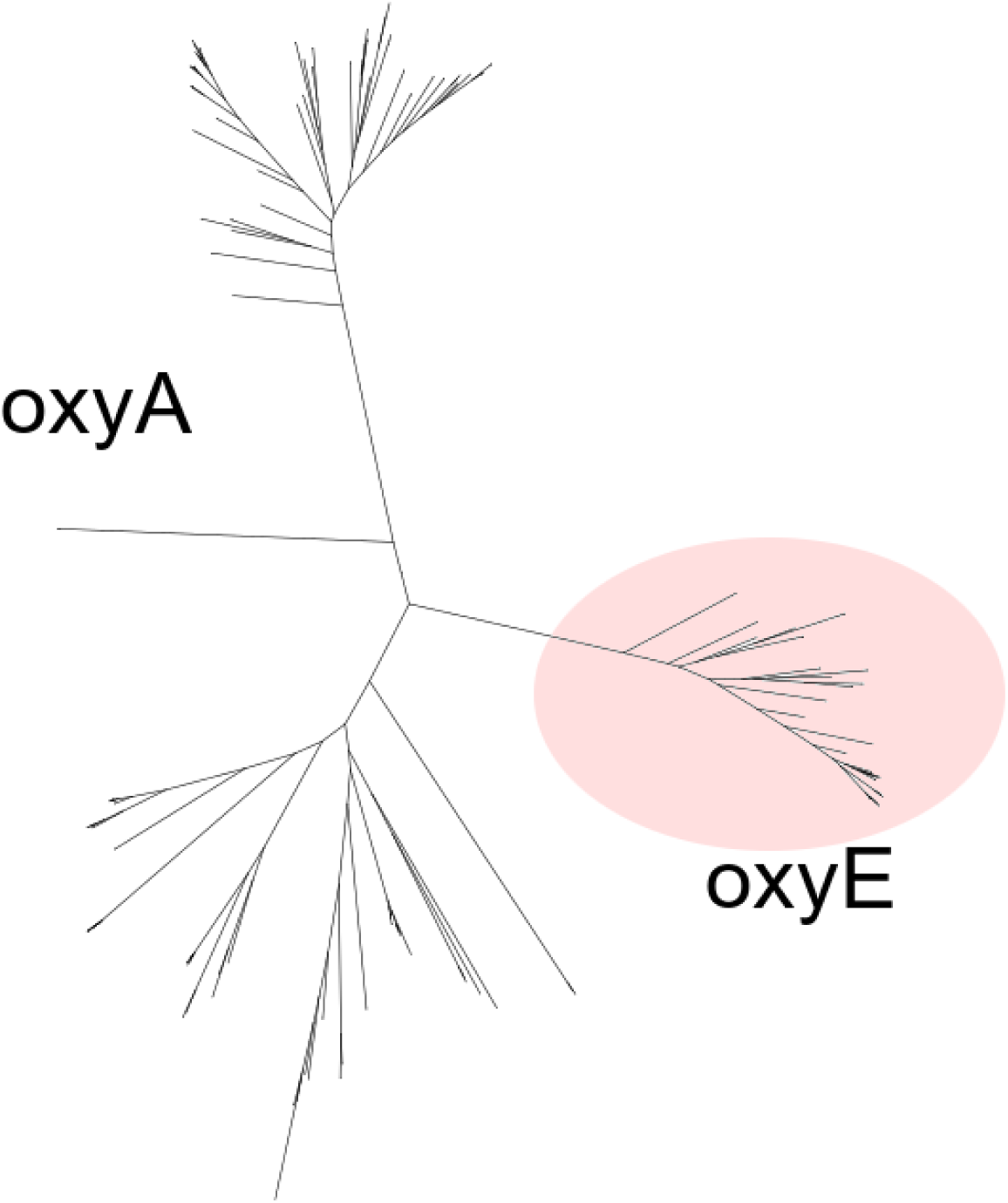
Phylogenetic tree (unrooted) of the mixed zol orthogroup OG3, which includes P450 monooxygenases *oxyA* and *oxyE*. The clades of the two genes can be separated after annotation of the corresponding enzymes with known function. The known oxyE genes all belong to the clade in the coloured circle and were removed from OG3 and assigned to the artificial group OG3e. Trees of both OG3 and OG3e can be explored in **Supplementary Data 1**.

**<u>Supplementary Figure 2:</u>** High-resolution concatenated phylogeny of dalabactin and murobactin encoding BGCs (**Supplementary Data 2**, **Supplementary Table 3**). Information including types, backbone amino acids and gene content of the BGCs are plotted next to the tree.

**Supplementary Figure 3:**
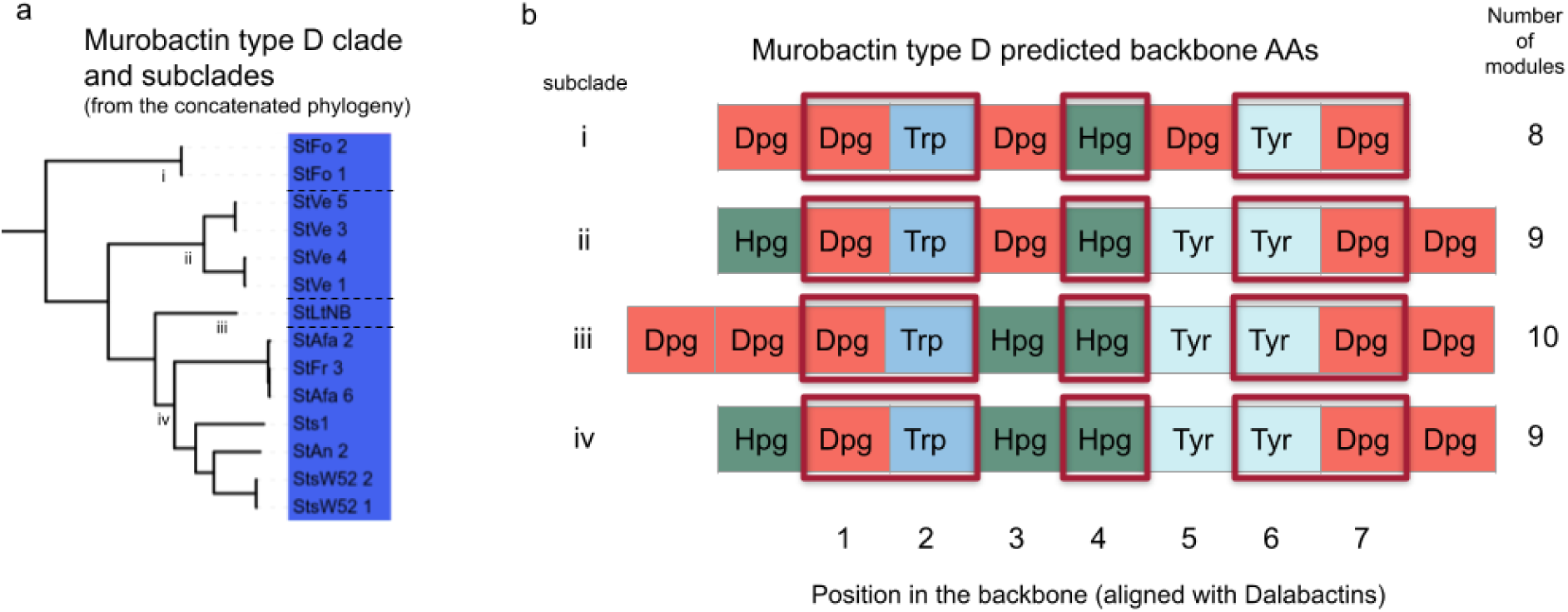
Murobactin type D subclades and backbone compositions. Panel a: clade of Murobactin type D, with marked subclades i-iv. BGC IDs are shown as leaf labels. Panel b: predicted backbone connected to its corresponding subclade. The red squares highlight the characteristic motif of the type.

**Supplementary Figure 4:**
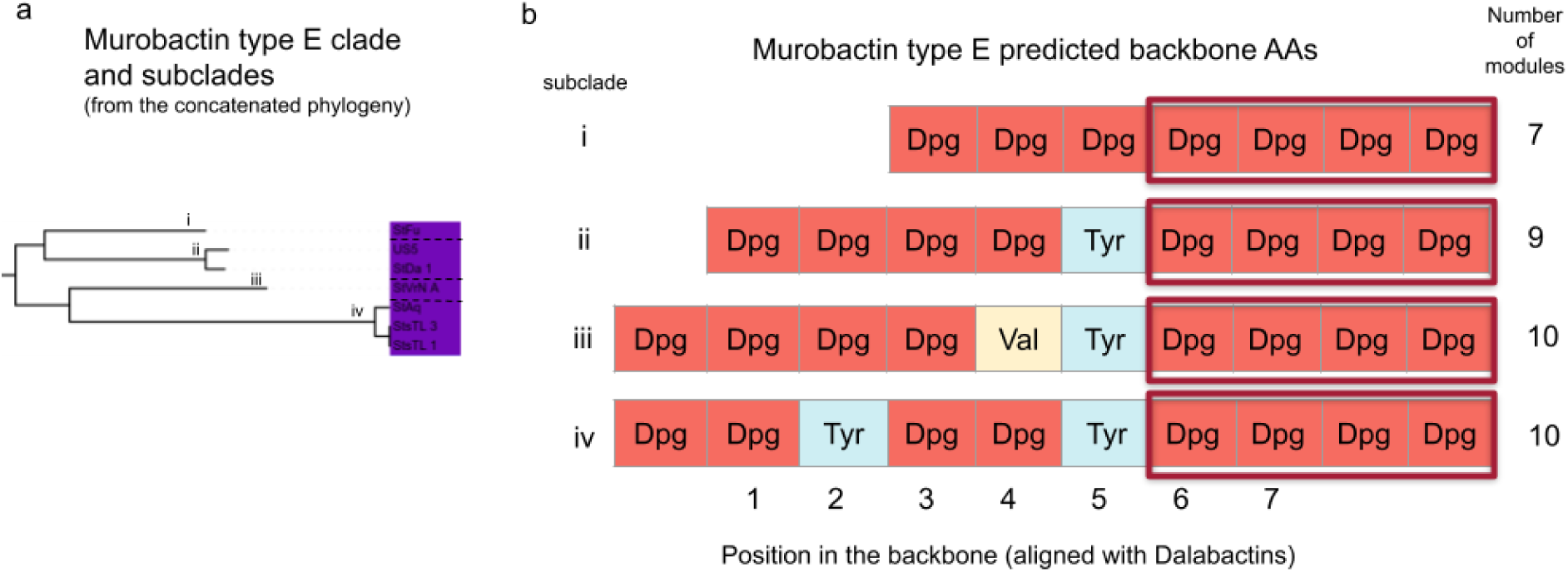
Murobactin type E subclades and backbone compositions. Panel a: clade of murobactin type E, with marked subclades i-iv. BGC IDs are shown as leaf labels. Panel b: predicted backbone connected to its corresponding subclade. The red squares highlight the characteristic motif of the type.

**Supplementary Figure 5:**
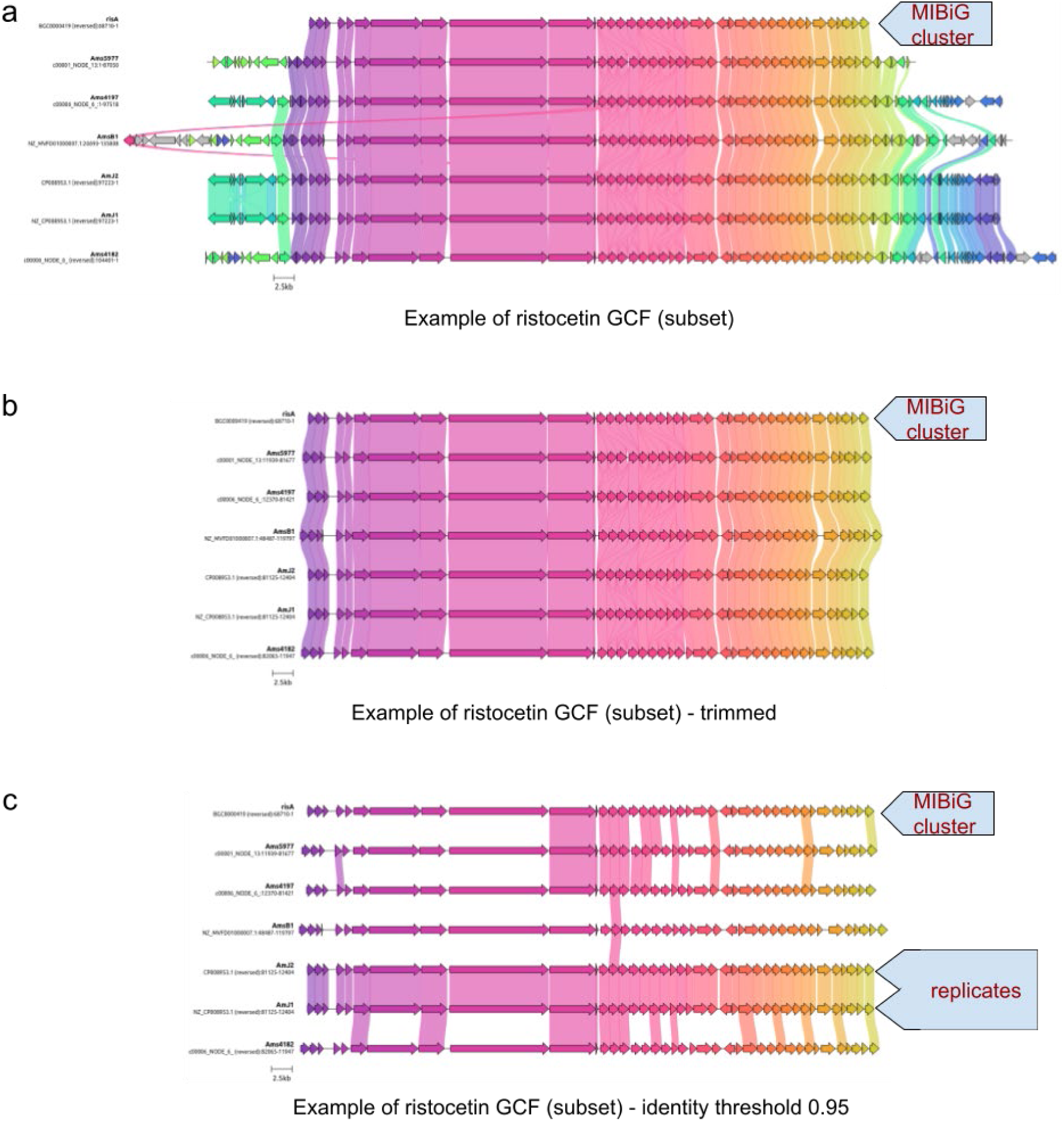
Example of clinker-based trimming and dereplication. Panel a: subset of the ristocetin gene cluster family (GCF), visualised with clinker. Genes are coloured the same if they were placed in the same gene group based on similarity by clinker and are connected by coloured bands if their identity is higher than 0.5. The MIBIG cluster is the top one (marked) and the trimming will be based on it. Panel b: the same visualisation as panel a, after the trimming is finished. Now all BGCs belonging to this GCF are uniformly trimmed. Panel c:

## Acknowledgements

AG and JR are grateful for the support of the Deutsche Forschungsgemeinschaft (DFG; Project ID No. 398967434-TRR 261). NZ and MA were supported by the German Center for Infection Research (DZIF) (TTU 09.716). NZ, MA, ES, NK, WW thank CMFI (Germany’s Excellence Strategy–EXC 2124–390838134) for funding and structural support. NK, NZ, MC, OG and ES were supported through the JPIAMR project AVOGADRO, with funding for the German partners provided by the Bundesministerium für Forschung, Technologie und Raumfahrt (BMFTR, formerly BMBF; grant no. 01KI2503). This work was supported by Monash University and EMBL Australia. This research was conducted by the Australian Research Council Centre of Excellence for Innovations in Peptide and Protein Science (CE200100012) and funded by the Australian Government. We thank Ana Monica Daneliuc for her assistance in conducting the HMM search. We also thank L. do Presti for invaluable comments on the manuscript. HMM search of X-domain in RefSeq and Genbank proteomes was performed on the IBMI nodes.

## Conflict of interest

The following authors have no conflict of interest to declare: Athina Gavriilidou, Wolfgang Wohlleben, Nadine Ziemert, Noel Kubach, Martina Adamek,

## Notes

### Competing Interest Statement

The authors have declared no competing interest.

### Summary of Updates

Complete restructuring of the manuscript with extended dataset and clearly defined nomenclature.

