## Supplementary Data 5 for "Structural and evolutionary analyses support reclassification of glycopeptide antibiotics as xyclopeptides": A_gpa_dataset_generation.ipynb

```
# %% [code]
##### IMPORTS or maybe specify also the anaconda environment
from pprint import pprint
import pandas as pd
import numpy as np
import os
import json

# %% [markdown]
# Creation of dataset of GPA encoding BGCs
#### Sources:
- publications
- database search

# %% [code]
##### file paths - at some point add script to create all of there if they don't exist yet!!!!!!!!!!!!!!
basedir = '/media/hdd_2tb/GPA_reclassification_clean'
dataset_creation_dir = os.path.join(basedir, 'A_dataset_creation')
publication_scripts_dir = os.path.join(basedir, 'publication_scripts')
### 0: HMMsearch
hmmsearch_dir = os.path.join(dataset_creation_dir, '0_HMMsearch')
hmmsearch_logs_dir = os.path.join(hmmsearch_dir, 'log_files')
hmm_Xdom_as6_file = os.path.join(hmmsearch_dir, 'as6_Xdomain.hmm')
### 1: downloads
downloads_dir = os.path.join(dataset_creation_dir, '1_positive_hits_downloads')
summary_downloads_table = os.path.join(downloads_dir, 'summary_hits_metadata.csv')
downloads_meta_tables_list = []
### 2: putative GPA BGCs
putative_gpas_dir = os.path.join(dataset_creation_dir, '2_putative_GPAs_analyses')
### 3: clustering and dereplication
dereplication_clustering_dir = os.path.join(dataset_creation_dir, '3_GPA_candidate_dereplication')
### 4: orthology inference
orthology_inf_dir = os.path.join(dataset_creation_dir, '4_orthology_inference')
### 5: basic phylogenetics
basicphylogenetics_dir = os.path.join(dataset_creation_dir, '5_basic_phylogenetics')
### 6: concatenated phylogeny
concatphylogeny_dir = os.path.join(dataset_creation_dir, '6_concat_phylogeny')


### older
hmmsearch_result = os.path.join(dataset_creation_dir, 'ncbi_hmmsearch_xdom_allT300_hits.csv')
todownload_table = os.path.join(dataset_creation_dir, 'GPA_clusters_for_download.csv')
todownload_table = os.path.join(dataset_creation_dir, 'GPA_clusters_for_download_withnew.tsv') ### for testing new genomes
print(todownload_table)
todownload_dir = os.path.join(dataset_creation_dir, 'raw_dataset')

# %% [markdown]
# Step 0: HMMsearch for X domain
Database search was conducted on multiple databases:
- MIBiG
- Literature (Culp2020)
- RefSeq
- Genbank
- Mgnify
- Local datasets (MAGs)
- Sequenced genome (Sva)?

An X-domain HMM from antiSMASH v6 was used with the hmmsearch (3.3) function on the protein sequences of the target databases (protein fasta files). A local cutoff of 300 was chosen based on known clusters.

# %% [markdown]
## Search of MIBiG database

# %% [code]
### X domain HMM search in MIBiG
mibig_hmmsearch_dir = os.path.join(hmmsearch_dir, 'MIBiG_hmmsearch')
mibig_hmm_out = os.path.join(mibig_hmmsearch_dir, 'mibig_xdom_hits')
mibig_hmm_tblout = os.path.join(mibig_hmmsearch_dir, 'mibig_xdom_seq_hits.tsv')
mibig_hmm_domtblout = os.path.join(mibig_hmmsearch_dir, 'mibig_xdom_dom_hits.tsv')
mibig_hmm_domtblout_xlsx = os.path.join(mibig_hmmsearch_dir, 'mibig_xdom_dom_hits.xlsx')
mibig_hmm_pfamtblout = os.path.join(mibig_hmmsearch_dir, 'mibig_xdom_pfam_hits.tsv')
mibig_fasta_file = os.path.join(mibig_hmmsearch_dir, 'mibig_prot_seqs_3.1.fasta')
cpus = 30
domT = 300
### commented out to avoid unnecessary download
if False:
    ### Download fasta file from database
    if not os.path.exists(mibig_fasta_file):
        mibig_download_command = 'wget https://dl.secondarymetabolites.org/mibig/mibig_prot_seqs_3.1.fasta'
        os.system(mibig_download_command)
    ### hmmsearch command for MIBiG
    mibig_hmmsearch_command = 'hmmsearch -o %s --tblout %s --domtblout %s --pfamtblout %s --cpu %s --domT %s %s %s' \
                              %(mibig_hmm_out, mibig_hmm_tblout, mibig_hmm_domtblout, mibig_hmm_pfamtblout, cpus, domT, hmm_Xdom_as6_file, mibig_fasta_file)
    os.system(mibig_hmmsearch_command)

### function to nicely visualize hmmsearch tblout files
def import_hmmsearch_tblout(filepath):
    # manually specify col limits
    dom_hits_colspecs = [(0, 252), (252, 264), (264, 270), (270, 291), (291, 302), (302, 308), (308, 318), (318, 325),
                         (325, 331), (331, 335), (335, 339), (339, 349), (349, 359), (359, 366), (366, 372), (372, 378),
                         (378, 384), (384, 390), (390, 396), (396, 402), (402, 408), (408, 413), (413, 435)]
    tblout_df = pd.read_fwf(filepath, skiprows=3, index_col=False,
                                             colspecs=dom_hits_colspecs,
                                             names=['target_name', 'accession1', 'tlen', 'query_name', 'accession2',
                                                    'full_sequence_qlen', 'full_sequence_E-value',
                                                    'full_sequence_score', 'full_sequence_bias', 'this_domain_#',
                                                    'this_domain_of', 'this_domain_c-Evalue', 'this_domain_i-Evalue',
                                                    'this_domain_score', 'this_domain_bias', 'hmm_coord_from',
                                                    'hmm_coord_to', 'ali_coord_from', 'ali_coord_to', 'env_coord_from',
                                                    'env_coord_to', 'acc', 'description_of_target'])
    ### skip version lines
    return tblout_df.iloc[:-10]

### preview of the results
mibig_hmmsearch_domhits_df = import_hmmsearch_tblout(mibig_hmm_domtblout)
#mibig_positive_hits_df = mibig_hmmsearch_domhits_df[mibig_hmmsearch_domhits_df.iloc[:,13]>domT] ### unnecessary since domT
print("Number of hits: %s" %(len(mibig_hmmsearch_domhits_df.index)))
mibig_hmmsearch_domhits_df.iloc[:5]

# %% [markdown]
## Search of known dataset (Culp et al 2020, Waglechner et al 2019)

# %% [code]
### info files needed can be downloaded here:
waglechner_stable1_link = 'https://static-content.springer.com/esm/art%3A10.1038%2Fs41564-019-0531-5/' \
                          'MediaObjects/41564_2019_531_MOESM3_ESM.xlsx'
waglechner_git_link = 'https://github.com/waglecn/GPA_evolution.git'
### paths
culp_hmmsearch_dir = os.path.join(hmmsearch_dir, 'Culp_dataset_hmmsearch')
waglechner_stable1_path = os.path.join(culp_hmmsearch_dir, '41564_2019_531_MOESM3_ESM.xlsx')
finished_table_path = os.path.join(culp_hmmsearch_dir, 'culp_finished_searched.csv')
### commented out to avoid unnecessary analysis
if False:
    %run hmmsearch_culp_dataset.py -o $culp_hmmsearch_dir -m $hmm_Xdom_as6_file -t $waglechner_stable1_path -f $finished_table_path

### preview of the results
culp_summary_domhitsfile = os.path.join(culp_hmmsearch_dir, "hmmsearch_summary_dom_hits.tsv")
culp_dataset_hmmsearch_domhits_df = pd.read_csv(culp_summary_domhitsfile, nrows=5)
culp_dataset_hmmsearch_domhits_df

# %% [markdown]
## Search of RefSeq database

# %% [code]
### paths
refseq_hmmsearch_dir = os.path.join(hmmsearch_dir, 'RefSeq_hmmsearch')
refseq_assembly_summary_table = os.path.join(refseq_hmmsearch_dir, 'assembly_summary_refseq.txt') # this could be > 90 MB
refseq_finished_table_path = os.path.join(refseq_hmmsearch_dir, 'refseq_finished_searched.csv')
### commented out to avoid unnecessary analysis (takes a few days time)
if False:
    %run hmmsearch_ncbi_ftp.py -i $refseq_assembly_summary_table -o $refseq_hmmsearch_dir -m $hmm_Xdom_as6_file -f $refseq_finished_table_path
### preview of the results
refseq_summary_domhitsfile = os.path.join(refseq_hmmsearch_dir, "hmmsearch_summary_dom_hits.tsv")
refseq_dataset_hmmsearch_domhits_df = pd.read_csv(refseq_summary_domhitsfile, nrows=5)
refseq_dataset_hmmsearch_domhits_df

# %% [markdown]
## Search of Genbank database

# %% [code]
### paths
genbank_hmmsearch_dir = os.path.join(hmmsearch_dir, 'Genbank_hmmsearch')
genbank_assembly_summary_table = os.path.join(genbank_hmmsearch_dir, 'assembly_summary_genbank.txt') # this could be > 750 MB
genbank_finished_table_path = os.path.join(genbank_hmmsearch_dir, 'genbank_finished_searched.csv')
### commented out to avoid unnecessary analysis (takes a few days time)
if False:
    %run hmmsearch_ncbi_ftp.py -i $genbank_assembly_summary_table -o $genbank_hmmsearch_dir -m $hmm_Xdom_as6_file -f $genbank_finished_table_path
### preview of the results
genbank_summary_domhitsfile = os.path.join(genbank_hmmsearch_dir, "hmmsearch_summary_dom_hits.tsv")
genbank_dataset_hmmsearch_domhits_df = pd.read_csv(genbank_summary_domhitsfile, nrows=5)
genbank_dataset_hmmsearch_domhits_df

# %% [markdown]
## Search of MGnify database

# %% [code]
### paths
mgnify_hmmsearch_dir = os.path.join(hmmsearch_dir, 'MGnify_hmmsearch')
mgnify_assembly_summary_table = os.path.join(mgnify_hmmsearch_dir, 'atlas_public_mgnify_asms_230531_toprocess.csv') # this file is manually extracted from the MGnify API and contains all the accession numbers that need to be processed. The date that the database was accessed is included in the filename.
mgnify_finished_table_path = os.path.join(mgnify_hmmsearch_dir, 'mgnify_finished_searched.csv')
### commented out to avoid unnecessary analysis (takes a couple days time)
if False:
    %run hmmsearch_mgnify_asms.py -i $mgnify_assembly_summary_table -f $mgnify_finished_table_path -o $mgnify_hmmsearch_dir -m $hmm_Xdom_as6_file
### alternatively:
print('conda activate gpas\npython %s/hmmsearch_mgnify_asms.py -i %s -f %s -o %s -m %s' %(publication_scripts_dir, mgnify_assembly_summary_table, mgnify_finished_table_path, mgnify_hmmsearch_dir, hmm_Xdom_as6_file))

### preview of the results
mgnify_summary_domhitsfile = os.path.join(mgnify_hmmsearch_dir, "hmmsearch_summary_dom_hits.tsv")
mgnify_dataset_hmmsearch_domhits_df = pd.read_csv(mgnify_summary_domhitsfile, nrows=5)
mgnify_dataset_hmmsearch_domhits_df

# %% [markdown]
## Search of sequenced genome (S. varsoviensis)

# %% [code]
### X domain HMM search in MIBiG
localseq_hmmsearch_dir = os.path.join(hmmsearch_dir, 'Local_Sequencing_hmmsearch')
localseq_hmm_out = os.path.join(localseq_hmmsearch_dir, 'localseq_xdom_hits')
localseq_hmm_tblout = os.path.join(localseq_hmmsearch_dir, 'localseq_xdom_seq_hits.tsv')
localseq_hmm_domtblout = os.path.join(localseq_hmmsearch_dir, 'localseq_xdom_dom_hits.tsv')
localseq_hmm_domtblout_xlsx = os.path.join(localseq_hmmsearch_dir, 'localseq_xdom_dom_hits.xlsx')
localseq_hmm_pfamtblout = os.path.join(localseq_hmmsearch_dir, 'localseq_xdom_pfam_hits.tsv')
localseq_fasta_file = os.path.join(localseq_hmmsearch_dir, 'Sva.faa') ### need faa actually !!!!!!! run prokka or sth and redo !!!!!!!!!!!!!!!!!!!!!!!!!!!!!!!!
cpus = 30
domT = 300
### commented out to avoid unnecessary analysis
if False:
    ### hmmsearch command for MIBiG
    localseq_hmmsearch_command = 'hmmsearch -o %s --tblout %s --domtblout %s --pfamtblout %s --cpu %s --domT %s %s %s' \
                              %(localseq_hmm_out, localseq_hmm_tblout, localseq_hmm_domtblout, localseq_hmm_pfamtblout, cpus, domT, hmm_Xdom_as6_file, localseq_fasta_file)
    os.system(localseq_hmmsearch_command)

### function to nicely visualize hmmsearch tblout files
def import_hmmsearch_tblout(filepath):
    # manually specify col limits --- this needs fixing for here - the first column is much shorter - goes up to 22 or so!!!!!!!!!!
    dom_hits_colspecs = [(0, 252), (252, 264), (264, 270), (270, 291), (291, 302), (302, 308), (308, 318), (318, 325),
                         (325, 331), (331, 335), (335, 339), (339, 349), (349, 359), (359, 366), (366, 372), (372, 378),
                         (378, 384), (384, 390), (390, 396), (396, 402), (402, 408), (408, 413), (413, 435)]
    tblout_df = pd.read_fwf(filepath, skiprows=3, index_col=False,
                                             colspecs=dom_hits_colspecs,
                                             names=['target_name', 'accession1', 'tlen', 'query_name', 'accession2',
                                                    'full_sequence_qlen', 'full_sequence_E-value',
                                                    'full_sequence_score', 'full_sequence_bias', 'this_domain_#',
                                                    'this_domain_of', 'this_domain_c-Evalue', 'this_domain_i-Evalue',
                                                    'this_domain_score', 'this_domain_bias', 'hmm_coord_from',
                                                    'hmm_coord_to', 'ali_coord_from', 'ali_coord_to', 'env_coord_from',
                                                    'env_coord_to', 'acc', 'description_of_target'])
    ### skip version lines
    return tblout_df.iloc[:-10]

### preview of the results
localseq_hmmsearch_domhits_df = import_hmmsearch_tblout(localseq_hmm_domtblout)
#mibig_positive_hits_df = mibig_hmmsearch_domhits_df[mibig_hmmsearch_domhits_df.iloc[:,13]>domT] ### unnecessary since domT
print("Number of hits: %s" %(len(localseq_hmmsearch_domhits_df.index)))
localseq_hmmsearch_domhits_df.iloc[:5]

# %% [markdown]
## Search of Stammsamlung Tuebingen 2022

# %% [code]
### Process was conducted on micromatrix server - hits extracted with samtools (see notes from 29.06)
### download hits, create metadata table and check manually after antismash!!!!!!!!!!!!!!!!!!!!!!!!!!!!!!!!!!!!!!!!!!!!!!!!!!!!!!!!!!!!!!!!!!!!!!!!!!!!!!!!!!!!!!!!!!!!!!
### paths
stammsamlung22_hmmsearch_dir = os.path.join(hmmsearch_dir, 'Stammsamlung2022_hmmsearch')
stammsamlung22_finished_table_path = os.path.join(stammsamlung22_hmmsearch_dir, 'stammsammlung2022_finished_searched.csv')
### commented out since input data is not made available - also it was run on the server
if False:
    %run hmmsearch_localdataset.py -i $stammsamlung22_faas_dir -f $stammsamlung22_finished_table_path -o $stammsamlung22_hmmsearch_dir -m $hmm_Xdom_as6_file
### preview of the results
stammsamlung22_summary_domhitsfile = os.path.join(stammsamlung22_hmmsearch_dir, "hmmsearch_summary_dom_hits.tsv")
stammsamlung22_dataset_hmmsearch_domhits_df = pd.read_csv(stammsamlung22_summary_domhitsfile, nrows=5)
stammsamlung22_dataset_hmmsearch_domhits_df

# %% [markdown]
## Search of Shrikant's old Stammsamlung (Nanopore) antismash5 results - turned faas

# %% [code]
### paths
shrikandold_hmmsearch_dir = os.path.join(hmmsearch_dir, 'Shrikant_olddata_hmmsearch')
shrikandold_finished_table_path = os.path.join(shrikandold_hmmsearch_dir, 'shrikantStammsamlung_assembliesAntismash_finished_searched.csv')
### commented out since input data is not made available
if False:
    %run hmmsearch_localdataset.py -i $shrikandold_assemblies_faas_dir -f $shrikandold_finished_table_path -o $shrikandold_hmmsearch_dir -m $hmm_Xdom_as6_file
### preview of the results
shrikandold_summary_domhitsfile = os.path.join(shrikandold_hmmsearch_dir, "hmmsearch_summary_dom_hits.tsv")
shrikandold_dataset_hmmsearch_domhits_df = pd.read_csv(shrikandold_summary_domhitsfile, nrows=5)
shrikandold_dataset_hmmsearch_domhits_df


# %% [markdown]
## Search of Shrikant's old Stammsamlung (TellSeq) data - after gene calling (prodigal)

# %% [code]
### paths:
shrikandold_telseq_hmmsearch_dir = os.path.join(hmmsearch_dir, 'Shrikantold_TellSeq_hmmsearch')
shrikandold_telseq_finished_table_path = os.path.join(shrikandold_telseq_hmmsearch_dir, 'shrikantStammsamlung_tellseq_prodigal_finished_searched.csv')
### commented out since input data is not made available
if False:
    %run hmmsearch_localdataset.py -i $shrikandold_telseq_hmmsearch_dir -f $shrikandold_telseq_finished_table_path -o $shrikandold_telseq_hmmsearch_dir -m $hmm_Xdom_as6_file -t fasta-files-fasta
### preview of the results
shrikandold_telseq_summary_domhitsfile = os.path.join(shrikandold_telseq_hmmsearch_dir, "hmmsearch_summary_dom_hits.tsv")
shrikandold_telseq_dataset_hmmsearch_domhits_df = pd.read_csv(shrikandold_telseq_summary_domhitsfile, nrows=5)
shrikandold_telseq_dataset_hmmsearch_domhits_df

# %% [markdown]
### Failed searches (0 hits):
* Caner's latest soil dataset (ran on IBMI server under A_dataset_creation/0_HMMsearch/Caner_data_hmmsearch)
* Shrikant's (published?) and Caner's soil long reads (Jan 2020) (ran on micromatrix server under /vol/tellseq/athina-Xdomain/shrikantsoil_longreads_hmmsearch)
* Shrikant's (published?) and Caner's soil long reads (2019) (ran on micromatrix server under /vol/tellseq/athina-Xdomain/shrikantsoil_2019_lr_hmmsearch_results)
* Shrikant's (published?) and Caner's soil short reads (ran on micromatrix server under /vol/tellseq/athina-Xdomain/shrikantsoil_shortreads_hmmsearch_results)
* Will's contigs from the Antarctic volcano sampling sites (antismash files and also contigs_faas - ran locally under /media/hdd_2tb/GPA_reclassification_clean/A_dataset_creation/0_HMMsearch/Will_dataset_hmmsearch/hmmsearch_output)

# %% [markdown]
# Step 1: Downloading related sequences

# %% [markdown]
## Downloading putative GPA clusters from MIBiG

# %% [code]
### Downloading MIBiG clusters and generating metadata table
mibig_download_folder = os.path.join(downloads_dir, 'MIBiG_hits_downloads')
mibig_file_suffix = 'gbk'
mibig_download_metadata_table = os.path.join(mibig_download_folder, 'mibig_hits_metadata.csv')
downloads_meta_tables_list.append(mibig_download_metadata_table)
if False:
    %run download_mibig_dataset.py -i $mibig_hmm_domtblout -o $mibig_download_folder -t $mibig_download_metadata_table
mibig_download_df = pd.read_csv(mibig_download_metadata_table, nrows=5)
mibig_download_df

# %% [markdown]
## Downloading literature dataset (Culp et al, 2020; Waglechner et al, 2019)

# %% [code]
### Downloading table of information on dataset (Waglechner et al, 2019: Supplementary table 1)
waglechner_stable1_link = 'https://static-content.springer.com/esm/art%3A10.1038%2Fs41564-019-0531-5/MediaObjects/41564_2019_531_MOESM3_ESM.xlsx'
waglechner_git_link = 'https://github.com/waglecn/GPA_evolution.git'
culp_download_dir = os.path.join(downloads_dir, 'Culp_dataset_downloads')
culp_file_suffix = 'gbk'
waglechner_stable1_path = os.path.join(culp_download_dir, '41564_2019_531_MOESM3_ESM.xlsx')
culp_download_metadata_table = os.path.join(culp_download_dir, 'culp_dataset_metadata.csv')
downloads_meta_tables_list.append(culp_download_metadata_table)
culp_hmmsearch_result_summary_table = os.path.join(culp_hmmsearch_dir, 'hmmsearch_summary_dom_hits.tsv')
### Here add --genomes True to activate download of genomes when possible
if False:
    %run generate_culp_table.py -i $culp_download_dir -o $culp_download_metadata_table -t $waglechner_stable1_path -d $culp_summary_domhitsfile
culp_metadata_df = pd.read_csv(culp_download_metadata_table, nrows=5)
###
print("\nNOTE: Manually removed gene and CD with locus_tag A3CE_RS0142385 and A3CE_RS0142730 from the 05_cluster_AbalhimycinaFH1894.gbk file, because the feature translation extends out of record.\nNOTE: Manually removed gene and CD with locus_tag ATP06_RS07260 and ATP06_RS07590 from the 11_cluster_AregGY080.gbk file, because the feature translation extends out of record.")
### show preview of table
culp_metadata_df

# %% [markdown]
## Downloading genomes with putative GPA clusters from RefSeq

# %% [markdown]
## Downloading genomes with putative GPA clusters from Genbank

# %% [code]
### paths
refseq_download_dir = os.path.join(downloads_dir, 'Refseq_hits_downloads')
refseq_file_suffix = 'gbff'
refseq_metadata_table = os.path.join(refseq_download_dir, 'refseq_hits_metadata.csv')
downloads_meta_tables_list.append(refseq_metadata_table)
refseq_hmmsearch_result_summary_table = os.path.join(refseq_hmmsearch_dir, 'hmmsearch_summary_dom_hits.tsv')
### commented out to avoid unnecessary download
if False:
    %run download_ncbi_dataset.py -i $refseq_finished_table_path -t $refseq_assembly_summary_table -o $refseq_download_dir -m $refseq_metadata_table -d $refseq_hmmsearch_result_summary_table
### show preview of table
refseq_metadata_df = pd.read_csv(refseq_metadata_table, nrows=5)
refseq_metadata_df#.iloc[:5]

# %% [code]
### paths
genbank_download_dir = os.path.join(downloads_dir, 'Genbank_hits_downloads')
genbank_file_suffix = 'gbff'
genbank_metadata_table = os.path.join(genbank_download_dir, 'genbank_hits_metadata.csv')
downloads_meta_tables_list.append(genbank_metadata_table)
genbank_hmmsearch_result_summary_table = os.path.join(genbank_hmmsearch_dir, 'hmmsearch_summary_dom_hits.tsv')
### commented out to avoid unnecessary download
if False:
    %run download_ncbi_dataset.py -i $genbank_finished_table_path -t $genbank_assembly_summary_table -o $genbank_download_dir -m $genbank_metadata_table -d $genbank_hmmsearch_result_summary_table
### show preview of table
genbank_metadata_df = pd.read_csv(genbank_metadata_table, nrows=5)
genbank_metadata_df

# %% [markdown]
## Downloading MAGs with putative GPA clusters from MGnify

# %% [code]
### paths
mgnify_download_dir = os.path.join(downloads_dir, 'MGnify_hits_downloads')
mgnify_file_suffix = 'fasta'
mgnify_metadata_table = os.path.join(mgnify_download_dir, 'mgnify_hits_metadata.csv')
downloads_meta_tables_list.append(mgnify_metadata_table)
mgnify_hmmsearch_result_summary_table = os.path.join(mgnify_hmmsearch_dir, 'hmmsearch_summary_dom_hits.tsv')
###
### commented out to avoid unnecessary download
if False:
    %run download_mgnify_dataset.py -i $mgnify_finished_table_path -t $mgnify_assembly_summary_table -o $mgnify_download_dir -m $mgnify_metadata_table -d $mgnify_hmmsearch_result_summary_table
### show preview of table
mgnify_metadata_df = pd.read_csv(mgnify_metadata_table, nrows=5)
mgnify_metadata_df

# %% [markdown]
## Extracting sequence of Local Sequencing project (Sva) with putative GPA clusters

# %% [code]
### paths
localseq_extract_dir = os.path.join(downloads_dir, 'Local_Sequencing_hits_exports')
localseq_file_suffix = 'fna'
localseq_metadata_table = os.path.join(localseq_extract_dir, 'localseq_hits_metadata.csv')
downloads_meta_tables_list.append(localseq_metadata_table)
### show preview of table
localseq_metadata_df = pd.read_csv(localseq_metadata_table, nrows=5)
localseq_metadata_df

# %% [markdown]
## Extracting sequences of Stammsamlung Tuebingen 2022 with putative GPA clusters

# %% [code]
### paths
stammsamlung2022_extract_dir = os.path.join(downloads_dir, 'Stammsamlung2022_hits_exports')
stammsamlung2022_file_suffix = 'fasta'
stammsamlung2022_metadata_table = os.path.join(stammsamlung2022_extract_dir, 'stammsamlung2022_hits_metadata.csv')
downloads_meta_tables_list.append(stammsamlung2022_metadata_table)
### show preview of table
stammsamlung2022_metadata_df = pd.read_csv(stammsamlung2022_metadata_table, nrows=5)
stammsamlung2022_metadata_df

# %% [markdown]
## Extracting sequences of Shrikant's old Stammsamlung (Nanopore) antismash5 results with putative GPA clusters

# %% [code]
### paths
shrikandold_extract_dir = os.path.join(downloads_dir, 'Shrikant_olddata_hits_exports')
shrikandold_file_suffix = 'fna'
shrikandold_metadata_table = os.path.join(shrikandold_extract_dir, 'shrikantolddata_hits_metadata.csv')
downloads_meta_tables_list.append(shrikandold_metadata_table)
### show preview of table
shrikandold_metadata_df = pd.read_csv(shrikandold_metadata_table, nrows=5)
shrikandold_metadata_df

# %% [markdown]
## Extracting sequences of Shrikant's old Stammsamlung (TelSeq) with putative GPA clusters

# %% [code]
### paths
shrikandold_telseq_extract_dir = os.path.join(downloads_dir, 'Shrikantold_TellSeq_hits_exports')
shrikandold_telseq_file_suffix = 'fasta'
shrikandold_telseq_metadata_table = os.path.join(shrikandold_telseq_extract_dir, 'shrikantoldtellseq_hits_metadata.csv')
downloads_meta_tables_list.append(shrikandold_telseq_metadata_table)
### show preview of table
shrikandold_telseq_metadata_df = pd.read_csv(shrikandold_telseq_metadata_table, nrows=5)
shrikandold_telseq_metadata_df

# %% [markdown]
## Summary of everything so far

# %% [code]
### read or make table
if os.path.exists(summary_downloads_table):# and False:
    summary_downloads_table_df = pd.read_csv(summary_downloads_table)
else:
    dfs_list = []
    for mtable in downloads_meta_tables_list:
        #print(mtable)
        #print(len(pd.read_csv(mtable)))
        dfs_list.append(pd.read_csv(mtable))
    summary_downloads_table_df = pd.concat(dfs_list).drop_duplicates().reset_index(drop=True)
    summary_downloads_table_df = pd.concat(dfs_list).reset_index(drop=True)
    #summary_downloads_table_df
    summary_downloads_table_df.to_csv(summary_downloads_table, index=False)
### show preview of table
#print("NOTE:\nThis table has been or should be manually edited to assign internal IDs!")
print("Total number of rows: %s" %(len(summary_downloads_table_df)))
summary_downloads_table_df#.iloc[:5]

# %% [markdown]
# Step 2: Isolation of putative GPA encoding clusters

# %% [markdown]
## Quick antiSMASH v7 analysis (all optional parameters off) of all sequences that had a positive hit

# %% [code]
### dictionary of download/extract directories and target files suffixes
hitdirs_to_suffix = {culp_download_dir:culp_file_suffix,genbank_download_dir:genbank_file_suffix, refseq_download_dir: refseq_file_suffix, localseq_extract_dir:localseq_file_suffix, mgnify_download_dir:mgnify_file_suffix, mibig_download_folder:mibig_file_suffix, shrikandold_extract_dir:shrikandold_file_suffix, stammsamlung2022_extract_dir:stammsamlung2022_file_suffix}
quick_antismash_results = os.path.join(putative_gpas_dir, '0_minimal_antismash7_full_results')
### run quick-and-dirty antiSMASH with all sequences that had a hit - commented out to avoid unnecessary analysis (takes a few hours minimum). Specify --overwrite True if necessary
if False:
    %run run_quick_antismash.py -i $downloads_dir -o $quick_antismash_results

### it may be better to run this script from the command line:
print('conda activate antismash7\npython %s/run_quick_antismash.py -i %s -o %s' %(publication_scripts_dir, downloads_dir, quick_antismash_results))

# %% [markdown]
## Detection of antiSMASH regions that include the X domain

# %% [code]
### detect the X-domain containing clusters only and move them to another folder for further analysis
xdomain_containing_regions_dir = os.path.join(putative_gpas_dir, '1_regions_with_Xdom')
cluster_summary_file = os.path.join(putative_gpas_dir, 'summary_hits_clusters_metadata.csv')
### commented out for speed, takes a few minutes
if False:
    %run detect_xdomain_regions.py -i $quick_antismash_results -t $summary_downloads_table -o $xdomain_containing_regions_dir -m $cluster_summary_file
#print("NOTE: eventually check if quick antiSMASH run of QPWMP018AI is finished (needed multiple days)!!!!!")
cluster_summary_table_df = pd.read_csv(cluster_summary_file, nrows=5)
cluster_summary_table_df

# %% [markdown]
## Full annotation of regions that include the X domain with bakta v1.8.1

# %% [code]
### here run Bakta on them - with parameters skip-trna and skip-tmrna to make sure no putative CDS are skipped
bakta_annotated_sequences_dir = os.path.join(putative_gpas_dir, '2_regions_with_Xdom_bakta_results')
bakta_db_dir = '/media/hdd_2tb/bakta_db/db'  ### replace with local database
### run bakta annotation with fnas of all sequences that had a hit - commented out to avoid unnecessary analysis (~19h for 324 clusters)
if False:
    %run run_bakta.py -i $xdomain_containing_regions_dir -o $bakta_annotated_sequences_dir -m $cluster_summary_file
### it may be better to run this script from the command line:
print('conda activate bakta\npython %s/run_bakta.py \\\n-i %s \\\n-o %s \\\n-m %s' %(publication_scripts_dir, xdomain_containing_regions_dir, bakta_annotated_sequences_dir, cluster_summary_file))


# %% [markdown]
## Full antiSMASH 7 analysis of only the relevant clusters

# %% [code]
full_antismash_results = os.path.join(putative_gpas_dir, '3_full_antismash7_regionsonly_results')
### full antiSMASH (all on) with all regions that contain an X domain - commented out to avoid unnecessary analysis (takes a few hours minimum). Specify --overwrite True if necessary
if False:
    %run run_full_antismash.py -i $xdomain_containing_regions_dir -o $full_antismash_results
### it may be better to run this script from the command line:
print('conda activate antismash7\npython %s/run_full_antismash.py \\\n-i %s \\\n-o %s' %(publication_scripts_dir, bakta_annotated_sequences_dir, full_antismash_results))

print("\nCheck here for full antiSMASH 7 results:\n%s" %(os.path.join(full_antismash_results, 'all_concatenated/index.html')))

# %% [markdown]
# Step 3: Similarity clustering and dereplication
BiG-SCAPE (latest) analysis (if mixed, take care of weights - we have an altered bigscape installation) of well annotated putative GPA clusters. Clusters with more than 99% similarity to each other that come from the same strain need to be dereplicated. Otherwise we keep them. We should divide the clusters in GCFs (we may need some threshold testing here) so that they are ready for the next step.

# %% [markdown]
## BiG-SCAPE (v1.1.5) MIBiG clusters in multiple thresholds to determine what to use for the full dataset

# %% [code]
### collect files in different folders for downstream analysis
mibig_only_annotated_as7_clusters = os.path.join(putative_gpas_dir, '4_mibig_annotated_as7_clusters')
all_annotated_as7_clusters = os.path.join(putative_gpas_dir, '5_all_candidate_as7_clusters')
### multi-threshold run for MIBiG clusters to figure out a good threshold that separates all groups (I-V)
bigscape_Ttesting_dir = os.path.join(dereplication_clustering_dir, '0_bigscape_mibigGPA_Ttesting_results')
### commented out to avoid unnecessary analysis
if False:
    %run run_bigscape.py -i $full_antismash_results -o $bigscape_Ttesting_dir -t "0.1,0.2,0.3,0.4,0.5,0.6,0.7,0.8,0.9"
### it may be better to run this script from the command line:
print('conda activate bigscape\npython %s/run_bigscape.py -i %s -o %s -t 0.1,0.2,0.3,0.4,0.5,0.6,0.7,0.8,0.9' %(publication_scripts_dir, full_antismash_results, bigscape_Ttesting_dir))
### Manually checked results here
print("\n Results can be visualized here: %s/index.html" %(bigscape_Ttesting_dir))
print("\nAfter manual inspection, T=0.2 seems to be a suitable threshold as it is the highest that separates types I-V")

# %% [markdown]
## BiG-SCAPE (v1.1.5) everything with chosen threshold

# %% [code]
### now run bigscape with defined threshold
bigscape_result_dir = os.path.join(dereplication_clustering_dir, '1_candidateBGC_bigscape_results')
all_annotated_as7_clusters = os.path.join(putative_gpas_dir, '5_all_candidate_as7_clusters')
### commented out to avoid unnecessary analysis
if False:
    %run run_bigscape.py -i $all_annotated_as7_clusters -o $bigscape_result_dir -t "0.2"
### it may be better to run this script from the command line:
print('conda activate bigscape\npython %s/run_bigscape.py \\\n-i %s \\\n-o %s \\\n-t 0.2' %(publication_scripts_dir, all_annotated_as7_clusters, bigscape_result_dir))
### Manually checked results here
print("\n Results can be visualized here: %s/index.html" %(bigscape_result_dir))

# %% [markdown]
## Visualization of Gene Cluster Families with clinker

# %% [code]
clinker_results_dir = os.path.join(dereplication_clustering_dir, '2_clinker_gcf_results')
### NOTE: latest version of run_clinker missed several clusters (did not include them in the analysis)!!!!!!!!!!!!!!!!!!!!!!!!!!!!!!!!!!!!!!!!!!!!!!!!1
### commented out to avoid unnecessary analysis (takes about 10 minutes) 
if False:
    %run run_clinker_on_clustergroups.py -i $all_annotated_as7_clusters -o $clinker_results_dir -m $bigscape_result_dir -cm $cluster_summary_file
### it may be better to run this script from the command line:
print('conda activate gpas\npython %s/run_clinker_on_clustergroups.py \\\n-i %s \\\n-o %s \\\n-m %s \\\n-cm %s' %(publication_scripts_dir, all_annotated_as7_clusters, clinker_results_dir, bigscape_result_dir, cluster_summary_file))
updatedsummaryfile = os.path.join(clinker_results_dir, 'summary_hits_clusters_metadata.csv')
updated_cluster_summary_table_df = pd.read_csv(updatedsummaryfile, nrows=5)
updated_cluster_summary_table_df

# %% [markdown]
## Manual trimming, filtering and dereplication per GCF
At this point each GCF annotated html file is visualized and the clusters are trimmed, incomplete ones are filtered out and a dereplication step is conducted to avoid using identical clusters. 
The information is stored in the clinker manually saved json files (e.g. gcf_56_part1_risA_trimmed.json) and in the metadata table (column considered_complete).
Also, a column GPA_type_original is added to the metadata table, to assign a GPA type according to the existing system.

Rules that we follow:
- If there is an MIBiG entry, we use it as a guide for trimming
- sometimes there are genes near the edges that are not included in MIBiG but based on more recent publications (ADD PUBLICATIONS ALSO HERE) may play a role in the biosynthesis. If detected, those are included in the cluster limits.
- If there is are multiple clusters from the same strain that are identical (all genes display > 0.95 identity), we keep only one of the clusters, preferably from RefSeq
- If a cluster seems to have noticeably fewer genes than the rest in its group, especially if it is on contig edge (but the decision is made on case-by-case) and even more if the existing genes have very high identity with other longer clusters in the same GCF, it is considered incomplete. That may include MIBiG clusters. Another rule may be also if it does not fulfill the Wright paper "necessary genes" (3 out of: oxyB,C, dpgC, hal and vanHAX).

Calculate about a day for a repeat of the manual process if new clusters are added

# %% [code]
### showing an example of trimming
from IPython.display import Image, display
listOfImageNames = ['clinker_example_original.png',
                    'clinker_example_trimmed.png',
                    'clinker_example_dereplication.png']
for imageName in listOfImageNames:
    display(Image(filename=imageName))
cluster_trimmed_summary_file = os.path.join(clinker_results_dir, 'summary_hits_clusters_metadata_trimmed.csv')
cluster_trimmed_summary_table_df = pd.read_csv(cluster_trimmed_summary_file, nrows=5)
cluster_trimmed_summary_table_df

# %% [markdown]
## Trimming the cluster files and adding the new info in the table

# %% [code]
trimmed_complete_clusters_dir = os.path.join(dereplication_clustering_dir, '3a_trimmed_complete_GPAclusters')
trimmed_incomplete_clusters_dir = os.path.join(dereplication_clustering_dir, '3b_trimmed_incomplete_GPAclusters')
untrimmed_complete_clusters_dir =os.path.join(dereplication_clustering_dir, '3c_untrimmed_complete_GPAclusters')
limits_clusters_metadata = os.path.join(dereplication_clustering_dir, 'summary_clusters_trimmed_metadata.csv')
### 1st step is json parser to get locus tags and locations of limits
### 2nd step is old gbk trimming script to get trimmed gbk, faa and fna files
### commented out to avoid unnecessary analysis
if False:
    %run extract_trimmed_limits.py -ic $clinker_results_dir -ia $all_annotated_as7_clusters -oc $trimmed_complete_clusters_dir -oi $trimmed_incomplete_clusters_dir -m $cluster_trimmed_summary_file -mo $limits_clusters_metadata -oco $untrimmed_complete_clusters_dir
### it may be better to run this script from the command line:
print('conda activate gpas\npython %s/extract_trimmed_limits.py \\\n-ic %s \\\n-ia %s \\\n-oc %s \\\n-oi %s \\\n-m %s \\\n-mo %s \\\n -oco %s' %(publication_scripts_dir,clinker_results_dir, all_annotated_as7_clusters, trimmed_complete_clusters_dir, trimmed_incomplete_clusters_dir, cluster_trimmed_summary_file, limits_clusters_metadata, untrimmed_complete_clusters_dir))
    

# %% [markdown]
## Visualizing trimmed clusters with antiSMASH

# %% [code]
trimmed_complete_clusters_antismashresults_dir = os.path.join(dereplication_clustering_dir, '4a_full_antismash7_completetrimmedGPAs_results')
### full antiSMASH (all on) with all regions that contain an X domain - commented out to avoid unnecessary analysis (takes a few hours minimum). Specify --overwrite True if necessary
if False:
    %run run_full_antismash.py -i $trimmed_complete_clusters_dir -o $trimmed_complete_clusters_antismashresults_dir
### it may be better to run this script from the command line:
print('conda activate antismash7\npython %s/run_full_antismash.py \\\n-i %s \\\n-o %s' %(publication_scripts_dir, trimmed_complete_clusters_dir, trimmed_complete_clusters_antismashresults_dir))

print("\nCheck here for full antiSMASH 7 results (only after the dataset is finalized):\n%s" %(os.path.join(trimmed_complete_clusters_antismashresults_dir, 'all_concatenated/index.html')))

# %% [markdown]
## Clustering trimmed clusters with BiG-SCAPE
Mostly for visualizing the difference.

# %% [code]
### now run bigscape with defined threshold
complete_trimmed_clusters_bigscape_result_dir = os.path.join(dereplication_clustering_dir, '4a_full_antismash7_completetrimmedGPAs_results')
### commented out to avoid unnecessary analysis
if False:
    %run run_bigscape.py -i $trimmed_complete_clusters_antismashresults_dir -o $complete_trimmed_clusters_bigscape_result_dir -t "0.2"
### it may be better to run this script from the command line:
print('conda activate bigscape\npython %s/run_bigscape.py \\\n-i %s \\\n-o %s \\\n-t 0.2' %(publication_scripts_dir, trimmed_complete_clusters_antismashresults_dir, complete_trimmed_clusters_bigscape_result_dir))
### Manually checked results here
print("\n Results can be visualized here: %s/index.html" %(complete_trimmed_clusters_bigscape_result_dir))

# %% [markdown]
# Step 4: Orthology inference

# %% [markdown]
## Using Zol to estimate orthogroups from MIBiG BGCs - using zol 1.3.9 but latest is 1.3.11!!!!!

# %% [code]
### now run Zol on mibig clusters first
mibig_zol_results_dir = os.path.join(orthology_inf_dir, '0a_untrimmed_mibig_ozol_results')
mibig_orthogroups_dir = os.path.join(mibig_zol_results_dir, 'Ortholog_Group_Processing')
mibig_ogs_faas_dir = os.path.join(mibig_orthogroups_dir, 'OG_Protein_Sequences')
### commented out to avoid unnecessary analysis
if False:
    %run run_zol.py -i $mibig_only_annotated_as7_clusters -o $mibig_zol_results_dir
### it may be better to run this script from the command line:
print('conda activate zol_env\npython %s/run_zol.py -i %s -o %s -as %s' %(publication_scripts_dir, mibig_only_annotated_as7_clusters, mibig_zol_results_dir, all_annotated_as7_clusters))
### Results folder here
print("\n Results can be found here:\n\n %s/Final_Results/Consolidated_Report.xlsx\n\n"
      "%s/Determine_Orthogroups/Orthogroups.tsv\n\n%s/Ortholog_Group_Processing/OG_Protein_Sequences" %(mibig_zol_results_dir, mibig_zol_results_dir, mibig_zol_results_dir))
#
mibig_orthogroupstable_dir = os.path.join(mibig_zol_results_dir, 'Determine_Orthogroups')
mibig_orthogroupscustomtable = os.path.join(mibig_orthogroupstable_dir, 'Orthogroups_custom.tsv')
mibig_orthogroupscustomtable_df = pd.read_csv(mibig_orthogroupscustomtable, sep='\t', nrows=5)
mibig_orthogroupscustomtable_df


# %% [markdown]
## Using Zol to estimate orthogroups from untrimmed complete BGCs

# %% [code]
### now run Zol on untrimmed complete clusters
originaluntrimmed_zol_results_dir = os.path.join(orthology_inf_dir, '1_untrimmed_complete_ozol_results')
### commented out to avoid unnecessary analysis - takes about 50 min
if False:
    %run run_zol.py -i $untrimmed_complete_clusters_dir -o $originaluntrimmed_zol_results_dir
### it may be better to run this script from the command line:
print('conda activate zol_env\npython %s/run_zol.py \\\n-i %s \\\n-o %s \\\n-as %s' %(publication_scripts_dir, untrimmed_complete_clusters_dir, originaluntrimmed_zol_results_dir, all_annotated_as7_clusters))
### Results folder here
print("\n Results can be found here:\n\n %s/Final_Results/Consolidated_Report.xlsx\n\n"
      "%s/Determine_Orthogroups/Orthogroups.tsv\n\n%s/Ortholog_Group_Processing/OG_Protein_Sequences" %(originaluntrimmed_zol_results_dir, originaluntrimmed_zol_results_dir, originaluntrimmed_zol_results_dir))

# %% [markdown]
## Using Zol to estimate orthogroups from trimmed complete BGCs
Note: this only makes sense to do once the trimmed dataset is finilized!!!!!!!!!!!!!!!!

# %% [code]
### now run Zol on trimmed complete clusters
trimmed_zol_results_dir = os.path.join(orthology_inf_dir, '0_trimmed_complete_ozol_results')
trimmed_orthogroups_dir = os.path.join(trimmed_zol_results_dir, 'Ortholog_Group_Processing')
trimmed_ogs_faas_dir = os.path.join(trimmed_orthogroups_dir, 'OG_Protein_Sequences')
### commented out to avoid unnecessary analysis - takes about 50 min
if False:
    %run run_zol.py -i $trimmed_complete_clusters_dir -o $trimmed_zol_results_dir
### it may be better to run this script from the command line:
print('conda activate zol_env\npython %s/run_zol.py \\\n-i %s \\\n-o %s \\\n-as %s' %(publication_scripts_dir, trimmed_complete_clusters_dir, trimmed_zol_results_dir, all_annotated_as7_clusters))
### Results folder here
print("\n Results can be found here:\n\n %s/Final_Results/Consolidated_Report.xlsx\n\n"
      "%s/Determine_Orthogroups/Orthogroups.tsv\n\n%s/Ortholog_Group_Processing/OG_Protein_Sequences" %(trimmed_zol_results_dir, trimmed_zol_results_dir, trimmed_zol_results_dir))
#
trimmed_orthogroupstable_dir = os.path.join(trimmed_zol_results_dir, 'Determine_Orthogroups')
trimmed_orthogroupscustomtable = os.path.join(trimmed_orthogroupstable_dir, 'Orthogroups_custom.tsv')
trimmed_orthogroupscustomtable_df = pd.read_csv(trimmed_orthogroupscustomtable, sep='\t', nrows=5)
trimmed_orthogroupscustomtable_df

# %% [markdown]
## Summarize zol results

# %% [code]
print("Summary of complete trimmed clusters:")
customtablefile = os.path.join(trimmed_zol_results_dir, 'Determine_Orthogroups/Orthogroups_custom.tsv')
zol_summary_table_df = pd.read_csv(customtablefile, sep='\t', nrows=5)
zol_summary_table_df

# %% [markdown]
## Extract NRPS domains and modules - IMAGE HERE IS OLD NEEDS REPLACING!!!!!!!!!!!!!!!!!!!!
Add here logic behind which modules/domains align with which (kind of included in extract_domain_seqs.py, otherwise in notes from 07.09.2023) 

Doesn't this happen before the prepare ogs?

# %% [code]
### mibig dataset
mibig_domain_analysis_overdir = os.path.join(orthology_inf_dir, '1a_untrimmed_mibig_domain_analysis')
mibig_backbone_analysis_dir = os.path.join(mibig_domain_analysis_overdir, '0_backbone_analysis')
mibig_modules_analysis_dir = os.path.join(mibig_domain_analysis_overdir, '1_modules_analysis')
mibig_domain_analysis_dir = os.path.join(mibig_domain_analysis_overdir, '2_domains_analysis')
mibig_nrpsgenes_tablefile = os.path.join(mibig_domain_analysis_overdir, 'nrpsgenes_metadata.csv')
### commented out to avoid unnecessary analysis
if False:
    %run extract_domain_seqs.py -i $mibig_only_annotated_as7_clusters -o $mibig_domain_analysis_overdir
### it may be better to run this script from the command line:
print('conda activate gpas\npython %s/extract_domain_seqs.py \\\n-i %s \\\n-o %s' %(publication_scripts_dir, mibig_only_annotated_as7_clusters, mibig_domain_analysis_overdir))

### showing an example of backbone amino acid conservation based on prediction
from IPython.display import Image, display
listOfImageNames = ['mibig_backbone_example_logo_a.png']
print("\n\nConservation pattern of MIBiG glycopeptide entries:")
for imageName in listOfImageNames:
    display(Image(filename=imageName))

# %% [code]
### complete trimmed dataset        - RUN ONLY WHEN TRIMMED DATASET IS COMPLETE!!!!!!!!!!!!!!!!!!!!!!!!!!!!
domain_analysis_overdir = os.path.join(orthology_inf_dir, '1_trimmed_complete_domain_analysis')
backbone_analysis_dir = os.path.join(domain_analysis_overdir, '0_backbone_analysis')
modules_analysis_dir = os.path.join(domain_analysis_overdir, '1_modules_analysis')
domain_analysis_dir = os.path.join(domain_analysis_overdir, '2_domains_analysis')
nrpsgenes_tablefile = os.path.join(domain_analysis_overdir, 'nrpsgenes_metadata.csv')
### commented out to avoid unnecessary analysis
if False:
    %run extract_domain_seqs.py -i $trimmed_complete_clusters_dir -o $domain_analysis_overdir
### it may be better to run this script from the command line:
print('conda activate gpas\npython %s/extract_domain_seqs.py \\\n-i %s \\\n-o %s' %(publication_scripts_dir, trimmed_complete_clusters_dir, domain_analysis_overdir))

# %% [markdown]
## Remove NRPS genes from OGs and replace by domains and modules
Also rename orthogroup files based on best annotation and change sequence headers to make sure results can be combined later. Any non-single-copy groups here will be filtered into single-copy.

# %% [code]
### make two versions, one where instead of NRPS genes there are domains and one where there are modules
mibig_tidy_groups_dir = os.path.join(orthology_inf_dir, '2a_untrimmed_mibig_tidy_groups')
mibig_modules_filtered_seqs_faas = os.path.join(mibig_tidy_groups_dir, '1_OGs_filtered_with_modules_faas')
mibig_domains_filtered_seqs_faas = os.path.join(mibig_tidy_groups_dir, '2_OGs_filtered_with_domains_faas')
### commented out to avoid unnecessary analysis
if False:
    %run prepare_ogs_for_phylogeny.py -i $mibig_ogs_faas_dir -o $mibig_tidy_groups_dir -nrps mibig_nrpsgenes_tablefile -ot mibig_orthogroupscustomtable -m mibig_modules_analysis_dir -d mibig_domain_analysis_dir
### it may be better to run this script from the command line:
print('conda activate gpas\npython %s/prepare_ogs_for_phylogeny.py \\\n-i %s \\\n-o %s \\\n-nrps %s \\\n-ot %s \\\n-m %s \\\n-d %s' %(publication_scripts_dir, mibig_ogs_faas_dir, mibig_tidy_groups_dir, mibig_nrpsgenes_tablefile, mibig_orthogroupscustomtable, mibig_modules_analysis_dir, mibig_domain_analysis_dir))
### show what we did
mibig_filtered_ogs_mods_table = os.path.join(mibig_tidy_groups_dir, '1_OGs_filtered_with_modules_metadata.tsv')
mibig_filtered_ogs_doms_table = os.path.join(mibig_tidy_groups_dir, '2_OGs_filtered_with_domains_metadata.tsv')
mibig_ogs_mods_df = pd.read_csv(mibig_filtered_ogs_mods_table, sep='\t')
mibig_ogs_doms_df = pd.read_csv(mibig_filtered_ogs_doms_table, sep='\t')
print("\nModules dataset includes %s groups.\nDomains dataset includes %s groups." %(len(mibig_ogs_mods_df), len(mibig_ogs_doms_df)))
mibig_ogs_doms_df.iloc[:5]

# %% [code]
### - RUN ONLY WHEN TRIMMED DATASET IS COMPLETE!!!!!!!!!!!!!!!!!!!!!!!!!!!!
### make two versions, one where instead of NRPS genes there are domains and one where there are modules
tidy_groups_dir = os.path.join(orthology_inf_dir, '2_trimmed_complete_tidy_groups')
modules_filtered_seqs_faas = os.path.join(tidy_groups_dir, '1_OGs_filtered_with_modules_faas')
domains_filtered_seqs_faas = os.path.join(tidy_groups_dir, '2_OGs_filtered_with_domains_faas')
### commented out to avoid unnecessary analysis
if False:
    %run prepare_ogs_for_phylogeny.py -i $trimmed_ogs_faas_dir -o $tidy_groups_dir -nrps nrpsgenes_tablefile -ot trimmed_orthogroupscustomtable -m modules_analysis_dir -d domain_analysis_dir
### it may be better to run this script from the command line:
print('conda activate gpas\npython %s/prepare_ogs_for_phylogeny.py \\\n-i %s \\\n-o %s \\\n-nrps %s \\\n-ot %s \\\n-m %s \\\n-d %s' %(publication_scripts_dir, trimmed_ogs_faas_dir, tidy_groups_dir, nrpsgenes_tablefile, trimmed_orthogroupscustomtable, modules_analysis_dir, domain_analysis_dir))
### show what we did
filtered_ogs_mods_table = os.path.join(tidy_groups_dir, '1_OGs_filtered_with_modules_metadata.tsv')
filtered_ogs_doms_table = os.path.join(tidy_groups_dir, '2_OGs_filtered_with_domains_metadata.tsv')
ogs_mods_df = pd.read_csv(filtered_ogs_mods_table, sep='\t')
ogs_doms_df = pd.read_csv(filtered_ogs_doms_table, sep='\t')
print("\nModules dataset includes %s groups.\nDomains dataset includes %s groups." %(len(ogs_mods_df), len(ogs_doms_df)))
ogs_doms_df.iloc[:5]

# %% [markdown]
# Step 5: Basic phylogeny
This includes the following:
- ### MSAs with mafft
mafft v7.520 (2023/Mar/22)
- ### Trimming MSAs with trimAl
trimAl v1.4.rev15 build[2013-12-17]
- ### Model testing untrimmed MSAs
IQ-TREE multicore version 2.2.5 COVID-edition for Linux 64-bit built Sep 15 2023
Standard Protein models considered: Dayhoff,DCMut,JTT,JTTDCMut,LG,Poisson,PMB,Q.pfam,VT,WAG
Mixed Protein models considered: C10,C20,C30,C40,C50,C60,EX2,EX3,EHO,UL2,UL3,EX_EHO,LG4M,LG4X,CF4
- ### Drawing bootstrapped ML tres with iqtree
IQ-TREE multicore version 2.2.5 COVID-edition for Linux 64-bit built Sep 15 2023
- ### Checking bootstrapping values?????? - maybe discuss with Huson???????
They are computed for all trees. For the concatenated phylogeny they are not relevant, since the MSAs are concatenated, not the trees. For the congruence analysis they could be relevant so there they can be manually checked since the number of trees is specific. For the gene network either they are not considered or they are automatically taken into account for the split weights.

# %% [markdown]
## MIBiG dataset

# %% [code]
def create_folder(path):
    if not os.path.exists(path):
        os.mkdir(path)
        print("Created folder: %s" %(path))
mibig_basicphylogeny_dir = os.path.join(basicphylogenetics_dir, '2a_untrimmed_mibig_basic_phylogeny')
create_folder(mibig_basicphylogeny_dir)
mibig_modules_basicphylogeny_dir = os.path.join(mibig_basicphylogeny_dir, '1_OGs_with_modules_phylogeny')
mibig_modules_modelT_summary_file = os.path.join(mibig_modules_basicphylogeny_dir, 'modeltesting_summary.csv')
mibig_modules_mafftMSAs = os.path.join(mibig_modules_basicphylogeny_dir, '1_mafft_MSAs')
mibig_modules_trimalMSAs = os.path.join(mibig_modules_basicphylogeny_dir, '2_trimal_MSAs')
mibig_modules_maffttrees = os.path.join(mibig_modules_basicphylogeny_dir, '4_mafft_iqtree_ML_trees')
mibig_modules_trimaltrees = os.path.join(mibig_modules_basicphylogeny_dir, '5_trimal_iqtree_ML_trees')
mibig_domains_basicphylogeny_dir = os.path.join(mibig_basicphylogeny_dir, '2_OGs_with_domains_phylogeny')
### commented out to avoid unnecessary analysis. modules one took 20-134 min until model testing, min in total. domains one took min.
if False:
    %run run_basic_phylogeny.py -i $mibig_modules_filtered_seqs_faas -o $mibig_modules_basicphylogeny_dir 
    %run run_basic_phylogeny.py -i $mibig_domains_filtered_seqs_faas -o $mibig_domains_basicphylogeny_dir 
### it may be better to run this script from the command line:
print('conda activate gpas\npython %s/run_basic_phylogeny.py \\\n-i %s \\\n-o %s\n\n' %(publication_scripts_dir, mibig_modules_filtered_seqs_faas, mibig_modules_basicphylogeny_dir))
print('conda activate gpas\npython %s/run_basic_phylogeny.py \\\n-i %s \\\n-o %s' %(publication_scripts_dir, mibig_domains_filtered_seqs_faas, mibig_domains_basicphylogeny_dir))
### show preview of results
mibig_modelT_modules_summary_table_df = pd.read_csv(mibig_modules_modelT_summary_file, nrows=5)
mibig_modelT_modules_summary_table_df

# %% [markdown]
## Complete trimmed BGCs dataset

# %% [code]
### - RUN ONLY WHEN TRIMMED DATASET IS COMPLETE!!!!!!!!!!!!!!!!!!!!!!!!!!!!
def create_folder(path):
    if not os.path.exists(path):
        os.mkdir(path)
        print("Created folder: %s" %(path))
basicphylogeny_dir = os.path.join(basicphylogenetics_dir, '2_trimmed_complete_basic_phylogeny')
create_folder(basicphylogeny_dir)
modules_basicphylogeny_dir = os.path.join(basicphylogeny_dir, '1_OGs_with_modules_phylogeny')
modules_modelT_summary_file = os.path.join(modules_basicphylogeny_dir, 'modeltesting_summary.csv')
modules_mafftMSAs = os.path.join(modules_basicphylogeny_dir, '1_mafft_MSAs')
modules_trimalMSAs = os.path.join(modules_basicphylogeny_dir, '2_trimal_MSAs')
modules_maffttrees = os.path.join(modules_basicphylogeny_dir, '4_mafft_iqtree_ML_trees')
modules_trimaltrees = os.path.join(modules_basicphylogeny_dir, '5_trimal_iqtree_ML_trees')
domains_basicphylogeny_dir = os.path.join(basicphylogeny_dir, '2_OGs_with_domains_phylogeny')
domains_modelT_summary_file = os.path.join(domains_basicphylogeny_dir, 'modeltesting_summary.csv')
domains_mafftMSAs = os.path.join(domains_basicphylogeny_dir, '1_mafft_MSAs')
domains_trimalMSAs = os.path.join(domains_basicphylogeny_dir, '2_trimal_MSAs')
domains_maffttrees = os.path.join(domains_basicphylogeny_dir, '4_mafft_iqtree_ML_trees')
domains_trimaltrees = os.path.join(domains_basicphylogeny_dir, '5_trimal_iqtree_ML_trees')
### commented out to avoid unnecessary analysis. modules one took 120+ min. domains one took min.
if False:
    %run run_basic_phylogeny.py -i $modules_filtered_seqs_faas -o $modules_basicphylogeny_dir 
    %run run_basic_phylogeny.py -i $domains_filtered_seqs_faas -o $domains_basicphylogeny_dir 
### it may be better to run this script from the command line:
print('conda activate gpas\npython %s/run_basic_phylogeny.py \\\n-i %s \\\n-o %s\n\n' %(publication_scripts_dir, modules_filtered_seqs_faas, modules_basicphylogeny_dir))
print('conda activate gpas\npython %s/run_basic_phylogeny.py \\\n-i %s \\\n-o %s' %(publication_scripts_dir, domains_filtered_seqs_faas, domains_basicphylogeny_dir))

print('\nNote: If sequences are fewer than 4 after removal of identical sequences, tree-building is aborted for that file (e.g. OG_153 is missing from the trimmed MSA trees because of that).')
### show preview of results
modelT_modules_summary_table_df = pd.read_csv(modules_modelT_summary_file, nrows=5)
modelT_modules_summary_table_df
### show preview of results
modelT_domains_summary_table_df = pd.read_csv(domains_modelT_summary_file, nrows=5)
modelT_domains_summary_table_df

# %% [markdown]
Checking bootstrap values?

# %% [markdown]
# Step 6: Concatenated representative phylogeny
This includes the following:
- ### Congruence analysis
-- Candidates include groups that are found in at least 90% of the clusters. Duplications are acceptable in maximum 5% of the clusters in a group, but they are filtered out before inclusion.
-- 100 random subsampled (70%) concatenated ML trees are built with FastTree v2.1.11 using a WAG + G model.
-- A consensus tree with split support values is calculated using dendropy's (v4.6.1) sumtrees function.
After identifying candidates, manually/automatically check bootstrap values???????????????????????????????????
-- The candidates' congruence value (normalised compatible split length, as described by Parks et al, 2017) was calculated with the genetreetk supported_splits function (github repo, accessed on 20.09.2023: https://github.com/dparks1134/GeneTreeTk)
- ### Concatenated phylogeny
Built from the MSAs of the candidates who passed the congruence check (assuming congruence threshold to be defined by 3rd module or 3rd A-domain) with model testing????? or with CAT + G as per Wang et al, 2008???????????????

- ### MAD rooting
The concatenated phylogenies are then rooted based on MAD (http://dx.doi.org/10.1038/s41559-017-0193)


# %% [markdown]
## MIBiG dataset

# %% [code]
def create_folder(path):
    if not os.path.exists(path):
        os.mkdir(path)
        print("Created folder: %s" %(path))
mibig_concatphylogeny_dir = os.path.join(concatphylogeny_dir, '2a_untrimmed_mibig_concat_phylogeny')
create_folder(mibig_concatphylogeny_dir)
### MIBIG dataset
mibig_modules_concatphyl_dir = os.path.join(mibig_concatphylogeny_dir, '1_OGs_with_modules_concatphylogeny')
mibig_domains_concatphyl_dir = os.path.join(mibig_concatphylogeny_dir, '2_OGs_with_domains_concatphylogeny')
mibig_modules_congruence_summarytable = os.path.join(mibig_modules_concatphyl_dir, '3_congruence_check_summary.csv')
### commented out to avoid unnecessary analysis. modules one took 120+ min. domains one took min.
if False:
    %run make_concat_phylogeny.py -i $mibig_modules_basicphylogeny_dir -o $mibig_modules_concatphyl_dir -m $mibig_filtered_ogs_mods_table 
    %run make_concat_phylogeny.py -i $mibig_domains_basicphylogeny_dir -o $mibig_domains_concatphyl_dir -m $mibig_filtered_ogs_doms_table 
### it may be better to run this script from the command line:
print('\n\nNOTE: this script can only be run on the command line!!!!!!!!!!!\n')
print('conda activate gpas\npython %s/make_concat_phylogeny.py \\\n-i %s \\\n-o %s \\\n -m %s\n\n' %(publication_scripts_dir, mibig_modules_basicphylogeny_dir, mibig_modules_concatphyl_dir, mibig_filtered_ogs_mods_table))
print('conda activate gpas\npython %s/make_concat_phylogeny.py \\\n-i %s \\\n-o %s \\\n-m %s' %(publication_scripts_dir, mibig_domains_basicphylogeny_dir, mibig_domains_concatphyl_dir, mibig_filtered_ogs_doms_table))
print('\n\nNOTE: these scripts can only be run on the command line!!!!!!!!!!!')
### show preview of results
mibig_modules_congruence_summary_table_df = pd.read_csv(mibig_modules_congruence_summarytable, nrows=5)
mibig_modules_congruence_summary_table_df

madpath = '/home/athina/Documents/programs/mad/mad'
### Now to MAD rooting
mibig_unrooted_modules_concat_tree = os.path.join(mibig_modules_concatphyl_dir, '5_concatenated_phylogeny.treefile')
mibig_MADrooted_modules_concat_tree = os.path.join(mibig_modules_concatphyl_dir, '5_concatenated_phylogeny.treefile.rooted')
if False:
    %run $madpath $mibig_unrooted_modules_concat_tree -t
print("\n%s %s -t" %(madpath, mibig_unrooted_modules_concat_tree))
### Now to MAD rooting
mibig_unrooted_domains_concat_tree = os.path.join(mibig_domains_concatphyl_dir, '5_concatenated_phylogeny.treefile')
mibig_MADrooted_domains_concat_tree = os.path.join(mibig_domains_concatphyl_dir, '5_concatenated_phylogeny.treefile.rooted')
if False:
    %run $madpath $mibig_unrooted_domains_concat_tree -t
print("\n\n%s %s -t" %(madpath, mibig_unrooted_domains_concat_tree))
print('\nRooted trees based on MAD')

# %% [markdown]
## Complete trimmed BGCs dataset

# %% [code]
##CAT + Γ model appropriate for large concatenated data sets (Wang et al, 2008)
def create_folder(path):
    if not os.path.exists(path):
        os.mkdir(path)
        print("Created folder: %s" %(path))
completetrimmedconcatphylogeny_dir = os.path.join(concatphylogeny_dir, '2_trimmed_complete_concat_phylogeny')
create_folder(completetrimmedconcatphylogeny_dir)
### 
modules_concatphyl_dir = os.path.join(completetrimmedconcatphylogeny_dir, '1_OGs_with_modules_concatphylogeny')
domains_concatphyl_dir = os.path.join(completetrimmedconcatphylogeny_dir, '2_OGs_with_domains_concatphylogeny')
modules_congruence_summarytable = os.path.join(modules_concatphyl_dir, '3_congruence_check_summary.csv')
### commented out to avoid unnecessary analysis. modules one took 120+ min. domains one took min.
if False:
    %run make_concat_phylogeny.py -i $modules_basicphylogeny_dir -o $modules_concatphyl_dir -m $filtered_ogs_mods_table 
    %run make_concat_phylogeny.py -i $domains_basicphylogeny_dir -o $domains_concatphyl_dir -m $filtered_ogs_doms_table 
### it may be better to run this script from the command line:
print('\n\nNOTE: this script can only be run on the command line!!!!!!!!!!!\n')
print('conda activate gpas\npython %s/make_concat_phylogeny.py \\\n-i %s \\\n-o %s \\\n -m %s\n\n' %(publication_scripts_dir, modules_basicphylogeny_dir, modules_concatphyl_dir, filtered_ogs_mods_table))
print('conda activate gpas\npython %s/make_concat_phylogeny.py \\\n-i %s \\\n-o %s \\\n-m %s' %(publication_scripts_dir, domains_basicphylogeny_dir, domains_concatphyl_dir, filtered_ogs_doms_table))
print('\n\nNOTE: this script can only be run on the command line!!!!!!!!!!!')
### show preview of results
modules_congruence_summary_table_df = pd.read_csv(modules_congruence_summarytable, nrows=5)
modules_congruence_summary_table_df


#print('\nAt this point, check for correct trimming')

# %% [markdown]
## MAD rooting

# %% [code]
madpath = '/home/athina/Documents/programs/mad/mad'
### Now to MAD rooting - imported also in the concat phylogeny script now
unrooted_modules_concat_tree = os.path.join(modules_concatphyl_dir, '5_concatenated_phylogeny.treefile')
MADrooted_modules_concat_tree = os.path.join(modules_concatphyl_dir, '5_concatenated_phylogeny.treefile.rooted')
if False:
    %run $madpath $unrooted_modules_concat_tree -t
print("\n%s %s -t" %(madpath, unrooted_modules_concat_tree))
### Now to MAD rooting
unrooted_domains_concat_tree = os.path.join(domains_concatphyl_dir, '5_concatenated_phylogeny.treefile')
MADrooted_domains_concat_tree = os.path.join(domains_concatphyl_dir, '5_concatenated_phylogeny.treefile.rooted')
if False:
    %run $madpath $unrooted_domains_concat_tree -t
print("\n\n%s %s -t" %(madpath, unrooted_domains_concat_tree))
print('\nRooted trees based on MAD')

# %% [markdown]
# Step 7: super network
Using SplitsTree CE version 6.1.14-beta:
- File > Import > Trees from multiple files
- Navigate to the step 5 folder 4_mafft_iqtree_ML_trees containing untrimmed mafft iqtree ML trees and select all of them
- Network > Super Network
- Go to the "Workflow" tab, right click on the "Splits" box (after "Super Network") and select "Add Algorithm" > "Splits Filter"
- Specify the "Filter Algorithm" as "GreedyWeaklyCompatible" and click the little green arrow to "Execute Algorithm"
- Optinally check the box next to "Recompute Cycle", to get a different angle of the network

# %% [code]
### showing an example of backbone amino acid conservation based on prediction
#from IPython.display import Image, display
#listOfImageNames = ['mibig_modules_supernetwork.png']
#print("\n\nMiBIG module-based super network with GreedyWeaklyCompatible splits filter:")
#for imageName in listOfImageNames:
#    display(Image(filename=imageName))

# %% [markdown]
## Visualizing super network

# %% [code]


# %% [markdown]
# NEXT SECTION HERE

# %% [markdown]
#### maybe eventually add log files to scripts?

# %% [code]
```
