## Supplementary Data 5 for "Structural and evolutionary analyses support reclassification of glycopeptide antibiotics as xyclopeptides": detect_xdomain_regions.py

```
#!/usr/bin/python3.6
# Created by Athina Gavriilidou at 13.07.23 11:54 #
# Imports #
import argparse
import os
import csv
from pprint import pprint
from Bio import SeqIO
import pandas as pd
import re
import math
import string
import warnings
import gbk_to_faa

from pandas.core.common import SettingWithCopyWarning

warnings.simplefilter(action='ignore', category=FutureWarning)
warnings.simplefilter(action='ignore', category=SettingWithCopyWarning)


# Functions #
def list_folders(directory):
    folders_list = [os.path.join(directory, o) for o in os.listdir(directory)
                    if os.path.isdir(os.path.join(directory,o))]
    return folders_list

def csv_to_dict(csvfile, key="", value=""):
    thedict ={}
    with open(csvfile, 'r') as infile:
        reader = csv.reader(infile)
        for headerline in reader:
            key_id = headerline.index(key)
            value_id = headerline.index(value)
            break
        for line in reader:
            if line[key_id] not in thedict.keys():
                thedict[line[key_id]] = line[value_id]
            else:
                thedict[line[key_id]] = "%s|%s" %(thedict[line[key_id]], line[value_id])
    return thedict

def locate_region_files(targetdir):
    files_list = [os.path.join(targetdir, o) for o in os.listdir(targetdir)
                  if os.path.isfile(os.path.join(targetdir, o)) and 'region' in o and o.endswith('.gbk')]
    if "QPWMP018AI_assembly_contig_304_region001" in targetdir:
        files_list = [x for x in files_list if not "QPWMP018AI_assembly_contig_304_region001" in os.path.basename(x)]
    return files_list

def grep_file(thefile, thestring):
    with open(thefile, 'r') as f:
        for line in f.readlines():
            if thestring in line:
                return True
    return False

'''for copying files. Make everything=True and then everything from the source will be copied to the destination'''
def execute_cp(source_fileordir, destinationfileordir, everything=False):
    if everything:
        cp_command = 'cp %s/* %s/.' %(source_fileordir,destinationfileordir)
    else:
        cp_command = 'cp \'%s\' %s' %(source_fileordir,destinationfileordir)
    try:
        os.system(cp_command)
    except:
        print("Warning. There was a problem with copying %s to %s" %(source_fileordir, destinationfileordir))

def create_folder(path):
    if not os.path.exists(path):
        os.mkdir(path)
        print("Created folder: %s" %(path))

def gbk_to_fna(gbkfile, fnafile):
    sequences_to_write = []
    with open(gbkfile) as input_handle, open(
            fnafile, "w"
    ) as output_handle:
        sequences = SeqIO.parse(input_handle, "genbank")
        #print(sequences)
        for seq_record in sequences:
            #print(seq_record)
            comments = seq_record.annotations['structured_comment']
            try:
                as_comments = comments['antiSMASH-Data']
                thestart = as_comments['Orig. start']
                theend = as_comments['Orig. end']
            except:
                thestart = '_'
                theend = '_'
                print("Could not infer start and end positions ")
            ### add new coordinates to fasta file
            seq_record.description = "%s_antiSMASH7_region_origstart_%s_origend_%s"\
                                     %(seq_record.description, thestart, theend)
            sequences_to_write.append(seq_record)
        count = SeqIO.write(sequences_to_write, output_handle, "fasta")
    #print("Converted %i records" % count)
    #exit()

def infer_bgcid(fullpathname, metadata_df, bgcid_to_xdomfile):#, species_to_char_dict,speciesname_to_char_dict):
    #if "GCF_000719255" in filename or "GCF_000718635" in filename:
    #    print(filename)
    filename = os.path.basename(fullpathname)
    foldername = os.path.dirname(fullpathname)
    #print(filename)
    ### culp dataset
    if filename[:2].isnumeric():
        source_df = metadata_df.loc[metadata_df['source'] == 'Culp']
        bgc_dict = source_df.loc[source_df['Database_ID'] == str(int(filename[:2]))].to_dict(orient='list')
        #print(bgc_dict)
        bgcid = bgc_dict['ID'][0]
        #print('culp',bgcid)
    ### MIBIG
    elif filename.startswith("BGC"):
        source_df = metadata_df.loc[metadata_df['source'] == 'MIBiG']
        bgc_dict = source_df.loc[source_df['Database_ID'] == filename.split("_")[0]].to_dict(orient='list')
        bgcid = bgc_dict['ID'][0]
    ### Genbank
    elif filename.startswith("GCA_"):
        source_df = metadata_df.loc[metadata_df['source'] == 'Genbank']
        indices = [m.start() for m in re.finditer('_', filename)]
        #print(filename[:indices[1]])
        bgc_dict = source_df.loc[source_df['Database_ID'].str.contains(filename[:indices[1]])].to_dict(orient='list')
        bgcid = bgc_dict['ID'][0]
    ### RefSeq
    elif filename.startswith("GCF_"):
        source_df = metadata_df.loc[metadata_df['source'] == 'RefSeq']
        indices = [m.start() for m in re.finditer('_', filename)]
        # print(filename[:indices[1]])
        bgc_dict = source_df.loc[source_df['Database_ID'].str.contains(filename[:indices[1]])].to_dict(orient='list')
        #if "GCF_000719255" in filename or "GCF_000718635" in filename:
            #print(filename)
            #print(bgc_dict)
        bgcid = bgc_dict['ID'][0]
        #print(bgc_dict)
    ### MGnify
    elif filename.startswith("MG"):
        source_df = metadata_df.loc[metadata_df['source'] == 'MGnify']
        bgc_dict = source_df.loc[source_df['Database_ID'] == filename.split("_")[0]].to_dict(orient='list')
        bgcid = bgc_dict['ID'][0]
    ### ShrikantOldTelSeq
    elif filename.startswith("QP"):
        if "assembly_contig_304_region001" in filename:
            source_df = metadata_df.loc[metadata_df['source'] == 'ShrikantOldData']
        else:
            source_df = metadata_df.loc[metadata_df['source'] == 'ShrikantOldTelSeq']
        bgc_dict = source_df.loc[source_df['Database_ID'] == filename.split("_")[0]].to_dict(orient='list')
        bgcid = bgc_dict['ID'][0]
    ### Stammsamlung2022
    elif filename.startswith("S00"):
        source_df = metadata_df.loc[metadata_df['source'] == 'Stammsamlung2022']
        bgc_dict = source_df.loc[source_df['Database_ID'] == filename.split("_")[0]].to_dict(orient='list')
        bgc_list = bgc_dict['ID']
        #bgcid = bgc_dict['ID'][0]
        i = 0
        bgcid = bgc_list[i]
        while bgcid in bgcid_to_xdomfile.keys():
            i+=1
            bgcid = bgc_list[i]
        #print(bgcid)
    ### LocalSeq - hardcoded for ease
    elif filename.startswith("Sva"):
        source_df = metadata_df.loc[metadata_df['source'] == 'LocalSeq']
        bgc_dict = source_df.loc[source_df['Database_ID'] == 'Sva'].to_dict(orient='list')
        bgcid = bgc_dict['ID'][0]
        #if 'region003' in filename:
        #    bgcid = 'SvN_1'
        #elif 'region022' in filename:
        #    bgcid = 'SvN_2'

    '''
    ### now if bgcid is nan, assign one
    genus_to_char_dict = {'Streptomyces':'St', 'Amycolatopsis':'Am', 'Microbispora':'Mb', 'uncultured':'U',
                          'Actinomadura': 'Ad', 'Actinobacteria':'Ac', 'Nonomuraea':'Nn', 'Nocardia':'Nc',
                          'Actinoplanes':'Ap','Umezawaea':'UM','Kitasatospora':'Kt','Kibdelosporangium':'Kb',
                          'Micromonosporaceae':'Mm', 'Kibdellosporangium':'Kb','Actinokineospora':'Ak',
                          'Sinosporangium':'Si','Kutzneria':'Ku','Uncultured':'U'}
    #species_to_char_dict = {}
    if str(bgcid).startswith('nan'):
        #print(bgc_dict)
        strain_str = bgc_dict['strain'][0]
        try:
            genus = strain_str.split(' ')[0]
            species = strain_str[len(genus) + 1:]
            strain = strain_str[len(genus) + len(species)+2:]
            if species.startswith('sp.'):
                is_sp = True
                species = species[4:]
            else:
                is_sp = False
            species = species.split(" ")[0]
        ### exception occurs when no strain is known
        except:
            genus = 'uncultured'
            is_sp = False
            species = bgc_dict['source'][0]
            strain = bgc_dict['Database_ID'][0]
        try:
            prefix = genus_to_char_dict[genus]
        ### exception occurs where the genus has no corresponding character hardcoded into the dictionary
        except:
            print("\nNew genus detected: %s. Update genus dictionary! \nAborting...\n" %(genus))
            pprint(genus_to_char_dict)
            exit()
        if genus not in species_to_char_dict:
            species_to_char_dict[genus] = {}
        if is_sp:
            prefix = prefix + 's'
        ### get species info in the name
        #speciesfirst = species.split(' ')[0]
        if species not in speciesname_to_char_dict.keys():
            midpart = species[:1].upper()
        else:
            midpart = speciesname_to_char_dict[species]
        ### check that a prefix of the name has not been generated due to duplications:
        for i in species_to_char_dict[genus].keys():
            #print([i])
            if midpart in i and len(midpart) != len(i):
                print(strain_str)
                midpart = midpart[:1] + species[1:len(i)]
        if midpart not in species_to_char_dict[genus].keys():
                species_to_char_dict[genus][midpart] = species
        ### if a duplicate name occured, change both
        elif species_to_char_dict[genus][midpart] != species:
            #print(genus, species, strain, species_to_char_dict[genus][midpart])
            speciesfirstprevious = species_to_char_dict[genus][midpart]
            midpartprevious = midpart + speciesfirstprevious[len(midpart)]
            species_to_char_dict[genus][midpartprevious] = speciesfirstprevious
            speciesname_to_char_dict[speciesfirstprevious] = midpartprevious
            del species_to_char_dict[genus][midpart]
            midpart = midpart + species[len(midpart)]
            species_to_char_dict[genus][midpart] = species
            speciesname_to_char_dict[species] = midpart
            #pprint(species_to_char_dict[genus])
        #if midpart == 'Vn':
        #    print(genus, speciesfirst, species_to_char_dict[genus][midpart])
        #    exit()
        ### assemble parts of the name
        bgcid = "%s%s" %(prefix, midpart)
        print("bgcid:",bgcid)
    pprint(species_to_char_dict['Streptomyces'])
    '''
    #if len(bgc_dict['ID']) > 1:
    #    print(filename, bgcid, foldername, bgc_dict)
    #print(bgcid_to_xdomfile.keys())
    ### extra check for unknown cases of two clusters in a genome
    '''
    if bgcid in bgcid_to_xdomfile.keys():
        #print(bgcid)
        bgc1 = "%s_1" %(bgcid)
        bgc2 = "%s_2" %(bgcid)
        bgcid_to_xdomfile[bgc1] = bgcid_to_xdomfile[bgcid]
        del bgcid_to_xdomfile[bgcid]
        bgcid = bgc2
        #pprint(bgcid_to_xdomfile)
        #print(filename)
        #exit()
    '''

    ### hardcoded special case where in a metagenomic sample we found a third hit
    #if "S009" in filename and 'contig_4262' in filename:
    #    bgcid = "US91_3"
    #pprint(bgcid_to_xdomfile)
    return bgcid, bgcid_to_xdomfile#, species_to_char_dict, speciesname_to_char_dict

def extract_info_from_gbk(gbkfile):
    #print("Extracting from %s" %(gbkfile))
    contig_edge = ''
    with open(gbkfile) as input_handle:
        sequences = SeqIO.parse(input_handle, "genbank")
        for seq_record in sequences:
            #print(seq_record)
            clusterstart =seq_record.annotations['structured_comment']['antiSMASH-Data']['Orig. start']
            clusterend =seq_record.annotations['structured_comment']['antiSMASH-Data']['Orig. end']
            bgc_locus = seq_record.id
            for seq_feature in seq_record.features:
                if seq_feature.type == "protocluster":
                    thiscontig_edge = seq_feature.qualifiers['contig_edge'][0]
                    if contig_edge == '' or thiscontig_edge == 'True':
                        contig_edge = thiscontig_edge
                if seq_feature.type == "cand_cluster":
                    thiscontig_edge = seq_feature.qualifiers['contig_edge'][0]
                    if contig_edge == '' or thiscontig_edge == 'True':
                        contig_edge = thiscontig_edge
                if seq_feature.type == "CDS":
                    #if 'BGC0000289' in gbkfile:
                    #    print(seq_feature)
                    ### to cover weird cases where no locus tag
                    try:
                        startgene = seq_feature.qualifiers['locus_tag'][0]
                    except:
                        startgene = seq_feature.qualifiers['protein_id'][0]
                    break
            for seq_feature in seq_record.features:
                if seq_feature.type == "protocluster":
                    thiscontig_edge = seq_feature.qualifiers['contig_edge'][0]
                    if contig_edge == '' or thiscontig_edge == 'True':
                        contig_edge = thiscontig_edge
                if seq_feature.type == "cand_cluster":
                    thiscontig_edge = seq_feature.qualifiers['contig_edge'][0]
                    if contig_edge == '' or thiscontig_edge == 'True':
                        contig_edge = thiscontig_edge
                if seq_feature.type == "CDS":
                    try:
                        endgene = seq_feature.qualifiers['locus_tag'][0]
                    except:
                        endgene = seq_feature.qualifiers['protein_id'][0]
    complete = ''
    return bgc_locus, startgene, endgene, clusterstart, clusterend, contig_edge, complete

'''given a directory, lists all files, with a certain suffix'''
def list_files(directory, suffix= "", suppress=False):
    the_files = []
    for file in os.listdir(directory):
        if file.endswith(suffix):
            the_files.append(os.path.join(directory, file))
    if not suppress:
        print("Found %s %s files" %(len(the_files), suffix))
    the_files.sort()
    return the_files

def fix_id(theid, index, fullname, list_of_ids):
    while theid in list_of_ids:
        if index == len(fullname):
            break
        theid = theid + fullname[index]
        index+=1
    while theid in list_of_ids:
        index -= 1
        #print(fullname)
        if index == -1:
            break
        theid = theid + fullname[index]
    return theid

def special_fix_id(theid, index, fullname, list_of_ids, suffixcount):
    while theid in list_of_ids:
        if index == len(fullname):
            break
        theid = theid[:-len(suffixcount)] + fullname[index] + theid[-len(suffixcount):]
        index+=1
    while theid in list_of_ids:
        index -= 1
        #print(fullname)
        if index == -1:
            break
        theid = theid[:-len(suffixcount)] + fullname[index] + theid[-len(suffixcount):]
    return theid

'''makes a list of bgcids based on strain and database ID'''
def generate_bgcids(metadata_df, strain_count_dict={}, bgcids_list=[], strain_to_bgcid_dict = {}, species_list = [],
                    species_char_dict = {'sp.':'s'}, strain_list = [], strain_char_dict = {},
                    fullspecies_count_dict = {}):
    genus_to_char_dict = {'Streptomyces': 'St', 'Amycolatopsis': 'Am', 'Microbispora': 'Mb', 'uncultured': 'U',
                          'Actinomadura': 'Ad', 'Actinobacteria': 'Ac', 'Nonomuraea': 'Nn', 'Nocardia': 'Nc',
                          'Actinoplanes': 'Ap', 'Umezawaea': 'UM', 'Kitasatospora': 'Kt', 'Kibdelosporangium': 'Kb',
                          'Micromonosporaceae': 'Mm', 'Kibdellosporangium': 'Kb', 'Actinokineospora': 'Ak',
                          'Sinosporangium': 'Si', 'Kutzneria': 'Ku', 'Uncultured': 'U', math.nan:'U', 'Frankia': 'F'}
    print("\nUsing genus naming conventions as follows:")
    pprint(genus_to_char_dict)
    special_cases_dict = {'15':'Uc37','16':'Uc878','17':'Uc915','21':'Ue15','22':'Ue16','24':'Ue26','41':'UsB',
                          '36':'UsD','contig_304_114':'UQ2','S001':'US1','S003':'US3','S005':'US5','S006':'US6',
                          'S008':'US8','GAEHKOJN_162442':'US91','QPWMP018AI.assembly|ctg1442_135':'UQ1',
                          'GAEHKOJN_183134':'US92','GAEHKOJN_145340':'US93', 'MGYA00375203':'Um',
                          'Sva_producer_05333':'StVrN_B','Sva_producer_00486':'StVrN_A'}
    print('\nSpecial cases:')
    pprint(special_cases_dict)
    print('')
    meta_dict = metadata_df[metadata_df['ID'].isna()].to_dict(orient='list')
    # bad petrformance but whatevs, data is still small:
    for item in meta_dict['strain']:
        ### special case for nan
        if type(item) is float:
            continue
        genus = item.split(' ')[0]
        ### identical strains are saved in a dict
        if item not in strain_count_dict.keys():
            if meta_dict['strain'].count(item) > 1:
                strain_count_dict[item] = meta_dict['strain'].count(item)
        try:
            ### ignore uncultured
            if 'ncultured' in item:
                continue
            species = item.split(' ')[1]
            species_list.append(species)
        ### ignore nan
        except:
            continue
        fullspecies = "%s %s" %(genus, species)
        if not item in strain_count_dict:
            if fullspecies not in fullspecies_count_dict.keys():
                fullspecies_count_dict[fullspecies] = 1
            else:
                fullspecies_count_dict[fullspecies] = fullspecies_count_dict[fullspecies] + 1
        strain = item[len(genus)+len(species)+2:]
        if 'subsp' in strain:
            subspecies = strain.split(' ')[-1]
            strain = strain.split(subspecies)[-1]
        ### if there is a strain, deal with that too
        if len(strain) > 0:
            strain_list.append(strain)
    species_list = list(set(species_list))
    species_list.sort()
    strain_list = list(set(strain_list))
    strain_list.sort()
    #print(strain_count_dict)
    #print(strain_list)
    ### make sure there are different chars for all species
    for species in species_list:
        if species not in species_char_dict.keys():
            index = 1
            char = species[:index]
            templist = [x[:index] for x in species_list if not x is species]
            while char in templist:
                #print(char)#, templist)
                index+=1
                char = species[:index]
                templist = [x[:index] for x in species_list if not x is species]
            if len(char) > 1:
                char = char[0].upper() + char[-1]
            else:
                char = char.upper()
            char = fix_id(char, index, species, species_char_dict.values())
            species_char_dict[species] = char
    #pprint(species_char_dict)
    ### make sure there are different chars for all species
    for strain in strain_list:
        if strain not in strain_char_dict.keys():
            index = 1
            char = strain[:index]
            templist = [x[:index] for x in strain_list if not x is strain]
            while char in templist:
                # print(char)#, templist)
                index += 1
                char = strain[:index]
                templist = [x[:index] for x in strain_list if not x is strain]
            if len(char) > 1:
                char = char[0].upper() + char[-1]
            else:
                char = char.upper()
            ### taking care of duplicates
            char = fix_id(char, index, strain, strain_char_dict.values())
            strain_char_dict[strain] = char
    #pprint(strain_char_dict)
    #pprint(strain_count_dict)
    #for strainstring in meta_dict['strain']:
    for idx, strainstring in metadata_df['strain'].iteritems():
        #print(idx)
        ### skip already assigned IDs
        if type(metadata_df.loc[idx, 'ID']) is str or not math.isnan(metadata_df.loc[idx, 'ID']):
            metadata_df.loc[idx, 'ID'] = metadata_df.loc[idx, 'ID']
            print('Keeping existing ID: %s' %(metadata_df.loc[idx, 'ID']))
        else:
            ### add number to identical bgcids
            if strainstring in strain_count_dict.keys():
                suffixcount = strain_count_dict[strainstring]
                strain_count_dict[strainstring] = suffixcount - 1
                suffixcount = '_%s' % (suffixcount)
            else:
                suffixcount = ''
            #print(strainstring)
            ### special case for nan
            if type(strainstring) is float or metadata_df.loc[idx, 'source']=='LocalSeq':
                try:
                    bgcid = special_cases_dict[metadata_df.loc[idx, 'Database_ID']]
                except:
                    bgcid = special_cases_dict[metadata_df.loc[idx, 'seq_with_Xdom']]
            else:
                genus = strainstring.split(' ')[0]
                prefix = genus_to_char_dict[genus]
                #print([prefix])
                ### special case for unknown
                if prefix == 'U':
                    try:
                        bgcid = special_cases_dict[metadata_df.loc[idx, 'Database_ID']]
                    except:
                        bgcid = special_cases_dict[metadata_df.loc[idx, 'seq_with_Xdom']]
                else:
                    speciessuffixstring = strainstring[len(genus)+1:]
                    #print([strainstring])
                    species = speciessuffixstring.split(' ')[0]
                    middix = species_char_dict[species]
                    suffixstring = speciessuffixstring[len(species)+1:]
                    fullspecies = "%s %s" % (genus, species)
                    if 'subsp' in suffixstring:
                        subspecies = suffixstring.split(' ')[-1]
                        #print(subspecies)
                        middix = middix + subspecies[0]
                        suffixstring = suffixstring.split(subspecies)[-1]
                        #suffixstring = suffixstring[1:]
                        #print(suffixstring)
                    ### if there is a strain, deal with that too
                    if len(suffixstring) > 0:
                        if strainstring in strain_count_dict.keys() or not fullspecies_count_dict[fullspecies] == 1:
                            suffix = strain_char_dict[suffixstring]
                    else:
                        suffix = ''
                    bgcid = "%s%s%s%s" %(prefix, middix, suffix, suffixcount)
                    bgcid = special_fix_id(theid=bgcid, index=len(fullspecies), fullname=strainstring, list_of_ids=bgcids_list,
                                           suffixcount=str(suffixcount))


            if bgcid not in bgcids_list or bgcid == 'SvN':
                #print('save id')
                bgcids_list.append(bgcid)
                metadata_df.loc[idx, 'ID'] = bgcid
                strain_to_bgcid_dict[strainstring] = bgcid
            else:
                print("\nERROR: BGC ID %s already in use! Check the function or your input list\n" %(bgcid))
                print(strainstring)
                print(bgcids_list)
                pprint(strain_to_bgcid_dict)
                exit()
            #pprint(strain_to_bgcid_dict)

    #with pd.option_context('display.max_rows', None,
    #                       ):
    #    print(metadata_df[metadata_df['ID'].isna()].iloc[:,:5])
    #    print(metadata_df.iloc[:, :5])
    #duplicated = metadata_df[metadata_df['strain'].isnotna()]
    #print(duplicated)
    duplicated = metadata_df[['strain','Database_ID']].duplicated().value_counts()
    #duplicated = duplicated.count(True)
    print("\nCounting duplicates:")
    print(duplicated)
    return metadata_df


# Main function #
def main(input, seqtable, output, metadata, overwrite=False):
    print("\nInput directory of antiSMASH results: %s\n"
          "Input table with seq_with_Xdom: %s\n"
          "Output folder: %s\n"
          "Output metadata table: %s\n"
          "Overwrite: %s\n"
          %(input, seqtable, output, metadata, overwrite))
    create_folder(output)
    as_result_folders_list = list_folders(input)
    print("Searching in %s antiSMASH results folders\n" %(len(as_result_folders_list)))
    dbid_to_seqX_dict = csv_to_dict(seqtable, 'Database_ID', 'seq_with_Xdom')
    #print(dbid_to_seqX_dict.keys())
    #exit()
    #print(len(dbid_to_seqX_dict))
    #print(len(set(dbid_to_seqX_dict.keys())))
    #exit()
    #print(len(dbid_to_seqX_dict.keys()))
    #pprint(dbid_to_seqX_dict)
    gpa_files_source_target_dict = {}
    metadata_df = pd.read_csv(seqtable)
    ### get sensible IDs
    metadata_df = generate_bgcids(metadata_df)
    #print(metadata_df[metadata_df['source'] == 'Culp'])
    #exit()
    bgcid_to_xdomfile = {}
    species_to_char_dict = {}
    speciesname_to_char_dict ={}

    ### iterate over result folders
    for resultfolder in as_result_folders_list:
        #if resultfolder.endswith('Sva'):
        #    print(resultfolder)
        resultfolder_hit_counter = 0
        thisfolder_bgc_dict = {}
        thename = os.path.basename(resultfolder)
        #print(thename)
        splitseqlabels = True
        if thename in dbid_to_seqX_dict.keys():
            theseqX = dbid_to_seqX_dict[thename]
        ### cases of RefSeq and Genbank that have a version suffix
        elif "%s.1"%(thename) in dbid_to_seqX_dict.keys():
            theseqX = dbid_to_seqX_dict["%s.1"%(thename)]
        elif "%s.2"%(thename) in dbid_to_seqX_dict.keys():
            theseqX = dbid_to_seqX_dict["%s.2"%(thename)]
        ### for Culp dataset
        elif thename[:2].isnumeric() and str(int(thename[:2])) in dbid_to_seqX_dict.keys():
            theseqX = dbid_to_seqX_dict[str(int(thename[:2]))]#.split("|")[-1]
            splitseqlabels = False
        ### for Mgnify hardcoded
        elif "_FASTA" in thename:
            theseqX = 'ctg29508_3'#dbid_to_seqX_dict[thename.split("_FASTA")[0]]
        ### for Stammsammlung:
        elif thename.startswith("S00"):
            theseqX = dbid_to_seqX_dict[thename.split("_contig")[0]]
        ### for ShrikantOldData
        elif thename.startswith('QP') and "assembly" in thename:
            theseqX = dbid_to_seqX_dict[thename.split("_assembly")[0]]
            splitseqlabels = False
        else:
            print('Error: could not find the database id for %s' %(thename))
            #exit()
            print('Skipping for now...')
            continue
        ### added cases where more than one X domain was found
        if "|" in theseqX and splitseqlabels:
            theseqXlist = theseqX.split("|")
        ### cover Culp cases where the name is part of the seqX divided by "|"
        #elif "%s|" %(thename) in theseqX:
        elif "%s" %(thename) in theseqX:
            theseqXlist = theseqX.split("|")
        ### special case for Shrikant's old data
        elif "QPWMP018AI.assembly|ctg1442_135" in theseqX:
            theseqXlist = ['ctg1_36']
        else:
            theseqXlist = [theseqX]
        ### collect region files
        theregionfiles = locate_region_files(resultfolder)
        #if thename == 'GCF_013317105':
        #    print(thename, theregionfiles)
        positive_hits = []
        #if thename == '42_cluster_WAC00631':
        #    print(theseqXlist)
        #    exit()
        ### check each region file
        for theseqX in theseqXlist:
            foundXdom = False
            for regiongbk in theregionfiles:
                ### to cover cases where multiple hits in one BGC
                if regiongbk in positive_hits:
                    continue
                if grep_file(regiongbk, theseqX):
                    foundXdom = True
                    positive_hits.append(regiongbk)
                    resultfolder_hit_counter+=1
                    thetargetfile = os.path.join(output, '%s_%s.fna' % (thename, os.path.basename(regiongbk)[:-4]))
                    bgcid, bgcid_to_xdomfile = infer_bgcid(thetargetfile, metadata_df, bgcid_to_xdomfile)
                    ### testing
                    #if 'US9' in bgcid:
                    #    print(regiongbk)
                    ### forgive me for the hardcoding, I had enough
                    #if bgcid == 'US91_3':
                    #    #print('gotcha')
                    #    bgcid = 'US91'
                    #    #exit()
                    #bgcid, bgcid_to_xdomfile, species_to_char_dict, speciesname_to_char_dict = infer_bgcid(thetargetfile,
                    #                                                                                       metadata_df,
                    #                                                                                       bgcid_to_xdomfile,
                    #                                                                                       species_to_char_dict,
                    #                                                                                       speciesname_to_char_dict)

                    #print(bgcid)
                    ### extra process for cases where more than one BGC was found in one folder
                    if resultfolder_hit_counter == 1:
                        original_bgcid = bgcid
                    ### assign letters to multiple hits within one folder
                    if resultfolder_hit_counter > 1:
                        if not resultfolder.endswith('Sva'):
                            #pprint(thisfolder_bgc_dict)
                            bgcid = "%s_%s" %(bgcid, string.ascii_uppercase[resultfolder_hit_counter - 1])
                            new_row_bool = metadata_df['ID'] == original_bgcid
                            new_row = metadata_df[new_row_bool]
                            new_row.loc[:,'ID'] = bgcid
                            new_xseqs = []
                            for xseq in theseqXlist:
                                if grep_file(regiongbk, xseq):
                                    new_xseqs.append(xseq)
                            new_xseqs_string = "|".join(new_xseqs)
                            new_row.loc[:,'seq_with_Xdom'] = new_xseqs_string
                            #print(new_row)
                            metadata_df = metadata_df.append(new_row, ignore_index=True)
                        ### for special case of Sva
                        else:
                            bgcid = bgcid.replace('_A', '_B')
                        #print(metadata_df)
                    '''
                    ### extra check for unknown cases of two clusters in a genome
                    if bgcid in bgcid_to_xdomfile.keys():
                        #print('duplicate',bgcid)
                        bgc1 = "%s_1" % (bgcid)
                        bgc2 = "%s_2" % (bgcid)
                        bgcid_to_xdomfile[bgc1] = bgcid_to_xdomfile[bgcid]
                        del bgcid_to_xdomfile[bgcid]
                        bgcid = bgc2
                    '''
                    ### save info to dicts
                    bgcid_to_xdomfile[bgcid] = [regiongbk, thetargetfile]
                    thisfolder_bgc_dict[bgcid] = [regiongbk, thetargetfile]
        #pprint(thisfolder_bgc_dict)

        ### if there is more than one, remove the original from dicts and dfs
        if resultfolder_hit_counter > 1 and not resultfolder.endswith('Sva'):
            if bgcid in thisfolder_bgc_dict.keys():
                firstbgcid = "%s_A" % (original_bgcid)
                thisfolder_bgc_dict[firstbgcid] = thisfolder_bgc_dict[original_bgcid]
                bgcid_to_xdomfile[firstbgcid] = thisfolder_bgc_dict[original_bgcid]
                new_row_bool = metadata_df['ID'] == original_bgcid
                new_row = metadata_df[new_row_bool]
                new_row.loc[:,'ID'] = firstbgcid
                new_xseqs = []
                regiongbk = thisfolder_bgc_dict[original_bgcid][0]
                for xseq in theseqXlist:
                    if grep_file(regiongbk, xseq):
                        new_xseqs.append(xseq)
                new_xseqs_string = "|".join(new_xseqs)
                #new_row['seq_with_Xdom'] = new_xseqs_string ### getting warning here to use .loc[row_indexer,col_indexer] =
                #print(new_row)
                new_row.loc[:,'seq_with_Xdom'] = new_xseqs_string ### getting warning here to use .loc[row_indexer,col_indexer] =
                #print(new_row)
                metadata_df = metadata_df.append(new_row, ignore_index=True)
                #metadata_df.loc[metadata_df['ID'] == original_bgcid, 'ID'] = firstbgcid
                metadata_df.drop(metadata_df[metadata_df.ID == original_bgcid].index, inplace=True)

            #print("Removing %s" %(original_bgcid))
            if not resultfolder.endswith('Sva'):
                del bgcid_to_xdomfile[original_bgcid]
            else:
                print(resultfolder)
            metadata_df = metadata_df.drop(metadata_df[metadata_df['ID'] == original_bgcid].index, axis=0)
        if not foundXdom and not "S00" in thename:  # and not "QPWMP018AI.assembly" == theseqX \
            # and not ("ctg1442_135" == theseqX and 'QPWMP018AI_assembly_contig_304_region001' in resultfolder) \
            # and not ("contig_304_114" == theseqX and not 'QPWMP018AI_assembly_contig_304_region001' in resultfolder):
            print("\nWarning: could not find Xdomain sequence %s in any of the files under:\n%s"
                  % (theseqX, resultfolder))
    #print(metadata_df[metadata_df['source']== 'Culp'])
    #pprint(bgcid_to_xdomfile.keys())
    #exit()
    #if "StVrN_B" in bgcid_to_xdomfile.keys():
    #    print('found')
    ### go overthe finished dict and make the copies
    for bgcid, [regiongbk, thetargetfile] in bgcid_to_xdomfile.items():
        thetargetfile = os.path.join(os.path.dirname(thetargetfile), "%s__%s" %(bgcid, os.path.basename(thetargetfile)))
        ### to make sure no conflicts with names later
        thetargetfile = thetargetfile.replace("<","_")
        thetargetgbkfile = thetargetfile[:-4] + '.gbk'
        #bgcid_to_xdomfile[bgcid] = regiongbk
        gpa_files_source_target_dict[regiongbk] = thetargetfile
        ### copy and rename file
        if not os.path.exists(thetargetfile) or overwrite:
            #execute_cp(regiongbk, thetargetfile)
            gbk_to_fna(regiongbk, thetargetfile)
        ### add a faa file from the gbk file
        faafile = thetargetfile.replace('.fna','.faa')
        #if not os.path.exists(faafile) or overwrite:
        ### changed it so script should run even if file exists, for checking
        gbk_to_faa.main(regiongbk, faafile, type='file', overwrite=overwrite)
        ### copy and rename gbk file too
        if not os.path.exists(thetargetgbkfile) or overwrite:
            execute_cp(regiongbk, thetargetgbkfile)
                #if not foundXdom and "QPWMP018AI" in thename:


    #print(metadata_df)
    #pprint(bgcid_to_xdomfile)
    print("\nDetected %s regions with X domains\nThey can be found under: %s\n" %(len(gpa_files_source_target_dict.keys()),
                                                                                output))
    ### now update metadata table
    metadata_df = metadata_df.set_index('ID', drop=False)
    #print(metadata_df)
    extra_info_dict = {}
    newcols_list = 'as7_BGC_locus as7_start_gene as7_end_gene as7_clusterstart as7_clusterend on_contig_edge ' \
                   'considered_complete'.split(' ')
    ### fill dictionaries - some ids are missing from here (do they have no X domain or is there a bug?)
    bgcids_list = list(bgcid_to_xdomfile.keys())
    bgcids_list.sort()
    #print([x for x in bgcids_list if str(x) != 'nan'])
    todelete_rowsIds = []
    for bgcid in bgcids_list:
        #print(bgcid)
        #if bgcid not in ['Sv1', 'Sv2']:
        #    continue
        xdomgbkfile = bgcid_to_xdomfile[bgcid][0]
        #print(bgcid, xdomgbkfile)
        bgc_locus, startgene, endgene, clusterstart, clusterend, contig_edge, complete = extract_info_from_gbk(xdomgbkfile)
        #if bgcid == 'StVrN_B':
        #    print(bgcid)
        #break
        #print(metadata_df.index.tolist())
        if bgcid in metadata_df.index:
            source = metadata_df.loc[bgcid, 'source']#.to_string()
        #commented out on 09.10.2023 because it should already be covered above
        '''
        elif bgcid.endswith("_1") or bgcid.endswith("_2") or bgcid.endswith("_3"):
            #print(bgcid)
            source = metadata_df.loc[bgcid[:-2], 'source']
            ### new bgcid needs to be incorporated in the metadata_df
            metadata_df.loc[bgcid,:] = metadata_df.loc[bgcid[:-2], :].copy().replace(bgcid[:-2], bgcid)
            #print(type(metadata_df.loc[bgcid[:-2], :]))
            ### old bgcid needs to be removed in the end
            todelete_rowsIds.append(bgcid[:-2])
            #print(metadata_df.loc[bgcid[:-2], :])
            #print(metadata_df.loc[bgcid])
        '''
        #print([source])
        ### we don't know about Culp and MIBiG because they were already isolated BGCs before the antiSMASH run
        if source in ['Culp','MIBiG']:
            contig_edge = ''
        extra_info_dict[bgcid] = [bgc_locus, startgene, endgene, clusterstart, clusterend, contig_edge, complete]
    todelete_rowsIds = list(set(todelete_rowsIds))
    #print(metadata_df)
    metadata_df = metadata_df.drop(todelete_rowsIds)
    #exit()
    extra_df = pd.DataFrame.from_dict(extra_info_dict, orient='index', columns=newcols_list)
    metadata_df = pd.concat([metadata_df, extra_df], axis=1)
    metadata_df.to_csv(metadata, index=False)
    print("Created metadata table:\n%s" %(metadata))
    print("\nDone")

#'''
# Parsing arguments & Execution #
parser = argparse.ArgumentParser(description='Given the directory containing the antismash result folders and the '
                                             'table that includes the seq_with_Xdom table, detect the antiSMASH '
                                             'regions that contain an X domain and output their DNA sequence to'
                                             ' the output folder.')
parser.add_argument('-i', '--input', help='Input directory of antiSMASH result folders', required=True)
parser.add_argument('-t', '--table', help='Input table (csv) connecting the Database_ID to the seq_with_Xdom', required=True)
parser.add_argument('-o', '--output', help='Output folder where the positive hits will be found', required=True)
parser.add_argument('-m', '--metadata', help='Output table (csv) - will have same info as argument table but with '
                                             'antiSMASH cluster information too', required=True)
parser.add_argument('-r', '--overwrite', help='Optional: overwrite existing results (folders). Default: False',
                    required=False, default=False)
if __name__ == "__main__":
    args = parser.parse_args()
    main(args.input, args.table, args.output, args.metadata, args.overwrite)
#'''

'''
### special case only for testing
gbkdir = '/media/hdd_2tb/GPA_reclassification_clean/preprint_testdata/2_putative_GPAs_analyses/1_regions_withXdom'
gbklist = list_files(gbkdir, suffix='.gbk')
for gbkfile in gbklist:
    if not "SsAg" in gbkfile:
        continue
    print(os.path.basename(gbkfile))
    outfna = gbkfile.replace('.gbk','.fna')
    print(outfna)
    gbk_to_fna(gbkfile, outfna)
'''
'''
### special case for debugging:
inputdir = '/media/hdd_2tb/GPA_reclassification_clean/A_dataset_creation/2_putative_GPAs_analyses/0_minimal_antismash7_full_results'
outputdir = '/media/hdd_2tb/GPA_reclassification_clean/A_dataset_creation/2_putative_GPAs_analyses/1_regions_with_Xdom'
table = '/media/hdd_2tb/GPA_reclassification_clean/A_dataset_creation/1_positive_hits_downloads/summary_hits_metadata.csv'
out_metadata = '/media/hdd_2tb/GPA_reclassification_clean/A_dataset_creation/2_putative_GPAs_analyses/summary_hits_clusters_metadata.csv'
main(inputdir, table, outputdir, out_metadata)
'''

'''
### special case for undetected file:
regiongbk = '/media/hdd_2tb/GPA_reclassification_clean/A_dataset_creation/2_putative_GPAs_analyses/1_regions_with_Xdom/StJs_A__GCF_015163075_NZ_JADDXU010000021.1.region001.gbk'
thetargetfile = '/media/hdd_2tb/GPA_reclassification_clean/A_dataset_creation/2_putative_GPAs_analyses/1_regions_with_Xdom/StJs_A__GCF_015163075_NZ_JADDXU010000021.1.region001.fna'
gbk_to_fna(regiongbk, thetargetfile)
'''
```
