## Supplementary Data 5 for "Structural and evolutionary analyses support reclassification of glycopeptide antibiotics as xyclopeptides": download_mgnify_dataset.py

```
#!/usr/bin/python3.6
# Created by Athina Gavriilidou at 27.06.23 15:57 #
# Imports #
import argparse
from Bio import SeqIO
import os
from pprint import pprint
import csv
import subprocess
import pandas as pd
import math
import json


# Functions #
'''given a directory, lists all gbk files'''
def list_gbk_files(directory):
    gbk_files = []
    for file in os.listdir(directory):
        if file.endswith(".gbk"):
            gbk_files.append(os.path.join(directory, file))
    print("Found " + str(len(gbk_files)) + " .gbk files")
    return gbk_files

def append_lines_csv(lines, output):
    with open(output, 'a') as f:
        tsv_output = csv.writer(f)
        for line in lines:
            tsv_output.writerow(line)
    print("\nAppended to file: " + output)

'''Takes a gbk file and creates an faafile with all AA seqs indexed by the locus tag. taken from source above'''
def get_acc_from_gbk(gbkfile):
    accession = False
    input_handle = open(gbkfile, "r")
    bgc_filename = gbkfile.split("/")[-1]
    bgc_filename = bgc_filename[:-4]
    #print(bgc_filename)
    for seq_record in SeqIO.parse(input_handle, "genbank"):
        if accession:
            break
        #print("Dealing with GenBank record %s" % seq_record.id)
        dbxrefs = seq_record.dbxrefs
        for xref in dbxrefs:
            if xref.startswith("Assembly"):
                accession = xref.split(":")[-1]
                #print(accession)
    input_handle.close()
    if not accession:
        #print("No assembly accession available for %s\n" %(bgc_filename))
        accession = ""
        print("Manual search required for %s" % (bgc_filename))
        '''
        input_handle = open(gbkfile, "r")
        for seq_record in SeqIO.parse(input_handle, "genbank"):
            # print("Dealing with GenBank record %s" % seq_record.id)
            dbxrefs = seq_record.dbxrefs
            for xref in dbxrefs:
                if xref.startswith("BioProject"):
                    bioproject = xref.split(":")[-1]
                elif xref.startswith("BioSample"):
                    biosample = xref.split(":")[-1]
            print("BioProject: %s\tBioSample: %s" %(bioproject, biosample))
            break
        input_handle.close()
        '''
    return [bgc_filename, accession]

def csv_to_dict(csvfile, key="", value=""):
    thedict ={}
    with open(csvfile, 'r') as infile:
        reader = csv.reader(infile)
        for headerline in reader:
            key_id = headerline.index(key)
            value_id = headerline.index(value)
            break
        for line in reader:
            thedict[line[key_id]] = line[value_id]
    return thedict

'''taken from https://stackoverflow.com/questions/2170900/get-first-list-index-containing-sub-string'''
def index_containing_substring(the_list, substring):
    for i, s in enumerate(the_list):
        #print([s, substring, i])
        if substring in s or substring == s:
              return i
    return -1

def filtered_csv_to_dict(filepath, key="", value="", delim=',', skiplines=1, filterlist=[]):
    if delim is None:
        if filepath.endswith(".csv"):
            delim = ","
        else:
            delim = "\t"
    content = {}
    if len(filterlist) > 0:
        applyfilter = True
    else:
        applyfilter = False
    with open(filepath) as fd:
        rd = csv.reader(fd, delimiter=delim, quotechar='"')
        while skiplines != 0:
            for skipline in rd:
                break
            skiplines -= 1
        for headline in rd:
            if not isinstance(key, int):
                key = index_containing_substring(headline, key)
                #print(headline, key)
            if not isinstance(value, int):
                value = index_containing_substring(headline, value)
                #print(value)
            break
        for line in rd:
            if applyfilter and line[key] not in filterlist:
                continue
            if type(value) == list:
                this_value = []
                for v in value:
                    this_value.append(line[v])
                content[line[key]] = this_value
            else:
                content[line[key]] = line[value]
    return content

def get_all_genome_files(accession, theoutput):
    home_directory = os.path.expanduser('~')
    ncbitoolpath = os.path.join(home_directory, 'anaconda3/bin/ncbi-genome-download')
    command = "%s --parallel 30 --format all --assembly-accessions " \
              "%s -o %s bacteria" %(ncbitoolpath, accession, theoutput)
    print("Getting  %s" %(accession))
    subprocess.run(command, shell=True)
    new_loc = os.path.join(theoutput, accession)
    old_loc = os.path.join(theoutput, "refseq/bacteria/%s" %(accession))
    mvcommand = "mv %s %s" %(old_loc, new_loc)
    subprocess.run(mvcommand, shell=True)
    if not os.path.exists(new_loc):
        print("Failed to download %s" %(accession))


def gunzip_copy(new_loc, correct_loc, filesuffix, name):
    gzfile = ""
    for file in os.listdir(new_loc):
        if file.endswith(filesuffix):
            gzfile = os.path.join(new_loc, file)
            break
    correct_file = os.path.join(correct_loc, "%s_strain_proteome.faa" %(name))
    if os.path.exists(correct_file):
        return
    if gzfile != "":
        gzcommand = "gunzip -c %s > %s" %(gzfile, correct_file)
        subprocess.run(gzcommand, shell=True)
    else:
        print("No %s file found in %s" %(filesuffix, new_loc))

'''given a df, get list of refseq/genbank ids by locating correct column'''
def get_ids_from_df(thedf, id_keywords, ids_dict={}):
    if len(ids_dict.keys()) == 0:
        id_col = ''
        for thecol in thedf.columns:
            if id_col != '':
                break
            for id_kw in id_keywords:
                if id_kw in thecol:
                    id_col = thecol
                    print('ID column detected: %s' %(id_col))
                    break
        ids_dict = thedf[id_col].to_dict()
    else:
        ### find correct column
        id_col = ''
        for thecol in thedf.columns:
            if id_col != '':
                break
            for id_kw in id_keywords:
                if id_kw in thecol:
                    id_col = thecol
                    print('ID column detected: %s' % (id_col))
                    break
        for gpa, id in ids_dict.items():
            ### in cases where there was no id found for the genome use the cluster id
            if not isinstance(id, str):  # and math.isnan(id):
                ids_dict[gpa] = thedf.loc[gpa, id_col].split(":")[0] ### added this to cover some weird cases
        ### remove also the sequenced ones (they dont have an id)
        ids_dict = {gpa: id for gpa, id in ids_dict.items() if id != 'sequenced'}
        ### fix cases of two clusters from single genome
        fixcases = {}
        for gpa in ids_dict.keys():
            if gpa.endswith("_1"):
                newkey = gpa[:-2]
                fixcases[newkey] = ids_dict[gpa]
        for key, value in fixcases.items():
            ids_dict[key] = value
        ids_dict = {gpa: id for gpa, id in ids_dict.items() if not gpa.endswith('_1') and not gpa.endswith('_2')}
    return ids_dict

'''creates folder in given path, if it does not exist already'''
def create_folder(path):
    if not os.path.exists(path):
        os.mkdir(path)

def run_download_command(seqid, outfolder):
    print('seqid', seqid)
    ### first try refseq
    if seqid.startswith('GCF'):
        command = 'ncbi-genome-download --parallel 30 --format all --assembly-accessions %s -o %s ' \
                  'bacteria' %(seqid, outfolder)
        #create_folder(outfolder)
        subprocess.run(command, shell=True)
        final_dir = os.path.join(os.path.join(os.path.join(outfolder,'refseq'), 'bacteria'), seqid)
        command = 'mv %s/* %s/.' %(final_dir, outfolder)
        subprocess.run(command, shell=True)
    ### then try genbank
    elif seqid.startswith('GCA'):
        command = 'ncbi-genome-download --section genbank --parallel 30 --format all --assembly-accessions %s ' \
                  '-o %s bacteria' % (seqid, outfolder)
        #create_folder(outfolder)
        subprocess.run(command, shell=True)
        final_dir = os.path.join(os.path.join(os.path.join(outfolder, 'genbank'), 'bacteria'), seqid)
        command = 'mv %s/* %s/.' % (final_dir, outfolder)
        subprocess.run(command, shell=True)
    ### then try nucleotide record of the cluster
    else:
        ### we have one JGI genome
        if seqid == 'Gp0217399':
            return
        outfolder = outfolder.replace("strain", 'cluster')
        create_folder(outfolder)
        for molecule in ['nucleotide', 'protein']:
            print('\nTrying to download %s %s records' %(seqid, molecule))
            for format in 'fasta,genbank'.split(','):
                outfile = os.path.join(outfolder, "%s_%s.%s" %(seqid, molecule[:4], format))
                command = 'ncbi-acc-download --molecule %s --extended-validation all --format %s --out %s ' \
                           '%s' %(molecule,format, outfile, seqid)
                subprocess.run(command, shell=True)

def detect_file(directory, fileprefix):
    onlyfiles = [f for f in os.listdir(directory) if os.path.isfile(os.path.join(directory, f))
                 and f.startswith(fileprefix)]
    if len(onlyfiles) == 1:
        return onlyfiles[0]
    else:
        print('Found multiple files with prefix %s' %(fileprefix))
        pprint(onlyfiles)

def run_datasetsdownload_command(seqid, outfolder):
    command = '/home/athina/anaconda3/envs/ncbi_datasets/bin/datasets download genome accession %s ' \
              '--filename %s' %(seqid, outfolder)
    subprocess.run(command, shell=True)
    command = 'unzip %s -d %s' %(outfolder, outfolder[:-4])
    subprocess.run(command, shell=True)
    command = 'rm %s' % (outfolder)
    subprocess.run(command, shell=True)
    outfolder = outfolder[:-4]
    datafolder1 = os.path.join(os.path.join(outfolder, 'ncbi_dataset'), 'data')
    datafolder2 = os.path.join(datafolder1, seqid)
    genomefastafile = detect_file(datafolder2, seqid)
    fullprefix = genomefastafile.split('_genomic.fna')[0]
    print("Renaming files using prefix %s" %(fullprefix))
    level1files = [os.path.join(datafolder1, f) for f in os.listdir(datafolder1)
                   if os.path.isfile(os.path.join(datafolder1, f))]
    level2files = [os.path.join(datafolder2, f) for f in os.listdir(datafolder2)
                   if os.path.isfile(os.path.join(datafolder2, f))]
    for thefile in level1files + level2files:
        if thefile.startswith(fullprefix):
            command = 'mv %s %s/.' %(thefile, outfolder)
        else:
            command = 'mv %s %s/%s_%s' %(thefile, outfolder, fullprefix, os.path.basename(thefile))
        subprocess.run(command, shell=True)

def extracheck(outputdir):
    useddir = '/media/hdd_2tb/phd_main/GPA_producer_genomes/0_genomes_full'
    if not os.path.exists(useddir):
        return
    usedfolders = [f for f in os.listdir(useddir)]
    usedfolders.sort()
    outfolders = [f for f in os.listdir(outputdir) if '_cluster_' not in f]
    outfolders.sort()
    for outf in outfolders:
        if outf not in usedfolders:
            print("WARNING: Folder %s not found!" %(outf))
    for usedf in usedfolders:
        if usedf not in outfolders:
            print("NOTE: should folder %s also be present?" %(usedf))
    print("")

def list_files(directory, suffix=''):
    if suffix == '':
        files_list = [os.path.join(directory, o) for o in os.listdir(directory)
                        if os.path.isfile(os.path.join(directory,o))]
    else:
        files_list = [os.path.join(directory, o) for o in os.listdir(directory)
                      if os.path.isfile(os.path.join(directory, o)) and o.endswith(suffix)]
    files_list.sort()
    return files_list

def run_gunzip(filelist):
    ### addition 24.11.22 to not decompress if everything is there already
    dorun = False
    for f in filelist:
        if not os.path.exists(f[:-3]):
            dorun = True
    if not dorun:
        return
    if len(filelist) > 0:
        print("Decompressing files in %s" %(os.path.dirname(filelist[0])))
    for f in filelist:
        if not os.path.exists(f[:-3]):
            command = 'gunzip -d -k %s' %(f)
            subprocess.run(command, shell=True)


def get_positive_hits(csvfile, id_keywords):
    positive_hits = []
    with open(csvfile, 'r') as infile:
        reader = csv.reader(infile)
        for headerline in reader:
            for i in range(len(headerline)):
                if headerline[i] in id_keywords:
                    key_id = i
            break
        try:
            key_value = headerline[i]
            print('Detected column %s' %(key_value))
        except:
            print("ERROR: No column with suitable header detected. Must be one of %s" %(id_keywords))
        for line in reader:
            ishit = line[-1]
            if ishit == 'True':
                positive_hits.append(line[key_id])
    print("Found %s positive hits" %(len(positive_hits)))
    return positive_hits

def execute_download(dlink, outfile):
    wget_command = 'wget -O %s %s' %(outfile, dlink)
    try:
        print("Using link: %s" %(dlink))
        os.system(wget_command)
    except:
        return dlink

def download_assembly_summary_table(input):
    ### get link
    if 'refseq' in input:
        thelink='https://ftp.ncbi.nlm.nih.gov/genomes/ASSEMBLY_REPORTS/assembly_summary_refseq.txt'
    elif 'genbank' in input:
        thelink = 'https://ftp.ncbi.nlm.nih.gov/genomes/ASSEMBLY_REPORTS/assembly_summary_genbank.txt'
    else:
        print("\n ERROR: could not recognize if genbank or refseq. Please provide the table or a file path with "
              "either of the two keywords (refseq|genbank) in the name\n")
        exit()
    execute_download(thelink, input)

def list_files_rec(output, list_of_suffixes):
    filelist = []
    folder_list = [os.path.join(output, o) for o in os.listdir(output)
                  if os.path.isdir(os.path.join(output, o))]
    for folder in folder_list:
        thisfolderfiles = []
        for suffix in list_of_suffixes:
            files_list = [os.path.join(folder, o) for o in os.listdir(folder)
                      if os.path.isfile(os.path.join(folder, o)) and o.endswith(suffix)]
            thisfolderfiles = thisfolderfiles + files_list
        filelist = filelist + thisfolderfiles
    filelist.sort()
    return filelist

def transform_series_to_string(seq_withXdom, keeponlyone=False):
    if type(seq_withXdom) is str:
        return seq_withXdom
    elif not keeponlyone:
        seq_withXdom = seq_withXdom.tolist()
        seq_withXdom = "|".join(x for x in seq_withXdom)
        return seq_withXdom
    else:
        seq_withXdom = seq_withXdom.tolist()
        return seq_withXdom[0]

def detect_json_link(tempjsonfile):
    with open(tempjsonfile, 'r') as jsonfile:
        jsoncontent = json.load(jsonfile)
        data = jsoncontent['data']
        for d in data:
            if "FASTA.fasta" in d['id'] :
                cdsfastalink = d['links']['self']
                cdsfastalink = cdsfastalink.rstrip('?format=json')
                return cdsfastalink

    return ''

def execute_delete(path):
    ### added recursive option for lists
    if type(path) == list:
        for el in path:
            execute_delete(el)
    ### redirect errors to /dev/null for clearer output
    rm_command = 'rm %s 2> /dev/null' %(path)
    try:
        os.system(rm_command)
    except:
        return path

def fill_meta_table(assemblies_df, positive_hits_ids, meta_df, input, outtable, hmmhits_table,
                    hits_accno_to_dlink_dict, output):
    lines_to_append = []  ### see download_mibig_dataset.py - same concept
    accno_to_assemblies_df = assemblies_df.copy()
    accno_to_assemblies_df.set_index("assembly", inplace=True)
    #print(accno_to_assemblies_df)
    source = 'MGnify'
    list_of_already_registered_accnos = meta_df['Database_ID'].tolist()
    list_of_failed_downloads = []
    print("\nStarting assemblies download...\n")
    ### iterate over assemblies
    for accno, downlink in hits_accno_to_dlink_dict.items():
        accnofolder = os.path.join(output, accno)
        create_folder(accnofolder)
        thefastafile = os.path.join(accnofolder, "%s_FASTA.fasta.gz" % (accno))
        if not os.path.exists(thefastafile[:-3]):
            if not os.path.exists(thefastafile):
                tempjsonfile = os.path.join(accnofolder, "%s.json" % (accno))
                execute_download(downlink, tempjsonfile)
                fastalink = detect_json_link(tempjsonfile)
                execute_delete(tempjsonfile)
            ### get fasta file
            execute_download(fastalink, thefastafile)
            run_gunzip([thefastafile])
        ### skip the appending to meta table if the accno is already registered there
        if accno in list_of_already_registered_accnos:
            continue
        ### get info row from assemblies df
        #this_row = accno_to_assemblies_df.loc[accno]
        strain_string = ''#this_row['organism_name']
        ncbi_accno = ''
        db_id = accno
        genus = strain_string.split(" ")[0]
        download_link = downlink
        product = ''
        note = ''
        ### getting search date
        try:
            dateofsearch = hmmhits_table.loc[db_id, 'search_date']
            dateofsearch = transform_series_to_string(dateofsearch, keeponlyone=True)
        except:
            dateofsearch = ''
        seq_withXdom = hmmhits_table.loc[db_id, 'target']
        seq_withXdom = transform_series_to_string(seq_withXdom)
        ### fill in row list columns='
        # strain ID source NCBI_AccNo Database_ID seq_with_Xdom product genus date_of_search download_link notes'
        row_list = [strain_string, '', source, ncbi_accno, db_id, seq_withXdom, product, genus, dateofsearch,
                    download_link, note]
        ### immediately append to growing table
        append_lines_csv([row_list], outtable)
        #lines_to_append.append(row_list)
    ### Maybe in the future we want to add assigned ID to these
    print("Warning: generation of assigned ID has not been implemented. Column will be empty.")

    if len(list_of_failed_downloads) > 0:
        print("\n %s downloads have failed:\n" % (len(list_of_failed_downloads)))
        pprint(list_of_failed_downloads)

    list_of_fasta_files = list_files_rec(output, ['.fna'])
    if len(list_of_fasta_files) == len(positive_hits_ids):
        print("\nAll %s genomic fasta files successfully downloaded." % (len(positive_hits_ids)))
    else:
        print("\nWarning: Irregular number of fasta files")
    '''
    ### join existing and new rows
    meta_df_table = meta_df.values.tolist()
    lines_to_append = meta_df_table + lines_to_append
    new_df = pd.DataFrame(lines_to_append,
                          columns='strain ID source NCBI_AccNo Database_ID seq_with_Xdom product genus '
                                  'date_of_search download_link notes'.split(' '))
    ### save to table
    new_df.to_csv(outtable, index=False)
    # print(new_df.iloc[:5])
    '''
    print("\nTable has been created/updated: %s\n" % (outtable))

# Main function #
def main(input, assemblytable, output, outtable, domtbloutfile):
    print("\nInput table of hmmsearch finished runs: %s\n"
          "Input table of MGnify accessions: %s\n"
          "Output folder: %s\n"
          "Output metadata table: %s\n"
          "HMMsearch domain summary file: %s\n"
          %(input, assemblytable, output, outtable, domtbloutfile))
    #exit()
    create_folder(output)
    id_keywords = 'assembly,accno,db_id'.split(',')
    positive_hits =  get_positive_hits(input, id_keywords)
    #print(positive_hits)
    #exit()

    hits_accno_to_dlink_dict = filtered_csv_to_dict(assemblytable, 'assembly', 'downloadlink', skiplines=0,
                                                    filterlist=positive_hits)
    #print(hits_accno_to_dlink_dict)
    #exit()
    if len(positive_hits) != len(hits_accno_to_dlink_dict):
        print("\nWarning: Number of positive hits (%s) does not correspond to number of download links found (%s)."
              " Manual check may be necessary.\n" %(len(positive_hits), len(hits_accno_to_dlink_dict)))

    ### read metadata table if it exists, otherwise create it
    if os.path.exists(outtable):
        meta_df = pd.read_csv(outtable)
    else:
        meta_df = pd.DataFrame(columns='strain ID source NCBI_AccNo Database_ID seq_with_Xdom product genus '
                                       'date_of_search download_link notes'.split(' '))
        meta_df.to_csv(outtable, index=False)

    ### import hmmsearch info and everything else needed
    assemblies_df = pd.read_csv(assemblytable)
    ### take note that this actually works!
    hmmhits_table = pd.read_csv(domtbloutfile, index_col=False, dtype='str')#, header=None,### comment out header is None here!!!!!!!!!!!!
                                #names='db_id	target	accession	c-Evalue	i-Evalue	score	bias	description'.split('\t'))
                                      #'search_date'.split('\t'))
    #print(hmmhits_table)
    ### to fix mismatch of accnos
    hmmhits_table['db_id_fixed'] = hmmhits_table.db_id.apply(lambda x: x[:15])
    hmmhits_table.set_index('db_id_fixed', inplace=True)
    positive_hits_ids = hmmhits_table.index.tolist()

    ### start filling the table before downloading genomes
    fill_meta_table(assemblies_df, positive_hits_ids, meta_df, input, outtable, hmmhits_table,
                    hits_accno_to_dlink_dict, output)

    #print(assemblies_df[['#assembly_accession', 'organism_name']])
    #print(assemblies_df.columns)
    #print(assemblies_df)
    #exit()

    #pprint(hits_accno_to_dlink_dict)
    #gpas_to_ids_dict = get_ids_from_df(todownload_df, id_keywords)
    #gpas_to_ids_dict = get_ids_from_df(todownload_df, ['cluster'], ids_dict=gpas_to_ids_dict)

    ### for testing only:
    #gpas_to_ids_dict = {'AkA': 'GCA_002934265.1', 'AmAl': 'GCF_000384215.1','AmSW1375': 'JX576190.1'}
    #if len(gpas_to_ids_dict.keys()) < len(todownload_df.index) - 3:
    #    print("\nNOTE: this is a sample dataset")


    '''
    ### check that everything was downloaded
    check = True
    print("\nChecking if everything was downloaded...")
    for gpa, id in gpas_to_ids_dict.items():
        id = id.rstrip()
        thisoutput = os.path.join(output, '%s_strain_%s' % (gpa, id))
        if (not os.path.exists(thisoutput) and not os.path.exists(thisoutput.replace("strain", 'cluster')))\
                or (os.path.exists(thisoutput) and
                    len([f for f in os.listdir(thisoutput)if os.path.isfile(os.path.join(thisoutput, f))]) == 0) \
                or (os.path.exists(thisoutput.replace("strain", 'cluster')) and
                    len([f for f in os.listdir(thisoutput.replace("strain", 'cluster'))
                          if os.path.isfile(os.path.join(thisoutput.replace("strain", 'cluster'), f))]) == 0):
            print("Failed to download %s with ID %s" %(gpa, id))
            if id.startswith("GC"):
                print('Retrying...')
                run_datasetsdownload_command(id, "%s.zip" %(thisoutput))
                if not os.path.exists(thisoutput):
                    print("Failed to download")
                    check = False
            else:
                print("NOTE: JGI-derived sequences CANNOT be automatically downloaded with this script.")
                check = False
    if check:
        print("All downloads successful! Dataset complete!\n")
    extracheck(output)
    '''


    '''
    ### unzip everything we need
    for gpa, id in gpas_to_ids_dict.items():
        thisoutput = os.path.join(output, '%s_strain_%s' % (gpa, id))
        ###only relevant for whole genomes
        if os.path.exists(thisoutput):
            compresedfiles = list_files(thisoutput, suffix='.gz')
            todecompress= [f for f in compresedfiles if ('_genomic.fna' in f and '_cds_from_' not in f)
                           or ('_feature_table.txt' in f) or ('genomic.gff' in f)]
            run_gunzip(todecompress)
    '''


    '''
    gbk_genomes = list_gbk_files(input)
    gbk_genomes.sort()
    genome_meta = os.path.join(output, "GPA_producer_genomes_metadata.csv")
    genome_to_acc_dict = csv_to_dict(genome_meta, "ID", "Refseq")
    name_to_acc_dict = {}
    for genome in gbk_genomes:
        bgc_filename, accession = get_acc_from_gbk(genome)
        name_to_acc_dict[bgc_filename] = accession
    for name, accession in name_to_acc_dict.items():
        print(name, accession)
        if accession == "":
            try:
                accession = genome_to_acc_dict[name]
                name_to_acc_dict[name] = accession
            except:
                print("No Accession found for %s" %(name))
                del name_to_acc_dict[name]
                continue
        ###try to download
        new_loc = os.path.join(output, "%s_strain_%s" %(name, accession))
        suppressed_assemblies = {"AmSW5977": "GCA_003977855.1", "KA16535": "GCA_000720375.1",
                                 "SsA": "GCA_001298575.1"}
        if name in suppressed_assemblies.keys(): ### exception, suppressed accession, manually downloaded
            new_loc = os.path.join(output, "%s_strain_%s" %(name, suppressed_assemblies[name]))
            print(new_loc)
        if not os.path.exists(new_loc):
            get_all_genome_files(name, accession, output)
        ### copy proteomes
        correct_loc = os.path.join(os.path.dirname(output), "proteomes_faa")
        gunzip_copy(new_loc, correct_loc, "_protein.faa.gz", name)

    '''
    print("\nDone")


#'''
# Parsing arguments & Execution #
parser = argparse.ArgumentParser(description='Given a table of Mgnify Accession numbers, downloads all '
                                             'files related to them')
parser.add_argument('-i', '--input', help='Input table (.csv) of mgnify accnos (csv with column label assembly '
                                          ' or accno or db_id)', required=True)
parser.add_argument('-t', '--table', help='Input mgnify accession number to download link table.', required=True)
parser.add_argument('-o', '--output', help='Output directory where data will be downloaded', required=True)
parser.add_argument('-m', '--metadata', help='Output metadata table with info on downloaded genomes. If exists, will '
                                             'be appended.', required=True)
parser.add_argument('-d', '--domtblout', help='Summary domtblout (tsv) of HMM search on the clusters', required=True)
if __name__ == "__main__":
    args = parser.parse_args()
    main(args.input, args.table, args.output, args.metadata, args.domtblout)
#'''
```
