## Supplementary Data 5 for "Structural and evolutionary analyses support reclassification of glycopeptide antibiotics as xyclopeptides": download_mibig_dataset.py

```
#!/usr/bin/python3.6
# Created by Athina Gavriilidou at 12.06.23 15:27 #
# Imports #
import argparse
import datetime
import os
import pandas as pd
import dateutil.parser as dparser
from collections import OrderedDict
import json
from pprint import pprint


# Functions #
def create_folder(path):
    if not os.path.exists(path):
        os.mkdir(path)

### get run date from last 10 lines of domtblout
def extract_date(last_tblout_lines):
    for line in last_tblout_lines.target_name:
        if "Date" in line:
            thedate = dparser.parse(line, fuzzy=True)
            return thedate.strftime('%d-%m-%Y')

### nice way to visualize hmmsearch domtblout
def import_hmmsearch_tblout(filepath):
    # manually specify col limits
    dom_hits_colspecs = [(0, 252), (252, 264), (264, 270), (270, 291), (291, 302), (302, 308), (308, 318), (318, 325),
                         (325, 331), (331, 335), (335, 339), (339, 349), (349, 359), (359, 366), (366, 372), (372, 378),
                         (378, 384), (384, 390), (390, 396), (396, 402), (402, 408), (408, 413), (413, 435)]
    tblout_df = pd.read_fwf(filepath, skiprows=3, index_col=False,
                                             colspecs=dom_hits_colspecs,
                                             names=['target_name', 'accession1', 'tlen', 'query_name', 'accession2',
                                                    'full_sequence_qlen', 'full_sequence_E-value',
                                                    'full_sequence_score', 'full_sequence_bias', 'this_domain_#',
                                                    'this_domain_of', 'this_domain_c-Evalue', 'this_domain_i-Evalue',
                                                    'this_domain_score', 'this_domain_bias', 'hmm_coord_from',
                                                    'hmm_coord_to', 'ali_coord_from', 'ali_coord_to', 'env_coord_from',
                                                    'env_coord_to', 'acc', 'description_of_target'])
    the_run_date = extract_date(tblout_df.iloc[-10:])
    ### skip version lines
    return tblout_df.iloc[:-10], the_run_date

def execute_delete(path):
    ### added recursive option for lists
    if type(path) == list:
        for el in path:
            execute_delete(el)
    ### redirect errors to /dev/null for clearer output
    rm_command = 'rm %s 2> /dev/null' %(path)
    try:
        os.system(rm_command)
    except:
        return path

def get_info_from_json(json_download_link, output, db_id):
    tempfilename = os.path.join(output, '%s_tempfile.json' %(db_id))
    download_command = 'wget %s --output-document %s' %(json_download_link, tempfilename)
    os.system(download_command)
    ### look for the info in the json file
    with open(tempfilename, 'r') as jsonfile:
        jsoncontent = json.load(jsonfile)
        cluster_info = jsoncontent['cluster']
        product = cluster_info['compounds'][0]['compound']
        strainname =  cluster_info['organism_name']
        genus = strainname.split(' ')[0]
    execute_delete(tempfilename)
    return strainname, product, genus

def execute_download(dlink, outfile):
    wget_command = 'wget -O %s %s' %(outfile, dlink)
    try:
        os.system(wget_command)
    except:
        return dlink

'''assign bgc ids based on the product made'''
def assign_ids(lines_to_append, old_products_list, new_products_list, old_meta_df_table):
    print("\nNow reassigning IDs...\n")
    product_col_index = 6
    id_col_index = 1
    db_id_index = 4
    product_to_occurence_count = {}
    lines_to_append = old_meta_df_table + lines_to_append
    list_of_ids = []
    '''
    ### if it was already in the table it already has been assigned an ID
    for product in old_products_list:
        if product not in product_to_occurence_count.keys():
            product_to_occurence_count[product] = [1, 1]
        else:
            temp_occ = product_to_occurence_count[product][0]
            total_occ = product_to_occurence_count[product][1]
            product_to_occurence_count[product] = [temp_occ + 1, total_occ + 1]
    ## if not but we have it in the new version we should be ready to assign an ID
    '''
    ### gather total occurences and start counters
    for product in new_products_list + old_products_list:
        if product not in product_to_occurence_count.keys():
            product_to_occurence_count[product] = [1, 0]
        else:
            temp_occ = product_to_occurence_count[product][0]
            product_to_occurence_count[product] = [temp_occ + 1, 0]
    #pprint(product_to_occurence_count)
    ### iterate over table
    for rowindex in range(len(lines_to_append)):
        row = lines_to_append[rowindex]
        product = row[product_col_index]    ### WARNING: here it is assumed that the product column has index 6
        standard_len = 3
        assigned_id = product.replace("-","")[:standard_len]
        ### cover the case of "iso"
        if product.startswith('iso'):
            assigned_id = "%s%s" %(assigned_id, product.replace("-","")[standard_len:2*standard_len])
        ### to avoid accidental duplicates
        while assigned_id in list_of_ids:
            if " " in product:
                assigned_id = "%s%s" %(assigned_id, product.split(" ")[-1])
                continue
            standard_len+=1
            assigned_id = product.replace("-", "")[:standard_len]

        total_occ = product_to_occurence_count[product][0]
        ### if there is more than one we need numbers
        if total_occ > 1:
            current_occ = product_to_occurence_count[product][1] + 1
            product_to_occurence_count[product] = [total_occ, current_occ]
            assigned_id = '%s_%s' % (assigned_id, current_occ)
        list_of_ids.append(assigned_id)
        print("Entry %s assigned ID %s" %(row[db_id_index], assigned_id))
        ### save new id in table
        row[id_col_index] = assigned_id
        lines_to_append[rowindex]= row
    print("\nWarning: IDs have been updated and may differ from the previous version of the table\n")
    ### now update meta_df and append to existing one
    new_df = pd.DataFrame(lines_to_append,
                          columns='strain ID source NCBI_AccNo Database_ID seq_with_Xdom product genus '
                                  'date_of_search download_link notes on_plasmid'.split(' '))
    return new_df


def fill_meta_table(meta_df, outtable, db_ids_seqs_df, the_run_date, output):
    database_ids_list = db_ids_seqs_df['database_id'].tolist()
    old_products_list = meta_df['product'].tolist()
    new_products_list = []
    db_ids_seqs_df = db_ids_seqs_df.set_index('database_id')
    #print(db_ids_seqs_df)
    lines_to_append = []
    ### iterate over database IDs
    for db_id in database_ids_list:
        ### example https://mibig.secondarymetabolites.org/repository/BGC0000322/BGC0000322.gbk
        download_link = 'https://mibig.secondarymetabolites.org/repository/%s/%s.gbk' % (db_id, db_id)
        download_file = os.path.join(output, '%s.gbk' %(db_id))
        ### download file
        if not os.path.exists(download_file):
            execute_download(download_link, download_file)
        ### skip ids already in the list
        if meta_df['Database_ID'].str.contains(db_id).any():
            print("Warning: entry %s already in table. Skipping..." %(db_id))
            continue
        ### else prepare line
        #'strain ID source NCBI_AccNo Database_ID seq_with_Xdom product genus '
        #'date_of_search download_link'
        seq_with_xdom = db_ids_seqs_df.loc[db_id]['seq_with_Xdom']
        ### example https://mibig.secondarymetabolites.org/repository/BGC0000322/annotations.json
        json_download_link = 'https://mibig.secondarymetabolites.org/repository/%s/annotations.json' %(db_id)
        strainname, product, genus = get_info_from_json(json_download_link, output, db_id)
        new_products_list.append(product)
        ### tidy up all info in a list with the correct column format
        db_id_list = [strainname, '', 'MIBiG', '', db_id, seq_with_xdom, product, genus, the_run_date,
                      download_link, '']
        lines_to_append.append(db_id_list)
    ### assign IDs to existing and new entries
    meta_df_table = meta_df.values.tolist()
    out_meta_df = assign_ids(lines_to_append, old_products_list, new_products_list, meta_df_table)
    #out_meta_df = pd.concat([meta_df, new_df])
    ### save to file
    out_meta_df.to_csv(outtable, index=False)
    return out_meta_df

# Main function #
def main(input, output, outtable):
    print("\nHmmsearch domain hits table: %s\n"
          "Output folder: %s\n"
          "Output table with files metadata:%s\n"
          %(input, output, outtable))
    if not "/" in outtable:
        print("Writing metadata table in output directory, since no full path was specified\n")
        outtable = os.path.join(output, outtable)
    ### import table of hmmsearch hits
    domhits_df, the_run_date = import_hmmsearch_tblout(input)
    ### get database ids and sequence ids
    domhits_df['database_id'] = domhits_df.target_name.str[:10]
    domhits_df['seq_with_Xdom'] = domhits_df.target_name.str.split("|").str[-1]
    ### read metadate table if it exists, otherwise create it
    if os.path.exists(outtable):
        meta_df = pd.read_csv(outtable)
    else:
        meta_df = pd.DataFrame(columns='strain ID source NCBI_AccNo Database_ID seq_with_Xdom product genus '
                                       'date_of_search download_link notes'.split(' '))
    ### extract all info
    fill_meta_table(meta_df, outtable, domhits_df[['database_id','seq_with_Xdom']], the_run_date, output)
    print("\nDone")


# Parsing arguments & Execution #
parser = argparse.ArgumentParser(description='Downloads all related files based on HMMsearch of MIBiG and creates a'
                                             ' table with all info')
parser.add_argument('-i', '--input', help='Input: Table (domtblout) of HMMsearch dom xits (assumed filtered by threshold)', required=True)
parser.add_argument('-o', '--output', help='Output folder where the files will be downloaded', required=True)
parser.add_argument('-t', '--table', help='Output table (csv) with all gathered information', required=True)
if __name__ == "__main__":
    args = parser.parse_args()
    main(args.input, args.output, args.table)
```
