## Supplementary Data 5 for "Structural and evolutionary analyses support reclassification of glycopeptide antibiotics as xyclopeptides": extract_domain_seqs.py

```
#!/usr/bin/python3.6
# Created by Athina Gavriilidou at 07.09.23 14:44 #
# Imports #
import argparse
import os
from Bio import SeqIO
from pprint import pprint
from Bio.SeqRecord import SeqRecord
from Bio.Seq import Seq
from Bio import motifs
from Bio import AlignIO
from Bio.Blast.Applications import NcbiblastpCommandline
import re
from collections import OrderedDict
import csv
import itertools
import pandas as pd
from Bio import pairwise2
from tqdm import tqdm
import numpy as np


# Functions #
def create_folder(path):
    if not os.path.exists(path):
        os.mkdir(path)
        print("Created folder: %s" %(path))

'''given a directory, lists all files, with a certain suffix'''
def list_files(directory, suffix= "", suppress=False):
    the_files = []
    for file in os.listdir(directory):
        if file.endswith(suffix):
            the_files.append(os.path.join(directory, file))
    if not suppress:
        print("Found %s %s files" %(len(the_files), suffix))
    the_files.sort()
    return the_files

'''given a table of info and a list of NRPS features, removes the ones not corresponding to core biosynthetic NRPS'''
def filter_out_extragenes(nrps_features, nrps_data_table, protocore_locations):
    toremove_locustags = []
    protocorestart, protocoreend = protocore_locations
    for i in range(len(nrps_features)):
        nrpsline = nrps_data_table[i]
        startloc, endloc = nrpsline[2], nrpsline[3]
        if not (startloc >= protocorestart and endloc <= protocoreend):
            toremove_locustags.append(nrpsline[1])
    new_nrps_features = []
    new_nrps_table = []
    #pprint(toremove_locustags)
    if len(toremove_locustags) > 0:
        for locustag in toremove_locustags:
            for nrpsline in nrps_data_table:
                if nrpsline[1] == locustag:
                    continue
                new_nrps_table.append(nrpsline)
            for nrpsfeature, cd_num  in nrps_features:
                #nrpsfeature = nrpsfeature[0]
                if nrpsfeature.qualifiers['locus_tag'][0] == locustag:
                    continue
                new_nrps_features.append(nrpsfeature)
        return new_nrps_features, new_nrps_table
    else:
        return nrps_features, nrps_data_table

'''assumes structure of table, checks if all entries of 4th column are the same'''
def check_if_all_genes_same_strand(nrps_data_table):
    latest_entry = False
    for nrpsline in nrps_data_table:
        if not latest_entry:
            latest_entry =nrpsline[3]
        elif latest_entry != nrpsline[3]:
            return False
    return latest_entry

'''split table with assumed structure into two categories'''
def split_into_groups(nrps_data_table):
    positive_group = []
    negative_group = []
    for nrpsline in nrps_data_table:
        if nrpsline[4] == 1:
            positive_group.append(nrpsline)
        elif nrpsline[4] == -1:
            negative_group.append(nrpsline)
        else:
            print('ERROR: uknown value where positive (1) or negative (-1) strand should be. Aborting...')
            print(nrpsline[4])
            pprint(nrps_data_table)
            exit()
    groups_dict = {1: positive_group, -1:negative_group}
    return groups_dict

'''quick and dirty method to detect X domain in gene based on Domain info'''
def checkifhasXdom(gene_domains_info):
    for el in gene_domains_info:
        if " X " in el:
            return True
    return False

'''fix order of NRPS genes as their modules would be'''
def reorder_based_onXdom(nrps_data_table, xdom_nrps_locus_tag):
    new_nrps_data_table = []
    for nrpsline in nrps_data_table:
        if nrpsline[1] == xdom_nrps_locus_tag:
            xdomnrpsline = nrpsline
            xdomstrand = nrpsline[4]
    #print(nrpsline, xdom_nrps_locus_tag)
    groups_dict = split_into_groups(nrps_data_table)
    lastgroup = groups_dict[xdomstrand]
    for strand, nrpssubtable in groups_dict.items():
        if strand != xdomstrand:
            firstgroup = nrpssubtable
            break
    if xdomstrand == -1:
        for line in firstgroup:
            new_nrps_data_table.append(line)
        for i in range(len(lastgroup)-1, -1, -1):
            line = lastgroup[i]
            new_nrps_data_table.append(line)
    else:
        for i in range(len(firstgroup)-1, -1, -1):
            line = firstgroup[i]
            new_nrps_data_table.append(line)
        for line in lastgroup:
            new_nrps_data_table.append(line)
    return new_nrps_data_table

def filter_protocore_locations(protocore_locations, gbkfile):
    records = list(SeqIO.parse(gbkfile, 'genbank'))
    for seq_record in records:
        for feature in seq_record.features:
            if feature.type == "aSDomain" and "X" in feature.qualifiers['aSDomain']:
                xleftlim = feature.location._start.position
                xrightlim = feature.location._start.position
                break
        try:
            bothlim = [xleftlim, xrightlim]
            break
        except:
            continue
    for protocore_candidate in protocore_locations:
        if xleftlim > protocore_candidate[0] and xrightlim < protocore_candidate[1]:
            return [protocore_candidate]

def check_for_inactive_module(locustag_to_modulefeatures_dict):
    inactive_module = False
    for locustag, modulefeaturelist in locustag_to_modulefeatures_dict.items():
        for modulefeature in modulefeaturelist:
            if 'final_module' in modulefeature.qualifiers.keys() and 'incomplete' in modulefeature.qualifiers.keys():
                domainlist = modulefeature.qualifiers['domains']
                for dom in domainlist:
                    if 'AMP-binding' in dom:
                        suffix = dom[-1]
                        break
                inactive_module = [locustag, suffix]
    return inactive_module

def reorder_based_onNRPStable(domtag_to_feature_dict, nrps_data_table, locustag_to_modulefeatures_dict):
    reordered_domtag_to_feature_dict=OrderedDict()
    inactive_module = check_for_inactive_module(locustag_to_modulefeatures_dict)
    for nrpsrow in nrps_data_table:
        nrpslocustag = nrpsrow[1]
        nrps_doms_dict = {}
        for domtag, featurelist in domtag_to_feature_dict.items():
            if nrpslocustag in domtag:
                nrps_doms_dict[domtag] = featurelist
        #nrps_doms_dict = {key:value for (key, value) in domtag_to_feature_dict.items() if nrpslocustag in key}
        max_suffix = len(nrps_doms_dict.keys())
        for i in range(max_suffix):
            expected_suffix = str(i+1)
            for domtag, featurelist in nrps_doms_dict.items():
                suffix = domtag[-1]
                ### exclude inactive a domain from this
                if inactive_module is not False and (nrpslocustag == inactive_module[0] and suffix == inactive_module[1]):
                    if 'inactive' not in featurelist:
                        featurelist.append('inactive')
                        nrps_doms_dict[domtag] = featurelist
                if nrpslocustag in domtag and expected_suffix == suffix:
                    reordered_domtag_to_feature_dict[domtag] = featurelist
                    expected_suffix = str(int(expected_suffix) + 1)
                    break
                #print(expected_suffix)
                #break
    return reordered_domtag_to_feature_dict

def execute_needle(faafile, knownstachfile, outmsafile):
    needlepath = '/usr/bin/needle'
    print("Assuming path to EMBOSS needle: %s" % (needlepath))
    needle_command = "%s -asequence %s -bsequence %s -outfile %s" %(needlepath, faafile, knownstachfile, outmsafile)
    print(needle_command)
    try:
        os.system(needle_command)
    except:
        print("\nERROR: could not run EMBOSS needle. Are you sure it is installed and in the path?\n")
        exit()

def execute_blastp(domfasta_file, dbfile, outfile):
    blast_path = '/home/athina/anaconda3/envs/zol_env/bin/blastp'
    #blastcommand = NcbiblastpCommandline(query=domfasta_file, db=manual_adom_annotation_fasta)
    blastcommand = '%s -query %s -db %s -out %s' %(blast_path, domfasta_file, dbfile, outfile)
    print(blastcommand)
    try:
        os.system(blastcommand)
    except:
        print("\nERROR: could not run blastp. Are you sure it is installed and in the path?\n")
        exit()

def make_blastp_db(manual_adom_annotation_fasta, dbfile):
    blast_path = '/home/athina/anaconda3/envs/zol_env/bin/makeblastdb'
    # blastcommand = NcbiblastpCommandline(query=domfasta_file, db=manual_adom_annotation_fasta)
    blastcommand = '%s -in %s -dbtype prot -parse_seqids -out %s' % (blast_path,manual_adom_annotation_fasta, dbfile)
    print(blastcommand)
    try:
        os.system(blastcommand)
    except:
        print("\nERROR: could not run blastp. Are you sure it is installed and in the path?\n")
        exit()

def featuredict_to_fasta(domtag_to_feature_dict, thesedoms_fasta, overwrite=False):
    if not os.path.exists(thesedoms_fasta) or overwrite:
        output_handle = open(thesedoms_fasta, "w")
        for header, featurelist in domtag_to_feature_dict.items():
            seq = featurelist[1]
            output_handle.write(">%s\n%s\n" % (header.replace("nrpspksdomains_",""), seq))
            #print(header, seq)
        output_handle.close()

def parse_blastpresult(dom_blastp_resultfile, domtag_to_feature_dict):
    #domtag_to_stach_match_dict = {}
    current_query = False
    countnow = False
    linecounter = 0
    with open(dom_blastp_resultfile, 'r') as infile:
        for line in infile:
            line = line.rstrip()

            if countnow and linecounter != 5:
                linecounter +=1
                continue
            if line.startswith('Query= '):
                current_query = line.split(" ")[1]
                linecounter = 0
                countnow = True
                #print(line)
            if countnow and linecounter == 5:
                fullname = line.split(" ")[0]
                stachmatch = fullname.split("|")[-1]
                stachmatch = stachmatch.split(":")[-1]
                score = line[70:74]
                evalue = line[78:83]
                #domtag_to_stach_match_dict[current_query] = stachmatch
                featurelist = domtag_to_feature_dict[current_query] + [stachmatch, score, evalue]
                #featurelist.append(stachmatch)
                domtag_to_feature_dict[current_query] = featurelist
                countnow = False
                #print(line)
    ### now add info to feature dict - or do it directly above
    #for domtag, featurelist in domtag_to_feature_dict.items():
    #    print(featurelist)
    #if "StsW2_2" in current_query:
    #    print()
    return domtag_to_feature_dict

def fix_backbones(domtag_to_feature_dict, dbfile, stachelhaus_alignments_dir, domfastaname, overwrite=False):
    #known_stach_codes = manual_adom_annotation_table['structure_based_specificity'].tolist()
    #known_stach_codes = manual_adom_annotation_table['stach_aa34'].tolist()
    ### create the fasta file for these domains
    thesedoms_fasta = os.path.join(stachelhaus_alignments_dir,f"adom_34stach_{domfastaname}")
    featuredict_to_fasta(domtag_to_feature_dict, thesedoms_fasta, overwrite=False)
    dom_blastp_resultfile = os.path.join(stachelhaus_alignments_dir, f'{domfastaname.split("__")[0]}_stach34_blastp_out.txt')
    #print(dom_blastp_resultfile)
    if not os.path.exists(dom_blastp_resultfile) or overwrite:
        execute_blastp(thesedoms_fasta, dbfile, dom_blastp_resultfile)
    ### now get results from blastp file
    domtag_to_feature_dict = parse_blastpresult(dom_blastp_resultfile, domtag_to_feature_dict)
    return domtag_to_feature_dict

def df_to_fasta(manual_adom_annotation_table,outfaafile,
                headercol,seq_col, unique_headers=True, overwrite=False):
    if not os.path.exists(outfaafile) or overwrite:
        output_handle = open(outfaafile, "w")
    else:
        return outfaafile
    loc_to_seq_dict = manual_adom_annotation_table[seq_col].to_dict()
    if not unique_headers:
        seq_index = 1
    header_to_seq_dict = {}
    for loc, seq in loc_to_seq_dict.items():
        header = manual_adom_annotation_table.loc[loc, headercol]
        if not unique_headers:
            header = f"{seq_index}|{header}"
            seq_index += 1
        header_to_seq_dict[header] = seq
    for header, seq in header_to_seq_dict.items():
        output_handle.write(">%s\n%s\n" % (header,seq))
    output_handle.close()
    return outfaafile


'''parses gbk file and identifies all necessary NRPS information'''
def get_nrps_data(gbkfile, manual_adom_annotation_table, manual_adom_annotation_fasta, stachelhaus_alignments_dir,
                  overwrite=False):
    ### check first that manual_adom_annotation_table is turned into fasta file
    if not os.path.exists(manual_adom_annotation_fasta) or overwrite:
        df_to_fasta(manual_adom_annotation_table, manual_adom_annotation_fasta,
                                                   headercol='structure_based_specificity',
                                                    seq_col='stach_aa34', unique_headers=False)
    ### make it into blast db
    dbfile = os.path.join(stachelhaus_alignments_dir, 'known_adom_stach34.db')
    if not os.path.exists(f"{dbfile}.pdb") or overwrite:
        make_blastp_db(manual_adom_annotation_fasta, dbfile)
    records = list(SeqIO.parse(gbkfile, 'genbank'))
    if len(records)>1:
        print('ERROR: BGC %s expected to have one record. Has multiple. Aborting...' %(os.path.basename(gbkfile)))
        exit()
    nrps_features = []
    protocore_locations = []
    xdom_nrps_locus_tag = ''
    domtag_to_feature_dict = {}
    x_dom_loc = []
    ### for better accuracy, start by detecting the X domain location and use to decect correct protocore
    for seq_record in records:
        if len(x_dom_loc) > 0:
            break
        for feature in seq_record.features:
            if feature.type == "aSDomain" and "X" in feature.qualifiers['aSDomain']:
                x_dom_loc = [int(feature.location.start), int(feature.location.end)]
                xdom_nrps_locus_tag = feature.qualifiers['locus_tag'][0]
                break
    if xdom_nrps_locus_tag == '':
        print("\nERROR: X domain locus detection failed for file: %s. \nAborting..." % (gbkfile))
        exit()
    nrps_locustags_list = []
    for seq_record in records:
        fnaseq = seq_record.seq
        cd_counter = 0
        for feature in seq_record.features:
            if feature.type == "CDS":
                cd_counter+=1
            elif feature.type == "proto_core" and "NRPS" in feature.qualifiers['product']:
                thislocstart = int(feature.location.start)
                thislocend = int(feature.location.end)
                if thislocstart < x_dom_loc[0] and thislocend > x_dom_loc[1]:
                    protocore_locations.append([thislocstart, thislocend])
            if (feature.type == "CDS" and "gene_kind" in list(feature.qualifiers.keys())
                    and "biosynthetic" in feature.qualifiers["gene_kind"] and "NRPS_PKS" in feature.qualifiers.keys()):
                #print(feature)
                nrps_features.append([feature, cd_counter])
                nrps_locustags_list.append(feature.qualifiers['locus_tag'][0])
            ### keep also domains
            if feature.type == "aSDomain" and "AMP-binding" == feature.qualifiers['aSDomain'][0]:
                try:
                    domtag_to_feature_dict[feature.qualifiers['domain_id'][0].replace("nrpspksdomains_","")] = [feature.qualifiers['stach_aa10'][0],
                                                                                  feature.qualifiers['stach_aa34'][0],
                                                                                  feature.qualifiers['aa10_score'][0],
                                                                                  feature.qualifiers['aa34_score'][0],
                                                                                  feature.qualifiers['most_likely_substrate'][0]]
                    #if "StsW2_2" in feature.qualifiers['domain_id'][0]:
                    #    print()
                ### some willnot have that info
                except:
                    continue

    ### 17.01.24 additional check for specificities based on known structures - should be done as soon as we have the stach codes
    #print(f"Fixing backbones for file {os.path.basename(gbkfile)} ...")
    domtag_to_feature_dict = fix_backbones(domtag_to_feature_dict, dbfile, stachelhaus_alignments_dir,
                                           os.path.basename(gbkfile).replace(".gbk", ".faa"))

    ### check for possible exceptions to the rule:
    if len(protocore_locations) >1:
        print("Warning: More than one NRPS cluster core detected for %s. Choosing based on X domain..."
              %(os.path.basename(gbkfile)))
        protocore_locations = filter_protocore_locations(protocore_locations, gbkfile)

    nrps_data_table = []
    nrps_feat_toremove = []
    for nrpsfeature, cd_num in nrps_features:
        startloc = int(nrpsfeature.location.start)
        endloc = int(nrpsfeature.location.end)
        locustag = nrpsfeature.qualifiers['locus_tag'][0]
        ### added on 22.10.2023 skip if not in protocore
        if not (startloc >= protocore_locations[0][0] and endloc <= protocore_locations[0][1]):
            nrps_feat_toremove.append(cd_num)
            continue
        strand = nrpsfeature.location.strand
        nrpsdata = [cd_num, locustag, startloc, endloc, strand]
        nrps_data_table.append(nrpsdata)
    ### remove filtered out based on protocore 22.20.2023
    nrps_features = [x for x in nrps_features if x[1] not in nrps_feat_toremove]

    ### here check if there is an NRPS that is not core biosynthetic (e.g. bpsD)
    nrps_features, nrps_data_table = filter_out_extragenes(nrps_features, nrps_data_table, protocore_locations[0])
    ### if the genes all have the same direction, the last one is the one containing the X domain, reordering here.
    nrps_data_table = reorder_based_onXdom(nrps_data_table, xdom_nrps_locus_tag)
    ### get module information
    locustag_to_modulefeatures_dict = {}
    for seq_record in records:
        for feature in seq_record.features:
            if feature.type == "aSModule":
                #if "StsW2_2" in feature.qualifiers['domains'][0]:
                #    print()
                ### extra check that we are focusing on the protocore 22.10.2023
                locationstart = feature.location._start
                locationend = feature.location._end
                if not (locationstart >= protocore_locations[0][0] and locationend < protocore_locations[0][1]):
                    continue
                thislocustag = feature.qualifiers['locus_tags'][0]
                if thislocustag in nrps_locustags_list:
                    if not thislocustag in locustag_to_modulefeatures_dict.keys():
                        locustag_to_modulefeatures_dict[thislocustag] = [feature]
                    else:
                        current_list = locustag_to_modulefeatures_dict[thislocustag]
                        if feature.location.strand == 1:
                            current_list.append(feature)
                        else:
                            current_list = [feature] + current_list
                        locustag_to_modulefeatures_dict[thislocustag] =current_list
    module_count = 0
    ### get full AA backbone from modules - here stachelhaus code results
    backbone_list = []

    for nrpsline in nrps_data_table:
        locustag = nrpsline[1]
        ### special case for avoparcin, may delete later
        if locustag == 'avo_00160':
            continue
        if locustag == 'UME_00135':
            print()
        try:
            modules = locustag_to_modulefeatures_dict[locustag]
        except:
            print("ERROR: locus tag not found: %s" %(locustag))
            pprint(locustag_to_modulefeatures_dict)
            exit()
        for i in range(len(modules)):
            m = modules[i]
            try:
                adom_id = [x.replace('nrpspksdomains_','') for x in m.qualifiers['domains'] if 'AMP-binding' in x][0]
            except:
                print("Warning: No AMP-binding domain found in a module of gene %s" %(locustag))
                adom_id = ''
            ### no monomer pairings in inactive A-domain
            if "monomer_pairings" in m.qualifiers.keys():
                module_count += 1
                ### 25.01.24 to replace with new Stachelhaus approach - taking blastp best match
                res, score, evalue = domtag_to_feature_dict[adom_id][5:8]
                m.qualifiers['stach_match'] = [res,score,evalue]
                backbone_list.append(res)
            modules[i] = m
        locustag_to_modulefeatures_dict[locustag] = modules
    #'''

    ### reorder also domtag_to_feature_dict based on nrps table
    domtag_to_feature_dict = reorder_based_onNRPStable(domtag_to_feature_dict, nrps_data_table, locustag_to_modulefeatures_dict)
    backbone_stach_list = []
    for domtag, featurelist in domtag_to_feature_dict.items():
        if 'inactive' not in featurelist:
            backbone_stach_list.append(featurelist[1])
        #exit()
    #print(backbone_list)
    ### here check if backbone length is as expected
    unexpected = False
    if module_count != 7:
        unexpected = module_count
    return (nrps_data_table, locustag_to_modulefeatures_dict, backbone_list, fnaseq, unexpected, domtag_to_feature_dict,
            backbone_stach_list)

def transform_res_list(backbone_list):
    res_to_letter = {'Tyr': 'Y', 'Val':'V','X':'X','Hpg':'H','Leu':'L','bOH-Tyr':'B','Asn':'N',
                     'Phe':'F','Dhpg':'R','Trp':'W','Asp':'D', 'Thr':'T', 'Ser':'S', 'Pro':'P',
                     'Ala':'A', 'Glu':'G'}
    l_str = ''
    for res in backbone_list:
        try:
            l_str = "%s%s" %(l_str, res_to_letter[res])
        except:
            continue
    return l_str

def backbones_to_fasta(backbones_table, outfaa):
    sequences_to_write = []
    with open(outfaa, "w") as output_handle:
        for line in backbones_table:
            record = SeqRecord(
                Seq(line[1]),
                id=line[0],
                name=line[0],
                description=line[0],
            )
            sequences_to_write.append(record)
        SeqIO.write(sequences_to_write, output_handle, "fasta")

def execute_quick_mafft(inputfasta, outputfasta):
    #mafft_command = 'mafft --auto %s > %s' %(inputfasta, outputfasta)
    mafft_command = 'mafft  --leavegappyregion --auto %s > %s' %(inputfasta, outputfasta)
    print(mafft_command)
    try:
        os.system(mafft_command)
    except:
        print("Warning. There was a problem with running mafft on the file: %s" %(inputfasta))

def execute_weblogo(inputfasta, outputpng):
    seqlogopath = ('/media/hdd_2tb/GPA_reclassification_clean/A_dataset_creation/4_orthology_inference/weblogo.2.8.2'
                   '/weblogo/seqlogo')
    print("\nNote: hardcoded seqlogo path")
    titletext = 'bOH-Tyr=B Dhpg=R Hpg=H'
    weblogo_command1 = '%s -c -k 0 -n -Y -p -M -w 30 -h 10 -F PNG -f %s > %s' %(seqlogopath, inputfasta, outputpng)
    weblogo_command2 = ('%s -c -k 0 -n -t \'%s\' -Y -a -p -M -w 30 -h 10 -F PNG -f %s > %s' %
                        (seqlogopath, titletext, inputfasta, outputpng[:-4] + '_a.png'))
    weblogo_command3 = ('%s -c -k 0 -n -Y -a -p -M -w 30 -h 10 -F EPS -f %s > %s' %
                        (seqlogopath, inputfasta, outputpng[:-4] + '_e.eps'))
    try:
        os.system(weblogo_command1)
        os.system(weblogo_command2)
        os.system(weblogo_command3)
    except:
        print("Warning. There was a problem with creating a weblogo out of the file: %s" %(inputfasta))

def simplify_msa(simplified_msa_file, backbones_aligned_fasta, backbones_table):
    stach_len = 34
    backbones_dict = {}
    simplified_recordlist = []
    for row in backbones_table:
        backbones_dict[row[0]] = row[1]
    with open(backbones_aligned_fasta) as inhandle:
        for record in SeqIO.parse(inhandle, "fasta"):
            bgcid = record.id
            msaseq = str(record.seq)
            postitions = [msaseq[i:i+stach_len] for i in range(0,len(msaseq),stach_len)]
            simplified_seq =backbones_dict[bgcid]
            simplified_msa_seq = ''
            msa_index = 0
            for res in simplified_seq:
                added = False
                while not added:
                    #if not '-' in postitions[msa_index]:
                    if postitions[msa_index].count('-') < 30:
                        simplified_msa_seq = simplified_msa_seq + res
                        added = True
                    else:
                        simplified_msa_seq = simplified_msa_seq + '-'
                    msa_index += 1
            while msa_index != len(postitions):
                simplified_msa_seq = simplified_msa_seq + '-'
                msa_index +=1
            simplified_recordlist.append([bgcid, simplified_msa_seq])
    ### now save to file
    output_handle = open(simplified_msa_file, "w")
    for rec in simplified_recordlist:
        output_handle.write(">%s\n%s\n" % (
            # seq_feature.qualifiers['locus_tag'][0],
            rec[0],
            #seq_feature.location,
            # bgc_filename,
            # seq_record.name,
            rec[1]))
    print(f"Created file: {simplified_msa_file}")

def return_longest_nongap(msa_file):
    gapstring = '-'
    colloc_to_gaps = OrderedDict()
    with open(msa_file, 'r') as inhandle:
        for record in SeqIO.parse(inhandle, "fasta"):
            msaseq = str(record.seq)
            for i in range(len(msaseq)):
                char = msaseq[i]
                if char == gapstring:
                    if i not in colloc_to_gaps.keys():
                        colloc_to_gaps[i] = 1
                    else:
                        colloc_to_gaps[i] = colloc_to_gaps[i] +1
    for i in range(len(msaseq)):
        if i not in colloc_to_gaps.keys():
            colloc_to_gaps[i] = 0
    ### keep only columns with no gaps at all
    mingap_locs = []
    mingap = min(colloc_to_gaps.values())
    for loc, gapcount in colloc_to_gaps.items():
        if gapcount == mingap:
            mingap_locs.append(loc)
    ### find longest incremental sublist
    start_loc = mingap_locs[0]
    previous_loc = mingap_locs[0]
    list_of_candidates = []
    if len(mingap_locs) == 1:
        return [mingap_locs[0], mingap_locs[0]]
    for loc in mingap_locs[1:]:
        if loc != previous_loc+1:
            list_of_candidates.append([start_loc, previous_loc])
            start_loc = loc
        previous_loc = loc
        if loc == mingap_locs[-1]:
            list_of_candidates.append([start_loc, previous_loc])
    len_to_loc_dict = {}
    for start, end in list_of_candidates:
        thelen = end - start +1
        len_to_loc_dict[thelen] = [start, end]
    best_candidate = len_to_loc_dict[max(len_to_loc_dict.keys())]
    print(f"The longest subsequence of most conserved positions with {mingap} gaps in the MSA were positions"
          f" {best_candidate[0]+1} to {best_candidate[1]+1}. \nNote: These positions will be used as the core. "
          f"All internal gaps will be removed, as no gaps can exist in the structure of the compounds.")
    return best_candidate

def fix_msa_file(simplified_msa, conserved_loc, fixed_msa_file):
    fixed_recordlist = []
    theconservedstart = conserved_loc[0]
    theconservedend = conserved_loc[1]
    with open(simplified_msa) as inhandle:
        for record in SeqIO.parse(inhandle, "fasta"):
            msaseq = str(record.seq)
            start_gaps_count = msaseq[:theconservedstart].count("-")
            start_gaps = "-"*start_gaps_count
            startseq = msaseq[:theconservedstart].replace("-","")
            if theconservedend != len(msaseq)-1:
                end_gaps_count = msaseq[theconservedend+1:].count("-")
                end_gaps = "-"*end_gaps_count
                endseq = msaseq[theconservedend+1:].replace("-","")
            else:
                end_gaps = ''
                endseq = ""
            fixed_seq = start_gaps + startseq + msaseq[theconservedstart:theconservedend+1] + endseq + end_gaps
            fixed_recordlist.append([record.id, fixed_seq])
    ### check if any columns now have only gaps, and remove them if so
    passed = []
    for id, msaseq in fixed_recordlist:
        for i in range(len(msaseq)):
            if i not in passed:
                char = msaseq[i]
                if char != '-':
                    passed.append(i)
    gaponly_cols = [x for x in range(len(msaseq)) if x not in passed]
    if len(gaponly_cols) > 0:
        for j in range(len(fixed_recordlist)):
            id, msaseq = fixed_recordlist[j]
            charlist = [msaseq[i] for i in range(len(msaseq)) if i not in gaponly_cols]
            msaseq = "".join(charlist)
            fixed_recordlist[j] = [id, msaseq]
    print(f"Note: after fixing the length of the MSA is: {len(msaseq)}")
    ### now save to file
    output_handle = open(fixed_msa_file, "w")
    for rec in fixed_recordlist:
        output_handle.write(">%s\n%s\n" % (
            # seq_feature.qualifiers['locus_tag'][0],
            rec[0],
            # seq_feature.location,
            # bgc_filename,
            # seq_record.name,
            rec[1]))
    print(f"Created file: {fixed_msa_file}")

def run_backbone_analysis(backbone_dir, backbones_stach_table, backbones_table):
    outbackbones_simple_fasta = os.path.join(backbone_dir, 'backbones_stach34concat.faa')
    if not os.path.exists(outbackbones_simple_fasta):
        backbones_to_fasta(backbones_stach_table, outbackbones_simple_fasta)
    ### make quick alignment
    backbones_aligned_fasta = "%s_mafftMSA.faa" % (outbackbones_simple_fasta[:-4])
    if not os.path.exists(backbones_aligned_fasta):
        execute_quick_mafft(outbackbones_simple_fasta, backbones_aligned_fasta)
    ### get simplified letters out of stachelhaus code alignment
    simplified_msa = "%s_mafftMSA_simplified.faa" % (outbackbones_simple_fasta[:-4])
    simplify_msa(simplified_msa, backbones_aligned_fasta, backbones_table)
    ### make weblogo out of alignment
    outlogo = os.path.join(backbone_dir, 'logo.png')
    execute_weblogo(simplified_msa, outlogo)
    ### find location of longest conserved (non gappy) region) and fix MSA file accordingly
    conserved_loc = return_longest_nongap(simplified_msa)
    fixed_msa_file = "%s_mafftMSA_simplified_fixed.faa" % (outbackbones_simple_fasta[:-4])
    fix_msa_file(simplified_msa, conserved_loc, fixed_msa_file)
    ### make weblogo out of fixed alignment
    outlogo = os.path.join(backbone_dir, 'fixed_logo.png')
    execute_weblogo(fixed_msa_file, outlogo)
    #exit()
    return fixed_msa_file

def save_moduleinfo_to_csv(bgcid_to_moduleinfotable_dict,moduleinfo_table_cols,outcsv):
    with open(outcsv, 'w') as f:
        csv_output = csv.writer(f)
        csv_output.writerow(['bgcid'] + moduleinfo_table_cols)
        for bgcid, moduletable in bgcid_to_moduleinfotable_dict.items():
            for line in moduletable:
                line = [bgcid] + line
                csv_output.writerow(line)
    print("\nCreated file: " + outcsv)

def save_domaininfo_to_csv(bgcid_to_domainsinfodicts_dict, bgcid_to_domainsinfo_cols, outcsv):
    with open(outcsv, 'w') as f:
        csv_output = csv.writer(f)
        csv_output.writerow(['bgcid'] + bgcid_to_domainsinfo_cols)
        for bgcid, domaintypedict in bgcid_to_domainsinfodicts_dict.items():
            for domaintype, domaintable in domaintypedict.items():
                for line in domaintable:
                    line = [bgcid] + line
                    csv_output.writerow(line)
    print("\nCreated file: " + outcsv)

def save_table_to_csv(table_list, outcsv):
    with open(outcsv, 'w') as f:
        csv_output = csv.writer(f)
        for line in table_list:
            csv_output.writerow(line)
    print("\nCreated file: " + outcsv)

def extract_module_seqs(modules_dir, bgcid_to_moduleinfotable_dict, bgc_to_fnaseq_dict):
    modules_faas_dir = os.path.join(modules_dir, 'modules_faas')
    all_modules_faa = os.path.join(modules_dir, 'all_modules.faa')
    create_folder(modules_faas_dir)
    modules_fnas_dir = os.path.join(modules_dir, 'modules_fnas')
    all_modules_fna = os.path.join(modules_dir,'all_modules.fna')
    create_folder(modules_fnas_dir)
    filenames_to_fnaseqrecords_dict = {}
    filenames_to_faaseqrecords_dict = {}
    domain_label_to_modulegenecount_dict = {}
    ### iterate over modules
    for bgcid, moduleinfotable in bgcid_to_moduleinfotable_dict.items():
        bgcid_dnaseq = bgc_to_fnaseq_dict[bgcid]
        gene_number = len(set([x[0] for x in moduleinfotable]))
        gene_number_letter = chr(ord('@')+ gene_number)
        maxmodules = len(moduleinfotable)
        gene_count = 1
        current_gene = moduleinfotable[0][0]
        current_gene_letter = chr(ord('@')+ gene_count)
        within_gene_mod_count = 1
        mod_count = 1
        #if "StsW2_2" in bgcid:
        #    print()
        for moduleinfo in moduleinfotable:
            #print(moduleinfo)
            startpos = moduleinfo[2]
            endpos = moduleinfo[3]
            strand = moduleinfo[4]
            ### normal counting for complete ones
            if moduleinfo[7] == 'yes':
                module_index_str = str(moduleinfo[1])
            ### annotating "e" for extra for incomplete ones
            else:
                module_index_str = 'e'
            if module_index_str == '-1':
                module_index_str = '00'
            module_faa_outfile = os.path.join(modules_faas_dir, 'modules_%s.faa' %(module_index_str))
            module_fna_outfile = os.path.join(modules_fnas_dir, 'modules_%s.fna' %(module_index_str))

            genelocustag = moduleinfo[0]
            if genelocustag != current_gene:
                gene_count+=1
                current_gene = genelocustag
                current_gene_letter = chr(ord('@')+ gene_count)
                within_gene_mod_count = 1
            within_gene_mod_letter = chr(ord('@')+ within_gene_mod_count).lower()
            ### save info for domains too
            for domainlabel in moduleinfo[6].split('|'):
                domainlabel = domainlabel.replace("nrpspksdomains_","")
                domainlabel = domainlabel.replace(" ","")
                domain_label_to_modulegenecount_dict[domainlabel] = [maxmodules, gene_number_letter,
                                                                     current_gene_letter, within_gene_mod_letter,
                                                                     mod_count, module_index_str]
            moduleheader = '%s__%s%s_%s%s%s' %(genelocustag,maxmodules, gene_number_letter,
                                               current_gene_letter, within_gene_mod_letter, mod_count)
            #print(moduleheader)
            module_fna_seq = bgcid_dnaseq[startpos:endpos]
            fna_module_rec = SeqRecord(module_fna_seq, id=moduleheader,
                                   name=moduleheader,
                                   description=moduleheader)
            if strand == 1:
                module_faa_seq = module_fna_seq.translate()
            ### case of negative strand, we need reverse complement
            else:
                module_fna_seq_compl = module_fna_seq.reverse_complement()
                module_faa_seq = module_fna_seq_compl.translate()
            faa_module_rec = SeqRecord(module_faa_seq, id=moduleheader,
                                       name=moduleheader,
                                       description=moduleheader)
            ### add to records list
            if module_faa_outfile not in filenames_to_faaseqrecords_dict:
                filenames_to_fnaseqrecords_dict[module_fna_outfile] = [fna_module_rec]
                filenames_to_faaseqrecords_dict[module_faa_outfile] = [faa_module_rec]
            else:
                fnarecordlist = filenames_to_fnaseqrecords_dict[module_fna_outfile]
                fnarecordlist.append(fna_module_rec)
                filenames_to_fnaseqrecords_dict[module_fna_outfile] = fnarecordlist
                faarecordlist = filenames_to_faaseqrecords_dict[module_faa_outfile]
                faarecordlist.append(faa_module_rec)
                filenames_to_faaseqrecords_dict[module_faa_outfile] = faarecordlist
                ### increase counters
            within_gene_mod_count+=1
            mod_count+=1
        #break
    with open(all_modules_fna, 'w') as handle:
        domainlist = list(itertools.chain.from_iterable(filenames_to_fnaseqrecords_dict.values()))
        SeqIO.write(domainlist, handle, 'fasta')
    with open(all_modules_faa, 'w') as handle:
        domainlist = list(itertools.chain.from_iterable(filenames_to_faaseqrecords_dict.values()))
        SeqIO.write(domainlist, handle, 'fasta')
    for filenames_to_seqrecords_dict_list in [filenames_to_fnaseqrecords_dict, filenames_to_faaseqrecords_dict]:
        for outfile, domainlist in filenames_to_seqrecords_dict_list.items():
            with open(outfile, 'w') as handle:
                SeqIO.write(domainlist, handle, 'fasta')
    return domain_label_to_modulegenecount_dict

'''for domains we need to find their location info and add it to the dict before extracting the actual sequences'''
def extract_domain_seqs(domains_dir, dtype_to_bgcid_to_domainsinfo_dict, bgc_to_fnaseq_dict,bgc_gbks_list,
                        domain_label_to_modulegenecount_dict, bgcid_to_domainsinfodicts_dict):
    #pprint(domain_label_to_modulegenecount_dict)
    ### get domain info from gbks first
    label_to_info_dict = {}
    for gbkfile in bgc_gbks_list:
        #bgcid = os.path.basename(gbkfile)[:-4]
        records = list(SeqIO.parse(gbkfile, 'genbank'))
        if len(records) > 1:
            print('ERROR: BGC %s expected to have one record. Has multiple. Aborting...' % (os.path.basename(gbkfile)))
            exit()
        for seq_record in records:
            for feature in seq_record.features:
                if feature.type == "aSDomain":
                    if "label" in feature.qualifiers.keys():
                        thislabel = feature.qualifiers['label'][0]
                        thisfaa = feature.qualifiers['translation'][0]
                        label_to_info_dict[thislabel] = [feature.location, thisfaa]
    ### then similar process as for modules, but for each type separately
    for dtype, bgcid_to_domaininfo_dict in dtype_to_bgcid_to_domainsinfo_dict.items():
        dtypestring = dtype.replace("-","_")
        modules_faas_dir = os.path.join(domains_dir, '%s_faas' %(dtypestring))
        all_modules_faa = os.path.join(domains_dir, 'all_%s_domains.faa' %(dtypestring))
        create_folder(modules_faas_dir)
        modules_fnas_dir = os.path.join(domains_dir, '%s_fnas' %(dtypestring))
        all_modules_fna = os.path.join(domains_dir, 'all_%s_domains.fna' %(dtypestring))
        create_folder(modules_fnas_dir)
        filenames_to_fnaseqrecords_dict = {}
        filenames_to_faaseqrecords_dict = {}
        for bgcid, domaininfotable in bgcid_to_domaininfo_dict.items():
            bgcid_dnaseq = bgc_to_fnaseq_dict[bgcid]
            for domaininfo in domaininfotable:
                domainlabel = domaininfo[3]
                #print(domainlabel)
                startpos = int(label_to_info_dict[domainlabel][0].start)
                endpos = int(label_to_info_dict[domainlabel][0].end)
                strand = int(label_to_info_dict[domainlabel][0].strand)
                thisbgcthistypetable = bgcid_to_domainsinfodicts_dict[bgcid][dtype]
                extracounter = 0

                #pprint(thisbgcthistypetable)
                #exit()
                module_index_str = str(domaininfo[1])
                if module_index_str == '-1':
                    module_index_str = '00'

                genelocustag = domaininfo[0]
                #if bgcid == 'StAfJ3_2':
                #    pprint(domain_label_to_modulegenecount_dict)
                maxmodules, gene_number_letter, current_gene_letter, within_gene_mod_letter, mod_count, module_index_str = domain_label_to_modulegenecount_dict[domainlabel]
                moduleheader = '%s__%s%s_%s%s%s' % (genelocustag, maxmodules, gene_number_letter,
                                                    current_gene_letter, within_gene_mod_letter, mod_count)
                bgcid_to_domainsinfodicts_dict[bgcid][dtype] = thisbgcthistypetable
                if dtypestring not in ['X', 'Condensation_Starter', 'Thioesterase']:
                    module_faa_outfile = os.path.join(modules_faas_dir,'%s_domains_%s.faa' % (dtypestring,
                                                                                              module_index_str))
                    module_fna_outfile = os.path.join(modules_fnas_dir,'%s_domains_%s.fna' % (dtypestring,
                                                                                              module_index_str))
                    ### all X, TE and Cstarter domains should be together in one file
                else:
                    module_faa_outfile = os.path.join(modules_faas_dir,'%s_domains.faa' % (dtypestring))
                    module_fna_outfile = os.path.join(modules_fnas_dir,'%s_domains.fna' % (dtypestring))

                ### 26.01.24 moved this loop here to make sure moduleheader is also saved in the domain table
                for domainothertableline in thisbgcthistypetable:
                    if domainlabel == domainothertableline[3]:
                        domainothertableline = domainothertableline + [startpos, endpos, strand, moduleheader]
                        thisbgcthistypetable[extracounter] = domainothertableline
                    extracounter += 1
                # print(moduleheader)
                module_fna_seq = bgcid_dnaseq[startpos:endpos]
                fna_module_rec = SeqRecord(module_fna_seq, id=moduleheader,
                                           name=moduleheader,
                                           description=moduleheader)
                module_faa_seq = label_to_info_dict[domainlabel][1]
                #print(module_faa_seq)
                faa_module_rec = SeqRecord(Seq(module_faa_seq), id=moduleheader,
                                           name=moduleheader,
                                           description=moduleheader)
                ### add to records list
                if module_faa_outfile not in filenames_to_faaseqrecords_dict:
                    filenames_to_fnaseqrecords_dict[module_fna_outfile] = [fna_module_rec]
                    filenames_to_faaseqrecords_dict[module_faa_outfile] = [faa_module_rec]
                else:
                    fnarecordlist = filenames_to_fnaseqrecords_dict[module_fna_outfile]
                    fnarecordlist.append(fna_module_rec)
                    filenames_to_fnaseqrecords_dict[module_fna_outfile] = fnarecordlist
                    faarecordlist = filenames_to_faaseqrecords_dict[module_faa_outfile]
                    faarecordlist.append(faa_module_rec)
                    filenames_to_faaseqrecords_dict[module_faa_outfile] = faarecordlist
            # break
        with open(all_modules_fna, 'w') as handle:
            domainlist = list(itertools.chain.from_iterable(filenames_to_fnaseqrecords_dict.values()))
            SeqIO.write(domainlist, handle, 'fasta')
        with open(all_modules_faa, 'w') as handle:
            domainlist = list(itertools.chain.from_iterable(filenames_to_faaseqrecords_dict.values()))
            SeqIO.write(domainlist, handle, 'fasta')
        for filenames_to_seqrecords_dict_list in [filenames_to_fnaseqrecords_dict, filenames_to_faaseqrecords_dict]:
            for outfile, domainlist in filenames_to_seqrecords_dict_list.items():
                with open(outfile, 'w') as handle:
                    SeqIO.write(domainlist, handle, 'fasta')
    return bgcid_to_domainsinfodicts_dict

def make_itol_formatted_table(domainsouttable):
    colorscheme = {'Leu':'#8fce00','Val':'#ffe599','Trp':'#2986cc','Tyr':'#9fdde8','bOH-Tyr':'#6a329f','Hpg':'#296b51',
                   'Dhpg':'#f44336','Phe':'#b6d7a8','Asn':'#a64d79','unknown':'#999999','-':'#ffffff', np.NaN:'#ffffff',
                   'Glu':'#ce7e00','Ala':'#d5a6bd','Ser':'#f1c232','Asp':'#cc0000','Thr':'#d46217'}
    ### NaN is for the case of StsY_B_AMP-binding.2 which should be labelled inactive but isn't.
    outdir = os.path.dirname(domainsouttable)
    itolouttable = os.path.join(outdir, 'domains_metadata_foriTOL.xlsx')
    domains_df = pd.read_csv(domainsouttable)
    ### fixing here also the issue with the label that iTOL can't handle
    domains_df.replace("StRir_1", "StRirr_1", inplace=True)
    #print(domains_df)
    activeamps_df = domains_df[(domains_df['module_active'] != 'inactive') & (domains_df['type'] == 'AMP-binding')][['bgcid', 'module_index','beststachscore_substrate','beststach_score']]
    activeamps_df.rename(columns={'bgcid':'Tree node ID','beststachscore_substrate':'Label'},inplace=True)

    bgcid_list = activeamps_df[activeamps_df['module_index'] == 4]['Tree node ID'].tolist()
    base_df = pd.DataFrame({'Tree node ID':bgcid_list, 'Tree node label':bgcid_list})
    base_df.set_index('Tree node ID', inplace=True)
    with pd.ExcelWriter(itolouttable) as writer:
    #if True:
        ### going over each position of the backbone
        for i in range(activeamps_df['module_index'].min(), activeamps_df['module_index'].max()+1):
            sheetname = f'backbone_pos_{i}'
            position_df = activeamps_df[activeamps_df['module_index'] == i]
            position_df.set_index('Tree node ID', inplace=True)
            position_df = pd.concat([base_df, position_df[['Label']]], axis=1)
            position_df.fillna('-', inplace=True)
            position_df['Color'] = position_df['Label'].map(colorscheme)
            cols = position_df.columns.tolist()
            cols = cols[:-2] + [cols[-1]] + [cols[-2]]
            position_df = position_df[cols]
            #print(position_df[position_df['text_label']!='-'])
            position_df.to_excel(writer, sheet_name=sheetname, index_label='Tree node ID')
            #break

        ### make sheets also for other domains - start with inactive AMPs
        domain_for_heatmatp_df = domains_df[(domains_df['module_active'] == 'inactive') &
                                            (domains_df['type'] == 'AMP-binding')][['bgcid', 'module_index', 'type']]
        uncommon_domains_df = base_df.copy()
        domain_for_heatmatp_df.rename(columns={'bgcid': 'Tree node ID'}, inplace=True)
        thefieldlabel = "inactive_AMP-binding"
        domain_for_heatmatp_df = domain_for_heatmatp_df.groupby('Tree node ID').count()
        domain_for_heatmatp_df.rename(columns={'module_index': thefieldlabel}, inplace=True)
        uncommon_domains_df = pd.concat([uncommon_domains_df, domain_for_heatmatp_df[[thefieldlabel]]], axis=1)
        uncommon_domains_df.fillna(0, inplace=True)
        ### for C starters also do it separately
        domain_for_heatmatp_df = domains_df[domains_df['type'] == 'Condensation_Starter'][['bgcid',
                                                                                           'module_index', 'type']]
        domain_for_heatmatp_df.rename(columns={'bgcid': 'Tree node ID'}, inplace=True)
        thefieldlabel = "Condensation_Starter"
        domain_for_heatmatp_df = domain_for_heatmatp_df.groupby('Tree node ID').count()
        domain_for_heatmatp_df.rename(columns={'module_index': thefieldlabel}, inplace=True)
        uncommon_domains_df = pd.concat([uncommon_domains_df, domain_for_heatmatp_df[[thefieldlabel]]], axis=1)
        uncommon_domains_df.fillna(0, inplace=True)
        ### for all the other domains, go one by one
        uncommon_domain_list = ['Epimerization','nMT','TIGR01720']
        for domtype in uncommon_domain_list:
            domain_for_heatmatp_df = domains_df[domains_df['type'] == domtype][['bgcid','module_index', 'type']]
            domain_for_heatmatp_df.rename(columns={'bgcid': 'Tree node ID'}, inplace=True)
            ### going over each position of the backbone
            for i in range(-1, 9):
                thisposition_df = domain_for_heatmatp_df[domain_for_heatmatp_df['module_index'] == i]
                if len(thisposition_df) == 0:
                    continue
                thefieldlabel = f"{thisposition_df.iloc[0, 2]}_pos{thisposition_df.iloc[0, 1]}"
                thisposition_df = thisposition_df.groupby('Tree node ID').count()
                thisposition_df.rename(columns={'module_index': thefieldlabel}, inplace=True)
                uncommon_domains_df = pd.concat([uncommon_domains_df, thisposition_df[[thefieldlabel]]], axis=1)
                uncommon_domains_df.fillna(0, inplace=True)
        uncommon_domains_df.to_excel(writer, sheet_name='uncommon_domains_heatmap', index_label='Tree node ID')
    print(f"Created file: {itolouttable}")
    return itolouttable
    #print(activeamps_df)


    # Main function #
def main(input_as_gbks_folder, output, manual_adom_annotation_file = False, replace=False):
    print("\nInput folder with gbk files: %s\n"
          "Output folder: %s\n"
          "Manual A-dom annotation file: %s\n"
          % (input_as_gbks_folder, output, manual_adom_annotation_file))
    create_folder(output)
    bgc_gbks_list = list_files(input_as_gbks_folder, suffix='.gbk')
    print("\nAssumption: the core NRPS genes are either all on the same strand or they are split into two groups, "
          "each on a different strand. This method does not cover the cases of an intermediate NRPS gene being on a "
          "different strand and every other NRPS gene being on the same strand.\n")
    print("Analyzing %s files..." %(len(bgc_gbks_list)))
    ###create folders and declare variables
    backbone_dir =os.path.join(output, '0_backbone_analysis')
    stachelhaus_alignments_dir = os.path.join(backbone_dir, 'stachelhaus_alignments')
    modules_dir = os.path.join(output, '1_modules_analysis')
    domains_dir = os.path.join(output, '2_domains_analysis')
    create_folder(backbone_dir)
    create_folder(stachelhaus_alignments_dir)
    create_folder(modules_dir)
    create_folder(domains_dir)
    total_backbone_res_list = []
    total_backbone_stach_list = []
    tranlsated_backbones_list = []
    backbones_table = []
    backbones_stach_table = []
    bgcid_to_nrspsdatatable_dict = OrderedDict()
    bgcid_to_locustagmodules_dict = {}
    bgc_to_fnaseq_dict = {}
    unexpected_list = []
    warningsfile = os.path.join(output, 'warnings.txt')
    warnings_table = [['###Table of important warnings']]
    bgcid_to_domsdict_dict = {}

    if not manual_adom_annotation_file is False:
        print("Manual A-domain annotation table provided:")
        manual_adom_annotation_table = pd.read_csv(manual_adom_annotation_file, index_col='label',
                                                   usecols=['label','structure_based_specificity',
                                                            'stach_aa10', 'stach_aa34'], sep='\t')
        #print(manual_adom_annotation_table)
    else:
        #if os.path.exists(os.path.join(backbone_dir,'backbone_exceptions_manual.tsv')):
        if os.path.exists(os.path.join(backbone_dir,'adomains_known_specs.csv')):
            manual_adom_annotation_file = os.path.join(backbone_dir,'adomains_known_specs.tsv')
            print("Found manual A-domain annotation file: %s" %(manual_adom_annotation_file))
            manual_adom_annotation_table = pd.read_csv(manual_adom_annotation_file, index_col='label',
                                                       usecols=['label','structure_based_specificity',
                                                                'stach_aa10', 'stach_aa34'], sep='\t')
            #print(manual_adom_annotation_table)
        elif os.path.exists(os.path.join(os.path.dirname(input_as_gbks_folder),'adomains_known_specs.tsv')):
            manual_adom_annotation_file = os.path.join(os.path.dirname(input_as_gbks_folder),'adomains_known_specs.csv')
            print("Found manual A-domain annotation file: %s" % (manual_adom_annotation_file))
            #manual_adom_annotation_table = pd.read_csv(manual_adom_annotation_file, sep='\t', header=None, index_col=0)
            manual_adom_annotation_table = pd.read_csv(manual_adom_annotation_file, index_col='label',
                                                       usecols=['label','structure_based_specificity',
                                                                'stach_aa10', 'stach_aa34'], sep='\t')
            #print(manual_adom_annotation_table)
        else:
            print("Did not detect manual A-domain annotation file and none was provided")
            manual_adom_annotation_table = False
    print(manual_adom_annotation_table)
    ### start iterating over gbks
    for gbkfile in tqdm(bgc_gbks_list):
        #if os.path.basename(gbkfile)[:-4] not in ['a40','bal','a47']:
        #    continue
        #if not os.path.basename(gbkfile)[0].islower():
        #    continue
        bgcid = os.path.basename(gbkfile)[:-4]
        if bgcid == 'AdOg':
            print("ERROR: This BGC shouldn't be here")
            exit()
        #print("\nWorking with %s" %(bgcid))
        (nrps_data_table, locustag_to_modulefeatures_dict,
         backbone_list, fnaseq, unexpected, domtag_to_feature_dict,
         backbone_stach_list) = get_nrps_data(gbkfile, manual_adom_annotation_table,
                                              manual_adom_annotation_file.replace(".tsv", '.faa'),
                                              stachelhaus_alignments_dir)
        if unexpected:
            print("Warning: Unexpected number of modules (%s) detected in BGC %s"
                  %(unexpected,bgcid))
            unexpected_list.append([bgcid, unexpected])
        bgc_to_fnaseq_dict[bgcid] = fnaseq
        bgcid_to_nrspsdatatable_dict[bgcid] = nrps_data_table
        bgcid_to_locustagmodules_dict[bgcid] = locustag_to_modulefeatures_dict
        #pprint(nrps_data_table)
        #pprint(locustag_to_modulefeatures_dict)
        total_backbone_res_list = list(set(total_backbone_res_list + backbone_list))
        tranlsated_backbones_list.append(transform_res_list(backbone_list))
        concat_stach = "".join(backbone_stach_list)
        backbones_table.append([bgcid, transform_res_list(backbone_list)])
        backbones_stach_table.append([bgcid, concat_stach])
        bgcid_to_domsdict_dict[bgcid] = domtag_to_feature_dict
        #break
    #pprint(bgcid_to_domsdict_dict)
    ### At this point the bgcid_to_nrspsdatatable_dict has for each BGC an ordered (according to X domain position)
    ### table with NRPS genes (columns: gene_num, locus_tag, startpos, endpos, strand).
    #pprint(bgcid_to_nrspsdatatable_dict)
    ### At this point the bgcid_to_locustagmodules_dict has for each BGC a dictionary where the NRPS genes locustags
    ### are keys and a correctly ordered list of module SeqFeatures are values
    #pprint(bgcid_to_locustagmodules_dict)
    #print(total_backbone_res_list)
    #pprint(tranlsated_backbones_list)


    ### saving info for checking in the end
    warnings_table.append(['BGC_ID','unexpected_no_of_modules'])
    warnings_table = warnings_table + unexpected_list

    ### do backbone analysis
    print("\nConducting backbone analysis based on Stachelhaus codes...")
    #backbones_aligned_fasta = run_backbone_analysis(backbone_dir, backbones_table)
    backbones_aligned_fasta = run_backbone_analysis(backbone_dir, backbones_stach_table, backbones_table)


    ### define starting point for the first module of each cluster according to the backbone alignment
    bgcid_to_1stmodulepos_dict = SeqIO.to_dict(SeqIO.parse(backbones_aligned_fasta, "fasta"))
    bgcid_to_1stmodulepos_dict = {k: str(v.seq) for (k, v) in bgcid_to_1stmodulepos_dict.items()}
    print("Considering all C-domain types as one category except Condensation_Starter. That includes Cglyc, "
          "Condensation_DCL, Condensation_LCL")
    for bgcid, msastring in bgcid_to_1stmodulepos_dict.items():
        if type(msastring) is int:
            continue
        #print(msastring)
        startindex = re.search(r'[^-]', msastring).start()
        #print(startindex)
        bgcid_to_1stmodulepos_dict[bgcid] = startindex
        ### declare balhimycin as the global standard
        if bgcid == 'bal':
            globalone = startindex

    print("\nGlobal position 1 of modules is in MSA position %s (before correction), based on balhimycin.\n" %(globalone))
    bgcid_to_1stmodulepos_dict = {k: v-globalone+1 for (k, v) in bgcid_to_1stmodulepos_dict.items()}
    #pprint(bgcid_to_1stmodulepos_dict)

    bgcid_to_moduleinfotable_dict = OrderedDict()
    bgcid_to_domainsinfodicts_dict = OrderedDict()
    dtype_to_bgcid_to_domainsinfo_dict = OrderedDict()
    moduleinfo_table_cols = ('nrps_locus_tag,module_index,startpos,endpos,strand,'
                             'a_dom_specificity,domains_list,complete').split(',')
    bgcid_to_domainsinfo_cols = ('nrps_locus_tag,module_index,type,label,a_dom_specificity,'
                                 'stach_aa10,stach_aa34,aa10_score,aa34_score,'
                                 'most_likely_substrate,beststachscore_substrate,beststach_score,beststach_evalue,'
                                 'module_active,startpos,endpos,strand,domainlabel,comment').split(',')
    ### iterate over all modules to set them in order
    print("\nSorting modules and domains...")
    for bgcid, locustag_to_modulefeatures_dict in bgcid_to_locustagmodules_dict.items():
        ### adding special check for known incomplete mibig clusters - removed avo on 03.12
        #if bgcid in ['avo','chl']:
        if bgcid in ['chl']:
            print("\nBGC %s not complete. Skipping..." %(bgcid))
            continue
        #print("\n", bgcid)
        nrps_data_table = bgcid_to_nrspsdatatable_dict[bgcid]
        locustags = [x[1] for x in nrps_data_table]
        module_index= bgcid_to_1stmodulepos_dict[bgcid]
        thisbgc_moduleinfo_table = []
        thisbgc_domaintypes_info_dict = OrderedDict()
        domtag_to_feature_dict = bgcid_to_domsdict_dict[bgcid]
        #pprint(domtag_to_feature_dict)
        for locustag in locustags:
            modulefeatureslist = locustag_to_modulefeatures_dict[locustag]
            #pprint(modulefeatureslist)
            for modulefeature in modulefeatureslist:
                #print(modulefeature)
                if 'complete' in modulefeature.qualifiers.keys():
                    completemodule='yes'
                else:
                    completemodule='no'
                thislocustag = modulefeature.qualifiers['locus_tags'][0]
                startpos = int(modulefeature.location.start)
                endpos = int(modulefeature.location.end)
                strand = int(modulefeature.location.strand)
                if "stach_match" in modulefeature.qualifiers:
                    monpairing = modulefeature.qualifiers['stach_match'][0]
                elif "manual_monomer_pairings" in modulefeature.qualifiers:
                    monpairing = modulefeature.qualifiers['manual_monomer_pairings'][0].split(" ")[0]
                elif "monomer_pairings" in modulefeature.qualifiers:
                    monpairing = modulefeature.qualifiers['monomer_pairings'][0].split(" ")[0]
                else:
                    monpairing = 'X'
                domains_list = modulefeature.qualifiers['domains']
                #print(module_index, domains_list)
                ### here extract info for domain labels and their types and index!!!!!!!!!!!!!!!!!!!!!!!!!!!!
                for domainid in domains_list:
                    ### covering case where a space appeared in the domain id
                    domainid = domainid.replace(' ','')
                    domainlabel = domainid.replace("nrpspksdomains_","")
                    thetype = domainlabel.replace(thislocustag, "")[1:-2]
                    ### added on 26.01.24 - keeping condensation starters separate.
                    if thetype == 'Condensation_Starte r':
                        thetype = "Condensation_Starter"
                    if thetype in ('Cglyc,Condensation_DCL,Condensation_LCL').split(','):
                        thetype = 'Condensation'
                    if thetype == 'AMP-binding':
                        spec = monpairing
                        try:
                            extra_info= domtag_to_feature_dict[domainlabel]
                            ### making sure lines are equal
                            if len(extra_info)<9:
                                extra_info.append('')
                        except:
                            #print('failed to get stach info for %s' %(domainlabel))
                            #print(domtag_to_feature_dict.keys())
                            extra_info=['','','','','','','','','']
                        #pprint(extra_info)
                    else:
                        spec = ''
                        extra_info = ['','','','','','','','','']
                    domaininfo = [thislocustag, module_index, thetype, domainlabel, spec] + extra_info
                    #if thislocustag == 'StAfJ3_2_00100':
                    #    print()
                    if thetype in thisbgc_domaintypes_info_dict.keys():
                        thetable = thisbgc_domaintypes_info_dict[thetype]
                        thetable.append(domaininfo)
                        thisbgc_domaintypes_info_dict[thetype] = thetable
                    else:
                        thisbgc_domaintypes_info_dict[thetype] = [domaininfo]
                    if thetype in dtype_to_bgcid_to_domainsinfo_dict.keys():
                        if bgcid in dtype_to_bgcid_to_domainsinfo_dict[thetype].keys():
                            theetable = dtype_to_bgcid_to_domainsinfo_dict[thetype][bgcid]
                            theetable.append(domaininfo)
                            dtype_to_bgcid_to_domainsinfo_dict[thetype][bgcid] = theetable
                        else:
                            dtype_to_bgcid_to_domainsinfo_dict[thetype][bgcid] = [domaininfo]
                    else:
                        dtype_to_bgcid_to_domainsinfo_dict[thetype] = {bgcid:[domaininfo]}

                ### continue saving info for modules
                domainslist_string = '|'.join(domains_list)
                moduleline = [thislocustag,module_index,startpos,endpos,strand, monpairing,domainslist_string, completemodule]
                thisbgc_moduleinfo_table.append(moduleline)
                module_index+=1
                #break
            #break
        bgcid_to_domainsinfodicts_dict[bgcid] = thisbgc_domaintypes_info_dict
        bgcid_to_moduleinfotable_dict[bgcid] = thisbgc_moduleinfo_table
        #break
    #pprint(bgcid_to_domainsinfodicts_dict)
    #pprint(bgcid_to_moduleinfotable_dict)

    ### save module info to csv - and seqs to fastas?
    modulesouttable = os.path.join(modules_dir, 'modules_metadata.csv')
    save_moduleinfo_to_csv(bgcid_to_moduleinfotable_dict,moduleinfo_table_cols, modulesouttable)
    domain_label_to_modulegenecount_dict = extract_module_seqs(modules_dir, bgcid_to_moduleinfotable_dict, bgc_to_fnaseq_dict)
    ### save also NRPS genes info in the same way
    outnrpstable = os.path.join(output, 'nrpsgenes_metadata.csv')
    save_moduleinfo_to_csv(bgcid_to_nrspsdatatable_dict,
                           ['gene_num_in_cluster', 'locus_tag', 'start_pos', 'end_pos', 'strand'],
                           outnrpstable)
    ### save domain info to csv - and seqs to fastas?
    bgcid_to_domainsinfodicts_dict = extract_domain_seqs(domains_dir, dtype_to_bgcid_to_domainsinfo_dict,
                                                         bgc_to_fnaseq_dict,bgc_gbks_list,
                                                         domain_label_to_modulegenecount_dict,
                        bgcid_to_domainsinfodicts_dict)
    domainsouttable = os.path.join(domains_dir, 'domains_metadata.csv')
    save_domaininfo_to_csv(bgcid_to_domainsinfodicts_dict, bgcid_to_domainsinfo_cols, domainsouttable)
    ### added on 29.04.24 to make it easier to work with in iTOL
    make_itol_formatted_table(domainsouttable)
    ### save warnings too
    save_table_to_csv(warnings_table, warningsfile)
    print("\nDone")

#'''
# Parsing arguments & Execution #
parser = argparse.ArgumentParser(description='Given a folder of antiSMASH-generated gbk files of glycopeptide '
                                             'BGCs, detect NRPS genes, align predicted AA backbones to determine'
                                             ' the central residue, then extract modules and domains naming them'
                                             ' based on the central residue.')
parser.add_argument('-i', '--input', help='Input folder of antiSMASH-generated gbk files', required=True)
parser.add_argument('-o', '--output', help='Output folder where all files will be created', required=True)
parser.add_argument('-a', '--annotationfile', help='Table of A domain specificiites', required=False,
                    default=False)
parser.add_argument('-r', '--replace', help='Replace existing files', required=False,
                    default=False)

if __name__ == "__main__":
    args = parser.parse_args()
    main(args.input, args.output, args.annotationfile, args.replace)
#'''

#### just make the iTOL table
#make_itol_formatted_table('/media/hdd_2tb/GPA_reclassification_clean/A_dataset_creation/4_orthology_inference/1_trimmed_complete_domain_analysis/2_domains_analysis/domains_metadata.csv')

### just make the fixed weblogo
#execute_weblogo('/media/hdd_2tb/GPA_reclassification_clean/A_dataset_creation/4_orthology_inference/1_trimmed_complete_domain_analysis/0_backbone_analysis/backbones_stach34concat_mafftMSA_simplified_fixed.faa',
#                '/media/hdd_2tb/GPA_reclassification_clean/A_dataset_creation/4_orthology_inference/1_trimmed_complete_domain_analysis/0_backbone_analysis/fixed_logo.png')

'''
### for debugging - take care
input='/media/hdd_2tb/GPA_reclassification_clean/A_dataset_creation/3_GPA_candidate_dereplication/3a_trimmed_complete_GPAclusters'
output='/media/hdd_2tb/GPA_reclassification_clean/A_dataset_creation/4_orthology_inference/1_trimmed_complete_domain_analysis'
#annotationfile = '/media/hdd_2tb/GPA_reclassification_clean/A_dataset_creation/3_GPA_candidate_dereplication/backbone_exceptions_manual.tsv'
annotationfile = '/media/hdd_2tb/GPA_reclassification_clean/A_dataset_creation/3_GPA_candidate_dereplication/adomains_known_specs.tsv'
main(input, output, annotationfile)
'''
```
