## Supplementary Data 5 for "Structural and evolutionary analyses support reclassification of glycopeptide antibiotics as xyclopeptides": extract_trimmed_limits.py

```
#!/usr/bin/python3.6
# Created by Athina Gavriilidou at 02.08.23 18:20 #
# Imports #
import argparse
import os
from pprint import pprint
import pandas as pd
import json
import math
from Bio import SeqIO
import csv

# Functions #
def create_folder(path):
    if not os.path.exists(path):
        os.mkdir(path)

'''given a directory, lists all files in it with the given suffix'''
def list_files(directory, suffix):
    files = [os.path.join(directory, o) for o in os.listdir(directory)
               if os.path.isfile(os.path.join(directory, o)) and o.endswith(suffix)]
    print("Found %s %s files" % (len(files), suffix))
    files.sort()
    return files

def get_limits_from_json(clinkerjson, bgc_to_limits_dict):
    with open(clinkerjson, 'r') as jsonfile:
        jsoncontent = json.load(jsonfile)
        data = jsoncontent['data']
        clusters = data['clusters']
        ### go into the clusters
        for c in clusters:
            bgcid = c['name']
            #bgcclinkerid = c['uid']
            bgclocus = c['loci'][0]
            lefttrim = bgclocus['_trimLeft']
            righttrim = bgclocus['_trimRight']
            genes_list = bgclocus['genes']
            if not lefttrim is None:
                left_locustag = lefttrim['names']['locus_tag']
                left_location = lefttrim['start']
            else:
                left_locustag = genes_list[0]['names']['locus_tag']
                left_location = genes_list[0]['start']
            if not righttrim is None:
                right_locustag = righttrim['names']['locus_tag']
                right_location = righttrim['end']
            else:
                right_locustag = genes_list[-1]['names']['locus_tag']
                right_location = genes_list[-1]['end']
            ### making sure even reversed clusters are in order in the table
            if left_location > right_location:
                templeftloc = left_location
                left_location = right_location
                right_location = templeftloc
                #bgc_to_limits_dict[bgcid] = [left_locustag, right_locustag, left_location, right_location]
            leftlocusid = left_locustag.split("_")[-1]
            rightlocusid = right_locustag.split("_")[-1]
            if int(leftlocusid) > int(rightlocusid):
                templeftlocutag = left_locustag
                left_locustag = right_locustag
                right_locustag = templeftlocutag
            ### now save info - 22.10.2023 - json limits have proven unreliable, will switch to using locus_tags
            bgc_to_limits_dict[bgcid] = [left_locustag, right_locustag, left_location, right_location]
            #if bgcid == "AmLr_2":
            #    print()
    return bgc_to_limits_dict

def list_to_string(thelist, connector='_'):
    thestring = ''
    for e in thelist:
        thestring = thestring + str(e) + connector
    thestring = thestring[:-len(connector)]
    return thestring

def extract_gbkandfaaandfna(untrimmedgbk, left_locustag, right_locustag, outgbk,outfaa, outfna, bgcid,
                            thisbgc_domainto_stachelhaus_dict):
    input_handle = open(untrimmedgbk, 'r')
    records = SeqIO.parse(input_handle, "genbank")
    output_handlefaa = open(outfaa, "w")
    output_handlefna = open(outfna, "w")
    tokeep = False
    toremovefeatures = []
    #bgcid, label, feature.location, original_locus_tag, bgclocus,feature.location._start.position,feature.location._end.position
    genes_meta_table = ['bgcid,locus_tag,location,original_locus_tag,bgc_locus,infile_startpos,infile_endpos,'
                        'name,gene,gene_kind,gene_functions,product,db_xref,note'.split(',')]
    reclist = []

    foundleftlim=False
    ### first find start and end locations or take everything if right and left are false
    for seq_record in records:
        for feature in seq_record.features:
            if feature.type == "CDS":
                if not foundleftlim and left_locustag is False or feature.qualifiers['locus_tag'][0] == left_locustag:
                    leftlim = feature.location._start.position
                    foundleftlim = True
                if right_locustag is not False and feature.qualifiers['locus_tag'][0] == right_locustag: # no elif because of very fragmented clusters
                    rightlim = feature.location._end.position
                    break
        if right_locustag is False:
            rightlim = feature.location._end.position
    # for testing
    #if left_locustag is False and right_locustag is False:
    #    print([leftlim, rightlim])
    #    print(seq_record)
    try:
        bothlims = [leftlim, rightlim]
    except:
        print(left_locustag, right_locustag)
        print("ERROR: could not find limits for %s. Aborting..." %(untrimmedgbk))
        exit()

    #if "AmBlD4" in untrimmedgbk:
    #    print()

    ### for Stachelhaus code qualifiers
    aspec_cols = 'stach_aa10	stach_aa34	aa10_score	aa34_score	most_likely_substrate'.split("\t")

    ### then the rest
    input_handle = open(untrimmedgbk, 'r')
    records = SeqIO.parse(input_handle, "genbank")
    for seq_record in records:
        featlist=[]
        bgclocus = seq_record.id
        for feature in seq_record.features:
            try:
                ### exception
                #if (bgcid == 'Stsf0' and 'locus_tag' in feature.qualifiers.keys()
                #        and feature.qualifiers['locus_tag'][0] == 'StsF0__61'):
                #    print("Special case of CDS StsF0__61: hardcoded position complement(97664..98916)")
                #    feature.location._start.position = 97664
                #    feature.location._end.position = 98916
                #    feature.location.strand = -1
                if feature.type in ['proto_core','cand_cluster', 'protocluster']:
                    featlist.append(feature) ### changed on 20.10.2023
                    continue
                ### add stachelhaus code info
                if feature.type == 'aSDomain' and 'AMP-binding' == feature.qualifiers['aSDomain'][0]:
                    thistag = feature.qualifiers['domain_id'][0]
                    try:
                        stach_info = thisbgc_domainto_stachelhaus_dict[thistag]
                    except:
                        stach_info = []
                    if len(stach_info) > 0:
                        for i in range(len(aspec_cols)):
                            label = aspec_cols[i]
                            feature.qualifiers[label] = [stach_info[i]]
                if not tokeep:
                    try:
                        start_pos = feature.location._start.position
                    except:
                        start_pos = min(feature.location)
                    if start_pos >= leftlim:
                        tokeep = True
                if tokeep:
                    try:
                        end_pos = feature.location._end.position
                    except:
                        end_pos = max(feature.location)
                    if end_pos > rightlim:
                        tokeep = False
                if not tokeep:
                    toremovefeatures.append(feature)
                else:
                    featlist.append(feature)
                ### for getting the protein sequences
                if tokeep and feature.type == "CDS":
                    try:    ### added on 29.11.21
                        if len(feature.qualifiers['translation']) != 1:
                            print("Something wrong with feature %s. \nSkipping" %(feature))
                            continue
                    except: ### added on 29.11.21
                        print("Could not even find tranlsation in feature %s.\n Attempting to retrieve it from DNA seq"
                              %(feature.qualifiers['locus_tag'][0]))
                        gene_sequence = feature.extract(seq_record.seq)
                        feature.qualifiers['translation'] = [gene_sequence.translate()]
                    ### collecting metadata
                    label = feature.qualifiers['locus_tag'][0]
                    try:
                        original_locus_tag = feature.qualifiers['original_locus_tag'][0]
                    except:
                        original_locus_tag = ""
                    try:
                        gene_meta_list = [bgcid, label, feature.location, original_locus_tag, bgclocus,
                                          feature.location._start.position,feature.location._end.position]
                    ### exception necessary for split gene locations e.g. location: join{[98633:98916](-), [97663:98634](-)}
                    except:
                        gene_meta_list = [bgcid, label, feature.location, original_locus_tag, bgclocus,
                                          min(feature.location), max(feature.location)]

                    for qual in 'Name,gene,gene_kind,gene_functions,product,db_xref,note'.split(','):
                        try:
                            if qual == 'gene_functions' or qual == 'product' or qual == 'db_xref':
                                thisqual = feature.qualifiers[qual]
                                thisqual = list_to_string(thisqual, connector='|')
                            else:
                                thisqual = feature.qualifiers[qual][0]
                        except:
                            thisqual = ''
                        gene_meta_list.append(thisqual)
                    gene_meta_list.append(thisqual)
                    #print(gene_meta_list)
                    #gene_meta_list.append(original_locus_tag)
                    #gene_meta_list.append(bgclocus)
                    #gene_meta_list.append(feature.location._start.position)
                    #gene_meta_list.append(feature.location._end.position)
                    genes_meta_table.append(gene_meta_list)
                    #print(genes_meta_table)
                    output_handlefaa.write(">%s |%s\n%s\n" % (
                        # seq_feature.qualifiers['locus_tag'][0],
                        label,
                        feature.location,
                        # bgc_filename,
                        # seq_record.name,
                        feature.qualifiers['translation'][0]))
            except:
                print("ERROR: There was some kind of issue with BGC %s!\n" %(bgcid))
                print(feature)
                #print(min(feature.location), max(feature.location))
                exit()

        #for feature in toremovefeatures:
        #    seq_record.features.remove(feature)
        seq_record.features = featlist
        reclist.append(seq_record)
    #print(seq_record.features[0])
    ### create a fixed copy
    with open(outgbk, 'w') as output_handle:
        SeqIO.write(seq_record, output_handle, 'genbank')
    ### save also dna sequence
    dna_records = []
    #print(outgbk)
    #print(seq_record)
    #print(type(leftlim), type(rightlim))
    dnaseq = seq_record.seq[leftlim:rightlim]
    output_handlefna.write(">%s |[%s:%s]\n%s\n" % (
                    bgcid,
                    leftlim,
                    rightlim,
                    dnaseq))
    return genes_meta_table, leftlim, rightlim

'''save lines in csv format in the given output file'''
def save_lines_csv(lines, output):
    with open(output, 'w') as f:
        tsv_output = csv.writer(f)
        for line in lines:
            tsv_output.writerow(line)
    print("\nCreated file: " + output)

def extract_trimmed_clusters(clustermetadata_df, untrimmed_gbk_files_list, outputcomplete, outputincomplete,
                             bgc_to_limits_dict, bgcid_to_domainspecinfo_dict):
    genes_meta_table_full = ['bgcid,locus_tag,location,original_locus_tag,bgc_locus,infile_startpos,infile_endpos,'
                             'locus_startpos,locus_endpos,name,gene,gene_kind,gene_functions,product,db_xref,note'.split(',')]
                             #'original_locus_tag,bgc_locus,infile_startpos,infile_endpos,locus_startpos,locus_endpos'.split(',')]
    for untrimmedgbk in untrimmed_gbk_files_list:
        bgcid = os.path.basename(untrimmedgbk)[:-4]
        if bgcid not in clustermetadata_df.index.tolist():
            continue
        iscomplete = clustermetadata_df.loc[bgcid, 'considered_complete']
        ### 15.01.24 getting more info for reproducibility
        #as7_BGC_locus = clustermetadata_df.loc[bgcid, 'as7_BGC_locus']
        try:
            as7_clusterstart = int(clustermetadata_df.loc[bgcid, 'as7_clusterstart'])
        except:
            if clustermetadata_df.loc[bgcid, 'as7_clusterstart'].startswith('<'):
                as7_clusterstart = int(clustermetadata_df.loc[bgcid, 'as7_clusterstart'][1:])
                print(f"Uncommon starting location for {bgcid}: {clustermetadata_df.loc[bgcid, 'as7_clusterstart']}"
                      f"\nAdjusting it to {as7_clusterstart}")

        ### go through with it only if the right and left limits are specified
        try:
            #leftlim = int(trimmed_metadata_df.loc[bgcid, 'trimmed_clusterstart'])
            #[left_locustag, right_locustag, left_location, right_location]
            left_locustag = bgc_to_limits_dict[bgcid][0]
            #rightlim = int(trimmed_metadata_df.loc[bgcid, 'trimmed_clusterend'])
            right_locustag = bgc_to_limits_dict[bgcid][1]
        except:
            print("No limits found for %s. Skipping..." %(bgcid))
            continue
        if iscomplete == 'yes':
            outgbk = os.path.join(outputcomplete, '%s.gbk' %(bgcid))
        else:
            outgbk = os.path.join(outputincomplete, '%s.gbk' % (bgcid))
        outfaa = outgbk[:-4] + '.faa'
        outfna = outgbk[:-4] + '.fna'
        #if bgcid == 'AmLr_2':
        #    print()
        thisbgc_domainto_stachelhaus_dict = bgcid_to_domainspecinfo_dict[bgcid]
        genes_meta_table, left_location, right_location = extract_gbkandfaaandfna(untrimmedgbk, left_locustag,
                                                                                  right_locustag, outgbk,outfaa,
                                                                                  outfna, bgcid,
                                                                                  thisbgc_domainto_stachelhaus_dict)
        ### 15.01.24 calculate location in original locus
        #if as7_clusterstart != 0:
        locusstart_location = left_location + as7_clusterstart
        locusend_location = right_location + as7_clusterstart
        #print(genes_meta_table)
        #print(genes_meta_table[0])
        for i in range(1,len(genes_meta_table)):
            linelist = genes_meta_table[i]
            #pprint([genes_meta_table[0],linelist])
            locusgenestart = int(linelist[5]) + as7_clusterstart
            #linelist.append(locusgenestart)
            locusgeneend = int(linelist[6]) + as7_clusterstart
            #linelist.append(locusgeneend)
            linelist = linelist[:7] + [locusgenestart,locusgeneend] + linelist[7:]
            genes_meta_table[i] = linelist

        bgc_to_limits_dict[bgcid] = [left_locustag, right_locustag, left_location, right_location,
                                     locusstart_location, locusend_location]
        genes_meta_table_full = genes_meta_table_full + genes_meta_table[1:]
        #break
    return genes_meta_table_full, bgc_to_limits_dict

'''for copying files. Make everything=True and then everything from the source will be copied to the destination'''
def execute_cp(source_fileordir, destinationfileordir, everything=False):
    if everything:
        cp_command = 'cp %s/* %s/.' %(source_fileordir,destinationfileordir)
    else:
        cp_command = 'cp \'%s\' %s' %(source_fileordir,destinationfileordir)
    try:
        os.system(cp_command)
    except:
        print("Warning. There was a problem with copying %s to %s" %(source_fileordir, destinationfileordir))

'''added on 24.11.2023 - we need to double check Stachelhaus codes later'''
def getextraStachelhausinfo(antismashinput):
    json_files = list_files(antismashinput, '.json')
    bgcid_to_domainspecinfo_dict = {}
    for jsonfile in json_files:
        bgcid = os.path.basename(jsonfile)[:-5]
        domain_tag_to_stachelhausinfo_dict = {}
        #print(jsonfile)
        with open(jsonfile,'r') as injson:
            jsoncontent = json.load(injson)
            record = jsoncontent['records'][0]
            #print(record.keys())
            modules_dict = record['modules']
            #print(modules_dict.keys())
            nrps_pks_domains_predictions = modules_dict['antismash.modules.nrps_pks']['domain_predictions']
            #print(len(record['features'])) #445
            domain_locus_tags_list = nrps_pks_domains_predictions.keys()
            for dom_tag in domain_locus_tags_list:
                if 'AMP-binding' in dom_tag:
                    try:
                        thisdomain_info = nrps_pks_domains_predictions[dom_tag]['nrpys']
                    ### if the whole thing is empty, it is probably a broken BGC so skip
                    except:
                        continue
                        #print(dom_tag)
                        #print(nrps_pks_domains_predictions[dom_tag])
                        #exit()
                    thisdomain_aa10 = thisdomain_info['aa10']
                    thisdomain_aa34 = thisdomain_info['aa34']
                    try:
                        thisdomain_stachelhausinfo = thisdomain_info['stachelhaus_matches'][0]
                        thisdomain_aa10score = thisdomain_stachelhausinfo['aa10_score']
                        thisdomain_aa34score = thisdomain_stachelhausinfo['aa34_score']
                        ### there may be more than one but the first should be the highest score
                        thisdomain_substrate = thisdomain_stachelhausinfo['substrates'][0]['norine']
                    ### if matches is empty we have no idea what it is
                    except:
                        #print(dom_tag)
                        #print(thisdomain_info)
                        thisdomain_aa10score = ''
                        thisdomain_aa34score = ''
                        thisdomain_substrate = 'X'
                        #exit()
                    #print(thisdomain_substrate)
                    domain_tag_to_stachelhausinfo_dict[dom_tag] = [thisdomain_aa10, thisdomain_aa34, thisdomain_aa10score,
                                                                   thisdomain_aa34score, thisdomain_substrate]
            #print(nrps_pks_domains_predictions.keys())
            #print(nrps_pks_domains['domain_predictions'])

        #pprint(domain_tag_to_stachelhausinfo_dict)
        bgcid_to_domainspecinfo_dict[bgcid] = domain_tag_to_stachelhausinfo_dict
    #pprint(bgcid_to_domainspecinfo_dict)
    #exit()
    #print(bgcid_to_domainspecinfo_dict.keys())
    return bgcid_to_domainspecinfo_dict


# Main function #
def main(clinkerinput, antismashinput, metadata, outputcomplete, outputincomplete, metadataout, outputcompleteoriginal):
    print("\nInput clinker folder: %s\n"
          "Input antiSMASH generated gbk files: %s\n"
          "Input metadata table: %s\n"
          "Output folder for complete clusters: %s\n"
          "Output folder for incomplete clusters: %s\n"
          "Output updated metadata table: %s\n"
          %(clinkerinput, antismashinput, metadata, outputcomplete, outputincomplete, metadataout))
    if outputcompleteoriginal:
        print("Output folder for complete original (non-trimmed) clusters: %s\n" %(outputcompleteoriginal))
        create_folder(outputcompleteoriginal)
    clustermetadata_df = pd.read_csv(metadata)
    clustermetadata_df.set_index('ID', drop=False, inplace=True)
    untrimmed_gbk_files_list = list_files(antismashinput, '.gbk')
    bgcid_to_domainspecinfo_dict = getextraStachelhausinfo(antismashinput)

    #'''
    create_folder(outputcomplete)
    create_folder(outputincomplete)
    trimmed_json_files_list = list_files(clinkerinput, '_trimmed.json')
    ### may change it later, work only with entries that have been manually trimmed
    if False:#True:
        originalrows = clustermetadata_df.shape[0]
        clustermetadata_df = clustermetadata_df.dropna(subset=['considered_complete'])
        ### drop also all with no GCF assignment (the ones where antismash failed)
        clustermetadata_df = clustermetadata_df.dropna(subset=['GCF_T0.2'])
        clustermetadata_df['GCF_T0.2'] = clustermetadata_df['GCF_T0.2'].astype(int)
        print("\nKeeping only entries that have been manually curated based on the 'considered_complete' column."
              "\nProceeding with %s entries out of %s\n" %(clustermetadata_df.shape[0], originalrows))
        #print(clustermetadata_df)
    bgc_to_limits_dict = {}
    ### iterate over trimmed json files
    for clinkerjson in trimmed_json_files_list:
        gcfid = os.path.basename(clinkerjson).split('_')[1]
        bgc_to_limits_dict = get_limits_from_json(clinkerjson, bgc_to_limits_dict)


    print("\nExtracting complete trimmed clusters from files...")
    genes_meta_table_full, bgc_to_limits_dict = extract_trimmed_clusters(clustermetadata_df, untrimmed_gbk_files_list,
                                                                         outputcomplete,
                                                                         outputincomplete, bgc_to_limits_dict, bgcid_to_domainspecinfo_dict)
    save_lines_csv(genes_meta_table_full, os.path.join(os.path.dirname(metadataout), 'trimmed_clusters_gene_table.csv'))
    ### save trimmed summary after fixing start-end locations
    newcolsnames = ('trimmed_start_gene,trimmed_end_gene,trimmed_clusterstart,trimmed_clusterend,'
                    'locus_trimmed_clusterstart, locus_trimmed_clusterend').split(',')
    newcols_df = pd.DataFrame.from_dict(bgc_to_limits_dict, orient='index',
                                        columns=newcolsnames)
    trimmed_metadata_df = pd.concat([clustermetadata_df, newcols_df], axis=1)
    print(trimmed_metadata_df)
    trimmed_metadata_df.to_csv(metadataout, index=False)
    print("\nCreated file: %s\n" % (metadataout))
    #'''
    if outputcompleteoriginal:
        print("\nExtracting complete non trimmed clusters from files...")
        originalclstersdf = clustermetadata_df[(clustermetadata_df['considered_complete'] == 'yes')]
        print(originalclstersdf)
        #exit()
        for infile in untrimmed_gbk_files_list:
            thename = os.path.basename(infile)
            theid = thename[:-4]
            if theid in originalclstersdf.index.tolist():
            #if originalclstersdf['ID'].str == theid:
                newname = os.path.join(outputcompleteoriginal, thename)
                #print(theid, newname)
                #execute_cp(infile,newname)
                thisbgc_domainto_stachelhaus_dict = bgcid_to_domainspecinfo_dict[theid]
                extract_gbkandfaaandfna(infile,False, False, newname,'%s.faa' %(newname[:-4]),
                                        '%s.fna' %(newname[:-4]),theid,thisbgc_domainto_stachelhaus_dict)
    print("\nDone")

#'''
# Parsing arguments & Execution #
parser = argparse.ArgumentParser(description='Uses clinker trimmed.json files to infer trimmed cluster ')
parser.add_argument('-ic', '--clinkerinput', help='Input directory of clinker files, including '
                                                  'trimmed files (e.g. gcf_56_part1_risA_trimmed.json)',
                    required=True)
parser.add_argument('-ia', '--antismashinput', help='Input directory of gbk files of untrimmed clusters'
                                                    ' after antiSMASH annotation',
                    required=True)
parser.add_argument('-oc', '--outputcomplete', help='Output directory for complete trimmed dereplicated'
                                                    ' clusters', required=True)
parser.add_argument('-oi', '--outputincomplete', help='Output directory for non complete trimmed replicate'
                                                    ' clusters', required=True)
parser.add_argument('-oco', '--outputcompleteoriginal', help='Output directory for complete non trimmed '
                                                             ' clusters', required=False, default=False)
parser.add_argument('-m', '--metadata', help='Input metadata table of clusters', required=True)
parser.add_argument('-mo', '--metadataout', help='Output path of metadata table of clusters to be updated'
                                              ' with new info', required=True)
if __name__ == "__main__":
    args = parser.parse_args()
    main(args.clinkerinput, args.antismashinput, args.metadata, args.outputcomplete, args.outputincomplete,
         args.metadataout, args.outputcompleteoriginal)
#'''

'''
### for debugging
clinkerinput = '/media/hdd_2tb/GPA_reclassification_clean/A_dataset_creation/3_GPA_candidate_dereplication/2_clinker_gcf_results'
antismashinput = '/media/hdd_2tb/GPA_reclassification_clean/A_dataset_creation/2_putative_GPAs_analyses/5_all_candidate_as7_clusters'
outputcomplete = '/media/hdd_2tb/GPA_reclassification_clean/A_dataset_creation/3_GPA_candidate_dereplication/3a_trimmed_complete_GPAclusters'
outputincomplete = '/media/hdd_2tb/GPA_reclassification_clean/A_dataset_creation/3_GPA_candidate_dereplication/3b_trimmed_incomplete_GPAclusters'
outputcompleteoriginal = '/media/hdd_2tb/GPA_reclassification_clean/A_dataset_creation/3_GPA_candidate_dereplication/3c_untrimmed_complete_GPAclusters'
metadata = '/media/hdd_2tb/GPA_reclassification_clean/A_dataset_creation/3_GPA_candidate_dereplication/2_clinker_gcf_results/summary_hits_clusters_metadata_trimmed.csv'
### using old metadata for testnig
metadata = '/media/hdd_2tb/GPA_reclassification_clean/A_dataset_creation/3_GPA_candidate_dereplication/2_clinker_gcf_results_231023/summary_hits_clusters_metadata_trimmed.csv'
metadataout = '/media/hdd_2tb/GPA_reclassification_clean/A_dataset_creation/3_GPA_candidate_dereplication/summary_clusters_trimmed_metadata.csv'
main(clinkerinput, antismashinput, metadata, outputcomplete, outputincomplete, metadataout, outputcompleteoriginal)
'''
```
