## Supplementary Data 5 for "Structural and evolutionary analyses support reclassification of glycopeptide antibiotics as xyclopeptides": gbk_to_faa.py

```
#!/usr/bin/python3.6
# Created by Athina Gavriilidou at 27.02.20 10:18 #
### code segments taken from: https://warwick.ac.uk/fac/sci/moac/people/students/peter_cock/python/genbank2fasta/#gbk2faa
# Imports #
import argparse
from Bio import SeqIO
import os
from pprint import pprint
import csv
from datetime import datetime
from collections import OrderedDict
import random
import copy
import pandas as pd

# Functions #
'''save lines in csv format in the given output file'''
def save_lines_csv(lines, output):
    with open(output, 'w') as f:
        tsv_output = csv.writer(f)
        for line in lines:
            tsv_output.writerow(line)
    print("\nCreated file: " + output)

def list_to_string(thelist, connector='_'):
    thestring = ''
    for e in thelist:
        thestring = thestring + str(e) + connector
    thestring = thestring[:-len(connector)]
    return thestring

'''Takes a gbk file and creates an faafile with all AA seqs indexed by the locus tag. taken from source above'''
def gbkfile_to_faa(gbkfile, outfaafile, gpa=False, overwrite=False):
    input_handle = open(gbkfile, "r")
    if not os.path.exists(outfaafile) or overwrite:
        output_handle = open(outfaafile, "w")
        opened_output=True
    else:
        opened_output = False
    bgc_filename = gbkfile.split("/")[-1]
    bgc_filename = bgc_filename[:-4]
    gene_counter = 1
    cd_counter = 0
    #seq_counter = 0
    genes_meta_table = ['label,locus_tag,location,name,gene,gene_kind,gene_functions,product'.split(',')]
    newrecords=[]
    for seq_record in SeqIO.parse(input_handle, "genbank"):
        print("Dealing with GenBank record %s" % seq_record.id)
        for seq_feature in seq_record.features:
            if seq_feature.type == "CDS":
                cd_counter+=1
                #print(seq_feature)
                #assert len(seq_feature.qualifiers['translation']) == 1 ###commented out on 29.11.21
                try:    ### added on 29.11.21
                    if len(seq_feature.qualifiers['translation']) != 1:     ### added on 29.11.21
                        print("Something wrong with feature %s. \nSkipping" %(seq_feature))
                        continue
                except: ### added on 29.11.21
                    print("Could not even find tranlsation in feature %s.\n Attempting to retrieve it from DNA seq"
                          %(seq_feature.qualifiers['locus_tag'][0]))
                    gene_sequence = seq_feature.extract(seq_record.seq)
                    seq_feature.qualifiers['translation'] = [gene_sequence.translate()]
                    #exit()
                ### collecting metadata
                if gpa:
                    label = "%s_GPA_%s" %(bgc_filename, gene_counter)
                else:
                    try:
                        label = "%s|%s" %(bgc_filename, seq_feature.qualifiers['locus_tag'][0])
                    except:
                        try:
                            print('No locus_tag found for CDS at %s. Using gene name' %(seq_feature.location))
                            label = "%s|%s" %(bgc_filename, seq_feature.qualifiers['gene'][0])
                        except:
                            print('No locus_tag or gene name found for CDS at %s. Using protein_id' % (seq_feature.location))
                            label = "%s|%s" % (bgc_filename, seq_feature.qualifiers['protein_id'][0])
                try:
                    gene_meta_list = [label, seq_feature.qualifiers['locus_tag'][0], seq_feature.location]
                    ### special case with very short translation
                    if len(seq_feature.qualifiers['translation'][0]) <10:# is seq_feature.qualifiers['locus_tag'][0]:
                        print("Warning: skipping short sequence at %s" %(seq_feature.location))
                        cd_counter-=1
                        continue
                except:
                    try:
                        gene_meta_list = [label, seq_feature.qualifiers['gene'][0], seq_feature.location]
                    except:
                        gene_meta_list = [label, seq_feature.qualifiers['protein_id'][0], seq_feature.location]
                for qual in 'Name,gene,gene_kind,gene_functions,product'.split(','):
                    try:
                        if qual == 'gene_functions' or qual == 'product':
                            thisqual = seq_feature.qualifiers[qual]
                            thisqual = list_to_string(thisqual, connector='|')
                        else:
                            thisqual = seq_feature.qualifiers[qual][0]
                    except:
                        thisqual = ''
                    gene_meta_list.append(thisqual)
                genes_meta_table.append(gene_meta_list)
                #pprint(genes_meta_table)
                if opened_output:
                    output_handle.write(">%s|%s\n%s\n" % (
                        #seq_feature.qualifiers['locus_tag'][0],
                        label,
                        seq_feature.location,
                        #bgc_filename,
                        # seq_record.name,
                        seq_feature.qualifiers['translation'][0]))
                #seq_counter+=1
                ### fix annotations in gbk file
                try:
                    seq_feature.qualifiers['locus_tag_original'] = [seq_feature.qualifiers['locus_tag'][0]]
                    seq_feature.qualifiers['locus_tag'] = [label]
                except:
                    try:
                        seq_feature.qualifiers['locus_tag_original'] = [seq_feature.qualifiers['gene'][0]]
                        seq_feature.qualifiers['locus_tag'] = [label]
                    except:
                        seq_feature.qualifiers['locus_tag_original'] = [seq_feature.qualifiers['protein_id'][0]]
                        seq_feature.qualifiers['locus_tag'] = [label]
                gene_counter+=1
        newrecords.append(seq_record)
    if opened_output:
        output_handle.close()
    input_handle.close()
    ### added check to see if saved sequences are as many as they should be - can add exceptions as found
    seq_counter = len([1 for line in open(outfaafile, 'r') if line.startswith(">")])
    if cd_counter != seq_counter:
        print("\nERROR: Number of CDS in gbk file does not match number of sequences in fasta file (%s != %s)\n"
              "File: %s\n" %(cd_counter, seq_counter, gbkfile))
        exit()

'''given a directory, lists all gbk files'''
def list_gbk_files(directory):
    gbk_files = []
    for file in os.listdir(directory):
        if file.endswith(".gbk"):
            gbk_files.append(os.path.join(directory, file))
    print("Found " + str(len(gbk_files)) + " .gbk files")
    return gbk_files

'''creates folder in given path, if it does not exist already'''
def create_folder(path):
    if not os.path.exists(path):
        os.mkdir(path)

'''taken from https://warwick.ac.uk/fac/sci/moac/people/students/peter_cock/python/genbank2fasta/'''
def gbk_to_fna(gbk_filename, fna_filename):
    input_handle = open(gbk_filename, "r")
    output_handle = open(fna_filename, "w")

    for seq_record in SeqIO.parse(input_handle, "genbank"):
        print("Dealing with GenBank record %s" % seq_record.id)
        output_handle.write(">%s %s\n%s\n" % (
            seq_record.id,
            seq_record.description,
            seq_record.seq))

    output_handle.close()
    input_handle.close()

def extract_locus_from_gbk(locusname, gbk_filename):
    print("Extracting %s from %s" %(locusname, gbk_filename))
    outfilename = os.path.join(os.path.dirname(gbk_filename), os.path.basename(gbk_filename).replace("full", locusname))
    input_handle = open(gbk_filename, "r")
    newrecords = []
    for seq_record in SeqIO.parse(input_handle, "genbank"):
        #print("Dealing with GenBank record %s" % seq_record.id)
        if seq_record.id == locusname:
            newrecords.append(seq_record)
    input_handle.close()
    ### create a fixed copy
    with open(outfilename, 'w') as output_handle:
        SeqIO.write(newrecords, output_handle, 'genbank')


# Main function #
def main(input, output, type, overwrite=False):
    print("Input: %s\n"
          "Output: %s \n"
          "Type: %s\n"
          %(input, output, type))

    if type == "file":
        if output.endswith(".fna"):
            gbk_to_fna(input, output)
        else:
            gbkfile_to_faa(input, output, overwrite=overwrite)
    elif type == "folder":
        create_folder(output)
        print("Folder of gbk files:" + input)
        print("Folder for faa files: " + output)
        gbkfiles = list_gbk_files(input)
        gbkfiles.sort()
        for gbkfile in gbkfiles:
            name = gbkfile.split("/")[-1]
            name = name[:-4]
            print("Processing: " + name)
            outfaafile = os.path.join(output, name + ".faa")
            gbkfile_to_faa(gbkfile,outfaafile)
            #break
    else:
        print("ERROR: invalid type. Please choose one of: file, folder")
    print("\nDone")

#'''
# Parsing arguments & Execution #
parser = argparse.ArgumentParser(description='Turns a gbk file into an faa file containing all AA sequences.')
parser.add_argument('-i', '--input', help='Input gbk file or folder containing gbks (if folder type). Full path',
                    required=True)
parser.add_argument('-o', '--output', help='Output faa file or folder (if folder type). Full path', required=True)
parser.add_argument('-t', '--type', help='file or folder', required=True)
if __name__ == "__main__":
    args = parser.parse_args()
    main(args.input, args.output, args.type)
#'''

#main("/media/hdd_2tb/phd_main/trimming_practice_images/Aka_trimmed_shorter.gbk",
#     "/media/hdd_2tb/phd_main/4_GPA_trimmed_clusters/Aka_trimmed_shorter.faa",
#     "file")

#main("/media/hdd_2tb/phd_main/4_GPA_trimmed_clusters",
#     "/media/hdd_2tb/phd_main/4_GPA_trimmed_clusters",
#     "folder")

### 09.05.22
#inputdir = "/media/hdd_2tb/phd_main/GPA_clusters/0_trimmed_GPA_clusters"
#outputdir = "/media/hdd_2tb/phd_main/GPA_clusters/4_trimmed_annot_GPA_BGC_faas"
#type = "folder"
#main(inputdir, outputdir, type)

'''
### 18.07.23 - for extracting specific sequences from Stammsammlung files
thedir = '/media/hdd_2tb/GPA_reclassification_clean/A_dataset_creation/1_positive_hits_downloads/Stammsamlung2022_hits_exports'
locus_to_file_dict = {'contig_13251':os.path.join(thedir, 'S001_full.gbk'),
                      'contig_3960':os.path.join(thedir, 'S003_full.gbk'),
                      'contig_15':os.path.join(thedir, 'S005_full.gbk'),
                      'contig_2130':os.path.join(thedir, 'S006_full.gbk'),
                      'contig_9053':os.path.join(thedir, 'S008_full.gbk'),
                      'contig_1900':os.path.join(thedir, 'S009_full.gbk'),
                      'contig_4262':os.path.join(thedir, 'S009_full.gbk'),
                      'contig_788':os.path.join(thedir, 'S009_full.gbk')}
for key, value in locus_to_file_dict.items():
    extract_locus_from_gbk(key, value)
'''


'''
def csv_to_dict(csvfile, key="", value=""):
    thedict ={}
    with open(csvfile, 'r') as infile:
        reader = csv.reader(infile)
        for headerline in reader:
            key_id = headerline.index(key)
            value_id = headerline.index(value)
            break
        for line in reader:
            if line[key_id] not in thedict.keys():
                thedict[line[key_id]] = [line[value_id]]
            else:
                #thedict[line[key_id]] = "%s|%s" %(thedict[line[key_id]], line[value_id])
                thelist = thedict[line[key_id]]
                thelist.append(line[value_id])
                thedict[line[key_id]] = thelist
    return thedict

def csv_to_list(csvfile, key=""):
    thelist =[]
    with open(csvfile, 'r') as infile:
        reader = csv.reader(infile)
        for headerline in reader:
            key_id = headerline.index(key)
            break
        for line in reader:
            thelist.append(line[key_id])
    return thelist

ssd_incsv = '/media/hdd_2tb/Documents_HDD/IMBI_PhD_rep/IBMI_PhD_Retreat_SSD_RGs.csv'
out_csv = '/media/hdd_2tb/Documents_HDD/IMBI_PhD_rep/IBMI_PhD_Retreat_SSD_RGs_assigned.csv'
groups_dict = csv_to_dict(ssd_incsv, key='Group', value='Lastname')
names_list = csv_to_list(ssd_incsv, key='Lastname')

sessions_dict = {}
for name in names_list:
    sessions_dict[name] = [0,0,0,0,0,0]
### we need 6 sessions
for i in range(6):
    print("Session %s" %(i+1))
    #pairs = []
    #pairs_dict = {}
    current_participants = []
    #finished_groups = []
    session_groups_dict = copy.deepcopy(groups_dict)
    ### 13 pairs each session
    for j in range(1,13):
        print("\tPair %s" %(j))
        group1 = ''
        group2 = ''
        ### random 1st group
        #while group1 == '':
        group1, members1 = random.choice(list(session_groups_dict.items()))
            #if group1 in finished_groups:
                #group1 = ''
        ### random 2nd group, different than the first if possible, except the last pair, that can be from the same group
        while group2 == '':
            group2, members2 = random.choice(list(session_groups_dict.items()))
            if group2 == group1:# and j <14: #or group2 in finished_groups:
                group2 = ''
        ### random pair from group and save participants
        person1 = ''
        person2 = ''
        #while person1 == '':
        person1 = random.choice(members1)
            #if person1 in current_participants:
                #person1 = ''
        current_participants.append(person1)
        #while person2 == '':
        person2 = random.choice(members2)
            #if person2 == person1:#in current_participants:
                #person2 = ''
        current_participants.append(person2)
        ### remove current people from dict for speeding up
        members1.remove(person1)
        session_groups_dict[group1] = members1
        if len(members1) == 0:
            del session_groups_dict[group1]
        members2.remove(person2)
        session_groups_dict[group2] = members2
        if len(members2) == 0:
            del session_groups_dict[group2]
        #thispair=[person1, person2]
        #pairs.append(thispair)
        #pairs_dict[person1] = j
        #pairs_dict[person2] = j
        ### save to session dict
        sessions_dict[person1][i] = j
        sessions_dict[person2][i] = j
        ### mark group as finished if necessary
        #if (set(members1).issubset(set(current_participants))):
        #    finished_groups.append(group1)
        #if (set(members2).issubset(set(current_participants))):
        #    finished_groups.append(group2)
        #break
    #pprint(sessions_dict)
    ### make last group:
    extra_persons = list(set(names_list).difference(current_participants))
    #if len(extra_persons) > 1:
    #    print("More leftover people than expected")
    #    print(extra_persons)
    #    exit()
    #extra_person = extra_persons[0]
    #pairs_dict[extra_person] = random.randint(1,14)
    for extra_person in extra_persons:
        sessions_dict[extra_person][i] = 13
    ### save session's pairs
    #sessions_dict[name] =pairs_dict
    #break
#pprint(sessions_dict)

ssd_df = pd.DataFrame.from_dict(sessions_dict, orient='index', columns=['SSD1', 'SSD2', 'SSD3', 'SSD4', 'SSD5', 'SSD6'])
ssd_df.to_csv(out_csv)


'''
```
