## Supplementary Data 5 for "Structural and evolutionary analyses support reclassification of glycopeptide antibiotics as xyclopeptides": generate_culp_table.py

```
#!/usr/bin/python3.6
# Created by Athina Gavriilidou at 13.06.23 16:07 #
# Imports #
import argparse
import os
import pandas as pd
import subprocess
import shutil
import math
from os.path import dirname as up
import dateutil.parser as dparser

# Functions #
def create_folder(path):
    if not os.path.exists(path):
        os.mkdir(path)
        print("Created folder: %s" %(path))

'''given a directory, lists all gbk files'''
def list_gbk_files(directory):
    gbk_files = []
    for file in os.listdir(directory):
        if file.endswith(".gbk"):
            gbk_files.append(os.path.join(directory, file))
    print("Found " + str(len(gbk_files)) + " .gbk files")
    return gbk_files

def execute_download(dlink, outfile):
    wget_command = 'wget -O %s %s' %(outfile, dlink)
    try:
        print("Using link: %s" %(dlink))
        os.system(wget_command)
    except:
        return dlink

def execute_gitclone(dlink, outdir):
    git_clone_command = 'git clone -v %s %s' %(dlink, outdir)
    try:
        print("Cloning repository from: %s" %(dlink))
        os.system(git_clone_command)
    except:
        return dlink

'''for copying files. Make everything=True and then everything from the source will be copied to the destination'''
def execute_cp(source_fileordir, destinationfileordir, everything=False, filterlist=[]):
    if len(filterlist) != 0:
        filelist = list_gbk_files(source_fileordir)
        filelist = [f for f in filelist if os.path.basename(f).split('_')[0] in filterlist]
        for source_filename in filelist:
            cp_command = 'cp %s %s/.' % (source_filename, destinationfileordir)
            try:
                os.system(cp_command)
            except:
                print("Warning. There was a problem with copying %s to %s" % (source_filename, destinationfileordir))
    if everything:
        cp_command = 'cp %s/* %s/.' %(source_fileordir,destinationfileordir)
    else:
        cp_command = 'cp %s %s' %(source_fileordir,destinationfileordir)
    try:
        os.system(cp_command)
    except:
        print("Warning. There was a problem with copying %s to %s" %(source_fileordir, destinationfileordir))


def execute_mv(source_fileordir, destinationfileordir, everything=False):
    if everything:
        mv_command = 'mv %s/* %s/.' % (source_fileordir, destinationfileordir)
    else:
        mv_command = 'mv %s %s' % (source_fileordir, destinationfileordir)
    try:
        os.system(mv_command)
    except:
        print("Warning. There was a problem with moving %s to %s" % (source_fileordir, destinationfileordir))

def download_files(accno, outfolder, attempts=0):
    home_directory = os.path.expanduser('~')
    ncbi_tool_path = os.path.join(home_directory, 'anaconda3/bin/ncbi-acc-download')
    #possible_formats_list = 'fasta,genbank,featuretable,gff3'.split(',')
    print("Trying to download genome of %s" %(accno))
    if attempts == 5:
        print("ERROR: could not download %s files" %(accno))
        return
    outfile = os.path.join(outfolder, '%s.gbk' %(accno))
    temp_file_loc = os.path.join(up(up(outfile)), os.path.basename(outfile))
    ### for the cases of the straightforward ids
    if "." in accno:
        command = '%s --extended-validation all -v --format genbank ' \
                  '--out %s %s' %(ncbi_tool_path, temp_file_loc, accno)
        try:
            result = subprocess.check_output(command, shell=True, stderr=subprocess.STDOUT)
        except FileNotFoundError:
            print('ERROR: is ncbi-acc-download installed?\n')
            exit()
        result = result.decode('utf-8')
        result = result.split('\n')[0]
        download_link = result.split(' ')[-1]
        ### create folder after the file is downloaded and name it correctly
        create_folder(outfolder)
        execute_mv(temp_file_loc, outfile)
        return [download_link, accno]
    ### for the rest of the cases, where there is a master record
    command = '%s --extended-validation all -v --format genbank ' \
              '--out %s --recursive %s' %(ncbi_tool_path, temp_file_loc, accno)
    try:
        result = subprocess.check_output(command, shell=True, stderr=subprocess.STDOUT)
    except FileNotFoundError:
        print('ERROR: is ncbi-acc-download installed?\n')
        exit()
    except:
        print("")
    if not os.path.exists(temp_file_loc):
        theindex = accno.rindex('1')
        newaccno = accno[:theindex] + accno[theindex:].replace('1', '0')
        print("Warning: Download failed for %s. Retrying for %s" %(accno, newaccno))
        return download_files(newaccno, outfolder.replace(accno, newaccno), attempts=attempts+1)
    result = result.decode('utf-8')
    result = result.split('\n')[0]
    download_link = result.split(' ')[-1]
    ### create folder after the file is downloaded and name it correctly
    create_folder(outfolder)
    execute_mv(temp_file_loc, outfile)
    return [download_link, accno]

def check_if_nan(thestring):
    try:
        if math.isnan(thestring):
            return ''
    except:
        return thestring


def append_to_note(note, toappend):
    if len(note) == 0:
        return toappend
    else:
        return '%s. %s' %(note, toappend)

def transform_series_to_string(seq_withXdom, keeponlyone=False):
    if type(seq_withXdom) is str:
        return seq_withXdom
    elif not keeponlyone:
        seq_withXdom = seq_withXdom.tolist()
        seq_withXdom = "|".join(x for x in seq_withXdom)
        return seq_withXdom
    else:
        seq_withXdom = seq_withXdom.tolist()
        return seq_withXdom[0]

def fill_meta_table(stable_df, positive_hits_ids, meta_df, download_dir, outtable, download_link, hmmhits_table,
                    downloadgenomes=False):
    lines_to_append = [] ### see download_mibig_dataset.py - same concept
    source = 'Culp'
    if downloadgenomes:
        home_directory = os.path.expanduser('~')
        ncbi_tool_path = os.path.join(home_directory,'anaconda3/bin/ncbi-acc-download')
        print("\nUsing tool under: %s\n" %(ncbi_tool_path))
    ### iterate over table
    for x in range(1,len(positive_hits_ids)+1):
        db_id = positive_hits_ids[x-1]#.zfill(2)
        i = int(db_id)
        note = ''
        #print(stable_df)
        hasgenome = stable_df.loc[i,'genome?'] ### can be Y or N
        if hasgenome == 'Y':
            hasgenome = True
        elif hasgenome == 'N':
            hasgenome = False
        cluster_accno = stable_df.loc[i,'Cluster accession']
        if hasgenome:
            ncbi_accno = stable_df.loc[i,'genome acc']
            if not ":"in cluster_accno:
                note = append_to_note(note, 'BGC AccNo: %s' %(cluster_accno))
        else:
            ncbi_accno = cluster_accno
            note = append_to_note(note, 'No genome available, NCBI AccNo corresponds to the BGC.')
        ### if downloadgenomes is activated, take care of it
        if downloadgenomes and hasgenome:
            outfolder = os.path.join(download_dir, '%s_genome_%s' %(db_id, ncbi_accno))
            dresult= download_files(ncbi_accno, outfolder)
            #print(dresult)
            #print(len(dresult))
            genomedownloadlink, downloaded_ncbi_accno = dresult
            if ncbi_accno != downloaded_ncbi_accno:
                note= 'Original AccNo: %s' %(ncbi_accno)
                ncbi_accno = downloaded_ncbi_accno
        ### covering also case of missing strain, genus, species
        genus = check_if_nan(stable_df.loc[i,'Genus'])
        species = check_if_nan(stable_df.loc[i,'Species'])
        strain = check_if_nan(stable_df.loc[i,'Strain'])
        product = stable_df.loc[i,'product']
        if product == 'unknown':
            product = ''
        ### covering case of synonyms
        if ',' in strain:
            try:
                strain, synonym = strain.split(',')
                note = append_to_note(note, 'Synonym strain: %s' %(synonym))
            except:
                strain = strain.split(',')[0]
                synonyms_list = strain.split(',')[1:]
                note = append_to_note(note, '%sSynonym strain: %s' %(note, ' '.join(s for s in synonyms_list)))
        ### covering case of vague naming
        if 'Actinobacteria bacterium' in strain:
            strain = strain[len('Actinobacteria bacterium') + 1:]
            note = append_to_note(note, 'Original name: Actinobacteria bacterium %s' %(strain))
        ### covering case of extra info in parenthesis
        if "(" in strain:
            strain, appendtonote = strain.split('(')
            appendtonote = appendtonote.split(")")[0]
            note = append_to_note(note,appendtonote)
        ### covering case of D30
        if 'D30' == strain:
            strain = 'uncultured soil bacterium clone D30'
        ### creating full strain string
        strain_string = ' '.join(f for f in [genus, species, strain] if f!='')
        ### getting search date and sequence with Xdomain
        dateofsearch = hmmhits_table.loc[db_id, 'search_date']
        dateofsearch = transform_series_to_string(dateofsearch, keeponlyone=True)
        seq_withXdom = hmmhits_table.loc[db_id, 'target']
        seq_withXdom = transform_series_to_string(seq_withXdom)
        #print(db_id, dateofsearch)
        ### fill in row list columns='
        # strain ID source NCBI_AccNo Database_ID seq_with_Xdom product genus date_of_search download_link notes'
        row_list = [strain_string, '', source, ncbi_accno, db_id, seq_withXdom, product, genus, dateofsearch, download_link, note]
        lines_to_append.append(row_list)
    ### Maybe in the future we want to add assigned ID to these
    print("Warning: generation of assigned ID has not been implemented. Column will be empty.")
    ### join existing and new rows
    meta_df_table = meta_df.values.tolist()
    lines_to_append = meta_df_table + lines_to_append
    new_df = pd.DataFrame(lines_to_append,
                          columns='strain ID source NCBI_AccNo Database_ID seq_with_Xdom product genus '
                                  'date_of_search download_link notes'.split(' '))
    ### save to table
    new_df.to_csv(outtable, index=False)
    #print(new_df.iloc[:5])
    print("\nTable has been created/updated: %s\n" %(outtable))

### get run date from last 10 lines of domtblout
def extract_date(last_tblout_lines):
    for line in last_tblout_lines.target_name:
        if "Date" in line:
            thedate = dparser.parse(line, fuzzy=True)
            return thedate.strftime('%d-%m-%Y')

### nice way to visualize hmmsearch domtblout
def import_hmmsearch_tblout(filepath):
    # manually specify col limits
    dom_hits_colspecs = [(0, 252), (252, 264), (264, 270), (270, 291), (291, 302), (302, 308), (308, 318), (318, 325),
                         (325, 331), (331, 335), (335, 339), (339, 349), (349, 359), (359, 366), (366, 372), (372, 378),
                         (378, 384), (384, 390), (390, 396), (396, 402), (402, 408), (408, 413), (413, 435)]
    tblout_df = pd.read_fwf(filepath, skiprows=3, index_col=False,
                                             colspecs=dom_hits_colspecs,
                                             names=['target_name', 'accession1', 'tlen', 'query_name', 'accession2',
                                                    'full_sequence_qlen', 'full_sequence_E-value',
                                                    'full_sequence_score', 'full_sequence_bias', 'this_domain_#',
                                                    'this_domain_of', 'this_domain_c-Evalue', 'this_domain_i-Evalue',
                                                    'this_domain_score', 'this_domain_bias', 'hmm_coord_from',
                                                    'hmm_coord_to', 'ali_coord_from', 'ali_coord_to', 'env_coord_from',
                                                    'env_coord_to', 'acc', 'description_of_target'])
    the_run_date = extract_date(tblout_df.iloc[-10:])
    ### skip version lines
    return tblout_df.iloc[:-10], the_run_date

# Main function #
def main(input, outtable, stable, downloadgenomes, domtbloutfile):
    print("\nFolder with gbk files: %s\n"
          "Output table: %s\n"
          "Supplementary Table 1: %s\n"
          "Genome download: %s\n"
          "HMMsearch domain summary file: %s\n"
          % (input, outtable, stable, downloadgenomes, domtbloutfile))

    create_folder(input)
    ### files needed can be downloaded here:
    waglechner_stable1_link = 'https://static-content.springer.com/esm/art%3A10.1038%2Fs41564-019-0531-5/' \
                              'MediaObjects/41564_2019_531_MOESM3_ESM.xlsx'
    waglechner_git_link = 'https://github.com/waglecn/GPA_evolution.git'

    hmmhits_table = pd.read_csv(domtbloutfile, index_col=False, dtype='str')
    hmmhits_table.set_index('db_id',inplace=True)
    positive_hits_ids= hmmhits_table.index.tolist()
    ### get the Stable1 if not provided
    if not os.path.exists(stable):
        execute_download(waglechner_stable1_link, stable)

    stable_df = pd.read_excel(stable, skiprows=1)
    stable_df.set_index('Cluster', inplace=True)
    #print(stable_df.iloc[:5])

    ### download the cluster gbks if not in directory already
    list_of_gbk_files = list_gbk_files(input)
    if len(list_of_gbk_files) == len(positive_hits_ids):
        print("\n%s BGC gbk files detected as expected. Skipping download\n" %(len(list_of_gbk_files)))
    else:
        print("\nMissing %s BGC gbk file(s). Repeating download of all files" %(len(positive_hits_ids) - len(list_of_gbk_files)))
        git_dir = os.path.join(input, 'GPA_evolution')
        execute_gitclone(waglechner_git_link, git_dir)
        #execute_cp(os.path.join(git_dir, 'input_cluster_sequences'), input, everything=True)
        execute_cp(os.path.join(git_dir, 'input_cluster_sequences'), input, everything=False, filterlist=positive_hits_ids)
        print("The following directory is not being used and can be deleted:\n%s\n" %(git_dir))

    ### read metadata table if it exists, otherwise create it
    if os.path.exists(outtable):
        meta_df = pd.read_csv(outtable)
    else:
        meta_df = pd.DataFrame(columns='strain ID source NCBI_AccNo Database_ID seq_with_Xdom product genus '
                                       'date_of_search download_link notes'.split(' '))

    ### start downloading genomes and filling the table
    fill_meta_table(stable_df, positive_hits_ids, meta_df, input, outtable, waglechner_git_link,
                    hmmhits_table, downloadgenomes)


    '''
    ### maybe make a folder per entry and then have separate folders for full genome and for cluster!!!!!!!!!!!!
    ### OR NOT!!!! we already have all clusters directly from his github. maybe just for checking?
    firstacc = stable_df.iloc[0,2]
    testout = os.path.join(input, firstacc)
    result = download_files(firstacc, testout)
    result = result.split('\n')[0]
    result = result.split(' ')[-1]
    print("Detected download link: %s" %(result))
    '''
    print("\nDone")


# Parsing arguments & Execution #
parser = argparse.ArgumentParser(description='Generates a metadata table for the Culp dataset (Culp et al, 2020)')
parser.add_argument('-i', '--input', help='Input: Directory where all the gbk files will be downloaded (clusters '
                                          'from the Culp publication', required=True)
parser.add_argument('-o', '--output', help='Output metadata table', required=True)
parser.add_argument('-t', '--table', help='Supplementary table 1 (xlsx) of the Culp publication - will be downloaded '
                                          'in this path if not provided', required=True)
parser.add_argument('-d', '--domtblout', help='Summary domtblout (tsv) of HMM search on the clusters', required=True)
parser.add_argument('-g', '--genomes', help='Specify as True for genome download of NEW entries, where possible. '
                                            'Default is False',
                    required=False, default=False)
if __name__ == "__main__":
    args = parser.parse_args()
    main(args.input, args.output, args.table, args.genomes, args.domtblout)
```
