## Supplementary Data 5 for "Structural and evolutionary analyses support reclassification of glycopeptide antibiotics as xyclopeptides": hmmsearch_localdataset.py

```
#!/usr/bin/python3.6
# Created by Athina Gavriilidou at 29.06.23 15:56 #
# Imports #
import argparse
import os
import pandas as pd
import subprocess
import shutil
import math
from os.path import dirname as up
import csv
from pprint import pprint
import re
import dateutil.parser as dparser

# Functions #

def create_folder(path):
    if not os.path.exists(path):
        os.mkdir(path)
        print("Created folder: %s" %(path))

'''given a directory, lists all gbk files'''
def list_gbk_files(directory):
    gbk_files = []
    for file in os.listdir(directory):
        if file.endswith(".gbk"):
            gbk_files.append(os.path.join(directory, file))
    print("Found " + str(len(gbk_files)) + " .gbk files")
    gbk_files.sort()
    return gbk_files

'''given a directory, lists all faa files. I know it's redundant give me a break'''
def list_faa_files(directory):
    faa_files = []
    for file in os.listdir(directory):
        if file.endswith(".faa"):
            faa_files.append(os.path.join(directory, file))
    print("Found " + str(len(faa_files)) + " .faa files")
    faa_files.sort()
    return faa_files

def list_files(directory, suffix):
    files = []
    for file in os.listdir(directory):
        if file.endswith(suffix):
            files.append(os.path.join(directory, file))
    print("Found %s %s files" %(len(files), suffix))
    files.sort()
    return files


def execute_download(dlink, outfile):
    wget_command = 'wget -O %s %s' %(outfile, dlink)
    try:
        print("Using link: %s" %(dlink))
        os.system(wget_command)
    except:
        return dlink

def execute_gitclone(dlink, outdir):
    git_clone_command = 'git clone -v %s %s' %(dlink, outdir)
    try:
        print("Cloning repository from: %s" %(dlink))
        os.system(git_clone_command)
    except:
        return dlink

'''for copying files. Make everything=True and then everything from the source will be copied to the destination'''
def execute_cp(source_fileordir, destinationfileordir, everything=False):
    if everything:
        cp_command = 'cp %s/* %s/.' %(source_fileordir,destinationfileordir)
    else:
        cp_command = 'cp %s %s' %(source_fileordir,destinationfileordir)
    try:
        os.system(cp_command)
    except:
        print("Warning. There was a problem with copying %s to %s" %(source_fileordir, destinationfileordir))


def execute_mv(source_fileordir, destinationfileordir, everything=False):
    if everything:
        mv_command = 'mv %s/* %s/.' % (source_fileordir, destinationfileordir)
    else:
        mv_command = 'mv %s %s' % (source_fileordir, destinationfileordir)
    try:
        os.system(mv_command)
    except:
        print("Warning. There was a problem with moving %s to %s" % (source_fileordir, destinationfileordir))

'''save lines in csv format in the given output file'''
def save_lines_csv(lines, output):
    with open(output, 'w') as f:
        tsv_output = csv.writer(f)
        for line in lines:
            tsv_output.writerow(line)
    print("\nCreated file: " + output)

def append_to_file (sourcfile, destinationfile):
    os.system('cat %s >> %s' %(sourcfile, destinationfile))

def add_to_fullreport(hmmdomoutput, full_report_lines, accno):
    # [['accno','target_name','accession', 'c-evalue','i-evalue','score','bias']]
    ### get date info
    with open(hmmdomoutput, 'r') as infile:
        for line in infile:
            if line.startswith('# Date:'):
                thedate = dparser.parse(line, fuzzy=True)
                thedate = thedate.strftime('%d-%m-%Y')

    with open(hmmdomoutput, 'r') as infile:
        for skipline in infile:
            break
        for headline in infile:
            targetnameend, acc2start  = [m.start() for m in re.finditer('accession', headline)]
            acc2end = acc2start + len("accession")
            cvalstart = headline.find(' c-Evalue')
            cvalend = cvalstart + len(' c-Evalue')
            ivalstart = headline.find(' i-Evalue')
            ivalend = ivalstart + len(' i-Evalue')
            description_start = headline.find('description of target')
            score1start, score2start = [m.start() for m in re.finditer(' score', headline)]
            score2end = score2start + len(' score')
            bias1start, bias2start = [m.start() for m in re.finditer(' bias', headline)]
            bias2end = bias2start + len(' bias')
            break
        for skipline in infile:
            break
        for line in infile:
            ### if table finishes, break iteration
            if line.startswith('#'):
                break
            ### get info from line
            target_name = line[0:targetnameend]
            accession = line[acc2start:acc2end]
            cevalue = line[cvalstart:cvalend]
            ievalue = line[ivalstart:ivalend]
            score = line[score2start:score2end]
            bias = line[bias2start:bias2end]
            description = line[description_start:]
            summaryline = [accno, target_name, accession, cevalue, ievalue, score, bias, description, thedate]
            summaryline = [r.strip() for r in summaryline]
            full_report_lines.append(summaryline)
    return full_report_lines

def extract_domhits(domhitsfile, summary_domhitsfile, accno):
    result_lines = add_to_fullreport(domhitsfile, [], accno)
    with open(summary_domhitsfile, 'a') as infile:
        writer = csv.writer(infile)
        for line in result_lines:
            writer.writerow(line)
    return len(result_lines)

def execute_hmmsearch(proteinfasta, outputprefix, xdomain_hmm_path, output):
    summary_domhitsfile = os.path.join(output, "hmmsearch_summary_dom_hits.tsv")
    summary_concat_hitsfile = os.path.join(output, "hmmsearch_summary_concat_hits.tsv")
    summary_concat_tbloutfile = os.path.join(output, "hmmsearch_summary_concat_tblout.tsv")
    summary_concat_pfamhitsfile = os.path.join(output, "hmmsearch_summary_concat_pfamhits.tsv")
    accno = os.path.basename(outputprefix)[:-1]
    hitsfile = outputprefix + 'hits'
    seqhitsfile = outputprefix + 'seq_hits.tsv'
    domhitsfile = outputprefix + 'dom_hits.tsv'
    pfamhitsfile = outputprefix + 'pfam_hits.tsv'
    #xdomain_hmm_path = '/sfs/7/workspace/ws/bcbga01-gpa_newclasses-0/hmm_search_repeat_2023/as6_Xdomain.hmm'
    hmmsearch_command = 'hmmsearch -o %s --tblout %s --domtblout %s --pfamtblout %s --cpu 30 --domT 300 %s \%s' \
                        %(hitsfile, seqhitsfile, domhitsfile, pfamhitsfile, xdomain_hmm_path, proteinfasta)
    if not os.path.exists(outputprefix + 'hits'):
        try:
            os.system(hmmsearch_command)
        except:
            print("WARNING: hmmsearch failed for %s" %(proteinfasta))
            #return
    if os.path.exists(outputprefix + 'hits'):
        ### after hmmsearch is done, save results
        append_to_file(hitsfile, summary_concat_hitsfile)
        append_to_file(seqhitsfile, summary_concat_tbloutfile)
        append_to_file(pfamhitsfile, summary_concat_pfamhitsfile)
        hits_no = extract_domhits(domhitsfile, summary_domhitsfile, accno)
    else:
        print("WARNING: HMMSEARCH FAILED FOR %s" %(proteinfasta))
        return "Failed"
    ### report success or no:
    if hits_no == 0:
        return False
    else:
        return True

def append_lines_csv(lines, output):
    with open(output, 'a') as f:
        tsv_output = csv.writer(f)
        for line in lines:
            tsv_output.writerow(line)
    print("\nAppended to file: " + output)

def touch_file(filepath):
    os.system('touch %s' %filepath)

def print_list_with_sep(thelist, thesep):
    thestring = thesep.join(thelist)
    print(thestring)

def search_dataset(dict_of_label_to_faafiles, hmmsearch_output, xdommodelfile, output, finishedtable):
    if not os.path.exists(finishedtable):
        finished_df = pd.DataFrame(columns=['x_dom_hit'])
        save_lines_csv([['db_id','x_dom_hit']], finishedtable)
    else:
        finished_df = pd.read_csv(finishedtable, dtype={'db_id':str})
        finished_df.set_index('db_id', inplace=True)
    print(finished_df)
    ### making sure we only run for anything not yet checked
    finished_ids = finished_df.index.values.tolist()
    if len(finished_ids) > 0:
        print("\nSkipping following entries as they have been searched already:")
        print_list_with_sep(finished_ids, ' ')

    ### initialize dom summary file if there isn't one yet
    summary_domhitsfile = os.path.join(output, "hmmsearch_summary_dom_hits.tsv")
    if not os.path.exists(summary_domhitsfile):
        save_lines_csv(['db_id target accession c-Evalue i-Evalue score bias description search_date'.split(' ')],
                       summary_domhitsfile)

    ### iterate over fasta files:
    for label, faafile in dict_of_label_to_faafiles.items():
        ### skip if file has already been searched
        if label in finished_ids:
            continue
        ### do HMMsearch
        outputprefix = os.path.join(hmmsearch_output, '%s_' % (label))
        foundhit = execute_hmmsearch(faafile, outputprefix, xdommodelfile, output)
        ### save info to finished table
        append_lines_csv([[label, foundhit]], finishedtable)
    print("\nA summary of the results can be found under:\n%s\n" % (summary_domhitsfile))

def get_prokka_files(directory, suffix='faa', type='files'):
    dict_of_label_to_faafiles = {}
    if type == 'files':
        for file in os.listdir(directory):
            if os.path.isfile(os.path.join(directory, file)) and file.endswith(suffix):
                dict_of_label_to_faafiles[file[:-len(suffix) +1]] = os.path.join(directory, file)
    elif type == 'folders':
        for folder in os.listdir(directory):
            if os.path.isdir(os.path.join(directory, folder)):
                for file in os.listdir(os.path.join(directory, folder)):
                    if os.path.isfile(os.path.join(os.path.join(directory, folder), file)) and file.endswith(suffix):
                        dict_of_label_to_faafiles[folder] = os.path.join(os.path.join(directory, folder), file)
    else:
        print("Error: what is %s?" %(type))
        exit()
    return dict_of_label_to_faafiles


# Main function #
def main(output, xdommodelfile, input, dtype, finished):
    print("\nFolder of HMMsearch: %s\n"
          "X domain HMM file: %s\n"
          "Input directory: %s\n"
          "Data type: %s\n"
          "Finished table: %s"
          % (output, xdommodelfile, input, dtype, finished))
    create_folder(output)

    thetype, fileorfolder, suffix = dtype.split("-")
    if 'prokka' in thetype:
        dict_of_label_to_faafiles = get_prokka_files(input, suffix=suffix, type=fileorfolder)
    else:
        fileslist = list_files(input, suffix)
        dict_of_label_to_faafiles = {}
        for f in fileslist:
            label = os.path.splitext(f)[0]
            dict_of_label_to_faafiles[label] = f


    ### start with hmmsearch
    hmmsearch_output = os.path.join(output, 'hmmer_full_results')
    create_folder(hmmsearch_output)
    search_dataset(dict_of_label_to_faafiles, hmmsearch_output, xdommodelfile, output, finished)
    print("\nDone")


# Parsing arguments & Execution #
parser = argparse.ArgumentParser(description='Given a local dataset, runs hmmsearch (or prokka first)')
parser.add_argument('-o', '--output', help='Output directory of HMMsearch', required=True)
parser.add_argument('-m', '--model', help='X domain HMM to be used in the search', required=True)
parser.add_argument('-i', '--input', help='Input directory of faa files or prokka results or fna files', required=True)
parser.add_argument('-t', '--type', help='Type of input data (prokka results, faa files, fna files) and organisation.'
                                         ' Can be e.g. prokka-folders-faa or fasta-files-fna', required=True)
parser.add_argument('-f', '--finished', help='Output finished table', required=True)
if __name__ == "__main__":
    args = parser.parse_args()
    main(args.output, args.model, args.input, args.type, args.finished)
```
