## Supplementary Data 5 for "Structural and evolutionary analyses support reclassification of glycopeptide antibiotics as xyclopeptides": hmmsearch_mgnify_asms.py

```
#!/usr/bin/python3.6
# Created by Athina Gavriilidou at 02.05.23 18:08 #
# Imports #
import argparse
import csv
import numpy as np
import pandas as pd
import os
import json
import re
from pprint import pprint
import time
import dateutil.parser as dparser

# Functions #
'''creates folder in given path, if it does not exist already'''
def create_folder(path):
    if not os.path.exists(path):
        os.mkdir(path)

'''save lines in csv format in the given output file'''
def save_lines_csv(lines, output):
    with open(output, 'w') as f:
        tsv_output = csv.writer(f)
        for line in lines:
            tsv_output.writerow(line)
    print("\nCreated file: " + output)

def append_lines_csv(lines, output):
    with open(output, 'a') as f:
        tsv_output = csv.writer(f)
        for line in lines:
            tsv_output.writerow(line)
    print("\nAppended to file: " + output)

def execute_download(path, outfile, iter=0):
    wget_command = 'wget -O %s %s' %(outfile, path)
    try:
        os.system(wget_command)
    except:
        ### added on 05.06.23 to cover cases where connection is lost
        if iter<5:
            time.sleep(5)
            iter+=1
            return execute_download(path, outfile, iter=iter)
        return path
    ### cover also case where download looks like it succeeded but didnt
    if os.stat(outfile).st_size == 0 and iter <5:
        time.sleep(5)
        iter += 1
        return execute_download(path, outfile, iter=iter)

def detect_json_link(tempjsonfile):
    with open(tempjsonfile, 'r') as jsonfile:
        jsoncontent = json.load(jsonfile)
        data = jsoncontent['data']
        for d in data:
            if "FASTA" in d['id'] and "_CDS" in d['id'] and "faa" in d['id']:
                cdsfaalink = d['links']['self']
                cdsfaalink = cdsfaalink.rstrip('?format=json')
                return cdsfaalink

    return ''

def execute_delete(path):
    ### added recursive option for lists
    if type(path) == list:
        for el in path:
            execute_delete(el)
    ### redirect errors to /dev/null for clearer output
    rm_command = 'rm %s 2> /dev/null' %(path)
    try:
        os.system(rm_command)
    except:
        return path

def extract_domhits(domhitsfile, summary_domhitsfile, accno):
    result_lines = add_to_fullreport(domhitsfile, [], accno)
    with open(summary_domhitsfile, 'a') as infile:
        writer = csv.writer(infile)
        for line in result_lines:
            writer.writerow(line)
    return len(result_lines)

def append_to_file (sourcfile, destinationfile):
    os.system('cat %s >> %s' %(sourcfile, destinationfile))

def add_to_fullreport(hmmdomoutput, full_report_lines, accno):
    ### get date info
    with open(hmmdomoutput, 'r') as infile:
        for line in infile:
            if line.startswith('# Date:'):
                thedate = dparser.parse(line, fuzzy=True)
                thedate = thedate.strftime('%d-%m-%Y')
    # [['accno','target_name','accession', 'c-evalue','i-evalue','score','bias']]
    with open(hmmdomoutput, 'r') as infile:
        for skipline in infile:
            break
        for headline in infile:
            targetnameend, acc2start  = [m.start() for m in re.finditer('accession', headline)]
            acc2end = acc2start + len("accession")
            cvalstart = headline.find(' c-Evalue')
            cvalend = cvalstart + len(' c-Evalue')
            ivalstart = headline.find(' i-Evalue')
            ivalend = ivalstart + len(' i-Evalue')
            description_start = headline.find('description of target')
            score1start, score2start = [m.start() for m in re.finditer(' score', headline)]
            score2end = score2start + len(' score')
            bias1start, bias2start = [m.start() for m in re.finditer(' bias', headline)]
            bias2end = bias2start + len(' bias')
            break
        for skipline in infile:
            break
        for line in infile:
            ### if table finishes, break iteration
            if line.startswith('#'):
                break
            ### get info from line
            target_name = line[0:targetnameend]
            accession = line[acc2start:acc2end]
            cevalue = line[cvalstart:cvalend]
            ievalue = line[ivalstart:ivalend]
            score = line[score2start:score2end]
            bias = line[bias2start:bias2end]
            description = line[description_start:]
            summaryline = [accno, target_name, accession, cevalue, ievalue, score, bias, description, thedate]
            summaryline = [r.strip() for r in summaryline]
            full_report_lines.append(summaryline)
    return full_report_lines

def execute_hmmsearch(proteinfasta, outputprefix, xdomain_hmm_path, output):
    summary_domhitsfile = os.path.join(output, "hmmsearch_summary_dom_hits.tsv")
    summary_concat_hitsfile = os.path.join(output, "hmmsearch_summary_concat_hits.tsv")
    summary_concat_tbloutfile = os.path.join(output, "hmmsearch_summary_concat_tblout.tsv")
    summary_concat_pfamhitsfile = os.path.join(output, "hmmsearch_summary_concat_pfamhits.tsv")
    accno = os.path.basename(outputprefix)[:-1]
    hitsfile = outputprefix + 'hits'
    seqhitsfile = outputprefix + 'seq_hits.tsv'
    domhitsfile = outputprefix + 'dom_hits.tsv'
    pfamhitsfile = outputprefix + 'pfam_hits.tsv'
    #xdomain_hmm_path = '/sfs/7/workspace/ws/bcbga01-gpa_newclasses-0/hmm_search_repeat_2023/as6_Xdomain.hmm'
    hmmsearch_command = 'hmmsearch -o %s --tblout %s --domtblout %s --pfamtblout %s --cpu 30 --domT 300 %s \%s' \
                        %(hitsfile, seqhitsfile, domhitsfile, pfamhitsfile, xdomain_hmm_path, proteinfasta)
    if not os.path.exists(outputprefix + 'hits'):
        try:
            os.system(hmmsearch_command)
            execute_delete(proteinfasta)
        except:
            execute_delete(proteinfasta)
            #return
    if os.path.exists(outputprefix + 'hits'):
        ### after hmmsearch is done, save results
        append_to_file(hitsfile, summary_concat_hitsfile)
        append_to_file(seqhitsfile, summary_concat_tbloutfile)
        append_to_file(pfamhitsfile, summary_concat_pfamhitsfile)
        hits_no = extract_domhits(domhitsfile, summary_domhitsfile, accno)
    else:
        print("WARNING: HMMSEARCH FAILED FOR %s" %(proteinfasta))
        return "Failed"
    ### report success or no:
    if hits_no == 0:
        return False
    else:
        return True
    ### delete files
    #execute_delete([hitsfile, seqhitsfile, pfamhitsfile, domhitsfile])

def search_mgnify_fromlinks(filtered_accno_to_link_dict, output, model, finished, failed_file):
    accno_to_hit_lists = []
    output_tmp_fastas = os.path.join(output, 'temp_fastas')
    create_folder(output_tmp_fastas)
    output_hmmer_results = os.path.join(output, 'hmmer_full_results')
    create_folder(output_hmmer_results)
    output_tmp_jsons = os.path.join(output, 'temp_jsons')
    create_folder(output_tmp_jsons)
    list_of_not_found = []
    ### initialize dom summary file if there isn't one yet
    summary_domhitsfile = os.path.join(output, "hmmsearch_summary_dom_hits.tsv")
    if not os.path.exists(summary_domhitsfile):
        save_lines_csv(['db_id target accession c-Evalue i-Evalue score bias description search_date'.split(' ')],
                       summary_domhitsfile)
    ### parse links
    for accno, down_dict in filtered_accno_to_link_dict.items():
        downlink = down_dict['downloadlink']
        print(accno, downlink)
        ### get json file with info and link to fasta
        tempjsonfile = os.path.join(output_tmp_jsons, "%s.json" %(accno))
        execute_download(downlink, tempjsonfile)
        fastalink = detect_json_link(tempjsonfile)
        execute_delete(tempjsonfile)
        if fastalink == '':
            print("WARNING: LINK NOT FOUND FOR %s" %(accno))
            list_of_not_found.append(accno)
            append_lines_csv([[accno]], failed_file)
            continue
            #exit()
        ### get fasta file
        tempfastafile = os.path.join(output_tmp_fastas, "%s_FASTA_predicted_cds.faa.gz" %(accno))
        execute_download(fastalink, tempfastafile)
        ### do HMMsearch
        outputprefix = os.path.join(output_hmmer_results, '%s_' %(accno))
        foundhit = execute_hmmsearch(tempfastafile, outputprefix, model, output)
        execute_delete(tempfastafile)
        ### save info to finished table
        accno_to_hit_lists.append([accno, foundhit])
        append_lines_csv([[accno, foundhit]], finished)
        # for testing
        #break

    print("\n A summary of the results can be found under:\n%s\n" % (summary_domhitsfile))
    if len(list_of_not_found) > 0:
        print("\nThe following accession numbers did not have a protein fasta file:")
        pprint(list_of_not_found)
        notfound_file = os.path.join(output, 'notfoundfastaAccnos.csv')
        append_lines_csv(list_of_not_found, notfound_file)
    return accno_to_hit_lists

# Main function #
def main(input,finished, output, model):
    print("\nInput table: %s\n"
          "Finished table: %s\n"
          "Output folder: %s\n"
          "HMM model: %s\n"
          %(input, finished, output, model))
    create_folder(output)
    input_df = pd.read_csv(input, index_col=0)
    #print(input_df)
    if not os.path.exists(finished):
        finished_df = pd.DataFrame(columns=['x_dom_hit'])
        save_lines_csv([['assembly','x_dom_hit']], finished)
    else:
        finished_df = pd.read_csv(finished, index_col=0)
    ### added 05.06.23 to keep track of failed jobs
    failed_file = os.path.join(output, 'failed_runs_mgnify_accnos.csv')
    if not os.path.exists(failed_file):
        save_lines_csv([['assembly']], failed_file)
    # finished_df.loc['1'] = ['test']
    # finished_df.index.name = 'assembly'
    ### making sure we only run for anything not yet checked
    toremoveids = input_df.index.intersection(finished_df.index)
    filtered_df = input_df.drop(toremoveids)
    filtered_accno_to_link_dict = filtered_df.to_dict('index')
    ### starting the search in the assemblies
    accno_to_hit_lists = search_mgnify_fromlinks(filtered_accno_to_link_dict, output, model, finished,
                                                 failed_file)
    ### appending result to finished searches table
    #append_lines_csv(accno_to_hit_lists, finished)
    print("\nDone")


# Parsing arguments & Execution #
parser = argparse.ArgumentParser(description='')
parser.add_argument('-i', '--input', help='Input table of assembly IDs and download links', required=True)
parser.add_argument('-f', '--finished', help='Table of already checked assembly IDs', required=True)
parser.add_argument('-o', '--output', help='Output folder', required=True)
parser.add_argument('-m', '--model', help='Path to HMM file to be used for search', required=True)
if __name__ == "__main__":
    args = parser.parse_args()
    main(args.input, args.finished, args.output, args.model)
```
