## Supplementary Data 5 for "Structural and evolutionary analyses support reclassification of glycopeptide antibiotics as xyclopeptides": hmmsearch_ncbi_ftp.py

```
#!/usr/bin/python3.6
# Created by Athina Gavriilidou at 08.03.23 15:17 #
# Imports #
import argparse
import os
from pprint import pprint
import csv
import re
import pandas as pd
import dateutil.parser as dparser


# Functions #

def execute_delete(path):
    ### added recursive option for lists
    if type(path) == list:
        for el in path:
            execute_delete(el)
    ### redirect errors to /dev/null for clearer output
    rm_command = 'rm %s 2> /dev/null' %(path)
    try:
        os.system(rm_command)
    except:
        return path

def append_to_file (sourcfile, destinationfile):
    os.system('cat %s >> %s' %(sourcfile, destinationfile))

def extract_domhits(domhitsfile, summary_domhitsfile, accno):
    result_lines = add_to_fullreport(domhitsfile, [], accno)
    with open(summary_domhitsfile, 'a') as infile:
        writer = csv.writer(infile)
        for line in result_lines:
            writer.writerow(line)
    return len(result_lines)


def execute_hmmsearch(proteinfasta, outputprefix, xdomain_hmm_path, output):
    summary_domhitsfile = os.path.join(output, "hmmsearch_summary_dom_hits.tsv")
    summary_concat_hitsfile = os.path.join(output, "hmmsearch_summary_concat_hits.tsv")
    summary_concat_tbloutfile = os.path.join(output, "hmmsearch_summary_concat_tblout.tsv")
    summary_concat_pfamhitsfile = os.path.join(output, "hmmsearch_summary_concat_pfamhits.tsv")
    accno = os.path.basename(outputprefix)[:-1]
    hitsfile = outputprefix + 'hits'
    seqhitsfile = outputprefix + 'seq_hits.tsv'
    domhitsfile = outputprefix + 'dom_hits.tsv'
    pfamhitsfile = outputprefix + 'pfam_hits.tsv'
    #xdomain_hmm_path = '/sfs/7/workspace/ws/bcbga01-gpa_newclasses-0/hmm_search_repeat_2023/as6_Xdomain.hmm'
    hmmsearch_command = 'hmmsearch -o %s --tblout %s --domtblout %s --pfamtblout %s --cpu 30 --domT 300 %s \%s' \
                        %(hitsfile, seqhitsfile, domhitsfile, pfamhitsfile, xdomain_hmm_path, proteinfasta)
    if not os.path.exists(outputprefix + 'hits'):
        try:
            os.system(hmmsearch_command)
        except:
            print("WARNING: hmmsearch failed for %s" % (proteinfasta))
            # return
    if os.path.exists(outputprefix + 'hits'):
        ### after hmmsearch is done, save results
        append_to_file(hitsfile, summary_concat_hitsfile)
        append_to_file(seqhitsfile, summary_concat_tbloutfile)
        append_to_file(pfamhitsfile, summary_concat_pfamhitsfile)
        hits_no = extract_domhits(domhitsfile, summary_domhitsfile, accno)
    else:
        print("WARNING: HMMSEARCH FAILED FOR %s" % (proteinfasta))
        return "Failed"
    ### report success or no:
    if hits_no == 0:
        return False
    else:
        return True

'''save lines in csv format in the given output file'''
def save_lines_csv(lines, output):
    with open(output, 'w') as f:
        tsv_output = csv.writer(f)
        for line in lines:
            tsv_output.writerow(line)
    print("\nCreated file: " + output)

'''taken from https://stackoverflow.com/questions/2170900/get-first-list-index-containing-sub-string'''
def index_containing_substring(the_list, substring):
    for i, s in enumerate(the_list):
        #print([s, substring, i])
        if substring in s or substring == s:
              return i
    return -1

'''creates folder in given path, if it does not exist already'''
def create_folder(path):
    if not os.path.exists(path):
        os.mkdir(path)

'''creates dictionary out of table'''
def csv_to_dict(filepath, key, value, delim=None, skiplines=0):
    if delim is None:
        if filepath.endswith(".csv"):
            delim = ","
        else:
            delim = "\t"
    content = {}
    with open(filepath) as fd:
        rd = csv.reader(fd, delimiter=delim, quotechar='"')
        while skiplines !=0:
            for skipline in rd:
                break
            skiplines-=1
        for headline in rd:
            if not isinstance(key,int):
                key = index_containing_substring(headline, key)
                #print(key)
            if not isinstance(value, int):
                value = index_containing_substring(headline, value)
                #print(value)
            break
        for line in rd:
            if type(value) == list:
                this_value = []
                for v in value:
                    this_value.append(line[v])
                content[line[key]] = this_value
            else:
                content[line[key]] = line[value]
    return content

def add_to_fullreport(hmmdomoutput, full_report_lines, accno):
    # [['accno','target_name','accession', 'c-evalue','i-evalue','score','bias']]
    ### get date info
    with open(hmmdomoutput, 'r') as infile:
        for line in infile:
            if line.startswith('# Date:'):
                thedate = dparser.parse(line, fuzzy=True)
                thedate = thedate.strftime('%d-%m-%Y')

    with open(hmmdomoutput, 'r') as infile:
        for skipline in infile:
            break
        for headline in infile:
            targetnameend, acc2start  = [m.start() for m in re.finditer('accession', headline)]
            acc2end = acc2start + len("accession")
            cvalstart = headline.find(' c-Evalue')
            cvalend = cvalstart + len(' c-Evalue')
            ivalstart = headline.find(' i-Evalue')
            ivalend = ivalstart + len(' i-Evalue')
            description_start = headline.find('description of target')
            score1start, score2start = [m.start() for m in re.finditer(' score', headline)]
            score2end = score2start + len(' score')
            bias1start, bias2start = [m.start() for m in re.finditer(' bias', headline)]
            bias2end = bias2start + len(' bias')
            break
        for skipline in infile:
            break
        for line in infile:
            ### if table finishes, break iteration
            if line.startswith('#'):
                break
            ### get info from line
            target_name = line[0:targetnameend]
            accession = line[acc2start:acc2end]
            cevalue = line[cvalstart:cvalend]
            ievalue = line[ivalstart:ivalend]
            score = line[score2start:score2end]
            bias = line[bias2start:bias2end]
            description = line[description_start:]
            summaryline = [accno, target_name, accession, cevalue, ievalue, score, bias, description, thedate]
            summaryline = [r.strip() for r in summaryline]
            full_report_lines.append(summaryline)
    return full_report_lines

def execute_download(dlink, outfile):
    wget_command = 'wget -O %s %s' %(outfile, dlink)
    try:
        print("Using link: %s" %(dlink))
        os.system(wget_command)
    except:
        return dlink

def download_assembly_summary_table(input):
    ### get link
    if 'refseq' in input:
        thelink='https://ftp.ncbi.nlm.nih.gov/genomes/ASSEMBLY_REPORTS/assembly_summary_refseq.txt'
    elif 'genbank' in input:
        thelink = 'https://ftp.ncbi.nlm.nih.gov/genomes/ASSEMBLY_REPORTS/assembly_summary_genbank.txt'
    else:
        print("\n ERROR: could not recognize if genbank or refseq. Please provide the table or a file path with "
              "either of the two keywords (refseq|genbank) in the name\n")
        exit()
    execute_download(thelink, input)

def find_nth(haystack, needle, n):
    start = haystack.find(needle)
    while start >= 0 and n > 1:
        start = haystack.find(needle, start+len(needle))
        n -= 1
    return start


def touch_file(filepath):
    os.system('touch %s' %filepath)

'''very specific for test case'''
def filter_dict(original_dict, keys_to_keep):
    keys_to_keep = [k[:find_nth(k,'_',2)] for k in keys_to_keep]
    #pprint(keys_to_keep)
    new_dict = {}
    for key, value in original_dict.items():
        if key in keys_to_keep:
            new_dict[key] = value
    return new_dict

def print_list_with_sep(thelist, thesep):
    thestring = thesep.join(thelist)
    print(thestring)

def append_lines_csv(lines, output):
    with open(output, 'a') as f:
        tsv_output = csv.writer(f)
        for line in lines:
            tsv_output.writerow(line)
    print("\nAppended to file: " + output)

def search_ncbi_database(accno_to_folderpath_dict,output_tmp_fastas, output_hmmer_results,hmmmodel, output,
                         finishedtable):
    if not os.path.exists(finishedtable):
        finished_df = pd.DataFrame(columns=['x_dom_hit'])
        save_lines_csv([['db_id','x_dom_hit']], finishedtable)
    else:
        finished_df = pd.read_csv(finishedtable, dtype={'db_id':str})
        finished_df.set_index('db_id', inplace=True)
    print(finished_df)
    ### addition on 22.09.2023
    finished_df = finished_df[finished_df['x_dom_hit'] != 'Failed']
    ### making sure we only run for anything not yet checked
    finished_ids = finished_df.index.values.tolist()
    if len(finished_ids) > 0:
        print("\nSkipping following entries as they have been searched already:")
        print_list_with_sep(finished_ids, ' ')

    ### initialize dom summary file if there isn't one yet
    summary_domhitsfile = os.path.join(output, "hmmsearch_summary_dom_hits.tsv")
    if not os.path.exists(summary_domhitsfile):
        save_lines_csv(['db_id target accession c-Evalue i-Evalue score bias description search_date'.split(' ')],
                       summary_domhitsfile)

    ### initialize not found file if there isn't one yet
    notfoundfile = os.path.join(output, "notfound_assemblies.csv")
    if not os.path.exists(notfoundfile):
        save_lines_csv(['assembly_accno search_date'.split(' ')],
                       notfoundfile)
    notfound_already_list = list(csv_to_dict(notfoundfile, 'assembly_accno', 'search_date',',').keys())
    ### for testing only:
    counter = 0
    ### iterate over assemblies
    for accno, ftpfolderpath in accno_to_folderpath_dict.items():
        ### skip if file has already been searched
        if accno in finished_ids:
            continue
        foldername = ftpfolderpath.split("/")[-1]
        protein_fasta_path = ftpfolderpath + '/' + foldername + '_protein.faa.gz'
        tempfile = os.path.join(output_tmp_fastas, foldername + '_protein.faa.gz')
        hmmoutput_prefix = os.path.join(output_hmmer_results, foldername + "_")
        ### try to get their protein seqs
        if not os.path.exists(hmmoutput_prefix + 'hits'):
            try:
                execute_download(protein_fasta_path, tempfile)
            except:
                if accno not in notfound_already_list:
                    append_lines_csv([[accno]], notfoundfile)
                continue
        else:
            'hmmsearch already done. skipping download...'
        ### hmmsearch for X domain
        foundhit = execute_hmmsearch(tempfile, hmmoutput_prefix, hmmmodel, output)
        ### save info to finished table
        append_lines_csv([[accno, foundhit]], finishedtable)
        ### now remove downloaded file to avoid overload
        execute_delete(tempfile)
        ### for testing only
        if counter ==5:
           break

        print("\nA summary of the results can be found under:\n%s\n" % (summary_domhitsfile))

# Main function #
def main(input, output, hmmmodel, finishedtable):
    print("\nInput assembly summary table: %s\n"
          "Output folder of intermediate and final files: %s\n"
          "HMM model: %s\n"
          "Finished table: %s\n"
          %(input, output, hmmmodel, finishedtable))
    create_folder(output)
    output_tmp_fastas = os.path.join(output, 'temp_fastas')
    create_folder(output_tmp_fastas)
    output_hmmer_results = os.path.join(output, 'hmmer_full_results')
    create_folder(output_hmmer_results)
    ### if the table is not there, download it
    if not os.path.exists(input):
        download_assembly_summary_table(input)
    accno_to_folderpath_dict = csv_to_dict(input, 'assembly_accession', 'ftp_path', delim='\t', skiplines=1)
    ### for testing only
    #successful_subset_table = '/media/hdd_2tb/GPA_reclassification_clean/hmmsearch_tests/hmm_search_repeat_2023/refseq_hmmsearch/summary_dom_hits.tsv'
    #accno_to_targetname_dict = csv_to_dict(successful_subset_table, 'accno', 'targetname')
    #accno_to_folderpath_dict = filter_dict(accno_to_folderpath_dict, accno_to_targetname_dict.keys())

    ### start search
    search_ncbi_database(accno_to_folderpath_dict, output_tmp_fastas, output_hmmer_results,
                                             hmmmodel, output, finishedtable)

    #full_report_lines= [['accno','target_name','accession', 'c-evalue','i-evalue','score','bias','description']]
    #full_report_file = os.path.join(output, '0_full_report_hmmsearch_dom.csv')
    #counter = 1 #for testing
    ### start summary files
    #summary_domhitsfile = os.path.join(output, "summary_dom_hits.tsv")
    ##summary_concat_hitsfile = os.path.join(output, "summary_concat_hits.tsv")
    #summary_concat_tbloutfile = os.path.join(output, "summary_concat_tblout.tsv")
    #summary_concat_pfamhitsfile = os.path.join(output, "summary_concat_pfamhits.tsv")
    #touch_file(summary_domhitsfile)
    #touch_file(summary_concat_pfamhitsfile)
    #touch_file(summary_concat_hitsfile)
    #touch_file(summary_concat_tbloutfile)
    #save_lines_csv(full_report_lines, summary_domhitsfile)


    #print("\nNumber of hits: %s" %(len(full_report_lines) -1))
    #save_lines_csv(full_report_lines, full_report_file)

    #print("\n A summary of the results can be found under:\n%s\n" %(summary_domhitsfile))

    #if len(list_of_not_found) > 0:
    #    print("\nThe following accession numbers did not have a protein fasta file:")
    #    pprint(list_of_not_found)
    print("\nDone")


# Parsing arguments & Execution #
parser = argparse.ArgumentParser(description='Iteratively runs hmmsearch of X domain on refseq or genbank database '
                                             'and saves the result in tables.')
parser.add_argument('-i', '--input', help='Input assembly summary table of refseq or genbank (needs to include a '
                                          'keyword - refseq or genbank - and be tab-delimited). If '
                                          'the file does not exist, it will be downloaded on this path.', required=True)
parser.add_argument('-o', '--output', help='Output folder for temporary protein fasta files and hmmsearch result '
                                           'tables', required=True)
parser.add_argument('-m', '--model', help='Path to HMM file to be used for search', required=True)
parser.add_argument('-f', '--finished', help='Table of already checked entries. Will be created here if it does not'
                                             ' exist', required=True)
if __name__ == "__main__":
    args = parser.parse_args()
    main(args.input, args.output, args.model, args.finished)
```
