## Supplementary Data 5 for "Structural and evolutionary analyses support reclassification of glycopeptide antibiotics as xyclopeptides": make_concat_phylogeny.py

```
#!/usr/bin/python3.6
# Created by Athina Gavriilidou at 20.09.23 10:44 #
# Imports #
import argparse
import os
import pandas as pd
from pprint import pprint
from Bio import SeqIO
import random
from Bio.SeqRecord import SeqRecord
from Bio.Seq import Seq
import csv
import subprocess

### keeping process random but repeatable
random.seed(10)

# Functions #
def create_folder(path):
    if not os.path.exists(path):
        os.mkdir(path)
        print("Created folder: %s" %(path))

'''given a directory, lists all files, with a certain suffix'''
def list_files(directory, suffix= "", suppress=True):
    the_files = []
    for file in os.listdir(directory):
        if file.endswith(suffix):
            the_files.append(os.path.join(directory, file))
    if not suppress:
        print("Found %s %s files" %(len(the_files), suffix))
    the_files.sort()
    return the_files

def copy_file(sourcefile, destinationfile):
    command = "cp '%s' '%s'" %(sourcefile, destinationfile)
    os.system(command)
    #print(command)

def find_reffile(msa_list):
    #print("\nNote: assuming modules_7 or X_domains group exists and contains all bgcs")
    print("\nNote: assuming X_domains group exists and contains all bgcs")
    substrings = ['modules_7', 'X_domains']
    #substrings = ['X_domains']
    for substring in substrings:
        for el in msa_list:
            if substring in el:
                return el
    print("No file found with %s in name" %(substring))
    exit()

def get_ids(fastafile):
    fasta_ids = []
    fasta_sequences = SeqIO.parse(open(fastafile), 'fasta')
    for fasta in fasta_sequences:
        name = fasta.id
        fasta_ids.append(name)
    return fasta_ids

### randomly choose 70% of alignments to concatenate
def get_random_subsamples(list_of_files, percentage=0.7):
    print("Generating subsamples containing randomly %s per cent of alignments" %(percentage*100))
    k = round(percentage * len(list_of_files))
    list_of_100subsamples = []
    for i in range(100):
        included = False
        while not included:
            sampled_list = random.sample(list_of_files, k)
            if set(sampled_list) not in list_of_100subsamples:
                list_of_100subsamples.append(sampled_list)
                included = True
    if len(list_of_100subsamples) == 100:
        print("Generated 100 random subsamples\n")
        return list_of_100subsamples
    else:
        print("ERROR: 100 subsamples not created")
        exit()

def get_strlist_of_filenames(file_list, replace_str='_trimalMSA.faa'):
    filenames = [os.path.basename(f).replace(replace_str,'') for f in file_list]
    #filenamesstr = ''
    #for f in filenames:
    #    filenamesstr = filenamesstr + f + "|"
    filenamesstr = '|'.join(filenames)
    return filenamesstr

def fasta_to_dict(fastafile):
    fasta_id_to_seq_dict = {}
    fasta_sequences = SeqIO.parse(open(fastafile), 'fasta')
    for fasta in fasta_sequences:
        name, sequence = fasta.id, fasta.seq
        name = name.split("_GPA_")[0]
        fasta_id_to_seq_dict[name] = str(sequence)
    return fasta_id_to_seq_dict

def append_msa(msa_file, subsample_fasta_dict):
    msa_id_to_seq_dict = fasta_to_dict(msa_file)
    ### addition 20.01.2023 for cases where MSAs are not relabeled
    msa_fixedid_to_seq_dict = {k.split("_GPA_")[0]:v for k, v in msa_id_to_seq_dict.items()}
    new_subsample_fasta_dict = {}
    for id, seq in msa_id_to_seq_dict.items():
        msa_length = len(seq)
        break
    for id, finalseq in subsample_fasta_dict.items():
        try:
            seq = msa_fixedid_to_seq_dict[id] ### used to be msa_id_to_seq_dict
        except:
            print("%s seq not found in %s" %(id, os.path.basename(msa_file)))
            seq = '-'*msa_length
        new_subsample_fasta_dict[id] = finalseq + seq
    return new_subsample_fasta_dict

def dict_to_seqrecords(subsample_fasta_dict):
    seqrecords_list = []
    for id, seq in subsample_fasta_dict.items():
        sr = SeqRecord(Seq(seq), id, id, id)
        seqrecords_list.append(sr)
    return seqrecords_list

'''save lines in csv format in the given output file'''
def save_lines_csv(lines, output):
    with open(output, 'w') as f:
        tsv_output = csv.writer(f)
        for line in lines:
            tsv_output.writerow(line)
    print("\nCreated file: " + output)

'''generate random subsampled MSAs from the candidates.'''
def generate_random_subsampled_msas(metadata_df, candidates_trimmed_msas_list, output, random_subsample_table):
    xdom_msa_file = find_reffile(candidates_trimmed_msas_list)
    ids = get_ids(xdom_msa_file)
    #print(ids)
    list_of_100subsamples = get_random_subsamples(candidates_trimmed_msas_list)
    list_of_100concatmsas = []
    table_of_contents = [['subsample_ID', 'list_of_msa_files']]
    counter = 1
    ### creating MSA files
    for subsample in list_of_100subsamples:
        table_of_contents.append([counter, get_strlist_of_filenames(subsample)])
        ### create empty dict with all IDs
        subsample_fasta_dict = {}
        for id in ids:
            subsample_fasta_dict[id] = ''
        ### start adding seqs to dict
        for msa_file in subsample:
            subsample_fasta_dict = append_msa(msa_file, subsample_fasta_dict)
        ### turn dict into list of seqrecords
        subsample_seqrecords = dict_to_seqrecords(subsample_fasta_dict)
        ### save subsample values in a fasta file
        outsamplemsa = os.path.join(output, '%s_random_concat_msa.faa' % (counter))
        with open(outsamplemsa, "w") as output_handle:
            SeqIO.write(subsample_seqrecords, output_handle, "fasta")
            print("Created file %s" % (outsamplemsa))
            list_of_100concatmsas.append(outsamplemsa)
        counter += 1
    #pprint(table_of_contents)
    save_lines_csv(table_of_contents, random_subsample_table)
    taxon_file = os.path.join(os.path.dirname(random_subsample_table), '1_random_concatsubsampled_taxalist.txt')
    with open(taxon_file, 'w') as infile:
        writer = csv.writer(infile)
        for i in ids:
            writer.writerow([i])
    #
    return list_of_100concatmsas, taxon_file, ids

def execute_fasttree(protein_alignment, outtree):
    fasttree_path = '/home/athina/anaconda3/envs/fasttree/bin/fasttree'
    fasttree_command = "%s -wag -gamma -out %s %s" %(fasttree_path, outtree, protein_alignment)
    print(fasttree_command)
    try:
        os.system(fasttree_command)
    except:
        print("ERROR: could not execute fasttree. Check that the path is correct.")
        print(fasttree_path)
        exit()

def is_file_empty(file_path):
    """
    Check if an existing file has size zero.

    Parameters:
    - file_path (str): Path to the file.

    Returns:
    - bool: True if the file has size zero, False otherwise.
    """
    if not os.path.exists(file_path):
        print("File does not even exist: %s" % file_path)
        return True
    return os.path.getsize(file_path) == 0


def execute_sumtrees(inputdir, extoutput, taxonfilepath):
    ### conda installation:
    sumtrees_path = '/home/athina/anaconda3/envs/dendropy/bin/sumtrees.py'
    ### conda installation on python 3.5
    #sumtrees_path = '/home/athina/anaconda3/envs/sumtrees_special/bin/sumtrees_experimental.py'
    #sumtrees_path = '/home/athina/anaconda3/envs/sumtrees_special/bin/python /home/athina/anaconda3/envs/dendropy/bin/sumtrees.py'
    ### pip installation with added path to python 3.4:
    #sumtrees_path = '/home/athina/anaconda3/envs/sumtrees_special/bin/python /home/athina/anaconda3/bin/sumtrees.py'
    ### removed  -m 30
    sumtrees_command = (("%s   -m 30 --percentages --input-format newick --preserve-underscores "
                        "--extended-output %s --replace --taxon-name-filepath %s --force-unrooted "
                        "--output-tree-format newick --output-tree-filepath=%s.sumtrees %s/*.tree")
                        %(sumtrees_path, extoutput, taxonfilepath, extoutput, inputdir))
    print(sumtrees_command)
    try:
        os.system(sumtrees_command)
        print("\nCreated file: %s.sumtrees" %(extoutput))
    except:
        print("ERROR: could not execute sumtrees.py. Check that the path is correct.")
        print(sumtrees_path)
        exit()
    ### check if it worked and stop if not
    if is_file_empty("%s.sumtrees"%(extoutput)):
        print("ERROR: sumtrees process has failed. Output file is empty.")
        exit()

'''modified using chatgpt'''
def execute_sumtrees_chatgpt(inputdir, extoutput, taxonfilepath):
    sumtrees_path = '/home/athina/anaconda3/envs/dendropy/bin/sumtrees.py'

    # Read the content of the taxon file
    with open(taxonfilepath, 'r') as taxon_file:
        taxon_content = taxon_file.read()

    sumtrees_command = (
        f"{sumtrees_path} -m 30 --percentages --input-format newick --preserve-underscores "
        f"--extended-output {extoutput} --replace --taxon-names {taxon_content} --force-unrooted "
        f"--output-tree-format newick --output-tree-filepath={extoutput}.sumtrees {inputdir}/*.tree"
    )

    print(sumtrees_command)

    try:
        os.system(sumtrees_command)
        print(f"\nCreated file: {extoutput}.sumtrees")
    except Exception as e:
        print(f"ERROR: could not execute sumtrees.py. Check that the path is correct.")
        print(sumtrees_path)
        print(e)
        exit()


def execute_genetreetk(supported_tree, gene_tree, logfile):
    splits_tsv = logfile.replace('.log','.tsv')
    genetreetk_path = ('/media/hdd_2tb/GPA_reclassification_clean/A_dataset_creation/6_concat_phylogeny/GeneTreeTk'
                       '/bin/genetreetk')
    genetreetk_command = "%s supported_splits --tree1 %s --tree2 %s --split_file %s --min_support 80" \
                         %(genetreetk_path, supported_tree, gene_tree, splits_tsv)
    print(genetreetk_command)
    try:
        with open(logfile, 'w') as out:
            return_code = subprocess.call(genetreetk_command.split(' '), stdout=out)
        #os.system(genetreetk_command)
    except:
        print("ERROR: could not execute fasttree. Check that the path is correct.")
        print(genetreetk_path)
        exit()

'''parses genetreetk compare_trees log file for the Weight of well-support splits in common between T1 and T2'''
def get_splits_value(logfile):
    rfile = open(logfile, 'r')
    splits_value = ''
    for line in rfile:
        line = line.strip()
        if 'Fraction of common well-supported splits: ' in line:
            splits_value = line.replace('Fraction of common well-supported splits: ','')
            #print([splits_value])
            ### added try except on 23.01.23
            try:
                splits_value = float(splits_value)
            ### Exception after detecting 'Fraction of common well-supported splits' happens when 0 splits in common
            except:
                splits_value = 0
    rfile.close()
    if splits_value == '':
        print('Warning: could not detect Fraction of common well-supported splits in file: ',logfile)
        print("Setting value to 0 for failed process (happened in cases of 4 sequences only)")
        splits_value = 0
    return splits_value

'''For the preprint, this is what was chosen: -wag -gamma'''
def execute_iqtree_withtest(mfafile, outprefix, model='WAG+G'):
    iqtreepath = '/home/athina/anaconda3/envs/iqtree2/bin/iqtree2'
    print("Assuming path to iqtree: %s" % (iqtreepath))
    if model is 'test':
        model_str = ("-m MFP  --mset Dayhoff,DCMut,JTT,JTTDCMut,LG,Poisson,PMB,Q.pfam,VT,WAG,"
                     "C10,C20,C30,C40,C50,C60,EX2,EX3,EHO,UL2,UL3,EX_EHO,LG4M,LG4X,CF4 "
                     "--madd C10,C20,C30,C40,C50,C60,EX2,EX3,EHO,UL2,UL3,EX_EHO,LG4M,LG4X,CF4")
    else:
        print("Using given model: %s" %(model))
        model_str = '-m %s' %(model)
    # --msub nuclear
    iqtree_command = (("%s -s %s -pre %s -redo -bb 1000 -nt AUTO %s")
                      %(iqtreepath, mfafile, outprefix, model_str))
    print(iqtree_command)
    try:
        os.system(iqtree_command)
    except:
        print("\nERROR: could not run iqtree. Are you sure it is installed and in the path?\n")
        exit()

def execute_mad(unrootedtreefile):
    madpath = '/home/athina/Documents/programs/mad/mad'
    print("Assuming path to MAD: %s" % (madpath))
    madcommand = "%s %s -t" %(madpath, unrootedtreefile)
    try:
        os.system(madcommand)
        print("Creating rooted tree in file: %s.rooted" %(unrootedtreefile))
    except:
        print("\nERROR: could not run MAD. Are you sure it is installed and in the path?\n")
        exit()

'''copy passed msas into separate dir and use to create concatenated phylogeny'''
def create_concat_phylogeny_from_list(passed_msas_list, outputprefix, bgcids, replace=False):
    ### create empty dict with all IDs
    subsample_fasta_dict = {}
    for id in bgcids:
        subsample_fasta_dict[id] = ''
    ### start adding seqs to dict
    for msa_file in passed_msas_list:
        subsample_fasta_dict = append_msa(msa_file, subsample_fasta_dict)
    ### turn dict into list of seqrecords
    subsample_seqrecords = dict_to_seqrecords(subsample_fasta_dict)
    ### save subsample values in a fasta file
    outmsa = '%s_MSA.faa'%(outputprefix)
    with open(outmsa, "w") as output_handle:
        SeqIO.write(subsample_seqrecords, output_handle, "fasta")
        print("Created file %s\n" % (outmsa))
    ### then iqtree with model testing and manually check which cameout the best (one of C10-60 perhaps?)
    if not os.path.exists('%s.treefile' %(outputprefix)) or replace:
        execute_iqtree_withtest(outmsa, outputprefix)

def deal_with_problematic_id(folder, targetid, newid):
    print("Dealing with problematic ID. Changing %s to %s" % (targetid, newid))
    if os.path.isdir(folder):
        command = 'cd %s; for file in *; do echo $file; sed -i -e \'s/%s/%s/g\' $file; done' %(folder, targetid, newid)
    else:
        command = 'echo %s; sed -i -e \'s/%s/%s/g\' %s; done' %(folder, targetid, newid, folder)
    print(command)
    os.system(command)


# Main function #
def main(input, output, metadata, replace=False):
    print("\nInput directory of basic phylogeny: %s\n"
          "Output folder: %s\n"
          "Metadata table: %s\n"
          % (input, output, metadata))
    create_folder(os.path.dirname(output))
    create_folder(output)
    metadata_df = pd.read_csv(metadata, sep='\t', index_col=0)
    #print(metadata_df)
    ### filter table and keep only groups suitable for consideration
    print("\nConsidering only groups that are found in at least 90% of the clusters. Duplications are acceptable in"
          " maximum 5% of the clusters in a group, but they are filtered out before inclusion.\n")
    metadata_df = metadata_df[metadata_df['percent_BGCs_present'] > 0.9]
    metadata_df = metadata_df[metadata_df['percent_duplications'] < 0.05]
    candidates_list = metadata_df.index.to_list()
    #print(metadata_df)
    #exit()
    #print(candidates_list)
    ### collect candidate trimmed MSAs and copy in another folder in the output
    candidates_trimmed_msas_folder = os.path.join(output, '0_candidates_trimal_MSAs')
    create_folder(candidates_trimmed_msas_folder)
    trimmed_MSAs_dir = os.path.join(input, '2_trimal_MSAs')
    print("\nLooking for trimmed MSAs in: %s" %(trimmed_MSAs_dir))
    alltrimmed_msas_list = list_files(trimmed_MSAs_dir)
    candidates_trimmed_msas_list = []
    for tmsa in alltrimmed_msas_list:
        thename = os.path.basename(tmsa).replace('_trimalMSA.faa','')
        #print(thename)
        found_name = False
        for cand in candidates_list:
            if found_name:
                break
            if cand == thename:
                newmsafile = os.path.join(candidates_trimmed_msas_folder, os.path.basename(tmsa))
                candidates_trimmed_msas_list.append(newmsafile)
                copy_file(tmsa, newmsafile)
                found_name = True
                break
            else:
                if thename.startswith("%s_" %(cand)):
                    newmsafile = os.path.join(candidates_trimmed_msas_folder, os.path.basename(tmsa))
                    candidates_trimmed_msas_list.append(newmsafile)
                    copy_file(tmsa, newmsafile)
                    found_name = True
                    break
    if len(candidates_list) == len(candidates_trimmed_msas_list):
        print("\nFound and copied %s candidate trimmed MSAs" %(len(candidates_trimmed_msas_list)))
    else:
        print("ERROR: could not find all candidate trimmed MSAs (%s out of %s). Aborting..."
              %(len(candidates_trimmed_msas_list),len(candidates_list)))
        pprint(candidates_trimmed_msas_list)
        for cand in candidates_list:
            newmsafile = os.path.join(candidates_trimmed_msas_folder, os.path.basename(cand))
            if newmsafile not in candidates_trimmed_msas_list:
                print('Missing',cand)
        pprint(candidates_list)
        exit()
    ### detected problem with StRiR_1 / StRir_1 so fixing:
    deal_with_problematic_id(candidates_trimmed_msas_folder, 'StRir_1', 'StRirr_1')
    ### Create random subsampled trees based on trimmed MSAs
    random_subsamples_dir = os.path.join(output, '1_random_concatsubsampled_MSAs')
    create_folder(random_subsamples_dir)
    random_subsample_table = os.path.join(output, '1_random_concatsubsampled_membership.csv')
    list_of_100concatmsas, taxon_file, bgcids = generate_random_subsampled_msas(metadata_df,
                                                                                candidates_trimmed_msas_list,
                                                                                random_subsamples_dir,
                                                                                random_subsample_table)
    ### detected problem with StRiR_1 / StRir_1 so fixing:
    deal_with_problematic_id(taxon_file, 'StRir_1', 'StRirr_1')

    print("\nConcatenated MSA files prepared. Starting phylogenetic analyses with fasttree v2.1.11 ...\n")
    ### calculating phylogenies
    random_subsamples_trees_dir = os.path.join(output, '2_random_concatsubsampled_trees')
    create_folder(random_subsamples_trees_dir)
    print("\nNote: WAG + G model used for all concatenated ML phylogenies\n")
    for msa_fasta in list_of_100concatmsas:
        outtree = os.path.join(random_subsamples_trees_dir, os.path.basename(msa_fasta).replace('_msa.faa','.tree'))
        if not os.path.exists(outtree) or replace:
            execute_fasttree(msa_fasta, outtree)
    ### getting sumtrees from dendropy v4.6.1
    congruence_check_dir = os.path.join(output, '3_congruence_check')
    create_folder(congruence_check_dir)
    print("\nCalculating consensus tree with split support values using dendropy's (v4.6.1) sumtrees function...")
    sumtree_output_prefix = os.path.join(congruence_check_dir,'randomsubsample_consensus')
    sumtree_output_file = "%s.sumtrees" %(sumtree_output_prefix)
    if not os.path.exists(sumtree_output_file) or replace:
        execute_sumtrees(random_subsamples_trees_dir, sumtree_output_prefix, taxon_file)
    ### add check for sumtrees here - if file does not exist or is empty then stop the process
    if is_file_empty("%s.sumtrees" % (sumtree_output_prefix)):
        print("ERROR: sumtrees process has failed. Output file is empty.")
        exit()


    ### get also their gene trees
    candidates_trimmed_trees_folder = os.path.join(output, '0_candidates_trimal_iqtree_MLtrees')
    create_folder(candidates_trimmed_trees_folder)
    trimmed_trees_dir = os.path.join(input, '5_trimal_iqtree_ML_trees')
    print("\nLooking for trimmed MSA gene trees in: %s" % (trimmed_trees_dir))
    alltrimmed_trees_list = list_files(trimmed_trees_dir, suffix='.treefile')
    if len(alltrimmed_trees_list) < len(candidates_trimmed_msas_list):
        print("\nERROR: failed to locate all trimmed MSA gene trees. Aborting...")
        exit()
    candidates_genetrees_list = []
    for trimmedtree in alltrimmed_trees_list:
        thename = os.path.basename(trimmedtree).replace("_trimal_iqtree.treefile", "")
        found_name = False
        for cand in candidates_list:
            if found_name:
                break
            if thename in cand:
                # print(thename)
                newtrimmedtree = os.path.join(candidates_trimmed_trees_folder, os.path.basename(trimmedtree))
                copy_file(trimmedtree, newtrimmedtree)
                candidates_genetrees_list.append(newtrimmedtree)
                found_name = True
                break
            else:
                if thename.startswith("%s_" % (cand)):
                    # print(thename)
                    newtrimmedtree = os.path.join(candidates_trimmed_trees_folder, os.path.basename(trimmedtree))
                    copy_file(trimmedtree, newtrimmedtree)
                    candidates_genetrees_list.append(newtrimmedtree)
                    found_name = True
                    break
    # pprint(candidates_genetrees_list)
    ### detected problem with StRiR_1 / StRir_1 so fixing:
    deal_with_problematic_id(candidates_trimmed_trees_folder, 'StRir_1', 'StRirr_1')
    ### compare gene tree congruence
    print("\nCalculating congruence values using the genetreetk function supported_splits "
          "(gitgub repo https://github.com/dparks1134/GeneTreeTk cloned on 20.09.2023)...")
    logfilelist = []
    for genetree in candidates_genetrees_list:
        splitstsv = os.path.join(congruence_check_dir,
                               os.path.basename(genetree).replace(".treefile", '_ssplits.tsv'))
        logfile = splitstsv.replace(".tsv", '.log')
        if not os.path.exists(logfile) or replace:
            execute_genetreetk(sumtree_output_file, genetree, logfile)
        logfilelist.append(logfile)
    ### create summary file from all the output files
    print("\nCollecting info from log files...")
    summary_lines = [['gene_tree', 'fraction_of_splits', 'gene_passed']]
    for logfile in logfilelist:
        if 'modules_3' in os.path.basename(logfile) or 'AMP_binding_domains_3' in os.path.basename(logfile):
            con_threshold = get_splits_value(logfile)
            break
    #con_threshold = 0.5
    print(
        "IMPORTANT Note: assuming congruence metric (fraction of supported splits) threshold to be defined by 3rd module or "
        "3rd A-domain: %s" % (con_threshold))
    ### make a report and copy passed MSAs to another folder
    passed_msas_dir = os.path.join(output, '4_congruent_trimal_MSAs')
    create_folder(passed_msas_dir)
    passed_msas_list = []
    for logfile in logfilelist:
        gene_tree_name = os.path.basename(logfile).replace('_trimal_iqtree_ssplits.log', '')
        fraction_of_splits = get_splits_value(logfile)
        if fraction_of_splits > con_threshold:
            gene_passed = 'yes'
            gene_trimmed_msa = os.path.join(trimmed_MSAs_dir, '%s_trimalMSA.faa' %(gene_tree_name))
            newmsafile = os.path.join(passed_msas_dir, os.path.basename(gene_trimmed_msa))
            copy_file(gene_trimmed_msa, newmsafile)
            passed_msas_list.append(newmsafile)
        else:
            gene_passed = 'no'
        thisline = [gene_tree_name, fraction_of_splits, gene_passed]
        summary_lines.append(thisline)
    ### check that only the congruent MSAs are in the directory
    if len(passed_msas_list) != len(list_files(passed_msas_dir)):
        print("ERROR: Number of MSA files is not equivalent to congruent genes. Aborting...")
        pprint(passed_msas_list)
        exit()
    outcongruence_summarytable = os.path.join(output, '3_congruence_check_summary.csv')
    save_lines_csv(summary_lines, outcongruence_summarytable)
    ### detected problem with StRiR_1 / StRir_1 so fixing:
    deal_with_problematic_id(passed_msas_dir, 'StRir_1', 'StRirr_1')
    #### create concatenated phylogeny from all that passed check
    outputprefix = os.path.join(output, '5_concatenated_phylogeny')
    create_concat_phylogeny_from_list(passed_msas_list, outputprefix, bgcids)
    #### MAD-root the tree
    unrootedtreefile = "%s.treefile" %(outputprefix)
    execute_mad(unrootedtreefile)
    print("\nDone")


# Parsing arguments & Execution #
parser = argparse.ArgumentParser(description='Identifies groups suitable for a congruence check, conducts the check '
                                             'and builds the concatenated phylogeny')
parser.add_argument('-i', '--input', help='Input basic phylogeny folder. Assumed folder hierarchy as created by the '
                                          'script run_basic_phylogeny.py', required=True)
parser.add_argument('-o', '--output', help='Output folder', required=True)
parser.add_argument('-m', '--metadata', help='Input metadata table (.tsv) of the groups analysed in the input folder',
                    required=True)
if __name__ == "__main__":
    args = parser.parse_args()
    main(args.input, args.output, args.metadata)
```
