## Supplementary Data 5 for "Structural and evolutionary analyses support reclassification of glycopeptide antibiotics as xyclopeptides": prepare_ogs_for_phylogeny.py

```
#!/usr/bin/python3.6
# Created by Athina Gavriilidou at 15.09.23 09:53 #
# Imports #
import argparse
import os
import csv
from pprint import pprint
import pandas as pd
from Bio import SeqIO

# Functions #
def create_folder(path):
    if not os.path.exists(path):
        os.mkdir(path)
        print("Created folder: %s" %(path))

def csv_to_list(csvfile, colname):
    thelist = []
    with open(csvfile, 'r') as infile:
        reader = csv.reader(infile)
        for headerline in reader:
            if type(colname) is str:
                key_id = headerline.index(colname)
            else:
                key_id = colname
            break
        for line in reader:
            thelist.append(line[key_id])
    return thelist

def check_that_isnt_nrps(listtocheck, nrpslist):
    for el in listtocheck:
        if el in nrpslist:
            return False
    return True

def check_if_unique(bgcids_to_labels_dict):
    #pprint(bgcids_to_labels_dict)
    newdict = {}
    for bgcid, labelslist in bgcids_to_labels_dict.items():
        if len(labelslist) > 1:
            newdict[bgcid] = labelslist
    return newdict

def separate_mono_poly_seqs(fastafile, output, headersep="|", replace=False):
    #print(fastafile)
    records = list(SeqIO.parse(fastafile, 'fasta'))
    singles_file = os.path.join(output, os.path.basename(fastafile)[:-4] + '_singles.fa')
    doublesfile = os.path.join(output, os.path.basename(fastafile)[:-4] + '_multies.fa')
    presence_dict = {}
    record_count = 0
    ### figure out how many copies of the gene each cluster has
    for record in records:
        theid = record.id.split(headersep)[0]
        if theid not in presence_dict.keys():
            presence_dict[theid] = 1
        else:
            presence_dict[theid] = presence_dict[theid] + 1
        record_count += 1
    ### save all the singles and non singles in new fasta files
    singles_list = []
    doubleslist = []
    singles_id_list = []
    #print(presence_dict)
    for record in records:
        theid = record.id.split(headersep)[0]
        if presence_dict[theid] == 1:
            singles_list.append(record)
            singles_id_list.append(record.id)
        else:
            doubleslist.append(record)
    ### have a check for extreme cases
    if len(doubleslist)/2 > 0.1*record_count + 1:
        print("\nWARNING: too many duplicates in %s\n" % (fastafile))
    elif len(doubleslist) == 0:
        return
    ### save records
    if not os.path.exists(singles_file) or replace:
        SeqIO.write(singles_list, singles_file, 'fasta')
        print("Created file: %s" % (singles_file))
    else:
        print("File already exists: %s" %(singles_file))
    if not os.path.exists(doublesfile) or replace:
        SeqIO.write(doubleslist, doublesfile, 'fasta')
        print("Created file: %s" % (doublesfile))
    else:
        print("File already exists: %s" % (doublesfile))
    #print(singles_file, doublesfile, singles_id_list, record_count)
    return singles_file, doublesfile, singles_id_list, record_count


def execute_diamond_makedb(infasta, replace=False):
    outdb = "%s_db.dmnd" %(infasta[:-3])
    ### for saving some time in multiple runs:
    if not replace and os.path.exists(outdb):
        return outdb
    diamond_command = 'diamond makedb --in %s --db %s --threads 30' %(infasta, outdb)
    print(diamond_command)
    try:
        os.system(diamond_command)
    except:
        print("\nERROR: could not run diamond makedb. Are you sure diamond is installed and in the path?\n")
        exit()
    return outdb

def execute_diamond_blastp(dbfile, queryfile, replace=False):
    outtablefile = "%s_scores.tsv" %(queryfile[:-3])
    ### for saving some time in multiple runs:
    if not replace and os.path.exists(outtablefile):
        return outtablefile
    diamond_command = 'diamond blastp --db %s --query %s --out %s --outfmt 6 qseqid sseqid evalue positive ppos ' \
                      '--threads 30' %(dbfile, queryfile, outtablefile)
    print(diamond_command)
    try:
        os.system(diamond_command)
    except:
        print("\nERROR: could not run diamond blastp. Are you sure diamond is installed and in the path?\n")
        exit()
    return outtablefile

def calculate_avg_distances(scores_table_file, headersep='|', replace=False):
    #### get list(s) from file
    scores_table = pd.read_csv(scores_table_file, sep='\t', names='qseqid,sseqid,evalue,positive,ppos'.split(','))
    scores_table['bgc_id'] = scores_table.qseqid.apply(lambda x: x.split(headersep)[0])
    #print(scores_table)
    qseqid_to_bgc_id_df = scores_table[['qseqid', 'bgc_id']]
    qseqid_to_bgc_id_df = qseqid_to_bgc_id_df.drop_duplicates()
    #print(qseqid_to_bgc_id_df)
    avg_scores_table = scores_table[['qseqid','ppos']].groupby('qseqid').mean()
    avg_scores_table = avg_scores_table.reset_index(level=0)
    #print(avg_scores_table)
    ### organized results:
    avg_scores_table_full = pd.merge(avg_scores_table, qseqid_to_bgc_id_df, on='qseqid')
    avg_scores_table_full = avg_scores_table_full.set_index('qseqid', drop=True)
    #print(avg_scores_table_full)
    avg_scores_out = scores_table_file.replace("_multies_scores","_multies_avg_scores")
    if not os.path.exists(avg_scores_out) or replace:
        avg_scores_table_full.to_csv(avg_scores_out, sep='\t')
        print("Created file: %s" %(avg_scores_out))
    else:
        print("File already exists: %s" %(avg_scores_out))
    #print(avg_scores_table_full.groupby(['bgc_id'], sort=False)['ppos'].max())
    idx = avg_scores_table_full.groupby(['bgc_id'])['ppos'].transform(max) == avg_scores_table_full['ppos']
    #print(idx)
    chosen_seqs_list = idx[idx == True].index.to_list()
    ### save choice in file for future reference
    outchoice = scores_table_file.replace("_multies_scores","_kept_multies")
    #print(outchoice)
    #pprint(chosen_seqs_list)
    if not os.path.exists(outchoice) or replace:
        with open(outchoice, 'w') as outfile:
            writer = csv.writer(outfile,delimiter='\t')
            for seq in chosen_seqs_list:
                writer.writerow([seq])
        print("Created file: %s" %(outchoice))
    else:
        print("File already exists: %s" %(outchoice))
    ### 06.12.2032 added exception for manually filtered
    if ((os.path.basename(outchoice).startswith("OG_15_") or os.path.basename(outchoice).startswith("OG_16_"))
            and os.path.exists(outchoice.replace(".tsv", "_manual_v2.csv"))):
        manualoutfile = outchoice.replace(".tsv", "_manual_v2.csv")
        print("Warning! Manual override - using table: %s\n Assuming vanRS are OGs 15 and 16"
              %(manualoutfile))
        chosen_seqs_list = []
        with open(manualoutfile, 'r') as infile:
            reader = csv.reader(infile)
            for line in reader:
                chosen_seqs_list.append(line[0])

    #print(chosen_seqs_list)
    return chosen_seqs_list

def filter_duplicates(doubles_bgcid_to_labels_dict, bgcids_to_labels_dict, ogfaafile, filtering_dir, special_case="",
                      replace=False):
    #print(ogfaafile)
    singles_file, doublesfile, singles_id_list, record_count = separate_mono_poly_seqs(ogfaafile, filtering_dir,
                                                                                       replace=replace)
    singles_db = execute_diamond_makedb(singles_file, replace)
    doubles_scores_file = execute_diamond_blastp(singles_db, doublesfile, replace)
    try:
        chosen_doubles_list = calculate_avg_distances(doubles_scores_file, replace=replace)
    except:
        print("ERROR: Irregular scores table. This can happen due to very unequal sequence lengths. File: %s"
              %(doubles_scores_file))
        exit()
    if special_case == 'oxyE':
        print("Special case of oxyE - assuming OG_3")
        #print(chosen_doubles_list)
        #print(len(chosen_doubles_list))
        #print(len(doubles_bgcid_to_labels_dict))
        #print(len(bgcids_to_labels_dict))
        bgcids_to_labels_dict = {}
        for bgcid, current_labels in doubles_bgcid_to_labels_dict.items():
            #print(current_labels)
            for curlabel in current_labels:
                if curlabel not in chosen_doubles_list:
                    bgcids_to_labels_dict[bgcid] = [curlabel]
                    #pprint(bgcids_to_labels_dict)
                    break
        #print(len(bgcids_to_labels_dict))
        #exit()
    else:
        for bgcid in doubles_bgcid_to_labels_dict.keys():
            current_labels = bgcids_to_labels_dict[bgcid]
            for curlabel in current_labels:
                if curlabel in chosen_doubles_list:
                    bgcids_to_labels_dict[bgcid] = [curlabel]
                    break
    return bgcids_to_labels_dict

'''check if also OG trees are necessary for filtering out duplicates like in preprint'''
def filter_copy_file(ogfaafile, oglabel, modules_version_dir, domains_version_dir, filtering_dir, headersep='|',
                     replace=False, special_case=''):
    seqs_list = []
    warningslist = []
    bgcids_to_labels_dict = {}
    try:
        records = list(SeqIO.parse(ogfaafile, 'fasta'))
    except:
        print(f"Warning: could not detect file {ogfaafile}\nPossibly OG with only one sequence. Skipping...")
        return None, None, []
    #print(ogfaafile)
    for rec in records:
        seqs_list.append(rec.id)
        bgcid = rec.id.split(headersep)[0]
        if bgcid not in bgcids_to_labels_dict.keys():
            bgcids_to_labels_dict[bgcid] = [rec.id]
        else:
            labellist = bgcids_to_labels_dict[bgcid]
            labellist.append(rec.id)
            bgcids_to_labels_dict[bgcid] = labellist
    ### check if there are enough sequences to continue
    if len(seqs_list) < 4:
        return None, None, []
    ### check if there are duplicates
    doubles_bgcid_to_labels_dict = check_if_unique(bgcids_to_labels_dict)
    #pprint(doubles_bgcid_to_labels_dict)
    duplicates_num = 0
    #print(doubles_bgcid_to_labels_dict)
    ### eliminate duplicates if necessary
    if len(doubles_bgcid_to_labels_dict) > 0:
        duplicates_num = len(doubles_bgcid_to_labels_dict)
        ### added check for how many duplicates - if more than 10% stop the process and show it. Manual work will be necessary
        ### added condition to only check big OGs
        if len(bgcids_to_labels_dict.keys()) > 10 and duplicates_num > len(bgcids_to_labels_dict.keys()) * 0.1:
            warningslist.append("WARNING: Too many duplicates (%s out of %s BGCs) found in file: %s"#\nManual check may be necessary. Aborting..."
                  %(duplicates_num,len(bgcids_to_labels_dict.keys()),ogfaafile))
            #pprint(doubles_bgcid_to_labels_dict)
            #exit()
        #print('filter')
        bgcids_to_labels_dict = filter_duplicates(doubles_bgcid_to_labels_dict, bgcids_to_labels_dict, ogfaafile,
                                                    filtering_dir, special_case)
        if len(check_if_unique(bgcids_to_labels_dict)) > 0:
            print("\nERROR: filtering did not work for %s\n" %(ogfaafile))
            pprint(bgcids_to_labels_dict)
            exit()
    ### set new filenames
    if oglabel != "":
        oglabel = oglabel.replace(' ', '')
        oglabel = oglabel.replace(',','_')
        oglabel = oglabel.replace('/','_')
        oglabel = oglabel.replace('(', '_')
        oglabel = oglabel.replace(')', '_')
        outfilename = "%s_%s.faa"%(os.path.basename(ogfaafile)[:-4], oglabel)
    else:
        outfilename = "%s.faa" % (os.path.basename(ogfaafile)[:-4])
    if special_case == 'oxyE':
        outfilename = outfilename.replace("OG_3","OG_3a")
    ### get new records with fixed labels
    records = list(SeqIO.parse(ogfaafile, 'fasta'))
    # print(ogfaafile)
    new_records = []
    thisoggeneslist = []
    for rec in records:
        bgcid = rec.id.split(headersep)[0]
        try:
            correct_seq = bgcids_to_labels_dict[bgcid][0]
            if rec.id == correct_seq:
                rec.id = bgcid
                rec.name = bgcid
                rec.description = bgcid
                new_records.append(rec)
                thisoggeneslist.append(correct_seq)
        except:
            if not special_case == 'oxyE':
                print('ERROR: something went wrong with the filtering of %s' %(ogfaafile))
                exit()

    if modules_version_dir !='':
        outmodfilename = os.path.join(modules_version_dir, outfilename)
        if not os.path.exists(outmodfilename) or replace:
            with open(outmodfilename, 'w') as handle:
                SeqIO.write(new_records, handle,'fasta')
    if domains_version_dir != '':
        outdomfilename = os.path.join(domains_version_dir, outfilename.replace("all_",""))
        if not os.path.exists(outdomfilename) or replace:
            with open(outdomfilename, 'w') as handle:
                SeqIO.write(new_records, handle,'fasta')
    #print(ogfaafile, thisoggeneslist, duplicates_num)
    return thisoggeneslist, duplicates_num, warningslist

'''given a directory, lists all files, with a certain suffix'''
def list_files(directory, suffix= "", suppress=True):
    the_files = []
    for file in os.listdir(directory):
        if file.endswith(suffix):
            the_files.append(os.path.join(directory, file))
    if not suppress:
        print("Found %s %s files" %(len(the_files), suffix))
    the_files.sort()
    return the_files

def list_folders(directory):
    folders_list = [os.path.join(directory, o) for o in os.listdir(directory)
                    if os.path.isdir(os.path.join(directory,o))]
    return folders_list

# Main function #
def main(input, output, nrps_genes_table, orthogroups_table,
         modules_analysis_dir, domains_analysis_dir):
    print("\nInput directory of OGs sequences: %s\n"
          "Output directory: %s\n"
          "NRPS genes table: %s\n"
          "Orthogroups table: %s\n"
          "Modules analysis folder: %s\n"
          "Domains analysis folder: %s\n"
          %(input, output, nrps_genes_table, orthogroups_table,
         modules_analysis_dir, domains_analysis_dir))
    create_folder(output)
    modules_version_dir = os.path.join(output, '1_OGs_filtered_with_modules_faas')
    create_folder(modules_version_dir)
    domains_version_dir = os.path.join(output, '2_OGs_filtered_with_domains_faas')
    create_folder(domains_version_dir)
    nrps_genes_list = csv_to_list(nrps_genes_table, 'locus_tag')
    ogs_df = pd.read_csv(orthogroups_table, sep='\t', index_col=0)
    ogs_df['genes_list'] = ogs_df.iloc[:,5:].values.tolist()
    ogs_df['genes_list'] = ogs_df['genes_list'].apply(lambda x: [i for i in x if str(i) != "nan"])
    ogs_df['genes_list'] = ogs_df['genes_list'].apply(lambda x: [i.split("|")[-1] for i in x])
    ogs_df.sort_values(by=['Proportion of Total Gene Clusters with OG'], inplace=True)
    if ogs_df.iloc[-1,1] == 1:
        total_bgcs_num = len(ogs_df.iloc[-1,-1])
        print("Total numnber of BGCs detected: %s" %(total_bgcs_num))
    else:
        print(ogs_df.iloc[-1,1])
        print("\nWarning: no group found present in all BGCs. perect_BGCs_present column will now specify count\n")
        total_bgcs_num = 1
        #print('total', total_bgcs_num)
    #print(ogs_df)
    ogs_df_withoutnrps = ogs_df[ogs_df['genes_list'].map(lambda x: check_that_isnt_nrps(x,nrps_genes_list))]
    ogs_df_withoutnrps['label'] =ogs_df_withoutnrps.loc[:,'as_gene']
    ogs_df_withoutnrps.label.fillna(ogs_df_withoutnrps.Most_fitting_Annotation, inplace=True)
    #print(ogs_df_withoutnrps)
    ### adding manualy oxyEs
    oxyAErow = ogs_df_withoutnrps.xs('OG_3').copy()
    oxyAErow['as_gene'] = 'oxyE'
    oxyAErow['label'] = 'oxyE'
    oxyAErow.name = 'OG_3e'
    #ogs_df_withoutnrps.append(oxyAErow)
    ogs_df_withoutnrps.loc['OG_3e'] = oxyAErow
    #print(ogs_df_withoutnrps)
    #print(ogs_df_withoutnrps.columns)

    #exit()

    ogs_to_labels_dict = ogs_df_withoutnrps['label'].to_dict()
    #print(ogs_df_withoutnrps)
    ### sort out labels for OGs
    for og, label in ogs_to_labels_dict.items():
        if  str(label) == "nan":
            ogs_to_labels_dict[og] = ''
        elif len(label)> 10:
            ogs_to_labels_dict[og] = label[:10]
    #pprint(ogs_to_labels_dict)
    print("\nNote: Keeping only OGs and groups with at least 4 sequences after single-copy filtering.\n")
    filtering_dir = os.path.join(output, '0_multicopies_filtering')
    create_folder(filtering_dir)
    ogs_to_filtered_genes_dict = {}
    ogs_to_duplicate_num_dict = {}
    totalwarninglines = []
    ### adding manualy oxyEs
    #ogs_to_labels_dict['OG_3e'] = "oxyE"
    special_case = ''

    ### go through faa files, filter duplicates, rename sequences and copy to respective folders
    for og, oglabel in ogs_to_labels_dict.items():
        ogfaafile = os.path.join(input, '%s.faa' %(og))
        if og == "OG_3e":
            ogfaafile = os.path.join(input, '%s.faa' % ('OG_3'))
            special_case = 'oxyE'
        thisoggeneslist, duplicates_num, thiswarningslist = filter_copy_file(ogfaafile, oglabel, modules_version_dir,
                                                           domains_version_dir, filtering_dir,
                                                                             special_case=special_case)
        if len(thiswarningslist) > 0:
            totalwarninglines.append(thiswarningslist)
        ogs_to_filtered_genes_dict[og] = thisoggeneslist
        ### fixing oxyA and oxyE
        if og == "OG_3e" or og == "OG_3":
            ogs_to_duplicate_num_dict[og] = 0
        else:
            #print(og, duplicates_num, thisoggeneslist)
            try:
                ogs_to_duplicate_num_dict[og] = duplicates_num / total_bgcs_num
            ### in cases where None, they get filtered out later
            except:
                ogs_to_duplicate_num_dict[og] = duplicates_num
        special_case = ''
        '''
        ### 06.12.2023 added special case of oxyE - needs to be saved as extra file
        if og == "OG_3":
            extraog = og + "e"
            extralabel = "oxyE"
            thisoggeneslist, duplicates_num, thiswarningslist = filter_copy_file(ogfaafile, extralabel,
                                                                                 modules_version_dir,
                                                                                 domains_version_dir, filtering_dir,
                                                                                 special_case='oxyE')
            ogs_to_filtered_genes_dict[extraog] = thisoggeneslist
            try:
                ogs_to_duplicate_num_dict[extraog] = duplicates_num / total_bgcs_num
            ### in cases where None, they get filtered out later
            except:
                ogs_to_duplicate_num_dict[extraog] = duplicates_num
        '''
        #break
    ### testing
    #exit()
    ### now print possible issues
    #pprint(totalwarninglines)
    ### now add module and domain sequences
    moddom_sep="__"
    print("Assuming headerseparator for modules and domains to be %s" % (moddom_sep))
    mods_to_genes_dict = {}
    module_files_list = list_files(os.path.join(modules_analysis_dir, 'modules_faas'))
    for modfile in module_files_list:
        modgroupname = os.path.basename(modfile)[:-4]
        thisoggeneslist, duplicates_num, thiswarningslist = filter_copy_file(modfile, "", modules_version_dir, "",
                                           filtering_dir,headersep=moddom_sep)
        if len(thiswarningslist) > 0:
            totalwarninglines.append(thiswarningslist)
        if type(thisoggeneslist) is list:
            mods_to_genes_dict[modgroupname] = thisoggeneslist
    domain_folders_list = list_folders(domains_analysis_dir)
    doms_to_genes_dict = {}

    for domfolder in domain_folders_list:
        if domfolder.endswith('_faas'):
            if os.path.basename(domfolder) in ["X_faas","Thioesterase_faas"]:
                #print(domfolder)
                dtype = os.path.basename(domfolder).replace("_faas","")
                domslist = [os.path.join(os.path.dirname(domfolder),'all_%s_domains.faa' %(dtype))]
            else:
                domslist = list_files(domfolder, '.faa')
            for domfile in domslist:
                domgroupname = os.path.basename(domfile)[:-4]
                thisoggeneslist, duplicates_num, thiswarningslist = filter_copy_file(domfile, "", "",
                                                                   domains_version_dir,
                                                   filtering_dir, headersep=moddom_sep)
                if len(thiswarningslist) > 0:
                    totalwarninglines.append(thiswarningslist)
                if "X_domains" in domgroupname or "Thioesterase_domains" in domgroupname:
                    domgroupname = domgroupname.replace("all_","")
                if type(thisoggeneslist) is list:
                    doms_to_genes_dict[domgroupname] = thisoggeneslist
    ### now print possible issues
    pprint(totalwarninglines)
    ### make a table with group info
    ogs_df_withoutnrps['label'] = ogs_to_labels_dict.values()
    ogs_df_withoutnrps.rename(columns={"Proportion of Total Gene Clusters with OG": "percent_BGCs_present"},
                              inplace=True)

    #print(ogs_df_withoutnrps)
    #print(ogs_df_withoutnrps.columns)
    #exit()
    ogs_df_final = pd.concat([ogs_df_withoutnrps.iloc[:,1:5], ogs_df_withoutnrps.iloc[:,-2:]], axis=1)
    #print(ogs_df_final)
    ogs_df_final['genes_list'] = ogs_to_filtered_genes_dict.values()
    ogs_df_final['percent_duplications'] = ogs_to_duplicate_num_dict.values()
    ogs_df_final.dropna(subset=['genes_list'], inplace=True)
    ogs_df_final['bgc_count'] = ogs_df_final['genes_list'].apply(lambda x: len(x))
    ### fix for oxyE - added on 07.12.2023
    ogs_df_final.loc['OG_3e', 'percent_BGCs_present'] = ogs_df_final.loc['OG_3e', 'bgc_count'] / total_bgcs_num
    #final_cols = ogs_df_final.columns
    print(ogs_df_final)
    ### make df from modules and domains
    modrows = []
    for og, oggeneslist in mods_to_genes_dict.items():
        #print(og, oggeneslist)
        row_dict = {}
        row_dict['percent_BGCs_present'] = len(oggeneslist) / total_bgcs_num
        row_dict['Most_fitting_Annotation'] = og
        row_dict['as_gene'] = 'coreNRPS'
        row_dict['as_gene_kind'] = 'biosynthetic'
        row_dict['genes_list'] = oggeneslist
        row_dict['label'] = og
        row_dict['bgc_count'] = len(oggeneslist)
        row_dict['percent_duplications'] = 0
        modrows.append(row_dict)
    mod_df = pd.DataFrame.from_dict(modrows, orient='columns')
    mod_df.set_index('label', drop=False, inplace=True)
    print(mod_df)
    #exit()
    domrows = []
    for og, oggeneslist in doms_to_genes_dict.items():
        row_dict = {}
        row_dict['percent_BGCs_present'] = len(oggeneslist) / total_bgcs_num
        row_dict['Most_fitting_Annotation'] = og
        row_dict['as_gene'] = 'coreNRPS'
        row_dict['as_gene_kind'] = 'biosynthetic'
        row_dict['genes_list'] = oggeneslist
        row_dict['label'] = og
        row_dict['bgc_count'] = len(oggeneslist)
        row_dict['percent_duplications'] = 0
        domrows.append(row_dict)
    dom_df = pd.DataFrame.from_dict(domrows, orient='columns')
    dom_df.set_index('label',  drop=False, inplace=True)
    print(dom_df)
    ###full dfs and save
    full_mod_df = pd.concat([ogs_df_final, mod_df])
    full_mod_df.drop(['genes_list'], inplace=True, axis=1)
    out_mod_table = os.path.join(output, '1_OGs_filtered_with_modules_metadata.tsv')
    full_mod_df.to_csv(out_mod_table, sep='\t')
    full_dom_df = pd.concat([ogs_df_final, dom_df])
    full_dom_df.drop(['genes_list'], inplace=True, axis=1)
    out_dom_table = os.path.join(output, '2_OGs_filtered_with_domains_metadata.tsv')
    full_dom_df.to_csv(out_dom_table, sep='\t')

    ### now print possible issues
    print("")
    for warningsl in totalwarninglines:
        for warn in warningsl:
            print(warn)
    #pprint()
    print("\nDone")


# Parsing arguments & Execution #
parser = argparse.ArgumentParser(description='Makes all the filtering and renaming steps necessary before the basic '
                                             'phylogenetics can begin.')
parser.add_argument('-i', '--input', help='Input', required=True)
parser.add_argument('-o', '--output', help='Output', required=True)
parser.add_argument('-nrps', '--nrps_genes_table', help='Table with bgc IDs and their NRPS genes',
                    required=True)
parser.add_argument('-ot', '--orthogroups_table', help='Table with custom annotated orthogroups and '
                                                       'members', required=True)
parser.add_argument('-m', '--modules_analysis_dir', help='Folder of modules analysis', required=True)
parser.add_argument('-d', '--domains_analysis_dir', help='Folder of domains analysis', required=True)
if __name__ == "__main__":
    args = parser.parse_args()
    main(args.input, args.output, args.nrps_genes_table, args.orthogroups_table,
         args.modules_analysis_dir, args.domains_analysis_dir)
```
