## Supplementary Data 5 for "Structural and evolutionary analyses support reclassification of glycopeptide antibiotics as xyclopeptides": run_bakta.py

```
#!/usr/bin/python3.6
# Created by Athina Gavriilidou at 17.07.23 11:12 #
# Imports #
import argparse
import os
from pprint import pprint
import pandas as pd
import re
import fileinput
from Bio import SeqIO
import numpy as np

# Functions #
def create_folder(path):
    if not os.path.exists(path):
        os.mkdir(path)
        print("Created folder: %s" %(path))

'''given a directory, lists all files, with a certain suffix'''
def list_files(directory, suffix= "", suppress=False):
    the_files = []
    for file in os.listdir(directory):
        if file.endswith(suffix):
            the_files.append(os.path.join(directory, file))
    if not suppress:
        print("Found %s %s files" %(len(the_files), suffix))
    the_files.sort()
    return the_files

'''this version includes gene calling'''
def execute_bakta_full(fastafile, outputfolder, bgcid, taxonomy_string, databasepath, fileprefix, ismag=False):
    bakta_path = "/home/athina/anaconda3/envs/bakta/bin/bakta"
    print("Using bakta in path: %s\n" %(bakta_path))
    if not ismag:
        bakta_command = "%s --db %s --output %s --threads 30 --skip-trna --skip-tmrna --prefix %s --locus %s --locus-tag %s %s " \
                        "--keep-contig-headers %s" \
                        %(bakta_path, databasepath, outputfolder, fileprefix, bgcid, bgcid, taxonomy_string, fastafile)
    else:
        bakta_command = "%s --db %s --output %s --threads 30 --skip-trna --skip-tmrna --prefix %s --locus %s --locus-tag %s %s " \
                        "--keep-contig-headers --meta %s" \
                        %(bakta_path, databasepath, outputfolder, fileprefix, bgcid, bgcid, taxonomy_string, fastafile)
    print(bakta_command)
    #'''
    try:
        os.system(bakta_command)
    except:
        print("ERROR: bakta run failed for %s" %(fastafile))
        exit()
    #'''

'''this version reannotates a given faa file'''
def execute_bakta(fastafile, outputfolder, databasepath, fileprefix):
    bakta_path = "/home/athina/anaconda3/envs/bakta/bin/bakta_proteins"
    print("Using bakta_proteins in path: %s\n" %(bakta_path))
    bakta_command = "%s --db %s --output %s --threads 30  --prefix %s -v --force  %s" \
                    %(bakta_path, databasepath, outputfolder, fileprefix, fastafile)
    print(bakta_command)
    #'''
    try:
        os.system(bakta_command)
    except:
        print("ERROR: bakta run failed for %s" %(fastafile))
        exit()
    #'''

def infer_bgcid(filename, metadata_df, bgcidslist):
    ### latest change should allow easy bgcid inference
    if "__" in filename:
        bgcid = filename.split("__")[0]
        #print(bgcid)
        bgc_dict = metadata_df.loc[metadata_df['ID'] == bgcid].to_dict(orient='list')
    ### culp dataset
    elif filename[:2].isnumeric():
        source_df = metadata_df.loc[metadata_df['source'] == 'Culp']
        bgc_dict = source_df.loc[source_df['Database_ID'] == str(int(filename[:2]))].to_dict(orient='list')
        #print(bgc_dict)
        bgcid = bgc_dict['ID'][0]
    ### MIBIG
    elif filename.startswith("BGC"):
        source_df = metadata_df.loc[metadata_df['source'] == 'MIBiG']
        bgc_dict = source_df.loc[source_df['Database_ID'] == filename.split("_")[0]].to_dict(orient='list')
        bgcid = bgc_dict['ID'][0]
        #print(filename, bgcid)
    ### Genbank
    elif filename.startswith("GCA_"):
        source_df = metadata_df.loc[metadata_df['source'] == 'Genbank']
        indices = [m.start() for m in re.finditer('_', filename)]
        #print(filename[:indices[1]])
        bgc_dict = source_df.loc[source_df['Database_ID'].str.contains(filename[:indices[1]])].to_dict(orient='list')
        bgcid = bgc_dict['ID'][0]
    ### Refseq
    elif filename.startswith("GCF_"):
        source_df = metadata_df.loc[metadata_df['source'] == 'RefSeq']
        indices = [m.start() for m in re.finditer('_', filename)]
        # print(filename[:indices[1]])
        bgc_dict = source_df.loc[source_df['Database_ID'].str.contains(filename[:indices[1]])].to_dict(orient='list')
        #if "GCF_000718635" in filename:
        #    print(bgc_dict)
        bgcid = bgc_dict['ID'][0]
    ### MGnify
    elif filename.startswith("MG"):
        source_df = metadata_df.loc[metadata_df['source'] == 'MGnify']
        bgc_dict = source_df.loc[source_df['Database_ID'] == filename.split("_")[0]].to_dict(orient='list')
        bgcid = bgc_dict['ID'][0]
    ### ShrikantOldTelSeq
    elif filename.startswith("QP"):
        source_df = metadata_df.loc[metadata_df['source'] == 'ShrikantOldTelSeq']
        bgc_dict = source_df.loc[source_df['Database_ID'] == filename.split("_")[0]].to_dict(orient='list')
        bgcid = bgc_dict['ID'][0]
    ### Stammsamlung2022
    elif filename.startswith("S00"):
        source_df = metadata_df.loc[metadata_df['source'] == 'Stammsamlung2022']
        bgc_dict = source_df.loc[source_df['Database_ID'] == filename.split("_")[0]].to_dict(orient='list')
        bgcid = bgc_dict['ID'][0]
    ### LocalSeq - hardcoded for ease
    elif filename.startswith("Sva"):
        source_df = metadata_df.loc[metadata_df['source'] == 'LocalSeq']
        bgc_dict = source_df.loc[source_df['Database_ID'] == 'Sva'].to_dict(orient='list')
        if 'region003' in filename:
            bgcid = 'SvN_1'
        elif 'region022' in filename:
            bgcid = 'SvN_2'
    '''
    ### extra check for unknown cases of two clusters in a genome - unneccessary since another metadata table is input
    if bgcid in bgcidslist:
        bgc1 = "%s_1" % (bgcid)
        bgc2 = "%s_2" % (bgcid)
        bgcidslist.append(bgc1)
        bgcidslist.append(bgc2)
        bgcidslist.remove(bgcid)
        bgcid = bgc2
    '''
    if len(bgc_dict['ID']) != 1:
        print(bgcid, filename, bgc_dict)

    #print(bgcid)

    taxonomy_string = bgc_dict['strain'][0]
    if not type(taxonomy_string) is str:
        taxonomy_string = 'Uncultured organism'
    taxonomy_list = taxonomy_string.split(' ')
    if not 'Uncultured' in taxonomy_string:
        genus = taxonomy_list[0]
        species = taxonomy_list[1]
        strain = ''
        if len(taxonomy_list) > 2:
            strain = taxonomy_string[len(genus) + len(species) + 2:].replace(" ", "_")
        if strain != '':
            taxonomy_string = '--genus %s --species %s --strain %s' %(genus, species, strain)
        else:
            taxonomy_string = '--genus %s --species %s' %(genus, species)
    else:
        taxonomy_string = ''
    return bgcid, taxonomy_string, bgcidslist

### search for string in file and replace
def grep_replace(filename, text_to_search, replacement_text):
    with fileinput.FileInput(filename, inplace=True, backup='.bak') as file:
        for line in file:
            print(line.replace(text_to_search, replacement_text), end='')

def replace_def_in_gbk(gbk_filename, text_to_search, replacement_text):
    input_handle = open(gbk_filename, "r")
    newrecords = []
    overwrite = False
    for seq_record in SeqIO.parse(input_handle, "genbank"):
        defin = seq_record.description
        #print(defin)
        #print("Dealing with GenBank record %s" % seq_record.id)
        if text_to_search  in defin:
            #print([defin])
            defin = defin.replace(text_to_search, replacement_text)
            seq_record.description = defin
            overwrite = True
            #print(defin)
        newrecords.append(seq_record)
    input_handle.close()
    ### create a fixed copy
    if overwrite:
        print("Fixing %s" %(gbk_filename))
        with open(gbk_filename, 'w') as output_handle:
            SeqIO.write(newrecords, output_handle, 'genbank')

'''for copying files. Make everything=True and then everything from the source will be copied to the destination'''
def execute_cp(source_fileordir, destinationfileordir, everything=False):
    if everything:
        cp_command = 'cp %s/* %s/.' %(source_fileordir,destinationfileordir)
    else:
        cp_command = 'cp %s %s' %(source_fileordir,destinationfileordir)
    try:
        os.system(cp_command)
    except:
        print("Warning. There was a problem with copying %s to %s" %(source_fileordir, destinationfileordir))

def get_cd_locustags_locations(the_gbk_file):
    ### adding notfound case for later
    locustags_dict = {'':[]}
    ### getting info from genbank file
    records = list(SeqIO.parse(the_gbk_file, 'genbank'))
    cdscounter = 0
    for rec in records:
        this_seq = rec.id  ### takes it from Version???
        for seq_feature in rec.features:
            ### check cluster limits
            if seq_feature.type == "CDS":
                if len(seq_feature.qualifiers['translation'][0]) < 10:
                    print("Warning: skipping short sequence at %s" % (seq_feature.location))
                    continue
                cdscounter+=1
                try:
                    thislocustag = '|'.join(seq_feature.qualifiers['locus_tag'])
                except:
                    thislocustag = ''
                try:
                    thisproduct = '|'.join(seq_feature.qualifiers['product'])
                except:
                    thisproduct = ''
                try:
                    thisproteinid = '|'.join(seq_feature.qualifiers['protein_id'])
                except:
                    thisproteinid = ''
                try:
                    thisgene = '|'.join(seq_feature.qualifiers['gene'])
                except:
                    thisgene = ''
                #print(dir(seq_feature.location.start))
                thisstart = int(seq_feature.location.start)
                thisend = int(seq_feature.location.end)
                #print(seq_feature)
                ### special case of StavMA_1 - should be solved by bakta update?
                #if thislocustag == 'VK71_RS52500':
                #    continue
                #print(thislocustag, thisstart, thisend, thisproduct, thisproteinid)
                if thislocustag not in locustags_dict.keys():
                    locustags_dict[thislocustag] = [thisstart, thisend, thisproduct, thisproteinid, thisgene, thislocustag]
                ### to cover the cases where there are no locus tags
                else:
                    locustags_dict[cdscounter] = [thisstart, thisend, thisproduct, thisproteinid, thisgene, thislocustag]

    #print("Went through %s CDS" %(cdscounter))
    ### turn into df for easier parsing
    locustags_df = pd.DataFrame.from_dict(locustags_dict,orient='index',
                                          columns=['start','end','product','protein_id','gene','locus_tag'])
    locustags_df.drop(index='',inplace=True)
    return locustags_df

def add_original_tags(gbkfile, gene_annotations_tsv, reannotatedgbk, overwrite):
    bgcid = os.path.basename(reannotatedgbk).replace('.gbk','')
    gene_annotations_df = pd.read_csv(gene_annotations_tsv, sep='\t', skiprows=2)
    gene_annotations_df.replace(np.nan, '', regex=True, inplace=True)
    gene_annotations_df.index += 1
    bakta_cols = gene_annotations_df.columns
    col_dict = {}
    for col in bakta_cols:
        col_dict[col] = '%s_bakta' %(col)
    gene_annotations_df.rename(columns=col_dict, inplace=True)
    #gene_annotations_df[['filename', 'oneidentifier','location']] = gene_annotations_df['ID'].str.split('|', expand=True)
    original_locustags_df = get_cd_locustags_locations(gbkfile)
    #print(gene_annotations_df)
    #print(original_locustags_df)
    ### if the number of rows is the same, the order of genes should be the same too and the tables can be merged
    if len(gene_annotations_df) == len(original_locustags_df):
        original_locustags_df.reset_index(drop=True, inplace=True)
        gene_annotations_df.reset_index(drop=True, inplace=True)
        full_gene_annotations_df =pd.concat([original_locustags_df, gene_annotations_df], axis=1)
        if len(full_gene_annotations_df) != len(gene_annotations_df):
            print("\nERROR: something went wrong with the concatenation. Aborting...")
            print(full_gene_annotations_df)
            exit()
        #print(full_gene_annotations_df)
    else:
        print("ERROR: Number of genes in bakta table and provided table does not match! Aborting...")
        print(gbkfile)
        print(gene_annotations_tsv)
        exit()
    full_gene_annotations_df['BGC_ID'] = bgcid
    #full_gene_annotations_df = full_gene_annotations_df.columns.tolist()[-1:] + full_gene_annotations_df.columns.tolist()[1:]
    fixed_gbk_records_list = []
    #print("Getting info from: %s" %(original_gbkfile))
    ### getting info from genbank file
    try:
        records = list(SeqIO.parse(gbkfile, 'genbank'))
    except:
        print("ERROR: Could not extract records from %s" % (gbkfile))
        exit()
    cds_counter = 0
    originals_counter = 0
    new_locus_tags_list = []
    for rec in records:
        sequence_id = rec.id
        #print(sequence_id)
        for seq_feature in rec.features:
            ### add original info to CDS
            if seq_feature.type == "CDS":
                if len(seq_feature.qualifiers['translation'][0]) < 10:
                    #print("Warning: skipping short sequence at %s" % (seq_feature.location))
                    continue
                ### special case for StavMA_1
                #if 'locus_tag' in seq_feature.qualifiers and seq_feature.qualifiers['locus_tag'][0]  == 'VK71_RS52500':
                #    continue
                cds_counter+=1
                thisstart = int(seq_feature.location.start)
                thisend = int(seq_feature.location.end)
                #print(seq_feature)
                #print(thisstart, thisend)
                exactlocusrows = full_gene_annotations_df[(full_gene_annotations_df['start'] == thisstart) &
                                                       (full_gene_annotations_df['end'] == thisend)]

                #print(exactlocusrows)
                '''
                extratags=[]
                if len(exactlocusrows) == 0:
                    locusrowsleft = original_locustags_df[(original_locustags_df['start'] < thisstart) &
                                                          (original_locustags_df['end'] <= thisend) &
                                                          (thisstart < original_locustags_df['end'])]
                    if len(locusrowsleft) > 0:
                        extratags.append('partialleft')
                    locusrowsmid = original_locustags_df[(original_locustags_df['start'] >= thisstart) &
                                                            (original_locustags_df['end'] <= thisend)]
                    if len(locusrowsmid) > 0:
                        extratags.append('partialmid')
                    locusrowsright = original_locustags_df[(original_locustags_df['start'] >= thisstart) &
                                                           (original_locustags_df['end'] > thisend) &
                                                           (thisend > original_locustags_df['start'])]
                    if len(locusrowsright) > 0:
                        extratags.append('partialright')
                    if len(locusrowsmid) == 0:
                        locusrowsmid = original_locustags_df[(original_locustags_df['start'] > thisstart) &
                                                           (original_locustags_df['end'] < thisend) ]
                        if len(locusrowsmid) > 0:
                            extratags.append('partialboth')
                    exactlocusrows = pd.concat([locusrowsleft, locusrowsmid, locusrowsright])
                '''
                #print(exactlocusrows)
                #print(extratags)
                #print(exactlocusrows.columns)
                #exit()
                collist = exactlocusrows.columns.tolist()
                for col in collist[2:]:
                    if col == 'locus_tag' and len(exactlocusrows[col].values.tolist()[0]) != 0:
                        qualifier = 'original_locus_tag'
                        #print(exactlocusrows[col].values.tolist())
                        seq_feature.qualifiers[qualifier] = exactlocusrows[col].values.tolist()
                    if col not in seq_feature.qualifiers and len(str(exactlocusrows[col].values.tolist()[0])) != 0:
                        if not 'bakta' in col:
                            seq_feature.qualifiers[col] = exactlocusrows[col].values.tolist()
                        else:
                            qualifier = col.replace('_bakta','')
                            #for el in exactlocusrows[col].values.tolist():
                            seq_feature.qualifiers[qualifier] = exactlocusrows[col].values.tolist()
                    elif col in seq_feature.qualifiers and seq_feature.qualifiers[col] != exactlocusrows[col].values.tolist():
                        print("Warning: info does not match")
                        print(exactlocusrows, col)
                        exit()
                ### add new locus tag
                newlocus_tag = '%s__%s' %(bgcid, cds_counter)
                seq_feature.qualifiers['locus_tag'] = [newlocus_tag]
                new_locus_tags_list.append(newlocus_tag)

                '''
                seq_feature.qualifiers['original_locus_tag'] = exactlocusrows["locus_tag"].values.tolist()
                seq_feature.qualifiers['original_product'] = exactlocusrows["product"].values.tolist()
                seq_feature.qualifiers['original_protein_id'] = exactlocusrows["protein_id"].values.tolist()
                seq_feature.qualifiers['original_gene'] = exactlocusrows["gene"].values.tolist()
                '''
                #originals_counter +=1
                #if len(extratags) > 0:
                #    seq_feature.qualifiers['original_notes'] = extratags
                #break
        # break
        fixed_gbk_records_list.append(rec)
    #if cds_counter != originals_counter:
    #    print("\nERROR: Problem with reannotation of %s\n" %(gbkfile))
    #    exit()
    #else:
        #print("Went through %s CDS" %(cds_counter))
    ### save fixed records
    if not os.path.exists(reannotatedgbk) or overwrite:
        SeqIO.write(fixed_gbk_records_list, reannotatedgbk, 'genbank')
    #print("Created file: %s\n" % (reannotatedgbk))
    full_gene_annotations_df.rename(columns={'locus_tag':'original_locus_tag'}, inplace=True)
    #print(full_gene_annotations_df)
    full_gene_annotations_df.insert(0,'locus_tag',new_locus_tags_list,True)
    return full_gene_annotations_df


# Main function #
def main(input, output,metadata, overwrite=False):
    print("\nInput folder: %s"
          "\nOutput folder: %s"
          "\nMetadata table %s"
          "\nOverwrite files: %s\n"
          %(input,output,metadata, overwrite))
    create_folder(output)
    databasepath = '/media/hdd_2tb/bakta_db/db'
    print("Using bakta database under: %s" %(databasepath))
    #the_input_files = list_files(input, '.fna')
    the_input_files = list_files(input, '.faa')
    metadata_df = pd.read_csv(metadata)
    #print(metadata_df)
    bgcidslist = []
    gene_dfs_list = []
    for input_file in the_input_files:
        filename = os.path.basename(input_file)[:-4]
        ### for testing
        #if not filename.startswith('GCF'):
        #    continue
        ### special case for preprint dataset for testing - 31.08.23
        #if "preprint" in input_file:
        #    bgcid = filename.rstrip('.gbk')
        #    taxonomy_string = ""
        #else:
        input_gbkfile = input_file[:-4] + '.gbk'
        bgcid, taxonomy_string, bgcidslist = infer_bgcid(filename, metadata_df, bgcidslist)
        bgcidslist.append(bgcid)
        ### for testing
        #if not bgcid == 'bal':
        #    continue
        outputfolder = os.path.join(output, bgcid)
        #print(taxonomy_string)
        if not os.path.exists(os.path.join(outputfolder, '%s.tsv' %(bgcid))) or overwrite:
            execute_bakta(input_file, outputfolder, databasepath, bgcid)
            '''
            if not bgcid.startswith("U"):
                execute_bakta(input_file, outputfolder, bgcid, taxonomy_string, databasepath, bgcid, ismag=False)
            else:
                execute_bakta(input_file, outputfolder, bgcid, taxonomy_string, databasepath, bgcid, ismag=True)
            '''
        ### for testing
        #continue
        if len(list_files(outputfolder, suppress=True)) < 5:
            print("Warning: Bakta folder detected but contains fewer files than expected:\n%s" %(outputfolder))
        ### special case for preprint dataset for testing
        if "preprint" in input_file:
            continue

        ### fix definition in files
        gbkfile = os.path.join(outputfolder, '%s.gbff' %(bgcid))
        emblfile = os.path.join(outputfolder, '%s.gbff' %(bgcid))
        #replace_def_in_gbk(gbkfile, "whole genome shotgun sequence",
        #                   "BGC limits defined by antiSMASH 7.0.0")
        #replace_def_in_gbk(emblfile, "whole genome shotgun sequence",
        #                   "BGC limits defined by antiSMASH 7.0.0")
        ### add original annotations to gbff file - take info from antismash generated gbk file, since quick antismash
        ### was run with downloaded gbk files where possible
        reannotatedgbk = os.path.join(outputfolder, '%s.gbk' %(bgcid))
        gene_annotations_tsv = os.path.join(outputfolder,'%s.tsv' %(bgcid))
        #if not os.path.exists(reannotatedgbk) or overwrite:
        full_gene_annotations_df = add_original_tags(input_gbkfile, gene_annotations_tsv, reannotatedgbk, overwrite)
        gene_dfs_list.append(full_gene_annotations_df)
        ### for testing
        #break
    complete_genes_df = pd.concat(gene_dfs_list)
    print(complete_genes_df)
    print("\nNote: Altered descriptions in gbff and embl files in bakta results")
    print("\nDone")

#'''
# Parsing arguments & Execution #
parser = argparse.ArgumentParser(description='Takes a file or folder of fasta files and runs most comprehensive '
                                             'bakta annotation. Made for BGCs, so in a post-processing step, the '
                                             'created .gbff files are edited to avoid mislabeling them as genomes.')
parser.add_argument('-i', '--input', help='Input folder with fasta files', required=True)
parser.add_argument('-o', '--output', help='Output folder for bakta results', required=True)
parser.add_argument('-m', '--metadata', help='Metadata table including ID and strain', required=True)
parser.add_argument('-r', '--overwrite', help='Optional: overwrite existing results (folders). Default: False',
                    required=False, default=False)
if __name__ == "__main__":
    args = parser.parse_args()
    main(args.input, args.output, args.metadata, args.overwrite)
#'''
'''
### for debugging
inputdir = '/media/hdd_2tb/GPA_reclassification_clean/A_dataset_creation/2_putative_GPAs_analyses/1_regions_with_Xdom'
outputdir = '/media/hdd_2tb/GPA_reclassification_clean/A_dataset_creation/2_putative_GPAs_analyses/2_regions_with_Xdom_bakta_results'
metadata = '/media/hdd_2tb/GPA_reclassification_clean/A_dataset_creation/2_putative_GPAs_analyses/summary_hits_clusters_metadata.csv'
main(inputdir, outputdir, metadata)
'''
```
