## Supplementary Data 5 for "Structural and evolutionary analyses support reclassification of glycopeptide antibiotics as xyclopeptides": run_basic_phylogeny.py

```
#!/usr/bin/python3.6
# Created by Athina Gavriilidou at 18.09.23 17:16 #
# Imports #
import argparse
import csv
import os
from collections import OrderedDict
import sys
import time

# Functions #
def create_folder(path):
    if not os.path.exists(path):
        os.mkdir(path)
        print("Created folder: %s" %(path))

'''given a directory, lists all files, with a certain suffix'''
def list_files(directory, suffix= "", suppress=True):
    the_files = []
    for file in os.listdir(directory):
        if file.endswith(suffix):
            the_files.append(os.path.join(directory, file))
    if not suppress:
        print("Found %s %s files" %(len(the_files), suffix))
    the_files.sort()
    return the_files

def execute_mafft(faafile, outmsafile):
    mafftpath = '/home/athina/anaconda3/envs/mafft/bin/mafft'
    print("Assuming path to mafft: %s" % (mafftpath))
    mafft_command = "%s --auto %s > %s" %(mafftpath, faafile, outmsafile)
    print(mafft_command)
    try:
        os.system(mafft_command)
    except:
        print("\nERROR: could not run mafft. Are you sure it is installed and in the path?\n")
        exit()

def execute_trimal(mfafile, outtrimalfile):
    trimalpath = '/home/athina/anaconda3/envs/trimal/bin/trimal'
    print("Assuming path to trimal: %s" %(trimalpath))
    trimal_command = "%s -in %s -out %s -automated1" % (trimalpath, mfafile, outtrimalfile)
    print(trimal_command)
    try:
        os.system(trimal_command)
    except:
        print("\nERROR: could not run trimal. Are you sure it is installed and in the path?\n")
        exit()

def grep_file(thefile, thestring):
    with open(thefile, 'r') as f:
        for line in f.readlines():
            if thestring in line:
                return line
    return False


def execute_iqtree_modeltesting(mfafile, outprefix):
    iqtreepath = '/home/athina/anaconda3/envs/iqtree2/bin/iqtree2'
    print("Assuming path to iqtree: %s" % (iqtreepath))
    iqtree_command = (("%s -s %s -m MF -pre %s -redo --msub nuclear -nt AUTO "
                      "--mset Dayhoff,DCMut,JTT,JTTDCMut,LG,Poisson,PMB,Q.pfam,VT,WAG "
                       "--madd C10,C20,C30,C40,C50,C60,EX2,EX3,EHO,UL2,UL3,EX_EHO,LG4M,LG4X,CF4")
                      %(iqtreepath, mfafile, outprefix))
    print(iqtree_command)
    try:
        os.system(iqtree_command)
    except:
        print("\nERROR: could not run iqtree. Are you sure it is installed and in the path?\n")
        exit()

def execute_iqtree(mfafile, outprefix, thismodel):
    iqtreepath = '/home/athina/anaconda3/envs/iqtree2/bin/iqtree2'
    print("Assuming path to iqtree: %s" % (iqtreepath))
    iqtree_command = "%s -nt AUTO -s %s -m %s -pre %s -redo -bb 1000" % (iqtreepath, mfafile, thismodel, outprefix)
    print(iqtree_command)
    try:
        os.system(iqtree_command)
    except:
        print("\nERROR: could not run iqtree. Are you sure it is installed and in the path?\n")
        exit()


def execute_iqtree_keepident(mfafile, outprefix, thismodel):
    iqtreepath = '/home/athina/anaconda3/envs/iqtree2/bin/iqtree2'
    #print("Assuming path to iqtree: %s" % (iqtreepath))
    print("NOTE: Trying again to bootstrap by including identical sequences")
    iqtree_command = "%s -nt AUTO -s %s -m %s -pre %s -redo -bb 1000 -keep-ident" % (
    iqtreepath, mfafile, thismodel, outprefix)
    print(iqtree_command)
    try:
        os.system(iqtree_command)
    except:
        print("\nERROR: could not run iqtree. Are you sure it is installed and in the path?\n")
        exit()

def dict_to_csv(thedict, outcsv, thecols):
    with open(outcsv, 'w') as outfile:
        thewriter = csv.writer(outfile)
        thewriter.writerow(thecols)
        for key, value in thedict.items():
            therow = [key, value]
            thewriter.writerow(therow)
    print("Created file: %s" %(outcsv))

### checks if file was created in the last minute
def check_if_file_is_new(filepath):
    if time.time() - os.stat(filepath).st_mtime < 60:
        return True
    else:
        return False


# Main function #
def main(input, output, replace=False):
    print("\nInput directory of fasta files: %s\n"
          "Output folder: %s\n"
          %(input, output))
    create_folder(os.path.dirname(output))
    create_folder(output)
    mafft_output = os.path.join(output, '1_mafft_MSAs')
    create_folder(mafft_output)
    faa_files_list = list_files(input)
    ## create mafft MSAs
    print("\nStarting mafft MSAs")
    mafftmsas_list = []
    for faafile in faa_files_list:
        outmsafile = os.path.join(mafft_output, os.path.basename(faafile).replace('.faa','_mafftMSA.faa'))
        if not os.path.exists(outmsafile) or replace:
            execute_mafft(faafile, outmsafile)
        mafftmsas_list.append(outmsafile)
    ### create trimmed MSAs
    trimal_output = os.path.join(output, '2_trimal_MSAs')
    create_folder(trimal_output)
    trimalmsas_list = []
    print("\nStarting trimAl of mafft MSAs")
    for mfafile in mafftmsas_list:
        outtrimalfile = os.path.join(trimal_output, os.path.basename(mfafile).replace('_mafftMSA.faa',
                                                                                     '_trimalMSA.faa'))
        if not os.path.exists(outtrimalfile) or replace:
            execute_trimal(mfafile, outtrimalfile)
        trimalmsas_list.append(outtrimalfile)
    ### do model testing
    modelT_output = os.path.join(output, '3_modeltesting')
    create_folder(modelT_output)
    groupid_to_model = OrderedDict()
    model_testing_summary_table = os.path.join(output, 'modeltesting_summary.csv')
    print("\nStarting iqtree modeltesting of mafft MSAs\n")
    for mfafile in mafftmsas_list:
        groupid = os.path.basename(mfafile.replace("_mafftMSA.faa", ''))
        prefix = os.path.join(modelT_output, groupid)
        outmodelTlog = '%s.log' %(prefix)
        if not os.path.exists(outmodelTlog) or replace:
            execute_iqtree_modeltesting(mfafile, prefix)
        ### find model
        try:
            modelline = grep_file(outmodelTlog, 'Best-fit model:').rstrip('\n')
        except:
            print("ERROR: are you sure that model-testing was completed?\nSee file: %s" %(outmodelTlog))
            exit()
        print("%s %s" %(groupid, modelline))
        modelstring = modelline.replace('Best-fit model: ', '')
        modelstring = modelstring.split(" chosen according to")[0]
        #print(modelstring)
        groupid_to_model[groupid] = modelstring
        #break
    dict_to_csv(groupid_to_model, model_testing_summary_table, ['groupid','best_fit_model'])
    #exit()
    ### do ML of untrimmed with correct model
    trees_untrimmed_dir = os.path.join(output, '4_mafft_iqtree_ML_trees')
    create_folder(trees_untrimmed_dir)
    print("\nStarting iqtree ML phylogenies of mafft MSAs\n")
    for mfafile in mafftmsas_list:
        groupid = os.path.basename(mfafile.replace("_mafftMSA.faa", ''))
        thismodel = groupid_to_model[groupid]
        prefix = os.path.join(trees_untrimmed_dir, '%s_mafft_iqtree' %(groupid))
        outtree = '%s.treefile' %(prefix)
        if not os.path.exists(outtree) or replace:
            execute_iqtree(mfafile, prefix, thismodel)
        ## added second attempt at making bootstrapped tree for the genes with many identical sequences
        if not os.path.exists(outtree) or (replace and not check_if_file_is_new(outtree)):
            execute_iqtree_keepident(mfafile, prefix, thismodel)
    ### do ML of untrimmed with correct model
    trees_trimmed_dir = os.path.join(output, '5_trimal_iqtree_ML_trees')
    create_folder(trees_trimmed_dir)
    print("\nStarting iqtree ML phylogenies of trimmed MSAs\n")
    for mfafile in trimalmsas_list:
        groupid = os.path.basename(mfafile.replace("_trimalMSA.faa", ''))
        thismodel = groupid_to_model[groupid]
        prefix = os.path.join(trees_trimmed_dir, '%s_trimal_iqtree' % (groupid))
        outtree = '%s.treefile' % (prefix)
        if not os.path.exists(outtree) or replace:
            execute_iqtree(mfafile, prefix, thismodel)

    print("IMPORTANT NOTE: JTT model of amino acid substitution and a gamma-distributed rate variation with four categories")
    print("\nDone")


# Parsing arguments & Execution #
parser = argparse.ArgumentParser(description='Run basic phylogenetics for the input directory of fasta files.')
parser.add_argument('-i', '--input', help='Input folder with fasta files', required=True)
parser.add_argument('-o', '--output', help='Output folder', required=True)
if __name__ == "__main__":
    args = parser.parse_args()
    main(args.input, args.output)
```
