## Supplementary Data 5 for "Structural and evolutionary analyses support reclassification of glycopeptide antibiotics as xyclopeptides": run_bigscape.py

```
#!/usr/bin/python3.6
# Created by Athina Gavriilidou at 19.07.23 11:54 #
# Imports #
import argparse
import os

# Functions #
'''for copying files. Make everything=True and then everything from the source will be copied to the destination'''
def execute_cp(source_fileordir, destinationfileordir, everything=False):
    if everything:
        cp_command = 'cp %s/* %s/.' %(source_fileordir,destinationfileordir)
    else:
        cp_command = 'cp \'%s\' \'%s\'' %(source_fileordir,destinationfileordir)
    try:
        os.system(cp_command)
    except:
        print("Warning. There was a problem with copying %s to %s" %(source_fileordir, destinationfileordir))


def execute_bigscape(inputfolder, outputfolder, thresholdstring):
    bigscape_path = "/home/athina/Documents/programs/BiG-SCAPE-master/bigscape_GPAmode.py"
    truebigscape_path = "/home/athina/Documents/programs/BiG-SCAPE-master/bigscape.py"
    print("Using BiG-SCAPE in path: %s" %(bigscape_path))
    print("Modified to use NRPS weights also for mix\n")
    bigscape_command = "python %s -i %s -v --mix --include_singletons -o %s --no_classify --cutoffs %s " \
                        "--include_gbk_str gbk --mode glocal --cores 30" \
                        %(bigscape_path, inputfolder, outputfolder, thresholdstring)
    print(bigscape_command)
    try:
        os.system(bigscape_command)
    except:
        print("ERROR: Bigscape run failed for %s" %(inputfolder))
        exit()

def execute_true_bigscape(inputfolder, outputfolder, thresholdstring):
    bigscape_path = "/home/athina/Documents/programs/BiG-SCAPE-master/bigscape.py"
    print("Using BiG-SCAPE in path: %s" %(bigscape_path))
    bigscape_command = "python %s -i %s -v --include_singletons -o %s --cutoffs %s " \
                        "--include_gbk_str gbk --mode glocal --cores 30" \
                        %(bigscape_path, inputfolder, outputfolder, thresholdstring)
    print(bigscape_command)
    try:
        os.system(bigscape_command)
    except:
        print("ERROR: Bigscape run failed for %s" %(inputfolder))
        exit()

def create_folder(path):
    if not os.path.exists(path):
        os.mkdir(path)
        print("Created folder: %s" %(path))

'''given a directory, lists all files, with a certain suffix'''
def list_files(directory, suffix= "", suppress=False):
    the_files = []
    for file in os.listdir(directory):
        if file.endswith(suffix):
            the_files.append(os.path.join(directory, file))
    if not suppress:
        print("Found %s %s files" %(len(the_files), suffix))
    the_files.sort()
    return the_files

def tidy_antismash_gbks(input, mibig_only_annotated_as7_clusters, all_annotated_as7_clusters):
    folder_file_list = []
    folders_list = [os.path.join(input, o) for o
                    in os.listdir(input)
                    if os.path.isdir(os.path.join(input,o))]
    mibig_extra_ids = 'A40,A47,A50,GP6,UK6'.split(',')
    #print("Not tidying up mibig files anymore")
    for folder in folders_list:
        id = os.path.basename(folder)
        if 'all_concatenated' == id:
            continue
        thefiles = list_files(folder, suffix='gbk', suppress=True)
        for thefile in thefiles:
            if 'region' in thefile:
                break
        newfile = os.path.join(all_annotated_as7_clusters, '%s.gbk' %(id))
        execute_cp(thefile, newfile)

        ### added on 24.11.23 - take also json files
        jsonoriginal = '%s/%s.json' %(folder, id)
        newjson = '%s.json' %(newfile[:-4])
        execute_cp(jsonoriginal, newjson)

        #'''
        ### for mibig files only
        if id[0].islower() or id in mibig_extra_ids:
            newfile = os.path.join(mibig_only_annotated_as7_clusters, '%s.gbk' %(id))
            execute_cp(thefile, newfile)
        #'''


# Main function #
def main(input, output, threshold):
    threshold_string = threshold.replace(',', ' ')
    print("\nInput directory: %s\n"
          "Output directory: %s\n"
          "Threshold(s): %s\n"
          %(input, output, threshold_string))
    if len(threshold_string.split(' ')) > 1:
        print("Running in threshold checking mode")
        print("==================================")
        ### first tidy up the antismash results
        mibig_only_annotated_as7_clusters = os.path.join(os.path.dirname(input), '4_mibig_annotated_as7_clusters')
        all_annotated_as7_clusters = os.path.join(os.path.dirname(input), '5_all_candidate_as7_clusters')
        create_folder(mibig_only_annotated_as7_clusters)
        create_folder(all_annotated_as7_clusters)
        tidy_antismash_gbks(input, mibig_only_annotated_as7_clusters, all_annotated_as7_clusters)
        mibig_count = len(list_files(mibig_only_annotated_as7_clusters, suffix='.gbk'))
        total_count = len(list_files(all_annotated_as7_clusters,suffix='.gbk'))
        ### special case of preprint for testing
        if "preprint" in input:
            exit()
        print("\nFound %s GPA clusters in total.\nCollected %s MIBiG clusters" %(total_count,mibig_count))
        if os.path.exists(os.path.join(output, 'network_files')):
            print("Threshold testing analysis already conducted. Aborting...")
            exit()
        if mibig_count == 22:
            execute_bigscape(mibig_only_annotated_as7_clusters, output, threshold_string)
    elif len(threshold_string.split(' ')) == 1:
        print("Running in direct mode")
        print("======================")
        execute_bigscape(input, output, threshold_string)
        #execute_true_bigscape(input, output, threshold_string)

    print("\nDone")


# Parsing arguments & Execution #
parser = argparse.ArgumentParser(description='Run Bigscape (v1.1.5) for a given dataset with given threshold(s). '
                                             '\n Threshold checking mode: Giving multiple thresholds will cause the '
                                             'script to try to separate gbk'
                                             ' files into MIBiG and full files, and to run only the MIBiG in multiple'
                                             ' thesholds. '
                                             '\n Direct mode: Giving one threshold will cause the script to try to run '
                                             'bigscape with the files in the given input directly.')
parser.add_argument('-i', '--input', help='Input directory of gbk files or antismash result folders', required=True)
parser.add_argument('-o', '--output', help='Output directory of bigscape results', required=True)
parser.add_argument('-t', '--threshold', help='Threshold(s) for Bigscape. E.g. 0.1 or 0.1,0.2,0.3. '
                                              'Default is bigscape\'s default: 0.3',
                    required=False, default='0.3')
if __name__ == "__main__":
    args = parser.parse_args()
    main(args.input, args.output, args.threshold)
```
