## Supplementary Data 5 for "Structural and evolutionary analyses support reclassification of glycopeptide antibiotics as xyclopeptides": run_clinker_on_clustergroups.py

```
#!/usr/bin/python3.6
# Created by Athina Gavriilidou at 02.03.22 17:32 #
# Imports #
import argparse
import pandas as pd
import os
from datetime import date
import csv
from pprint import pprint
import json
import numpy as np
from collections import OrderedDict

# Functions #
def create_folder(path):
    if not os.path.exists(path):
        os.mkdir(path)

'''given a directory, lists all files in it with the given suffix'''
def list_files(directory, suffix):
    files = [os.path.join(directory, o) for o in os.listdir(directory)
               if os.path.isfile(os.path.join(directory, o)) and o.endswith(suffix)]
    print("Found %s %s files" % (len(files), suffix))
    files.sort()
    return files

def list_to_string(thelist, spacer):
    thestring = ""
    for el in thelist:
        thestring = thestring + el + spacer
    thestring = thestring[:-len(spacer)]
    return thestring

def execute_clinker(clusterfiles, prefile):
    jsonfile = "%s.json" %(prefile)
    htmlfile = "%s.html" %(prefile)
    clusterfiles_string = list_to_string(clusterfiles, ' ')
    clinkerpath = '/home/athina/anaconda3/envs/clinker/bin/clinker'
    print("Using clinker found under: %s" %(clinkerpath))
    theactualcommand = '%s --session %s --plot %s -ufo %s --jobs 30' \
                       %(clinkerpath, jsonfile, htmlfile, clusterfiles_string)
    print(theactualcommand)
    os.system(theactualcommand)

def list_folders(directory):
    folders_list = [os.path.join(directory, o) for o in os.listdir(directory)
                    if os.path.isdir(os.path.join(directory,o))]
    folders_list.sort()
    return folders_list

def detect_gcftable(bigscape_result_folder):
    thepath = os.path.join(bigscape_result_folder, 'network_files')
    therunfolder = list_folders(thepath)[-1]
    thegcftable = os.path.join(therunfolder, 'mix/mix_clustering_c0.20.tsv')
    if (os.path.exists(thegcftable) and os.path.isfile(thegcftable)):
        return thegcftable
    else:
        print("ERROR: could not find GCF assigment table. Check your parameters.")
        exit()

def csv_to_dict(csvfile, key="", value="", sep=','):
    thedict =OrderedDict()
    with open(csvfile, 'r') as infile:
        reader = csv.reader(infile, delimiter=sep)
        for headerline in reader:
            key_id = headerline.index(key)
            value_id = headerline.index(value)
            break
        for line in reader:
            if line[key_id] not in thedict.keys():
                thedict[line[key_id]] = [line[value_id]]
            else:
                thelist = thedict[line[key_id]]
                thelist.append(line[value_id])
                thedict[line[key_id]] = thelist
    return thedict

def detect_element(thelist, thesubstring):
    for el in thelist:
        if thesubstring in el:
            return el


### read json and create dictionary group : associated genes
def read_json(j_file):
    group_gene_dict = {}
    with open(j_file) as data:
        d = json.load(data)
        for i in d['groups']:
            if "label" in i.keys() and "genes" in i.keys():
                group_gene_dict.update({i["label"] : i["genes"]})
    return group_gene_dict

"""associate gene product to gene product which can then be used as label later
    go through json file and check whether current gene-uid extracted earlier (either from target or query entry (some are only in one)) is found
    if true, extract gene information: directory with key: original group, value: tuple of gene uid and "product" (alternatives are: gene_functions, locus_tag)
    """
def find_product(j_file):
    group_gene_dict = read_json(j_file)
    already_visited = []
    product_dict = {}
    with open(j_file) as data:
        d = json.load(data)
        for key in group_gene_dict:
            for gene in group_gene_dict[key]:
                for id, value in d['links'].items():
                    if 'query' in value.keys():
                        uid_query = value['query']['uid']
                        if uid_query == gene and uid_query not in already_visited:
                            already_visited.append(gene)
                            try:
                                fill_dict(product_dict, key, [gene, value["query"]["names"]["product"]])
                            ### covering case of error from 08.11.2023
                            except:
                                fill_dict(product_dict, key, [gene, value["target"]["names"]["Product"]])
                    if 'target' in value.keys():
                        uid_target = value['target']['uid']
                        if uid_target == gene and gene not in already_visited:
                            already_visited.append(gene)
                            try:
                                fill_dict(product_dict, key, [gene, value["target"]["names"]["product"]])
                            ### covering case of error from 08.11.2023
                            except:
                                fill_dict(product_dict, key, [gene, value["target"]["names"]["Product"]])
    return product_dict

def create_naming_dict(j_file):
    product_dict = find_product(j_file)
    naming_dict = {}
    for group in product_dict:
        product_list = []
        for gene, product in product_dict[group]:
            if product not in product_list:
                product_list.append(product)
        naming_dict.update({group : product_list})
    return naming_dict

### read html file, find inital group names and replace them in second html (so that it can be done multiple times)
def name_html(j_file, html_file, output):
    naming_dict = create_naming_dict(j_file)
    new_file = open(output, "w") # new file to rename groups
    used_keys = [] # necessary, if not used key "group 1" is also in "group 10" and then anything higher than 9 wont be called
    with open(html_file, 'r') as f:
        for line in f:
            for key in reversed(sorted(naming_dict)): # check if one of the group names are in the line (sorted and reversed as otherwise "Group 1" would be in "Group 11" and fuck it up
                if key in line:
                    used_keys.append(key)
                    line = line.replace(key, str(naming_dict[key]).replace("[", "").replace("]", "").replace("'", "")) # replace the label by wanted label (so far just all related genes)
            new_file.write(line)
    new_file.close()

### key: original group, value: list of tuples of gene with product,  I have to check if I have this connection
# already, if yes append to list, if not initialize a list.
def fill_dict(dictionary,key,value):
    if key in dictionary.keys():
        dictionary.get(key).append(value)
    else: dictionary.update({key: [value]})

def fix_html(clinkerjson, clinkerhtml):
    print("Fixing html...")
    newhtml = "%s_annot.html" %(clinkerhtml[:-5])
    name_html(clinkerjson, clinkerhtml, newhtml)

def detect_mibig_cluster(bgclist):
    mibig=False
    for el in bgclist:
        if el[0].islower():
            if not mibig:
                mibig = [el]
            else:
                mibig.append(el)
    return mibig

'''from https://stackoverflow.com/questions/2130016/splitting-a-list-into-n-parts-of-approximately-equal-length'''
def split(a, n):
    k, m = divmod(len(a), n)
    return (a[i*k+min(i, m):(i+1)*k+min(i+1, m)] for i in range(n))

def flatten_comprehension(matrix):
    return [item for row in matrix for item in row]

def detect_elements_withsubstring(thelist, thesubstring, index=-1):
    elements = []
    for el in thelist:
        if thesubstring == el[:index]:
            elements.append(el)
    return elements

'''makes sure that elements with the same prefix are not in different lists'''
def tidy_replicates(bgcmetalist):
    newbgcmetalist=[]
    allbgcslist = flatten_comprehension(bgcmetalist)
    prefixes = [x[:-1] for x in allbgcslist]
    ### keeping only prefixes that will cause problems
    relevantprefixes = [x for x in prefixes if prefixes.count(x) > 1]
    for i in range(1,len(bgcmetalist)):
        previouslist = list(bgcmetalist[i-1])
        currentlist = list(bgcmetalist[i])
        ### for each prefix that is an issue
        for prefix in relevantprefixes:
            previouscount = [x[:-1] for x in previouslist].count(prefix)
            currentcount =  [x[:-1] for x in currentlist].count(prefix)
            ### if found in two separate lists
            if previouscount !=0 and currentcount !=0:
                if previouscount > currentcount:
                    tobemoved = detect_elements_withsubstring(currentlist, prefix)
                    previouslist = previouslist + tobemoved
                    currentlist = [x for x in currentlist if x not in tobemoved]
                else:
                    tobemoved = detect_elements_withsubstring(previouslist, prefix)
                    currentlist = currentlist + tobemoved
                    previouslist = [x for x in previouslist if x not in tobemoved]
        ### after resolving all prefixes, save previous list
        newbgcmetalist.append(previouslist)
    ### after iteration is complete, save also current list
    newbgcmetalist.append(currentlist)
    return newbgcmetalist

# Main function #
def main(input, output, metatable, clustermetadata, overwrite=False):
    if not (os.path.exists(metatable) and os.path.isfile(metatable)):
        print("\nGCF assignment table not found. Trying to detect...")
        metatable = detect_gcftable(metatable)

    print("\nInput gbk files: %s\n"
          "Input GCF assignment table: %s\n"
          "Input cluster metadata table: %s\n"
          "Output folder: %s\n"
          "Overwrite: %s\n"
          %(input, metatable, clustermetadata, output, overwrite))
    clustergbkfiles = list_files(input, '.gbk')
    clustermetadata_df = pd.read_csv(clustermetadata)
    clustermetadata_df.set_index('ID', drop=False, inplace=True)
    bgc_to_gcf_dict = {}
    create_folder(output)
    gcf_to_bgc_dict = csv_to_dict(metatable, key='Family Number', value='#BGC Name', sep='\t')
    #gcf_to_bgc_dict['test']=['risA','bal','AmBf2']
    #pprint(gcf_to_bgc_dict)
    singletons = []
    ### iterate over gcfs and create corresponding clinker files
    for gcfid, bgclist in gcf_to_bgc_dict.items():
        for bgc in bgclist:
            bgc_to_gcf_dict[bgc] = gcfid
        if len(bgclist) > 1:
            ismibig = detect_mibig_cluster(bgclist)
            if len(bgclist) > 10:
                q, mod = divmod(len(bgclist), 10)
                if mod > 0:
                    q+=1
                #print("dividing", q, mod)
                #bgcmetalist = split(bgclist, q)
                bgcmetalist = np.array_split(bgclist, q)
                bgcmetalist = tidy_replicates(bgcmetalist)
            else:
                bgcmetalist = [bgclist]
            #print(gcfid)
            #pprint(bgclist)
            #pprint(bgcmetalist)
            count = 1
            for bgclist in bgcmetalist:
                bgclist = list(bgclist)
                if len(bgcmetalist) > 1:
                    thisgcfid = '%s_part%s' %(gcfid, count)
                    count+=1
                else:
                    thisgcfid = gcfid
                if ismibig:
                    print(bgclist, ismibig)
                    bgclist = list(set(bgclist + ismibig))
                    prefile = os.path.join(output, "gcf_%s_%s" % (thisgcfid, ismibig[0]))
                else:
                    prefile = os.path.join(output, "gcf_%s" % (thisgcfid))
                clusterfiles = [detect_element(clustergbkfiles, bgc) for bgc in bgclist]
                if not os.path.exists("%s.html" %(prefile)) or overwrite:
                    execute_clinker(clusterfiles, prefile)
                    fix_html("%s.json" %(prefile), "%s.html" %(prefile))
        else:
            print("GCF %s has only one member. Skipping..." %(gcfid))
            singletons = singletons + bgclist
    ### run also for singletons:
    gcfid = 'singles'
    if len(singletons) > 10:
        q, mod = divmod(len(singletons), 10)
        if mod > 0:
            q += 1
        bgcmetalist = np.array_split(singletons, q)
        bgcmetalist = tidy_replicates(bgcmetalist)
    else:
        bgcmetalist = singletons
    count = 1
    for bgclist in bgcmetalist:
        bgclist = list(bgclist)
        if len(bgcmetalist) > 1:
            thisgcfid = '%s_part%s' % (gcfid, count)
            count += 1
        else:
            thisgcfid = gcfid
        prefile = os.path.join(output, "gcf_%s" % (thisgcfid))
        clusterfiles = [detect_element(clustergbkfiles, bgc) for bgc in bgclist]
        if not os.path.exists("%s.html" % (prefile)) or overwrite:
            execute_clinker(clusterfiles, prefile)
        if not os.path.exists("%s_annot.html" % (prefile)) or overwrite:
            fix_html("%s.json" % (prefile), "%s.html" % (prefile))
    ### update also metadata table
    clustermetadata_df['GCF_T0.2'] = clustermetadata_df['ID'].map(bgc_to_gcf_dict)
    print(clustermetadata_df)
    newmetafile = os.path.join(output, os.path.basename(clustermetadata))
    clustermetadata_df.to_csv(newmetafile, index=False)

    ### check here for clusters without GCFs in the table and for clusters that are not in the clinker results
    ### (use method independent to clinker)
    #clustergbkfiles

    ### also what would fit is a function that compares previous to current GCFs (in the tables)

    print("\nDone")

#'''
# Parsing arguments & Execution #
parser = argparse.ArgumentParser(description='')
parser.add_argument('-i', '--input', help='Input directory of gbk files', required=True)
parser.add_argument('-o', '--output', help='Output directory of clinker results', required=True)
parser.add_argument('-m', '--metadata', help='Input bigscape clustering table (csv) with GCF assignment. Or bigscape'
                                             ' result folder (assuming only one analysis was run, will autodetect the'
                                             'file).',
                    required=True)
parser.add_argument('-cm', '--clustermetadata', help='Input table of cluster metadata. A column for GCFs'
                                                     ' will be added and the new table will be saved in the output '
                                                     'directory.',
                    required=True)
parser.add_argument('-r', '--overwrite', help='Overwrite results (default=False)', required=False, default=False)
if __name__ == "__main__":
    args = parser.parse_args()
    main(args.input, args.output, args.metadata, args.clustermetadata, args.overwrite )
#'''

'''
### for debugging
inputdir='/media/hdd_2tb/GPA_reclassification_clean/A_dataset_creation/2_putative_GPAs_analyses/5_all_candidate_as7_clusters'
outputdir='/media/hdd_2tb/GPA_reclassification_clean/A_dataset_creation/3_GPA_candidate_dereplication/2_clinker_gcf_results'
metadata='/media/hdd_2tb/GPA_reclassification_clean/A_dataset_creation/3_GPA_candidate_dereplication/1_candidateBGC_bigscape_results'
clustermetadata='/media/hdd_2tb/GPA_reclassification_clean/A_dataset_creation/2_putative_GPAs_analyses/summary_hits_clusters_metadata.csv'
main(inputdir,outputdir,metadata,clustermetadata)
'''

'''
### specific combo of tables
old_table = '/media/hdd_2tb/GPA_reclassification_clean/A_dataset_creation/3_GPA_candidate_dereplication/2_clinker_gcf_results_231023/summary_hits_clusters_metadata_trimmed.csv'
new_table = '/media/hdd_2tb/GPA_reclassification_clean/A_dataset_creation/3_GPA_candidate_dereplication/2_clinker_gcf_results/summary_hits_clusters_metadata.csv'
out_table = '/media/hdd_2tb/GPA_reclassification_clean/A_dataset_creation/3_GPA_candidate_dereplication/2_clinker_gcf_results/summary_hits_clusters_metadata_comparison231023.csv'
clustermetadata_df = pd.read_csv(new_table)
clustermetadata_df.set_index('ID', drop=False, inplace=True)
old_data_df = pd.read_csv(old_table, usecols=['ID','notes','considered_complete','GCF_T0.2'])
old_data_df.set_index('ID', drop=False, inplace=True)
old_data_df.rename(columns={'notes':'old_notes','considered_complete':'old_considered_complete',
                            'GCF_T0.2':'old_GCF_T0.2'}, inplace=True)
combo_df = pd.concat([clustermetadata_df, old_data_df], axis=1)
print(combo_df)
combo_df.to_csv(out_table)
'''
```
