## Supplementary Data 5 for "Structural and evolutionary analyses support reclassification of glycopeptide antibiotics as xyclopeptides": run_full_antismash.py

```
#!/usr/bin/python3.6
# Created by Athina Gavriilidou at 13.01.22 16:39 #
# Imports #
import argparse
import os
from pprint import pprint
from glob import glob
import json

# Functions #
'''given a directory, lists all files, with a certain suffix'''
def list_files(directory, suffix= "", suppress=False):
    the_files = []
    for file in os.listdir(directory):
        if file.endswith(suffix):
            the_files.append(os.path.join(directory, file))
    if not suppress:
        print("Found %s %s files" %(len(the_files), suffix))
    the_files.sort()
    return the_files


'''written on 08.02.22 because i kept needing to do this manually'''
def gunzip_file(filepath):
    command = "gunzip -k %s" %(filepath)
    os.system(command)

'''creates folder in given path, if it does not exist already'''
def create_folder(path):
    if not os.path.exists(path):
        os.mkdir(path)

def execute_full_antismash(fastafile, outputfolder, label):
    antismash_path = "/home/athina/anaconda3/envs/antismash7/bin/antismash"
    print("Using antismash in path: %s" %(antismash_path))
    antismash_command = "%s -c 30 --fullhmmer --clusterhmmer --tigrfam --asf --cc-mibig --cb-general " \
                            "--cb-subclusters --cb-knownclusters --pfam2go --rre --smcog-trees --tfbs " \
                            " --output-dir %s --output-basename %s " \
                            " %s" %(antismash_path, outputfolder, label, fastafile)
    print(antismash_command)
    try:
        os.system(antismash_command)
    except:
        print("ERROR: antismash run failed for %s" %(fastafile))
    #    exit()

'''using dictionary of file suffixes and directories to return list of files'''
def locate_files(hitdirs_to_suffix_dict):
    files_list = []
    ### try to find files with correct suffix in the directories
    for hitdir, suffix in hitdirs_to_suffix_dict.items():
        thisdir_files = [os.path.join(hitdir, o) for o in os.listdir(hitdir)
                         if os.path.isfile(os.path.join(hitdir, o)) and o.endswith(suffix)]
        ### if none are found, search one more level
        if len(thisdir_files) == 0:
            search_path = os.path.join(hitdir, '*/*%s'%(suffix))
            #print(search_path)
            thisdir_files = glob(search_path, recursive=True)
        #print('\n%s:\n%s' %(hitdir, len(thisdir_files)))
        files_list = files_list + thisdir_files
    return files_list

def run_full_antismash(inputfiles, outputdir, overwrite=False):
    #counter = 0
    ### iterate over files and run antiSMASH for each
    for [label, infile] in inputfiles:
        thisoutputfolder = os.path.join(outputdir, label)
        if not overwrite and os.path.exists(thisoutputfolder):
            print("Folder already exists: %s\nSkipping..." %(thisoutputfolder))
            continue
        execute_full_antismash(infile, thisoutputfolder, label)
        ### just for testing
        #counter +=1
        #if counter == 2:
        #    break


def create_folder(path):
    if not os.path.exists(path):
        os.mkdir(path)
        print("Created folder: %s" %(path))

def append_to_file (sourcfile, destinationfile):
    os.system('cat %s >> %s' %(sourcfile, destinationfile))

def list_folders_files(directory, suffix=''):
    folder_file_list = []
    folders_list = [os.path.join(directory, o) for o
                    in os.listdir(directory)
                    if os.path.isdir(os.path.join(directory,o))]
    folders_list.sort()
    singlefolder = False
    ### in case we ever use it for direct folder:
    if len(folders_list) == 0:
        print("\nWarning: no subfolders found. Working with input directory.\n")
        folders_list = [directory]
        singlefolder = True
    for folder in folders_list:
        id = os.path.basename(folder)
        thefiles = list_files(folder, suffix=suffix, suppress=True)
        thefiles.sort()
        #pprint(thefiles)
        for thefile in thefiles:
            if singlefolder:
                id =os.path.basename(thefile[:-len(suffix)].rstrip('.'))
            folder_file_list.append([id, thefile])
    return folder_file_list

def touch_file(filepath):
    os.system('touch %s' %filepath)

# Main function #
def main(input, outputdir,overwriteresults):
    print("\nInput directory: %s\n"
          "Output directory: %s\n"
          "Overwrite results: %s\n"
          %(input, outputdir, overwriteresults))
    create_folder(outputdir)
    bgcids_inputs_list = list_folders_files(input, suffix='.gbk')   ###changed to gbk due to reannotation with originals
    ### if gbk files arent found, continue with original gbffs (for test case)
    if len(bgcids_inputs_list) == 0:
        bgcids_inputs_list = list_folders_files(input, suffix='.gbff')
        print("\nWarning: No gbk files found. Proceeding with gbff files.\n")
        pprint(bgcids_inputs_list)
    run_full_antismash(bgcids_inputs_list, outputdir, overwriteresults)

    print("All concatenated should be generated and run only after all clusters have been computed")
    '''
    ### special all-in-one antismash run
    fullinfile = os.path.join(input, 'all_concatenated.gbff')
    if overwriteresults and os.path.exists(fullinfile):
        touch_file(fullinfile)
    elif not os.path.exists(fullinfile):
        for [bgcid, gbkfile] in bgcids_inputs_list:
            append_to_file(gbkfile, fullinfile)
        execute_full_antismash(fullinfile, os.path.join(outputdir,'all_concatenated'),'all_concatenated')
    '''
    print("\nDone")


# Parsing arguments & Execution #
#'''
parser = argparse.ArgumentParser(description='Script to run full antismash7 (all options on)')
parser.add_argument('-i', '--input', help='Input directory of bakta results folders with files to be analyzed',
                    required=True)
parser.add_argument('-o', '--output', help='Output directory for antismash results', required=True)
parser.add_argument('-r', '--overwrite', help='Optional: overwrite existing results (folders). Default: False',
                    required=False, default=False)

if __name__ == "__main__":
    args = parser.parse_args()
    main(args.input, args.output, args.overwrite)
#'''
```
