## Supplementary Data 5 for "Structural and evolutionary analyses support reclassification of glycopeptide antibiotics as xyclopeptides": run_quick_antismash.py

```
#!/usr/bin/python3.6
# Created by Athina Gavriilidou at 13.01.22 16:39 #
# Imports #
import argparse
import os
from pprint import pprint
from glob import glob
import json

# Functions #
def list_folders(directory):
    folders_list = [os.path.join(directory, o) for o in os.listdir(directory)
                    if os.path.isdir(os.path.join(directory,o))]
    return folders_list

def dict_folders(directory):
    folders_dict = {}
    folders_list = [os.path.join(directory, o) for o
                    in os.listdir(directory)
                    if os.path.isdir(os.path.join(directory,o))]

    folders_list.sort()
    for folder in folders_list:
        filename = os.path.basename(folder)
        id = filename.split("_strain_")[0]
        if id in folders_dict.keys():
            if '_fixed' in filename:
                folders_dict[id] = folder
        else:
            folders_dict[id] = folder
    return folders_dict

'''written on 08.02.22 because i kept needing to do this manually'''
def gunzip_file(filepath):
    command = "gunzip -k %s" %(filepath)
    os.system(command)

'''creates folder in given path, if it does not exist already'''
def create_folder(path):
    if not os.path.exists(path):
        os.mkdir(path)

def execute_quick_antismash(fastafile, outputfolder, label, ismag=False):
    antismash_path = "/home/athina/anaconda3/envs/antismash7/bin/antismash"
    print("Using antismash in path: %s" %(antismash_path))
    if fastafile.endswith('.gbk'):
        genefinding = ''
    elif ismag:
        genefinding = '--genefinding-tool prodigal-m'
    elif not ismag:
        genefinding = '--genefinding-tool prodigal'

    antismash_command = "%s -c 30 --cc-mibig --output-dir %s --output-basename %s %s" \
                            " %s" %(antismash_path, outputfolder, label, genefinding, fastafile)
    print(antismash_command)
    try:
        os.system(antismash_command)
    except:
        print("ERROR: antismash run failed for %s" %(fastafile))
        exit()

'''using dictionary of file suffixes and directories to return list of files'''
def locate_files(hitdirs_to_suffix_dict):
    files_list = []
    ### try to find files with correct suffix in the directories
    for hitdir, suffix in hitdirs_to_suffix_dict.items():
        thisdir_files = [os.path.join(hitdir, o) for o in os.listdir(hitdir)
                         if os.path.isfile(os.path.join(hitdir, o)) and o.endswith(suffix)]
        ### if none are found, search one more level
        if len(thisdir_files) == 0:
            search_path = os.path.join(hitdir, '*/*%s'%(suffix))
            #print(search_path)
            thisdir_files = glob(search_path, recursive=True)
        print(len(thisdir_files), hitdir)
        #print('\n%s:\n%s' %(hitdir, len(thisdir_files)))
        files_list = files_list + thisdir_files
    return files_list

def run_quick_antismash(hitdirs_to_suffix_dict, outputdir, overwrite=False):
    create_folder(outputdir)
    inputfiles = locate_files(hitdirs_to_suffix_dict)
    ### iterate over files and run antiSMASH for each
    for infile in inputfiles:
        label = os.path.basename(infile).split(".")[0]
        thisoutputfolder = os.path.join(outputdir, label)
        if 'Shrikant' in infile or 'Stammsamlung' in infile or 'MGnify' in infile:
            ismag = True
        else:
            ismag = False
        if not overwrite and os.path.exists(thisoutputfolder):
            print("Folder already exists: %s\nSkipping..." %(thisoutputfolder))
            continue
        execute_quick_antismash(infile, thisoutputfolder, label, ismag)

def create_folder(path):
    if not os.path.exists(path):
        os.mkdir(path)
        print("Created folder: %s" %(path))


# Main function #
def main(input, outputdir,overwriteresults):
    print("\nInput directory: %s\n"
          "Output directory: %s\n"
          "Overwrite results: %s\n"
          %(input, outputdir, overwriteresults))
    create_folder(outputdir)
    ### here function to turn inputdict string into a dictionary
    datasets_to_suffix_dict = {"Culp_dataset_downloads": "gbk","Genbank_hits_downloads": "gbff",
                               "Refseq_hits_downloads": "gbff", "Local_Sequencing_hits_exports": "gbk",
                               "MGnify_hits_downloads": "fasta", "MIBiG_hits_downloads": "gbk",
                               "Shrikant_olddata_hits_exports": "fna","Stammsamlung2022_hits_exports": "gbk",
                               "Shrikantold_TellSeq_hits_exports": "fasta"}
    hitdirs = list_folders(input)
    hitdirs_to_suffix_dict = {}
    for dataset, suffix in datasets_to_suffix_dict.items():
        for hitdir in hitdirs:
            if dataset in hitdir:
                hitdirs_to_suffix_dict[hitdir] = suffix
    pprint(hitdirs_to_suffix_dict)
    ### continue with the dict
    run_quick_antismash(hitdirs_to_suffix_dict, outputdir, overwriteresults)

    print("\nDone")


# Parsing arguments & Execution #
'''
parser = argparse.ArgumentParser(description='Script to run quick-and-dirty antismash7')
parser.add_argument('-i', '--input', help='Input download directory of all data (highest folder)', required=True)
parser.add_argument('-o', '--output', help='Output directory for antismash results', required=True)
parser.add_argument('-r', '--overwrite', help='Optional: overwrite existing results (folders). Default: False',
                    required=False, default=False)
if __name__ == "__main__":
    args = parser.parse_args()
    main(args.input, args.output, args.overwrite)
'''

### special case 16.10.2023
inputdir = '/media/hdd_2tb/side_projects/eva_panBGC_pipeline/0_input_data_preparation/0_amycolatopsis_genomes_gbks'
outputdir = '/media/hdd_2tb/side_projects/eva_panBGC_pipeline/0_input_data_preparation/1_minimal_antismash7_amycolatopsis_results'
inputfiles = locate_files({inputdir: 'gbk'})
### iterate over files and run antiSMASH for each
for infile in inputfiles:
    label = os.path.basename(infile).split(".")[0]
    thisoutputfolder = os.path.join(outputdir, label)
    if 'Shrikant' in infile or 'Stammsamlung' in infile or 'MGnify' in infile:
        ismag = True
    else:
        ismag = False
    if os.path.exists(thisoutputfolder):
        print("Folder already exists: %s\nSkipping..." %(thisoutputfolder))
        continue
    execute_quick_antismash(infile, thisoutputfolder, label, ismag)
```
