## Supplementary Data 5 for "Structural and evolutionary analyses support reclassification of glycopeptide antibiotics as xyclopeptides": run_zol.py

```
#!/usr/bin/python3.6
# Created by Athina Gavriilidou at 22.08.23 11:41 #
# Imports #
import argparse
import os

import numpy as np
import pandas as pd
from Bio import SeqIO

# Functions #
def create_folder(path):
    if not os.path.exists(path):
        os.mkdir(path)

def execute_zol(inputdir, outputdir):
    zol_path = "/home/athina/anaconda3/envs/zol_env/bin/zol"
    print("Using zol in path: %s\n" %(zol_path))
    zol_command = ("%s -i %s -o %s -c 30 --full_genbank_labels --comprehensive_evo_stats --width 100 --length 20 " #) \
                   #"--identity_threshold 0 --coverage_threshold 0") \
                    %(zol_path, inputdir, outputdir))
    print(zol_command)
    os.system(zol_command)


'''given a directory, lists all files in it with the given suffix'''
def list_files(directory, suffix):
    files = [os.path.join(directory, o) for o in os.listdir(directory)
               if os.path.isfile(os.path.join(directory, o)) and o.endswith(suffix)]
    print("Found %s %s files" % (len(files), suffix))
    files.sort()
    return files

def find_file_in_list(listoffiles, filename):
    for filepath in listoffiles:
        thename = os.path.basename(filepath)
        if filename in thename:
            return filepath

def get_locus_info(gbkfile, locustag):
    genekind = ''
    genename = ''
    #print('get locus info ',gbkfile)
    input_handle = open(gbkfile, "r")
    for seq_record in SeqIO.parse(input_handle, "genbank"):
        #print("Dealing with GenBank record %s" % seq_record.id)
        for seq_feature in seq_record.features:
            if seq_feature.type == "CDS" and locustag in seq_feature.qualifiers['locus_tag']:
                #print(seq_feature)
                if 'gene' in seq_feature.qualifiers.keys():
                    genename = seq_feature.qualifiers['gene'][0]
                elif 'original_gene' in seq_feature.qualifiers.keys():
                    genename = seq_feature.qualifiers['original_gene'][0]
                if 'gene_kind' in seq_feature.qualifiers.keys():
                    genekind = seq_feature.qualifiers['gene_kind'][0]
                break
    return genename, genekind

# Function to count the number of substrings separated by commas in a string
def count_commasubstrings(cell):
    if (type(cell) is float and np.isnan(cell)) or pd.isnull(cell) or len(cell) == 0 or not cell:  # If the string is empty, return 0
        return 0
    else:
        return len(cell.split(','))

# Main function #
def main(input, output, overwrite=False, antismashfolder=False):
    print("\nInput folder: %s\n"
          "Output folder: %s\n"
          "Overwrite: %s\n"
          "Reference folder: %s\n"
          %(input, output, overwrite, antismashfolder))
    if not os.path.exists(output) or overwrite:
        #create_folder(output)
        execute_zol(input, output)
    ### Combine info in orthogroups table
    reporttablepath = os.path.join(output, 'Final_Results/Consolidated_Report.tsv')
    ogstablepath = os.path.join(output, 'Determine_Orthogroups/Orthogroups.tsv')
    newogstablepath = "%s_custom.tsv" %(ogstablepath[:-4])
    important_cols = ['MI-BiG Annotation (E-value)', 'KO Annotation (E-value)', 'PGAP Annotation (E-value)']
    if not os.path.exists(newogstablepath) or overwrite:
        report_df = pd.read_csv(reporttablepath, sep='\t', index_col=0)
        annot_coldf = report_df[[important_cols[0], 'Proportion of Total Gene Clusters with OG']]
        #annot_coldf.rename(columns={important_cols[0]: 'Most_fitting_Annotation'}, inplace=True)
        annot_coldf['Most_fitting_Annotation'] = annot_coldf['MI-BiG Annotation (E-value)'].apply(
            lambda x : x.split('|')[5] if type(x) == str else x )
        #print(annot_coldf)
        annot_coldf.Most_fitting_Annotation.fillna(report_df[important_cols[1]], inplace=True)
        #print(annot_coldf)
        annot_coldf.Most_fitting_Annotation.fillna(report_df[important_cols[2]], inplace=True)
        #print(annot_coldf)
        ogs_df = pd.read_csv(ogstablepath, sep='\t', index_col=0)
        newogs_df = pd.concat([annot_coldf, ogs_df], axis=1)
        newogs_df.index.name = ogs_df.index.name
        ### if eligible, add more info
        if antismashfolder:
            ogs_list = newogs_df.index.tolist()
            ogs_dict_gene = {}
            ogs_dict_kind = {}
            mibig_clusters = list_files(antismashfolder, '.gbk')
            clusters_of_importance = 'bal,van,ris,a40,tei,kis,rim,isocomp,mis,cor,gp6'.split(',')
            ### iterate over ogs
            for og in ogs_list:
                #print(og)
                thisgene = ''
                thiskind = ''
                ### iterate over important clusters
                for mibigid in clusters_of_importance:
                    ### if the og is not saved yet
                    if og not in ogs_dict_gene.keys():
                        try:
                            thelocustag = newogs_df.loc[og,mibigid]
                        except:
                            continue
                        ### if we have a hit continue gathering info
                        if type(thelocustag) is str:
                            thelocustag = thelocustag.split("|")[-1]
                            mibigfile = find_file_in_list(mibig_clusters, mibigid)
                            if mibigfile is None:
                                print("Could not find gbk of %s" %(mibigid))
                                continue
                            #print('mibigfile',[mibigfile])
                            thisgene, thiskind = get_locus_info(mibigfile, thelocustag)
                            if len(thisgene) > 0:
                                ogs_dict_gene[og]=thisgene
                                ogs_dict_kind[og] =thiskind

                ### if there is no hit in MIBiG, get at least gene kind from any hit
                if og not in ogs_dict_gene.keys():
                    for id in ogs_df.columns:
                        #print(id)
                        thelocustag = newogs_df.loc[og,id]
                        ### if we have a hit continue gathering info
                        if type(thelocustag) is str:
                            thelocustag = thelocustag.split("|")[-1]
                            thefile = find_file_in_list(mibig_clusters, id)
                            thisgene, thiskindhere =get_locus_info(thefile, thelocustag)
                            if len(thiskind) == 0:
                                ogs_dict_gene[og]= thisgene
                                ogs_dict_kind[og] =thiskindhere
                            else:
                                ogs_dict_gene[og] = thisgene
                                ogs_dict_kind[og] = thiskind
                            break
                            #print(newogs_df)
            newogs_df['as_gene'] = newogs_df.index.map(ogs_dict_gene)
            newogs_df['as_gene_kind'] = newogs_df.index.map(ogs_dict_kind)
            cols = newogs_df.columns.tolist()
            cols = cols[:3] + cols[-2:] + cols[3:-2]
            newogs_df = newogs_df[cols]
        newogs_df.to_csv(newogstablepath, sep='\t')
        print(newogs_df)
        print("\nCreated file: %s" %(newogstablepath))
    else:
        print("File already exists: %s" %(newogstablepath))
        newogs_df = pd.read_csv(newogstablepath, sep='\t', index_col=0)
        print(newogs_df)
    ### create also presence absence table
    newogstablepath_counts = newogstablepath[:-4] + '_counts.tsv'
    if not os.path.exists(newogstablepath_counts) or overwrite:
        presence_absence_df = newogs_df.iloc[:,5:]
        annot_cols_df = newogs_df.iloc[:,:5]
        #print(presence_absence_df)
        # Applying the function to each cell in the DataFrame
        counts_df = presence_absence_df.applymap(count_commasubstrings)
        fullcounts_df = pd.concat([annot_cols_df, counts_df], axis=1)
        print(fullcounts_df)
        fullcounts_df.to_csv(newogstablepath_counts, sep='\t')
        print("\nCreated file: %s" % (newogstablepath_counts))
    else:
        print("File already exists: %s" % (newogstablepath_counts))
        fullcounts_df = pd.read_csv(newogstablepath_counts, sep='\t', index_col=0)
        print(fullcounts_df)
    print("\nDone")


# Parsing arguments & Execution #
parser = argparse.ArgumentParser(description='Runs zol with specific parameters for the BGCs in the input folder')
parser.add_argument('-i', '--input', help='Input directory of input BGCs (.gbk)', required=True)
parser.add_argument('-o', '--output', help='Output folder for files generated by zol', required=True)
parser.add_argument('-r', '--overwrite', help='Optional: overwrite existing results (folders). Default: False',
                    required=False, default=False)
parser.add_argument('-as', '--antismash', help='Optional: Folder with antiSMASH generated gbk files of'
                                               ' clusters, to be used as reference for inferring gene names and '
                                               'functions',
                    required=False, default=False)
if __name__ == "__main__":
    args = parser.parse_args()
    main(args.input, args.output, args.overwrite, args.antismash)
```
