## Supplementary figures and images for "Structural and evolutionary analyses support reclassification of glycopeptide antibiotics as xyclopeptides"

### clinker_example_dereplication.png

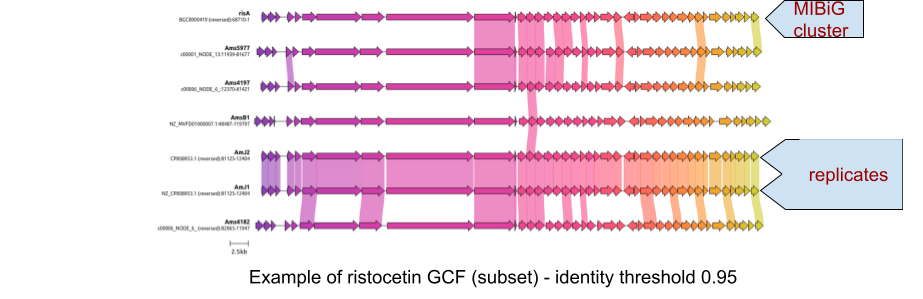

### clinker_example_original.png

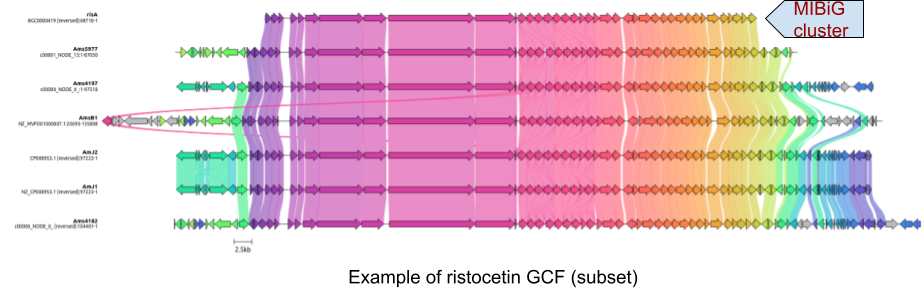

### clinker_example_trimmed.png

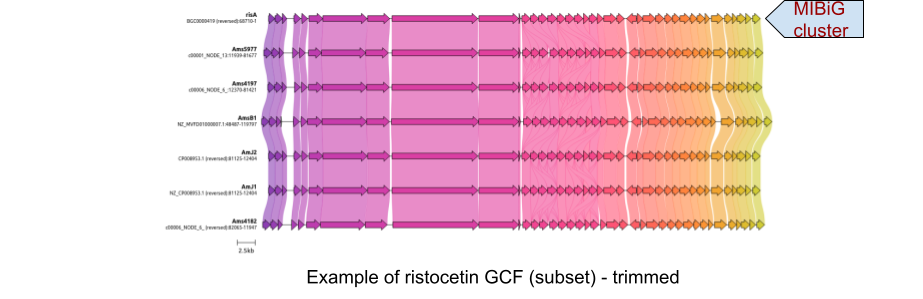

### mibig_backbone_example_logo_a.png

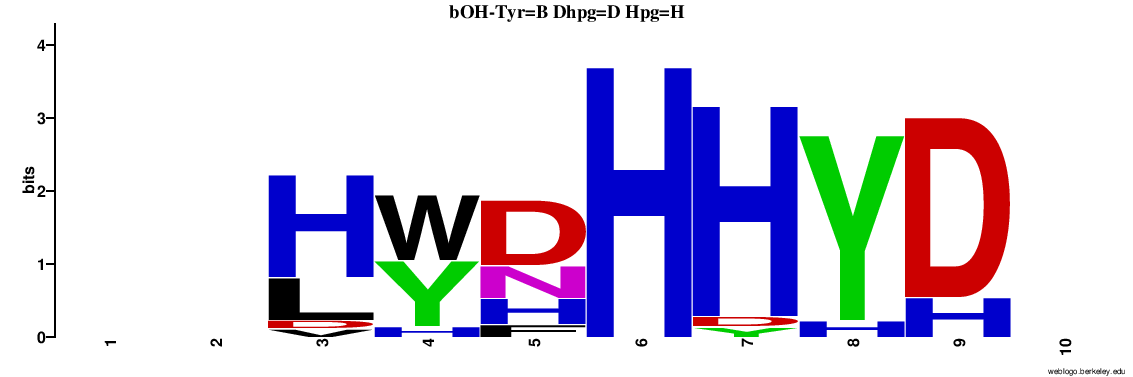

### SFig1_og3_oxyAE_tree.png

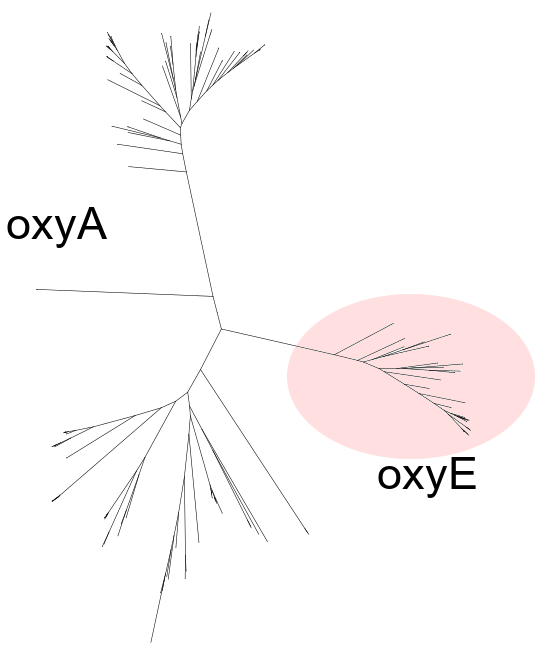

### Supplementary Figure 2

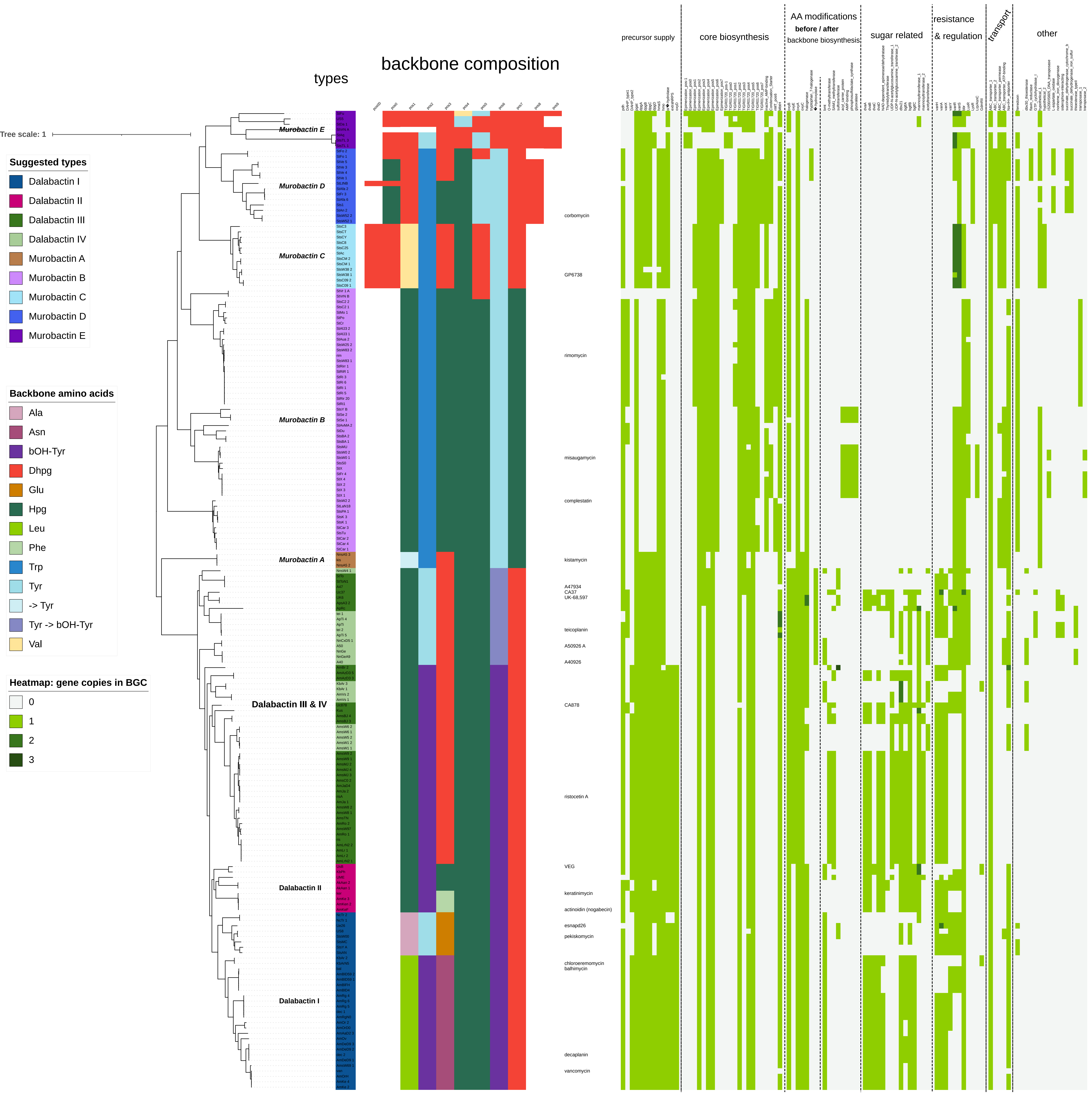

### Supplementary Figure 3

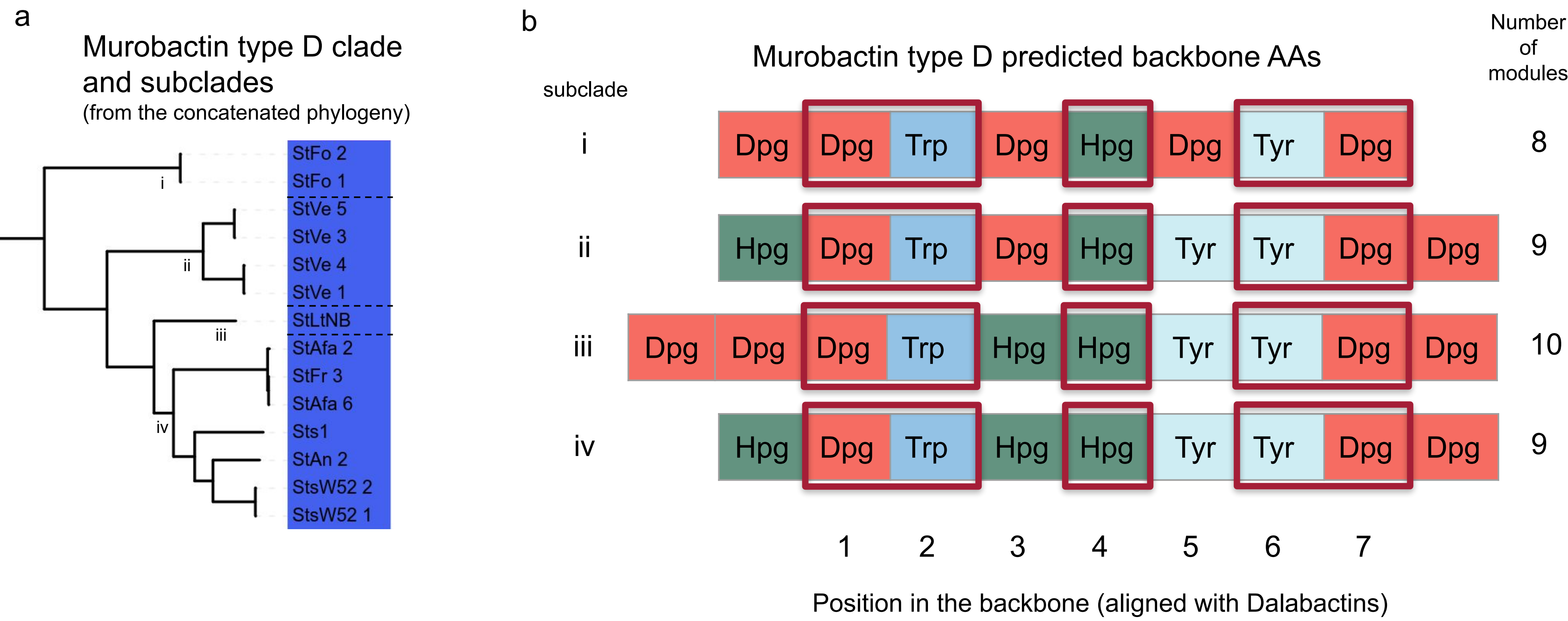

### Supplementary Figure 4

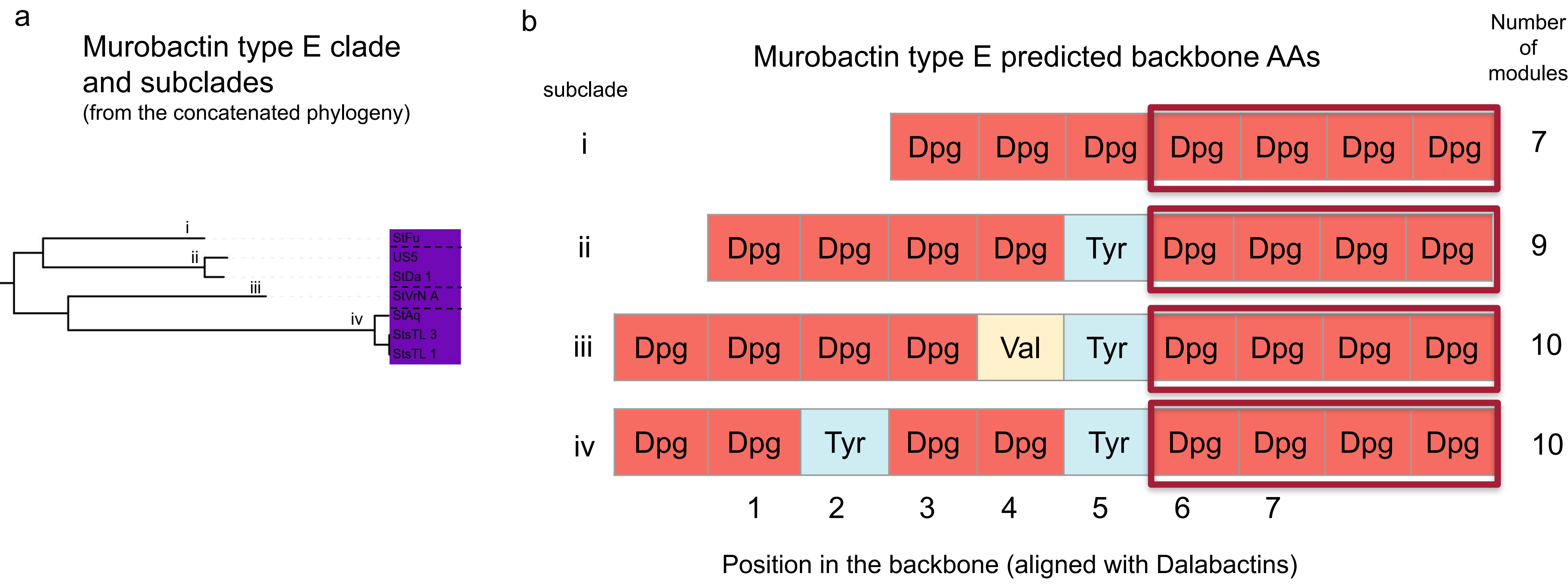

### Supplementary Figure 5

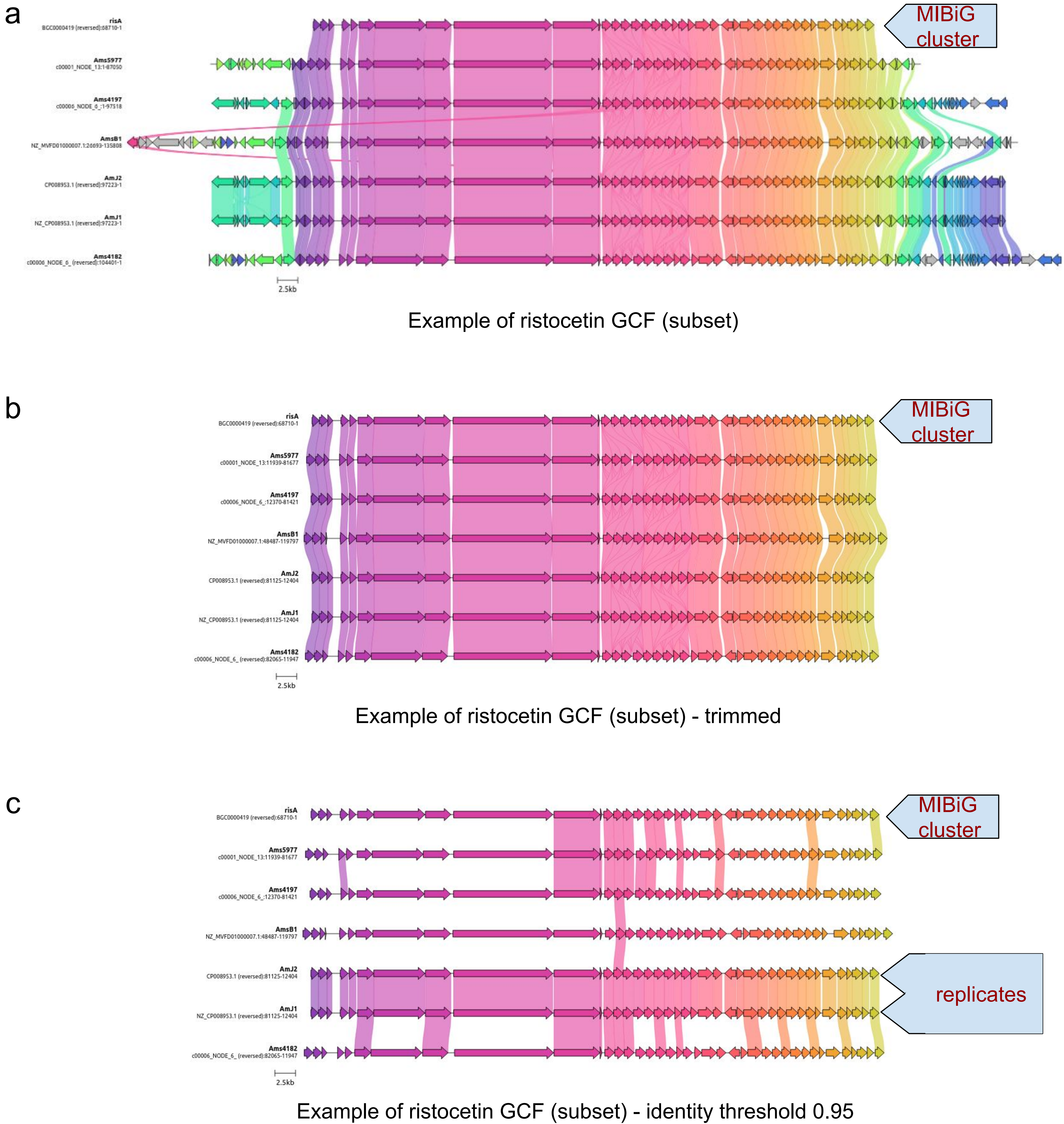
