## Supplementary Data 6 for "Structural and evolutionary analyses support reclassification of glycopeptide antibiotics as xyclopeptides"

### Run Info

| | |
|---|---|
| Host Name | **PC24B184 (localhost)** |
| Position | **1D** |
| Experiment Name | **2203-JR** |
| Sample ID | **pool2** |
| Run ID | **4e5d1aba-c944-4baa-a0fb-49c32540f66f** |
| Acquisition ID(s) | **245060d4f1b337c2d2f3b68b431bb26ab1eb18f9, 9d07cc83e95576775dd9d7d480403d0895b2ca10, dae01ab5a1921d3d05efa187d9c1544b7672e022, c432ab9e4f32ce2433363f22c538a7db066c7ba7** |
| Flow Cell Id | **PAH86725** |
| Start Time | **March 29, 12:04** |
| Run Length | **23h 59m** |

### Run Summary

| | |
|---|---|
| Reads Generated | **375.49 k** |
| Passed Bases | **2.24 Gb** |
| Failed Bases | **689.87 Mb** |
| Estimated Bases | **3.1 Gb** |

### Run Parameters

| | |
|---|---|
| Flow Cell Type | **FLO-PRO002** |
| Kit | **SQK-LSK109** |
| Initial bias voltage | **-165 mV** |
| FAST5 output | **Enabled** |
| FASTQ output | **Enabled** |
| BAM output | **Disabled** |
| Bulk file output | **Disabled** |
| Active channel selection | **Enabled** |
| Basecalling | **Enabled** |
| Specified run length | **72 hours** |
| FAST5 reads per file | **4000** |
| FAST5 output options | **vbz_compress,fastq,raw** |
| FASTQ reads per file | **4000** |
| FASTQ output options | **compress** |
| Mux scan period | **1 hour** |
| Reserved pores | **0 %** |
| Basecall model | **High-accuracy basecalling** |
| Barcoding | **barcoding_kits=["EXP-NBD104"],trim_barcodes="off",require_barcodes_both_ends="off", detect_mid_strand_barcodes="off",min_score=60** |
| Read filtering | **min_qscore=9** |

### Versions

| | |
|---|---|
| MinKNOW | **21.05.13** |
| MinKNOW Core | **4.3.8** |
| Bream | **6.2.5** |
| Guppy | **5.0.12** |

### Cumulative Output Reads

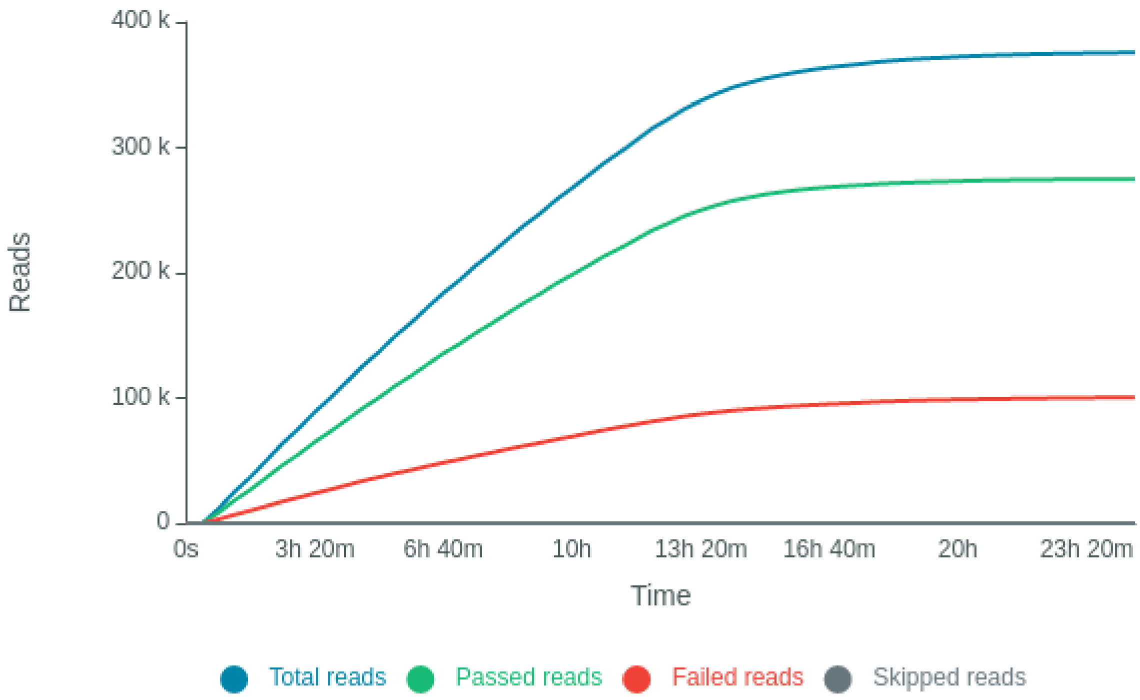


### Cumulative Output Bases

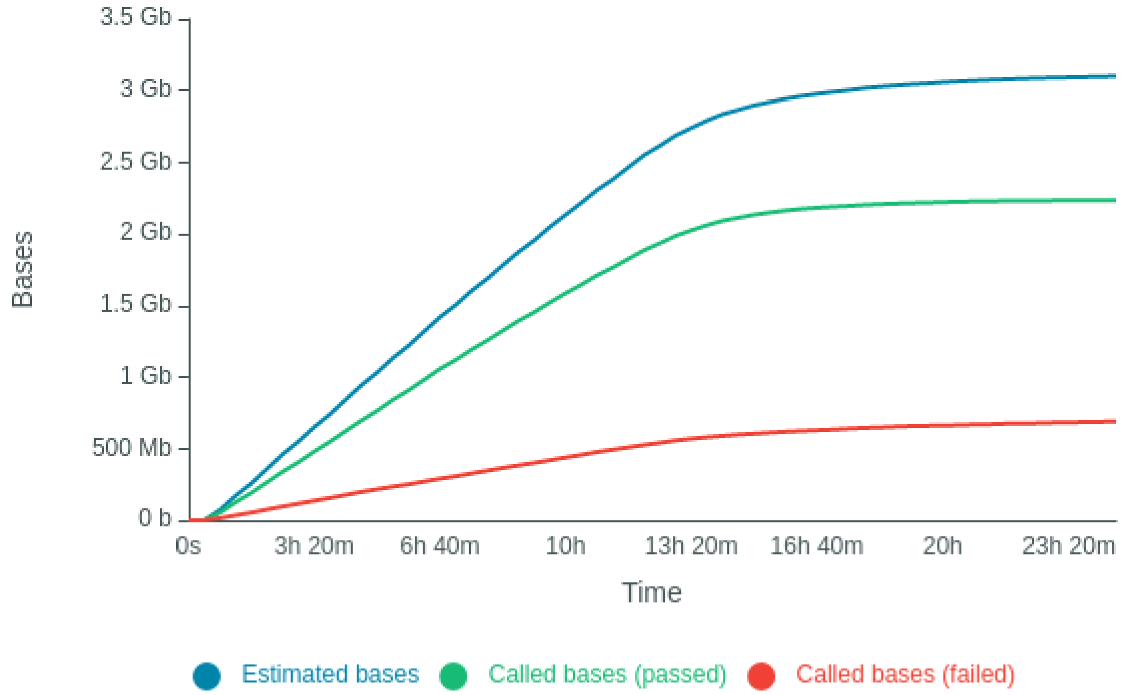

### Read Length Histogram Estimated Bases - Outliers Discarded

Estimated N50: 15.62 kb

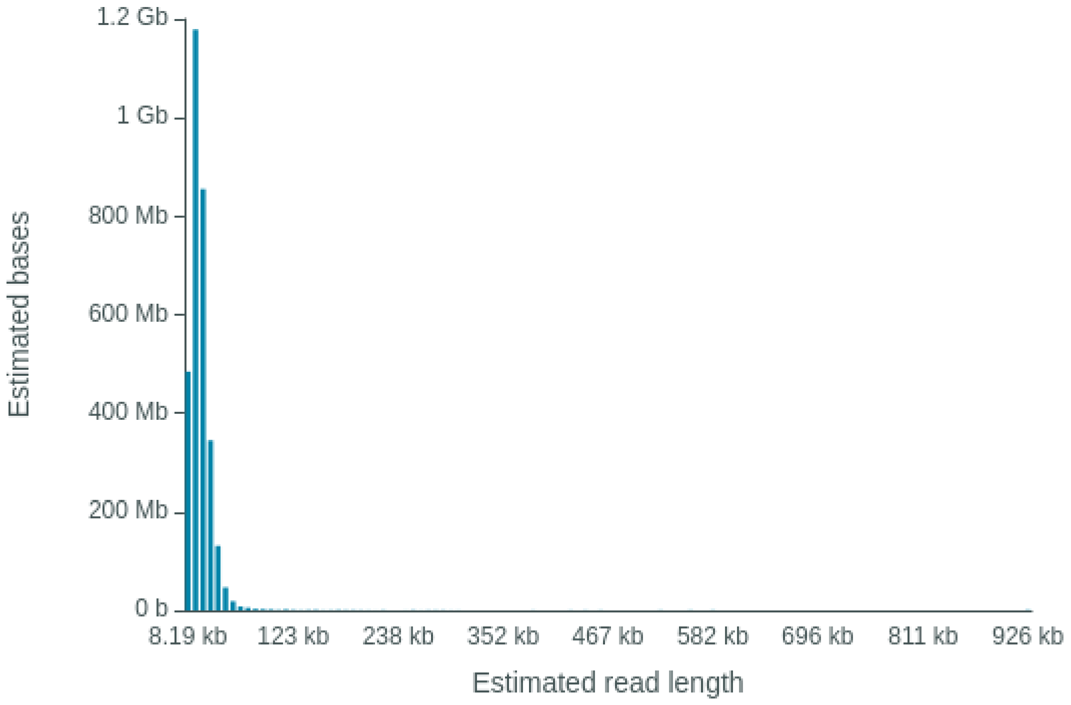


### Read Length Histogram Basecalled Bases - Outliers Discarded

Estimated N50: 14.76 kb

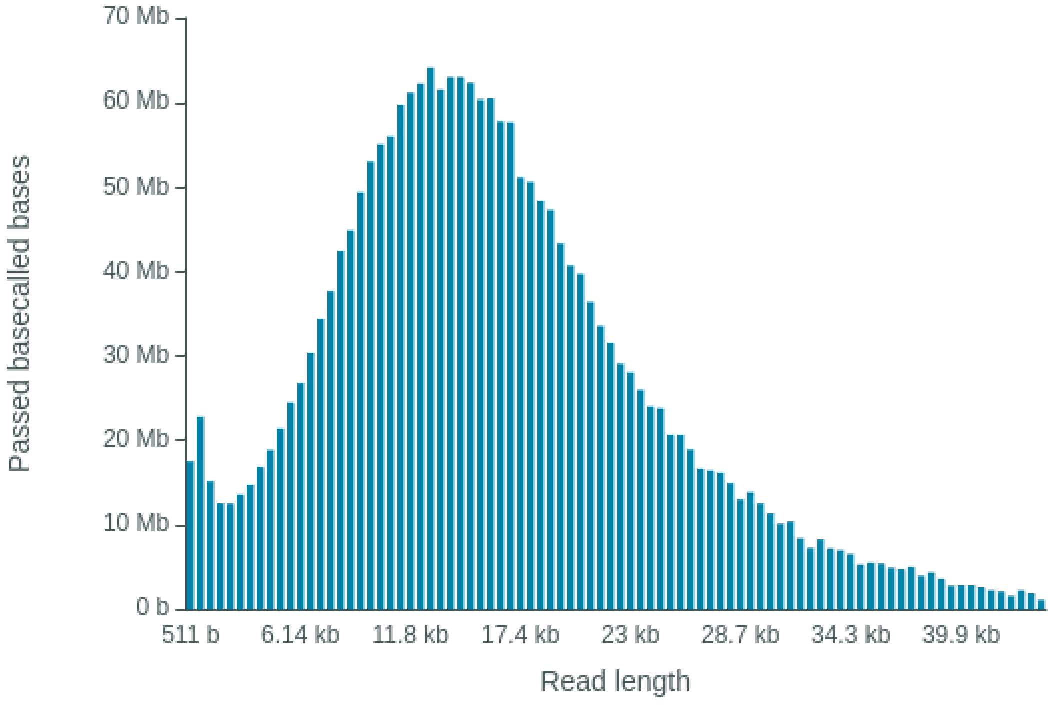

### Read Length Histogram Estimated Bases

Estimated N50: 15.62 kb

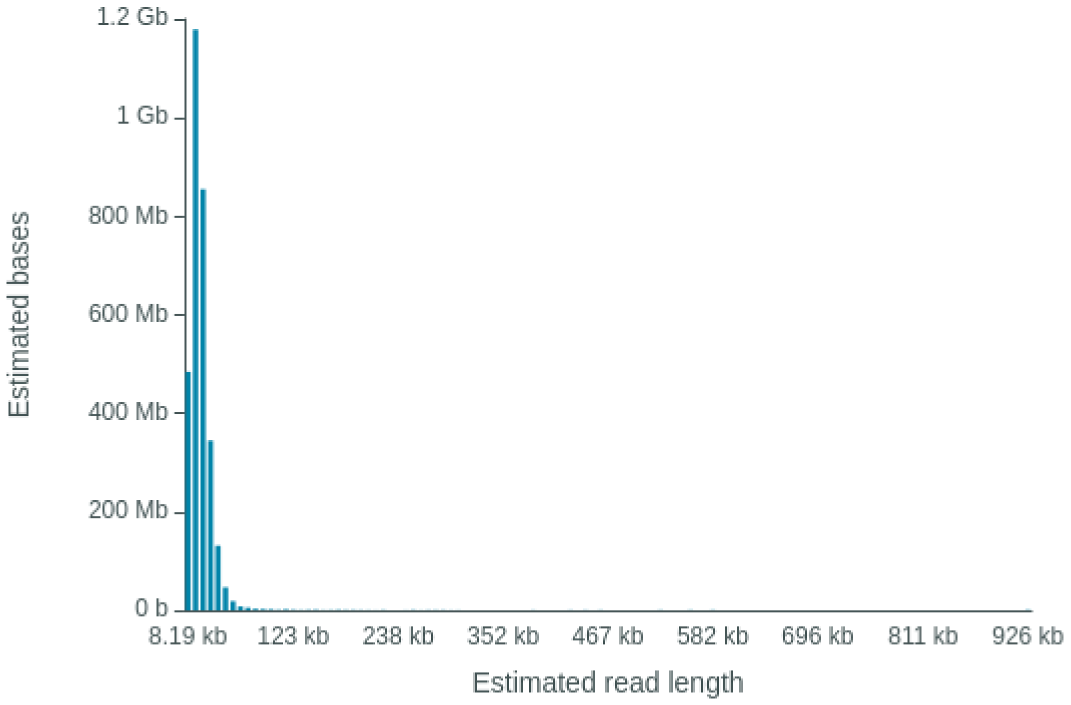


### Read Length Histogram Basecalled Bases

Estimated N50: 14.76 kb

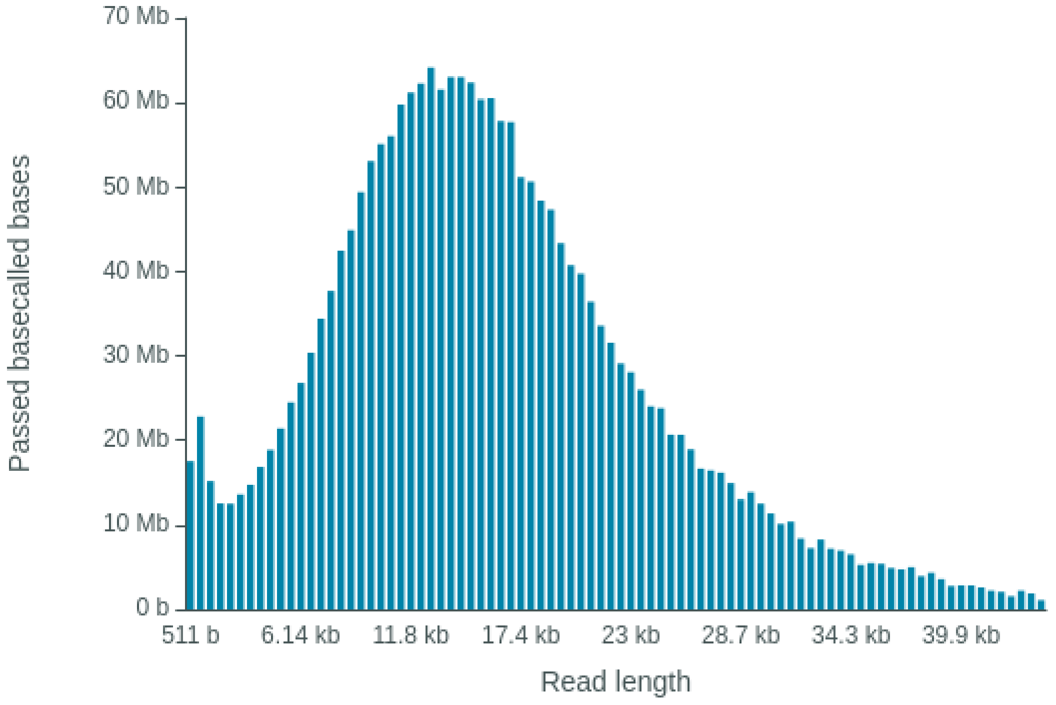

### Barcode Read Counts (reads)

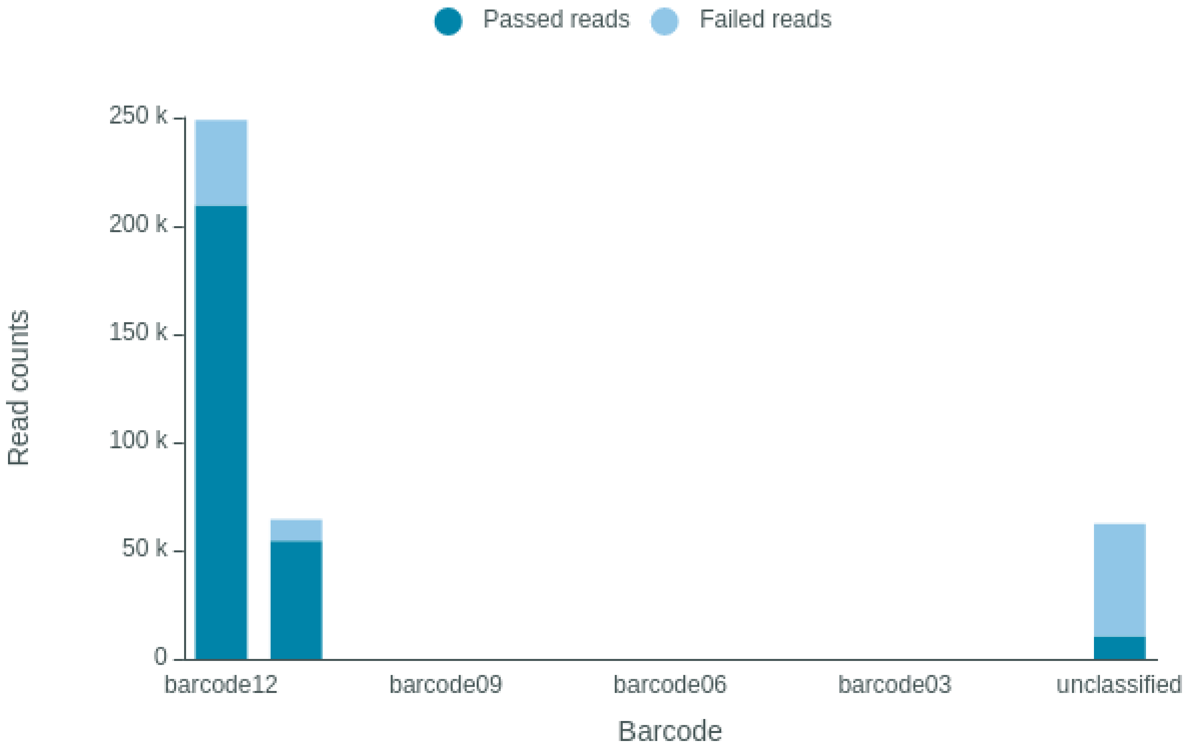


### Barcode Read Counts (bases)

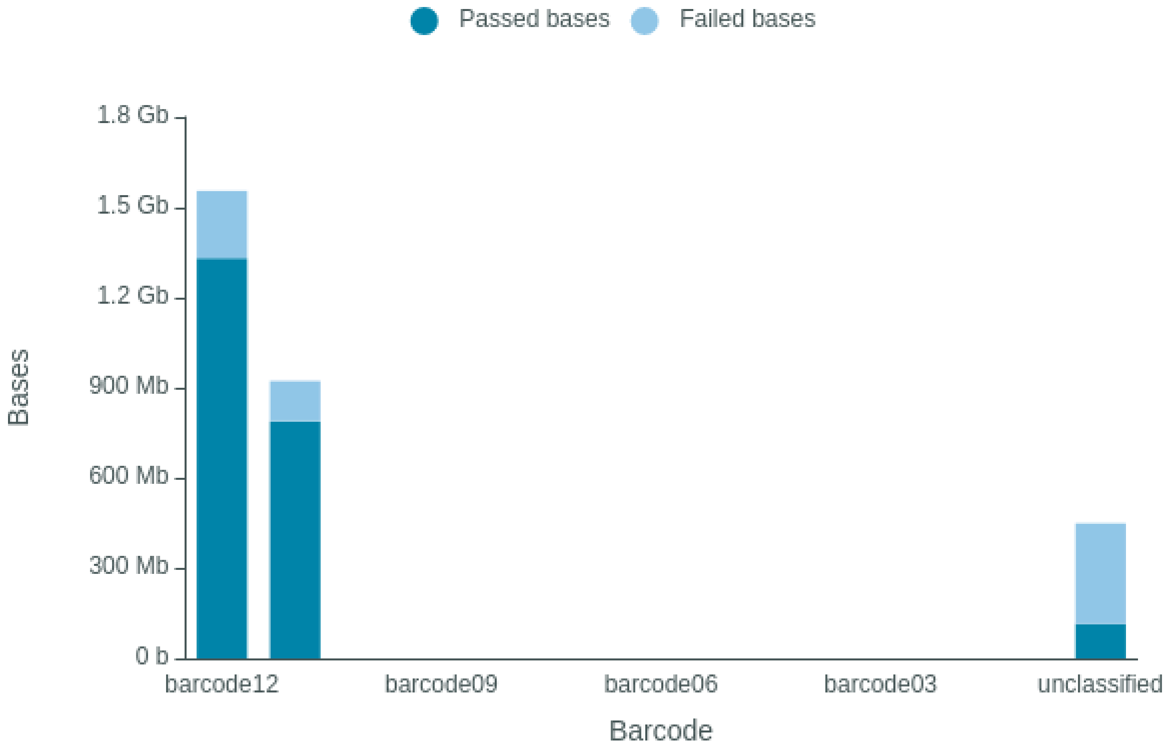

### Duty Time Grouped

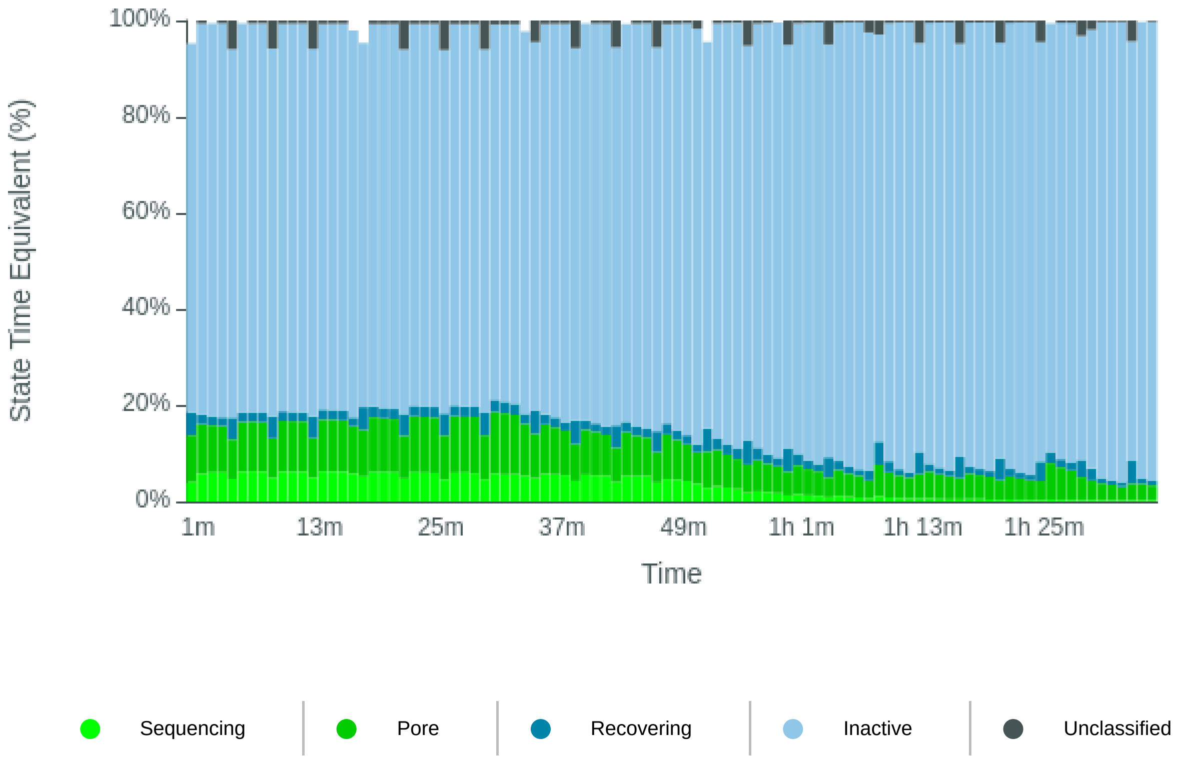


### Duty time Categorised

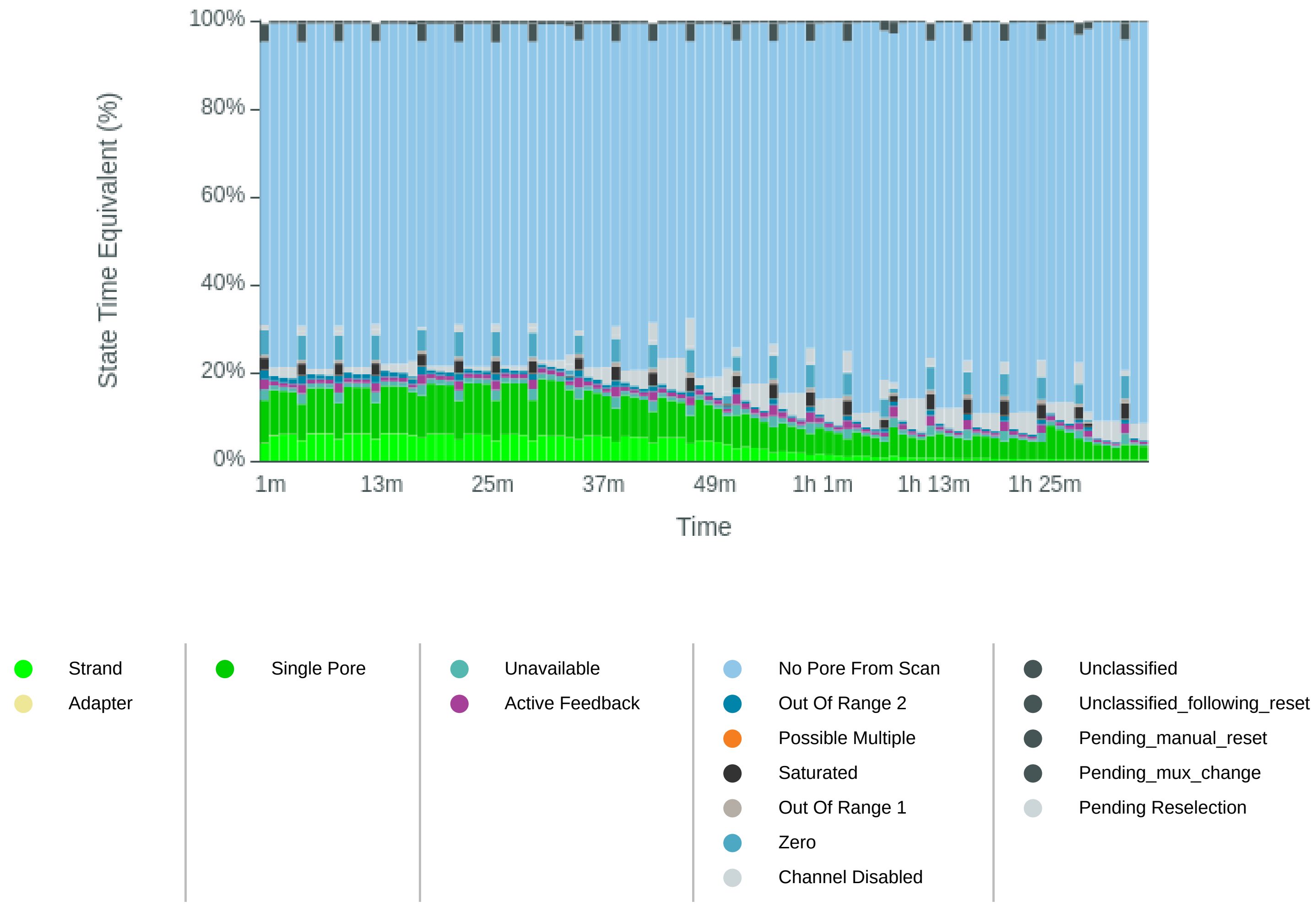

### Mux Scan Grouped

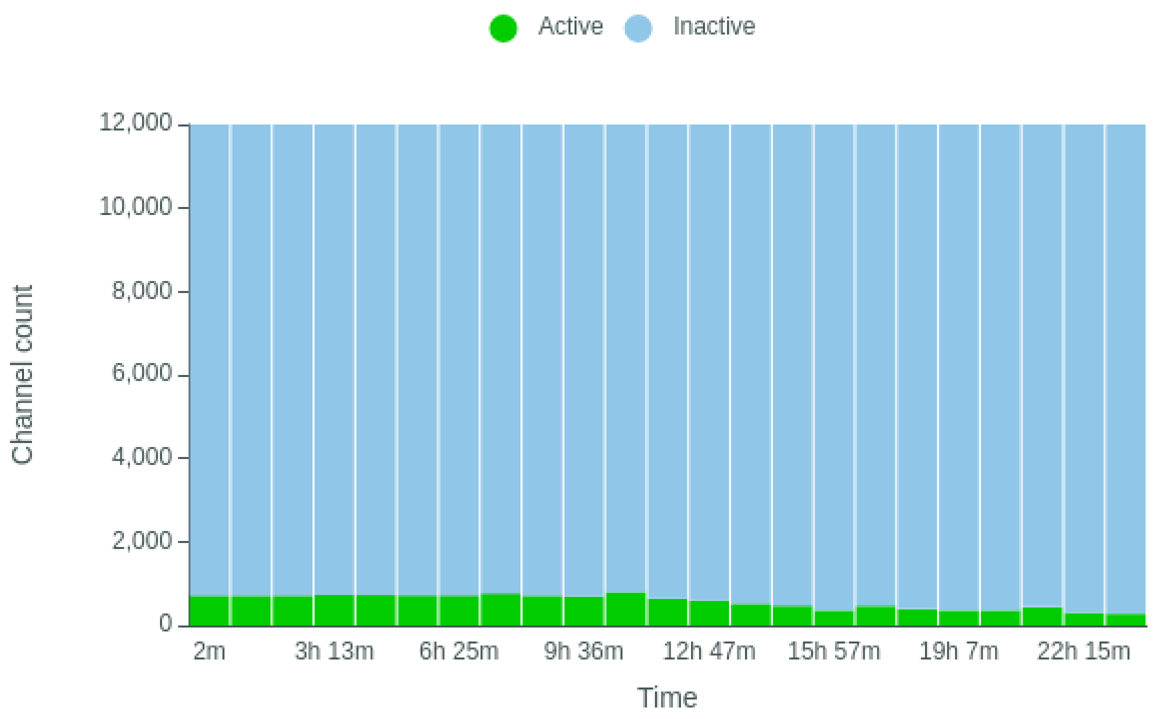


### Mux Scan Categorised

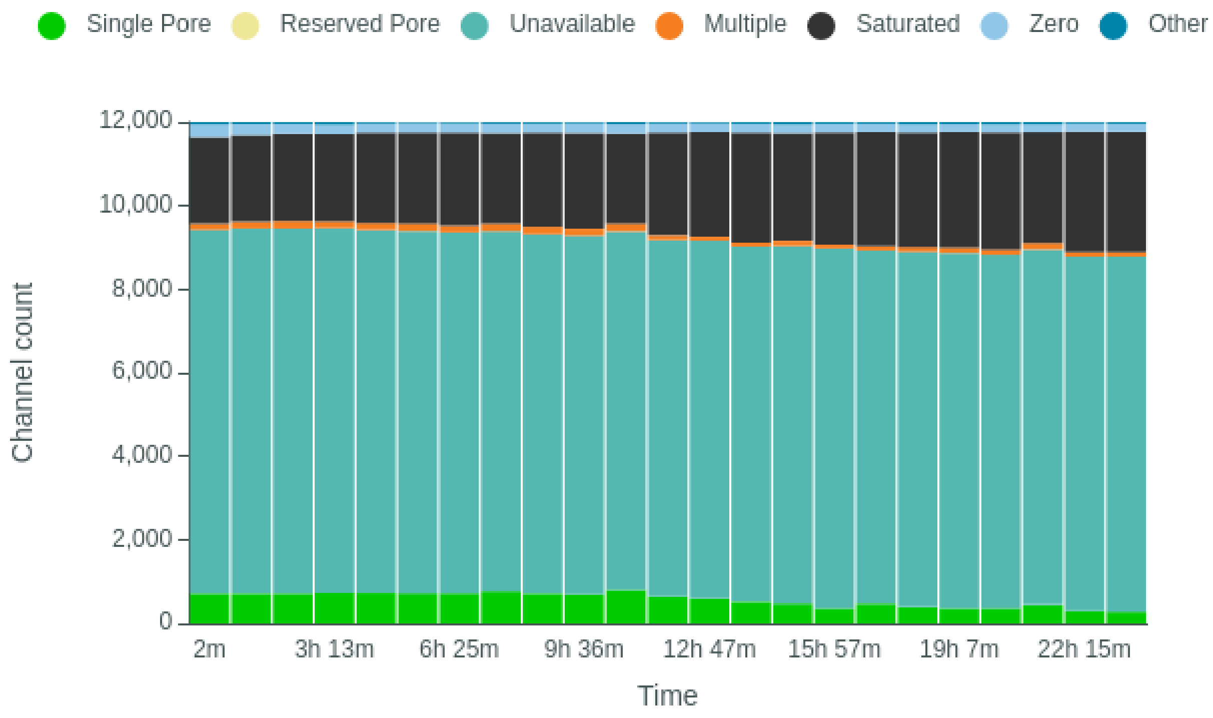

### Temperature History

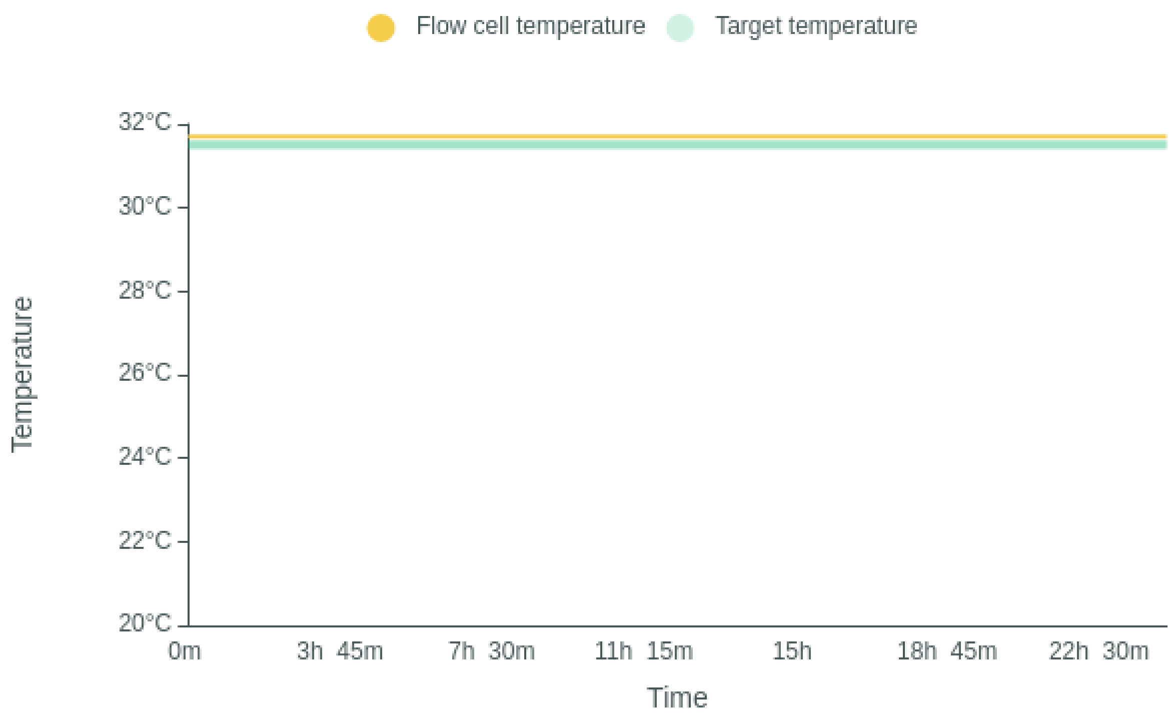


### Bias Voltage History

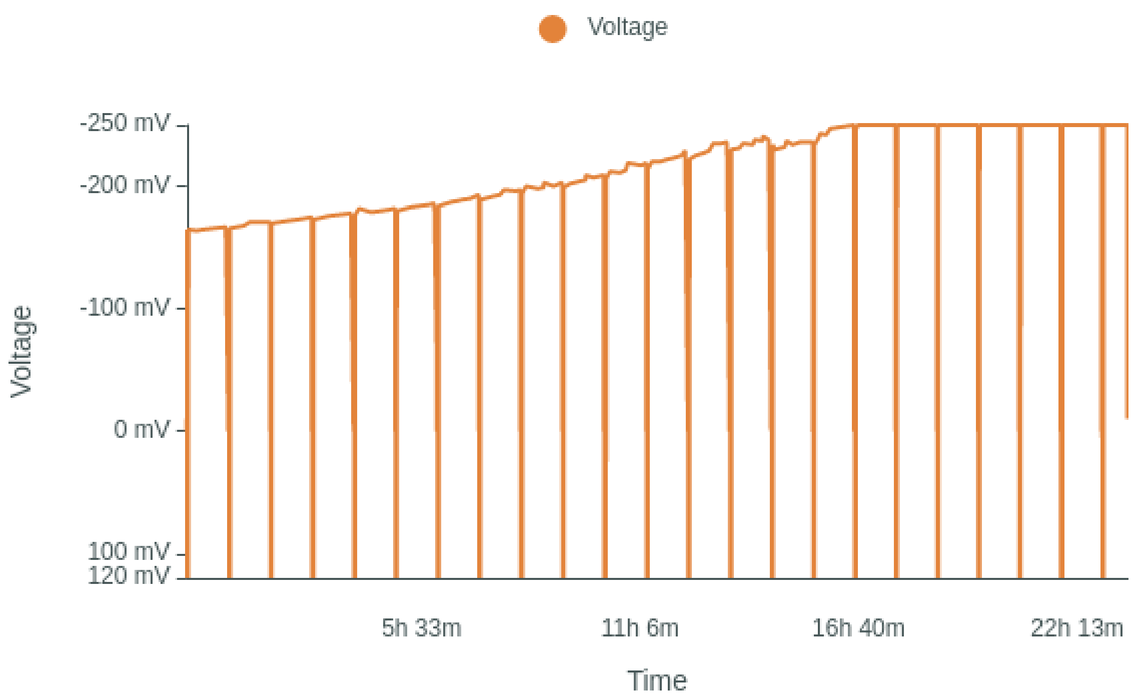

### Translocation Speed

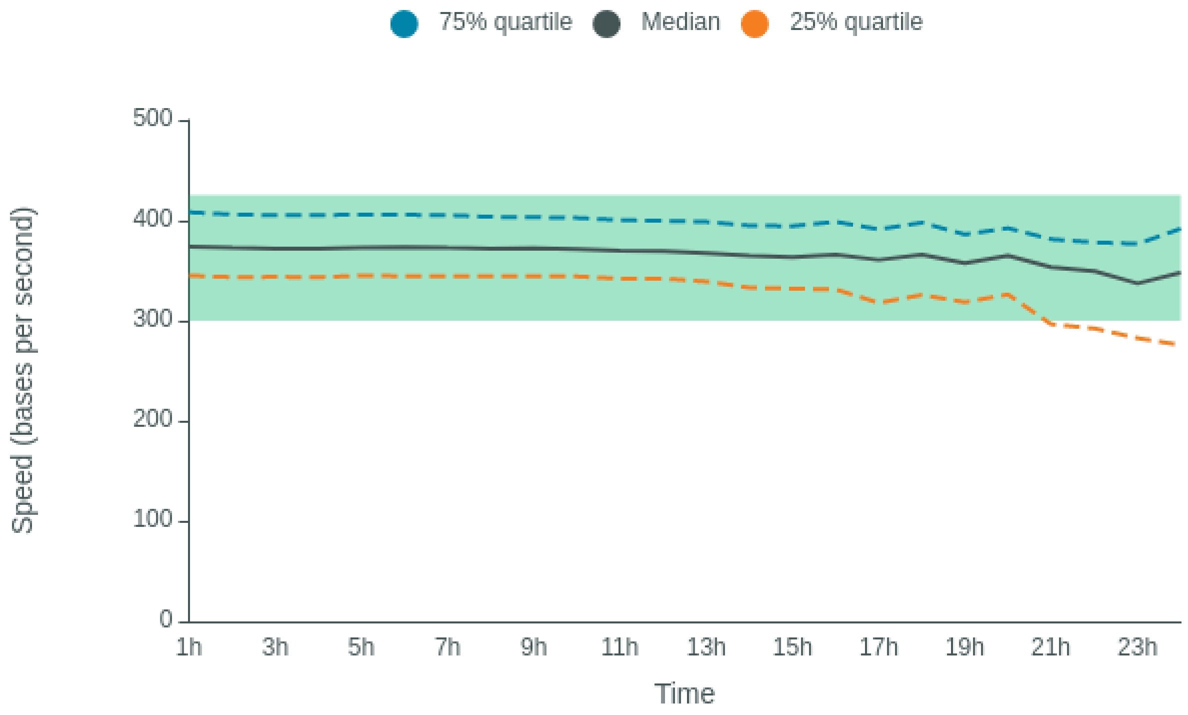


### QScore

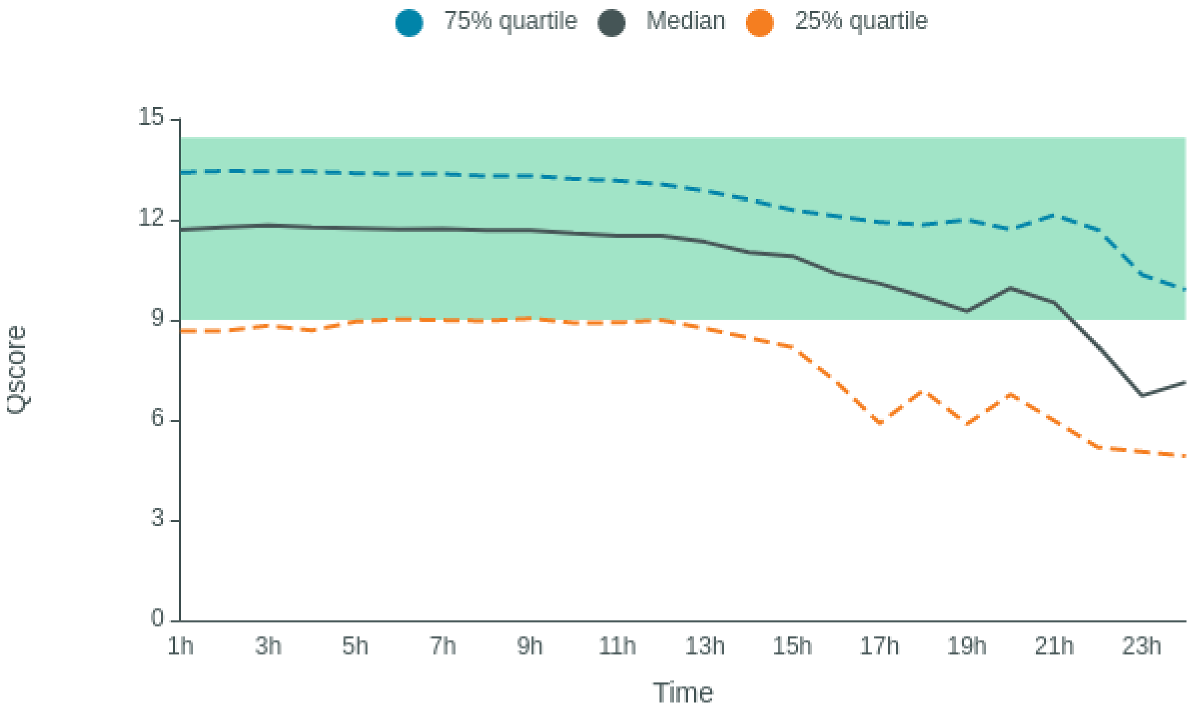

### Disk Write Performance

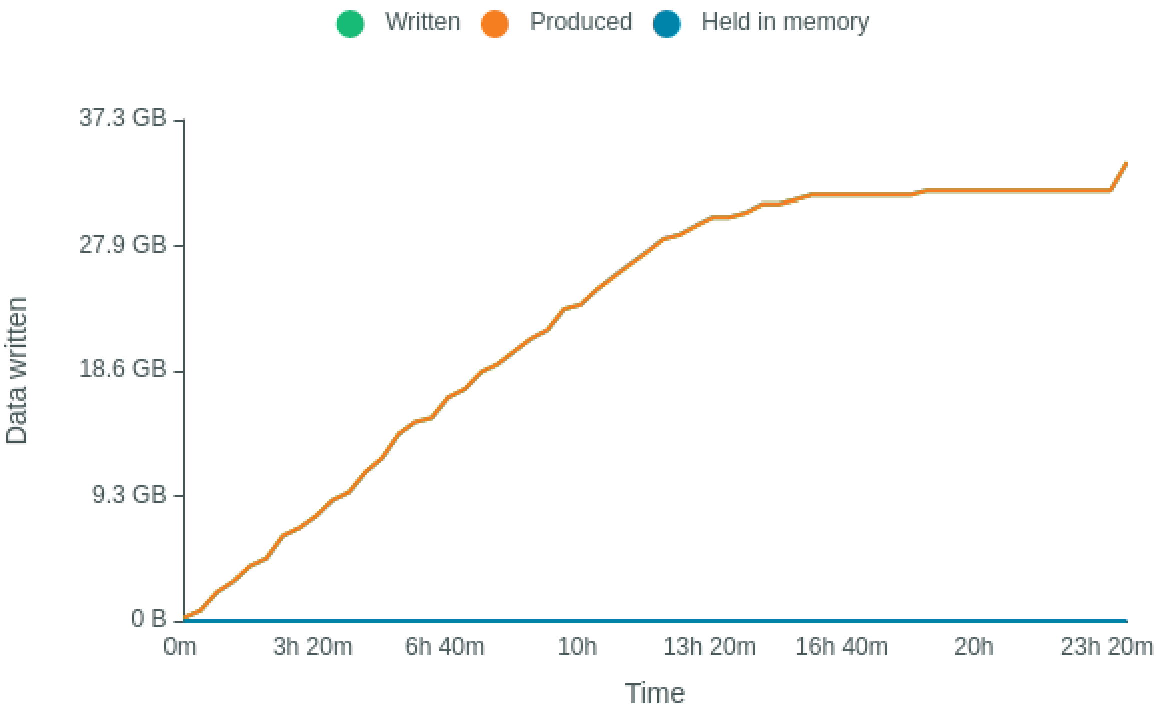

**Run Debug Messages**

- Mux scan for flow cell PAH86725 has found a total of 281 pores. 260 pores available for immediate sequencing March 30, 11:26
- Performing Mux Scan March 30, 11:23
- Mux scan for flow cell PAH86725 has found a total of 304 pores. 278 pores available for immediate sequencing March 30, 10:23
- Performing Mux Scan March 30, 10:20
- Mux scan for flow cell PAH86725 has found a total of 438 pores. 411 pores available for immediate sequencing March 30, 09:20
- Performing Mux Scan March 30, 09:18
- Mux scan for flow cell PAH86725 has found a total of 365 pores. 340 pores available for immediate sequencing March 30, 08:18
- Performing Mux Scan March 30, 08:15
- Mux scan for flow cell PAH86725 has found a total of 363 pores. 335 pores available for immediate sequencing March 30, 07:15
- Performing Mux Scan March 30, 07:12
- Mux scan for flow cell PAH86725 has found a total of 393 pores. 370 pores available for immediate sequencing March 30, 06:12
- Performing Mux Scan March 30, 06:09
- Mux scan for flow cell PAH86725 has found a total of 468 pores. 433 pores available for immediate sequencing March 30, 05:09
- Performing Mux Scan March 30, 05:06
- Mux scan for flow cell PAH86725 has found a total of 364 pores. 338 pores available for immediate sequencing March 30, 04:05
- Performing Mux Scan March 30, 04:03
- Mux scan for flow cell PAH86725 has found a total of 471 pores. 437 pores available for immediate sequencing March 30, 03:02
- Performing Mux Scan March 30, 02:59
- Mux scan for flow cell PAH86725 has found a total of 510 pores. 474 pores available for immediate sequencing March 30, 01:58
- Performing Mux Scan March 30, 01:56
- Mux scan for flow cell PAH86725 has found a total of 587 pores. 540 pores available for immediate sequencing March 30, 00:55
- Performing Mux Scan March 30, 00:52
- Mux scan for flow cell PAH86725 has found a total of 637 pores. 584 pores available for immediate sequencing March 29, 23:51
- Performing Mux Scan March 29, 23:48
- Mux scan for flow cell PAH86725 has found a total of 793 pores. 716 pores available for immediate sequencing March 29, 22:47
- Performing Mux Scan March 29, 22:45
- Mux scan for flow cell PAH86725 has found a total of 686 pores. 633 pores available for immediate sequencing March 29, 21:44
- Performing Mux Scan March 29, 21:41
- Mux scan for flow cell PAH86725 has found a total of 704 pores. 656 pores available for immediate sequencing March 29, 20:40
- Performing Mux Scan March 29, 20:37
- Mux scan for flow cell PAH86725 has found a total of 761 pores. 706 pores available for immediate sequencing March 29, 19:36
- Performing Mux Scan March 29, 19:33
- Mux scan for flow cell PAH86725 has found a total of 710 pores. 666 pores available for immediate sequencing March 29, 18:32

- Performing Mux Scan March 29, 18:30
- Mux scan for flow cell PAH86725 has found a total of 713 pores. 661 pores available for immediate sequencing March 29, 17:29
- Performing Mux Scan March 29, 17:26
- Mux scan for flow cell PAH86725 has found a total of 717 pores. 665 pores available for immediate sequencing March 29, 16:25
- Performing Mux Scan March 29, 16:22
- Mux scan for flow cell PAH86725 has found a total of 724 pores. 678 pores available for immediate sequencing March 29, 15:21
- Performing Mux Scan March 29, 15:19
- Mux scan for flow cell PAH86725 has found a total of 705 pores. 657 pores available for immediate sequencing March 29, 14:18
- Performing Mux Scan March 29, 14:15
- Mux scan for flow cell PAH86725 has found a total of 704 pores. 644 pores available for immediate sequencing March 29, 13:14
- Performing Mux Scan March 29, 13:11
- Mux scan for flow cell PAH86725 has found a total of 705 pores. 654 pores available for immediate sequencing March 29, 12:10
- Performing Mux Scan March 29, 12:07
- Starting sequencing procedure March 29, 12:07
- 180 seconds have elapsed. Experiment commencing. March 29, 12:07
- Waiting up to 180 seconds for temperature to stabilise at 31.5°C March 29, 12:04
- Disk /data has 27061 GB space remaining March 29, 12:04
